# A novel *Candida glabrata* protein regulated by mating signalling pathway shapes inter-species interaction

**DOI:** 10.1101/2024.12.04.621019

**Authors:** Venkata Anand Parnandi, Parth M. Trivedi, Harshita Panchal, Rajesh N. Patkar

**Author notes:** Author for correspondence: Rajesh N. Patkar; email –.

## Abstract

During mixed-species invasive candidiasis - an emerging threat in healthcare - presence of *Candida albicans* is near-essential for host colonisation by *C*. glabrata. However, the molecular communication underlying such intriguing inter-species interaction is largely unexplored. Here, we show that *C. glabrata* secretes a unique small protein Yhi1 that induces hyphal growth in *C. albicans*, which is essential for host tissue invasion. Notably, this Yhi1-based inter-species interaction is specific to *C. glabrata* and *C. albicans*, when compared with other common species of *Candida*.

Our structure-function analyses reveal a novel functional pentapeptide motif (AXVXH) required for the function of Yhi1. Interestingly, Yhi1 expression and efflux are regulated through the mating MAPK signalling pathway and the pheromone transporter *Cg*Ste6, respectively, in *C. glabrata*, despite its preferred asexual reproduction. Our findings shed light on how *C. glabrata* mating signalling pathway is repurposed to interact with *C. albicans*, and highlight the potential of Yhi1 as a biomarker and a template for a synthetic novel antifungal peptide.

---

Among the patients diagnosed with fungal infections globally every year, 1.56 million suffer from invasive candidiasis or *Candida* bloodstream infections, resulting in 995,000 deaths and a crude annual mortality of 63.6 % (Denning, 2024). To date, *Candida albicans* has been identified as the most predominant *Candida* species associated with candidiasis and is a widely used model organism to study fungal diseases and/or host-pathogen interactions (Coco et al., 2008; Kabir et al., 2012; Olsen et al., 2018; Cottier and Hall, 2019). *C. glabrata* has now emerged as the second most prevalent species of candidiasis worldwide and is considered as a high-priority pathogen (Katsipoulaki et al., 2024). Current research into candidiasis largely focuses on the pathogenesis and/or drug resistance of individual species of the fungal pathogen. However, the clinical scenario is more complex, involving multiple species as causative agents. Commonly occurring *Candida* species like *C. albicans*, *C. tropicalis*, *C. parapsilosis* and *C. krusei,* may be isolated as sole species from clinical biopsies, but isolation of *C. glabrata* alone is rare (Redding et al., 2002; Olsen et al., 2018). Recent clinical studies highlight the frequent co-occurrence of *C. glabrata*, along with *C. albicans*, from infection sites (Olsen et al., 2018; Hato et al., 2022). The frequent and potent co-infection by *C. glabrata* and *C. albicans*, along with their direct/physical contact demonstrated during mixed-species oropharyngeal candidiasis study, strongly suggests that crosstalk occurs between these species during pathogenesis (Tati et al., 2016). However, molecular mechanisms underlying the crosstalk between the species, especially during mixed-species invasive candidiasis, remain elusive.

Morphological plasticity is a key virulence trait for many fungal pathogens. *C. albicans* is a polymorphic fungus that can switch its morphology between yeast, pseudohyphae, and hyphal form (Kadosh, 2017; Noble et al., 2017). Various factors, including growth conditions, neutral pH, high temperature (≥37 °C), nutrient starvation, hypoxia, CO_2_, adherence, and molecules of different origins such as serum, N-acetyl glucosamine can trigger hyphal growth in *C. albicans* (Xu et al., 2008; Arkowitz and Bassilana, 2019). On the other hand, *C. glabrata* lacks the ability to switch morphologically from yeast to hyphal form; but is known to rapidly acquire resistance to first-line azole antifungal drugs (Borst et al., 2005; Barber et al., 2019; Ksiezopolska et al., 2021). Together, these two species can pose a serious threat, mainly due to their unique yet complementary abilities.

We hypothesized that the co-existence of the two *Candida* species within the host environment involves beneficial molecular communication that mutually enhances pathogenesis. Here, we show that *C. glabrata* can induce a key morphological change (hyphal growth) in *C. albicans*. We describe a novel small protein in *C. glabrata* that induces the hyphal morphology in *C. albicans*. This unique small protein, with a novel functional pentapeptide motif, is expressed under the MAP kinase (MAPK) mating signalling pathway in *C. glabrata*. We propose that, despite its preference for asexual reproduction, *C. glabrata* might have evolved to utilize its mating signalling pathway to express this protein, facilitating interaction with *C. albicans*. Overall, this study sheds light on a highly intriguing and impactful molecular communication that potentially occurs between different microbes for reciprocal benefit.

## Results and Discussion

### *C. glabrata* induces morphological change in *C. albicans*

To investigate whether *C. glabrata*, which does not exhibit morphological changes, can crosstalk with *C. albicans*, we examined the effect of *C. glabrata* cell-free supernatant (CFS) on *C. albicans* at 37 °C in a non-inducing glucose minimal medium (GMM). We observed that, the CFS from an 18 h-old culture of wild type (WT) *C. glabrata* had induced hyphal growth in *C. albicans* within 3 h of treatment, when compared with *C. albicans* grown in GMM alone or with CFS from an 18 h-old culture of WT *C. albicans* (Figure 1a). To determine if this activity was specific to a secreted molecule(s) of *C. glabrata*, we tested the effect of CFS from a 18 h-old culture of WT *Saccharomyces cerevisiae*, and observed no hyphal development in *C. albicans* (Figure 1a). The hyphal growth inducing activity in the *C. glabrata* CFS was significantly evident during the stationary phase i.e., 12 hour post inoculation (hpi) onwards (Figure 1b); whereas, none of the *S. cerevisiae* CFS samples induced hyphal growth in *C. albicans* (Figure 1c). This indicated that the hyphal growth inducing molecule(s) was accumulated and/or secreted specifically during the stationary growth phase of *C. glabrata*.

**Figure 1.**
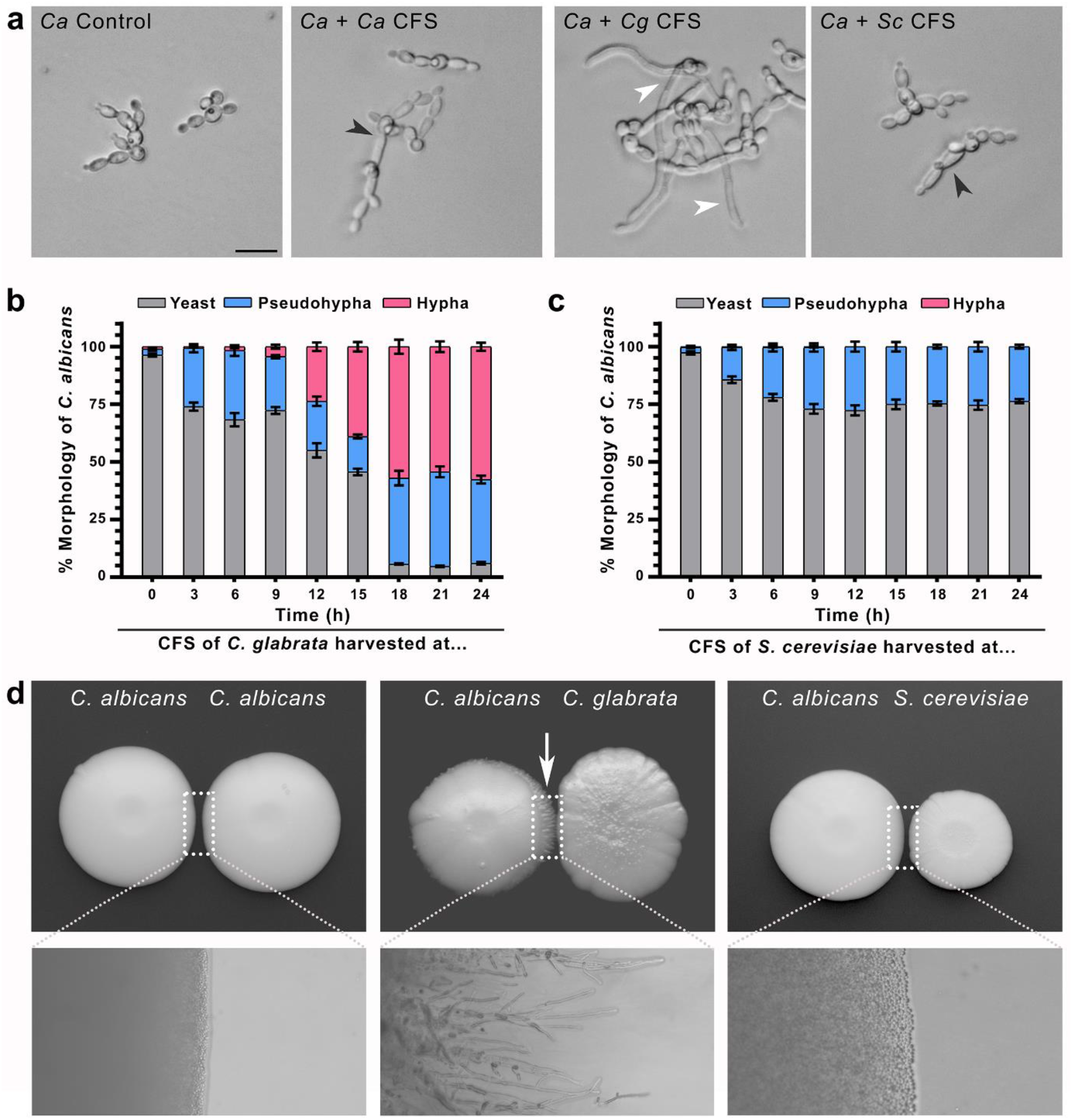
*Candida glabrata* has the ability to induce hyphal growth in *C. albicans*. **(a)** Representative brightfield micrographs showing the response of *C. albicans* to treatment with cell-free supernatant (CFS) from *C. albicans*, *C. glabrata* or *S. cerevisiae*. Black and white arrowheads depict the pseudohyphae and true hyphae, respectively. Scale bar, 10 µm. **(b)** and **(c)** Bar graphs showing quantification of percentage of different morphological forms of *C. albicans*, in response to treatment with *C. glabrata* CFS **(b)** or *S. cerevisiae* CFS **(c)** harvested at specified time intervals. The assay results were observed after 3 h of incubation at 37 °C. Data are presented as the mean ± S.E.M from three independent experiments each. n = 300 each; *P* < 0.05; unpaired *t*-test. **(d)** Co-culturing of *C. albicans* with *C. albicans*, *C. glabrata* or *S. cerevisiae* on YPD agar medium. The white arrow depicts hyphal growth induced in *C. albicans*, particularly at the interface of the two cultures. The lower panels depict micrographs of corresponding colony edges with the area of interest marked with the boxes (dashed lines).

We conducted an assay on YPD agar medium, where *C. albicans* culture was placed next to either *C. albicans*, *C. glabrata,* or *S. cerevisiae*, and incubated for 8-10 days to observe the colony characteristics. The assay showed hyphal growth only at the edge of the *C. albicans* colony facing that of *C. glabrata*, without making any physical contact, in 8-10 days post-inoculation (dpi) (Figure 1d). Notably, *C. albicans* did not form hyphae in response to *S. cerevisiae* or another *C. albicans* colony (Figure 1d).

To corroborate our finding, we used the transwell co-culture assay system to co-incubate the two fungal species in liquid GMM, without any physical contact. Consistent with our earlier results, the transwell co-culture assay showed that *C. glabrata* induced true hyphal growth in *C. albicans* within 3 h of co-incubation (Supplementary Movie 1). Conversely, *C. albicans* predominantly developed yeast cells or pseudohyphae in the presence of *S. cerevisiae* (Supplementary Movie 2).

We then investigated whether *C. glabrata* could induce hyphal growth in other polymorphic non-albicans *Candida* species, such as *C. tropicalis* and *C. dubliniensis*. Our findings showed that the hyphal growth induced by *C. glabrata* was specific to *C. albicans* (Supplementary Figure S1a). Although *C. tropicalis* developed hyphae in response to *C. glabrata*, it exhibited a similar response to *S. cerevisiae* and *C. albicans*, indicating that it was a non-specific response (Supplementary Figure S1a). We also assessed whether the response of *C. albicans* was specific to *C. glabrata*.

Our results show that the hyphal growth in *C. albicans* was induced only in the presence of *C. glabrata*, but not the other species tested here (Supplementary Figure S1b).

## Hyphal growth in *C. albicans* is induced by a mating MAPK signalling pathway-regulated novel small protein *Cg*Yhi1

Next, we investigated if the secretory molecule inducing hyphal growth was a protein or a small molecule of *C. glabrata*. We extracted the CFS of the WT *C. glabrata* culture with water (as earlier) or an organic solvent (acetonitrile or ethanol). We found that the hyphal growth inducing activity was retained only in the aqueous extract, indicating that the molecule of *C. glabrata* was likely a protein rather than a small molecule, based on the assumption that if the active molecule was a protein, its activity would be reduced or lost upon extraction with an organic solvent (Supplementary Figure S2). Altogether, these results show that *C. glabrata* can induce a morphological change in *C. albicans*.

To understand the genetic basis of the hyphal growth inducing activity, we screened an array of *C. glabrata* gene-deletion mutants (Schwarzmüller et al., 2014, Purohit and Gajjar, 2022; Supplementary Figure S3). The array consisted of 848 mutants from a gene-deletion library and 6 independently generated mutants, covering genes involved in key cellular functions such as transcriptional regulation, stress sensing/signalling, antifungal drug resistance, cell wall biogenesis/homeostasis, and a few uncharacterised genes lacking significant orthologues in *S. cerevisiae* (Supplementary Table S1). We identified 65 out of 854 *C. glabrata* mutants, which were impaired in inducing hyphal growth in *C. albicans* (Supplementary Table S2).

Given that physical contact was not required, we hypothesized that *C. glabrata* not only synthesizes but also secretes the inducer protein(s), to trigger hyphal growth in *C. albicans*. Among all the impaired mutants, we found two gene-deletion mutants relevant to the hypothesis – one with a deletion of a gene coding for an uncharacterised small protein (*CAGL0D06666g*) and the other with a deletion of a gene coding for a putative oligopeptide transporter (*CAGL0K00363g*). Therefore, to begin with, we characterized these two genes.

To assess if *C. glabrata CAGL0D06666g* was responsible for inducing hyphal growth in *C. albicans*, we expressed this small protein in *S. cerevisiae* to test whether it turned the non-inducing budding yeast into an inducer of the morphological change.

Interestingly, *S. cerevisiae* strain expressing the *C. glabrata* small protein induced hyphal growth in *C. albicans*, reminiscent of that induced by the WT *C. glabrata* (Figure 2a). This indicated that *CAGL0D06666g* may be associated with the ability of *C. glabrata* to induce hyphal growth in *C. albicans*. To corroborate our finding, we heterologously expressed (in *E. coli*) and purified *C. glabrata* small protein, and found that the purified *C. glabrata* small protein, verified by western blot hybridisation, (Supplementary Figure 4a) induced hyphal growth, with a minimal inducible concentration of ∼300 µM, in *C. albicans* within 3 hpi (Figure 2b; Supplementary Figure S4b). Next, we performed *in silico* analysis of *CAGL0D06666g* to identify the potential homolog of the novel small protein in other related fungal species. To our surprise, we could not find any sequence homolog of *CAGL0D06666g* in any known sequenced genomes across different domains of life. Thus, we identified that *C. glabrata* secretes a novel and likely unique small protein to induce a morphological change, which is a critical virulence determinant, in *C. albicans*. Hereafter, we refer to this *C. glabrata* novel small protein as Yhi1 (**<u>Y</u>**east to **<u>H</u>**ypha **I**nducer **<u>1</u>**).

**Figure 2.**
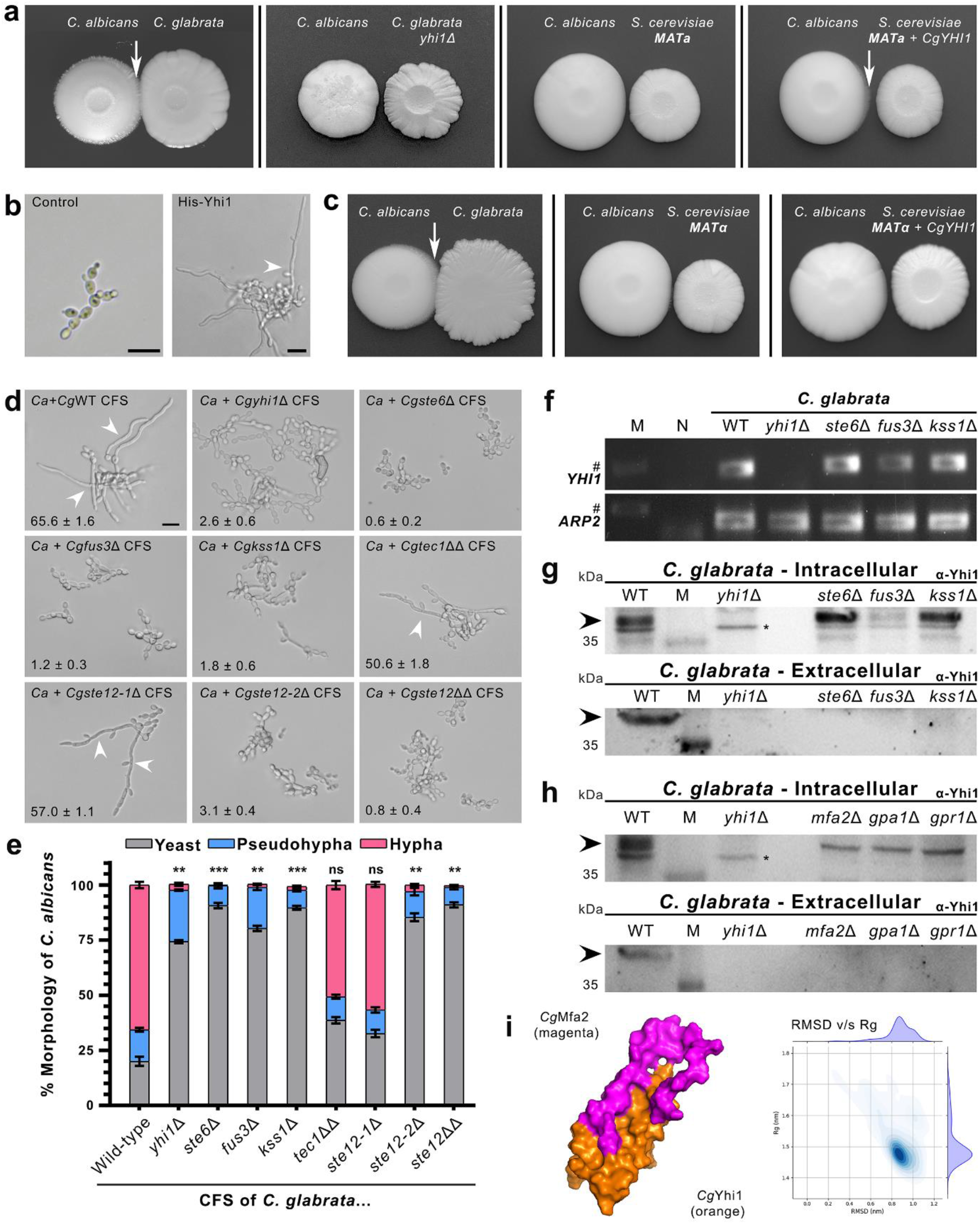
A novel small secretory protein *Cg*Yhi1, expressed and effluxed under the control of mating-related MAPK signalling pathway, induces hyphal growth in *C. albicans*. (a) Co-culturing of *C. albicans* with *C. glabrata* wild type, *C. glabrata yhi1*Δ, *S. cerevisiae MAT*a or *S. cerevisiae MAT*a expressing *CgYHI1* on solid Glucose Minimal Medium (GMM). The white arrows depict hyphal growth induced in *C. albicans*. **(b)** Brightfield micrographs depicting the hyphal growth induced in *C. albicans* by the purified Yhi1. *C. albicans* treated with buffer control or ∼300 µg/ml purified Yhi1. White arrowhead marks true hyphae developed by *C. albicans*. Scale bar, 5 µm. **(c)** Co-culturing of *C. albicans* with *C. glabrata*, *S. cerevisiae MAT*α or *S. cerevisiae MAT*α expressing *CgYHI1* on solid GMM. The white arrow depicts hyphal growth induced in *C. albicans* at the interface of the two cultures. **(d)** and **(e)** Representative brightfield micrographs and corresponding bar graph showing the qualitative and quantitative analyses, respectively, of the response of *C. albicans* to treatment with CFS from the wild type (WT) or specified gene-deletion mutants of *C. glabrata*. White arrowheads depict true hyphae developed by *C. albicans*. Inset values in (d) represent the percentage number of *C. albicans* hyphae observed upon treatment with the specified CFS. The data represents mean ± S.E.M from three independent experiments. n = 300. Scale bar, 5 µm. Statistical analysis was applied relative to *C. glabrata* WT. *P* > 0.05 (ns); *P* < 0.01 (**); *P* < 0.001 (***); two-way ANOVA with Tukey’s multiple comparisons. **(f)** Gene expression (RT-PCR) analysis of *CgYHI1* in the wild type or specified MAPK signalling pathway mutants of *C. glabrata*. *CgARP2* transcript was used as an internal control. M, Molecular size marker; N, No template control. The symbol # on the left marks the position of the 250 bp DNA fragment from the molecular size marker. **(g)** and **(h)** Immunoblot analyses, using α-Yhi1 antibody, showing the intracellular (cell lysate) and extracellular (CFS) levels of *Cg*Yhi1 in the WT and specified MAPK signalling pathway mutants of *C. glabrata*. The black arrowheads mark the positions of hybridisation with the α-Yhi1 antibody. The black asterisks denote the non-specific hybridisation. The numbers on the left mark the position of the proteins (M) of the specified molecular weight (kDa). M, Molecular weight marker. (i) *In silico* molecular docking depicting the interaction between *Cg*Yhi1 and factor **a** pheromone *Cg*Mfa2. The plot depicts the Root Mean Square Deviation (RMSD) versus the Radius of Gyration (Rg) of the *Cg*Yhi1-*Cg*Mfa2 complex.

Our *in silico* analysis of *CAGL0K00363g* showed that the putative oligopeptide transporter had a significant (∼40 %) sequence homology, and conserved domain architecture, with the pheromone transporter Ste6, which is required for the export of factor **a** pheromone in *S. cerevisiae* (Chan et al., 1983; Wilson and Herskowitz, 1984; Kuchler et al., 1989; Supplementary Figure S5). It is known that, in a haploid *S. cerevisiae* strain, specific mating-related genes are expressed depending upon the “mating type” of the strain. The yeast factor **a** mating peptide and its transporter Ste6 are specifically expressed only in mating type **a** (*MAT***a**) haploid cells but not in mating type **α** (*MAT***α**) cells (Wilson and Herskowitz, 1984; Kuchler et al., 1989; Supplementary Figure S6a). Thus, considering the aforementioned sequence homology and characteristics, we asked if the *C. glabrata* putative oligopeptide transporter (*CAGL0K00363g*) was required for efflux of *Cg*Yhi1, in a manner similar to Ste6 in *S. cerevisiae*. To address this, we expressed *Cg*Yhi1 separately in both the mating type strains of *S. cerevisiae*. We earlier showed that expression of *Cg*Yhi1 in *S. cerevisiae*, which was co-incidentally a *MAT***a** strain, induced hyphal growth in *C. albicans* (Figure 2a). Interestingly, the expression of *Cg*Yhi1 in the *MAT***α** strain of *S. cerevisiae* did not induce the characteristic hyphal growth in *C. albicans* (Figure 2c). These results show that *Cg*Yhi1 is effluxed via a Ste6-like oligopeptide transporter in *C. glabrata*.

We noticed that some of the impaired mutants, identified during the *C. glabrata* mutant screening, were of the genes, whose orthologs are involved in the mating or nutrient starvation related mitogen-activated protein kinase (MAPK) signalling pathway in budding yeast (Supplementary Table S2). We wondered if these *C. glabrata* putative MAPK signalling pathway genes (*CAGL0K01507g*, *CgGPR1*; *CAGL0F06677g*, *CgGPA1*; *CAGL0J04290g*, *CgFUS3*; *CAGL0K04169g*, *CgKSS1*; *CAGL0M01716g*, *CgTEC1-1*; *CAGL0F04081g*, *CgTEC1-2*; *CAGL0M01254g*, *CgSTE12-1*; *CAGL0H02145g*, *CgSTE12-2*) were involved in the *Cg*Yhi1-mediated interaction between *C. glabrata* and *C. albicans*. Our *in vitro* CFS-based hyphal induction assay showed that *fus3*Δ, *kss1*Δ, and *ste12-2*Δ mutants of *C. glabrata* were impaired in inducing hyphal growth in *C. albicans*, when compared with the WT. However, *CgTEC1-1*, *CgTEC1-2* and *CgSTE12-1* genes of *C. glabrata* were found to be dispensable for the hyphal induction in *C. albicans* (Figure 2d and 2e).

In *S. cerevisiae*, mating initiates when a pheromone, secreted by a particular mating type, binds to a specific G-Protein Coupled Receptor (GPCR) on the cell membrane of the opposite mating type. Such binding between the pheromone and GPCR leads to the activation of Gpa1 - α subunit of the heterotrimeric G-protein complex - and the downstream MAPK (Fus3) pathway in *S. cerevisiae* (Dietzel and Kurjan, 1987; Miyajima et al., 1987). On the other hand, during nutrient limitation, Gpa2 interacts with another GPCR (Gpr1) to trigger the MAPK (Kss1)-mediated starvation response in *S. cerevisiae* (Iyer et al., 2008). Intriguingly, our gene-deletion library screening showed that *Cg*Gpr1 and *Cg*Gpa1, but not *Cg*Gpa2, were required for the characteristic *Candida* inter-species interaction (Supplementary Table S2; Supplementary Figure S7a and S7b).

Although Fus3 and Kss1 MAPKs have specific roles to play during signalling for mating and nutrient starvation, respectively, in *S. cerevisiae* (Madhani et al., 1997), we did not find any differential roles for *Cg*Fus3 and *Cg*Kss1 homologs of *C. glabrata* with respect to inducing hyphal growth in *C. albicans*. Similarly, although Tec1 is also known to play an important role in pseudohyphal growth in response to nutrient starvation, but not during mating, in *S. cerevisiae*, its *C. glabrata* homologs did not show any role in *Candida* inter-species interaction (Figure 2d and 2e). Importantly, *Cg*Ste12-2, the homolog of Ste12 involved in mating signalling in *S. cerevisiae*, was required for inducing hyphal growth in *C. albicans* (Figure 2d and 2e). Our results show that the mating signalling pathway in *C. glabrata* is majorly required for its interaction with *C. albicans*.

To assess whether the mating signalling pathway regulated the expression and/or efflux of *Cg*Yhi1 during inter-species interaction, we studied the levels of *CgYHI1* transcript in the mutants of the MAPKs and the *Cg*Ste6 transporter. We observed that the transcription of *CgYHI1* was not affected in these mutants (Figure 2f). Given that the MAPK pathway is post-transcriptionally regulated in *S. cerevisiae*, we surmised that expression of *CgYHI1* could also be controlled at the level of translation. Thus, we studied the expression/accumulation of *Cg*Yhi1 protein in the aforementioned mutants, using a polyclonal anti-*Cg*Yhi1 antibody. Western blot analyses of the whole cell lysates (intracellular) showed that *Cg*Yhi1 was expressed in the absence of *Cg*Gpr1, *Cg*Gpa1, *Cg*Kss1 and *Cg*Ste6 function, but was undetectable in the absence of *Cg*Fus3 function (Figure 2g and 2h). This indicated that the expression and intracellular accumulation of *Cg*Yhi1 is post-transcriptionally regulated. This also supports our inference that the mating MAPK (*Cg*Fus3) signalling pathway was involved in *Candida* inter-species interaction. Importantly, *Cg*Yhi1 was not detected extracellularly in the absence of function of all these aforementioned individual proteins (Figure 2g and 2h). The absence of *Cg*Yhi1 in the extracellular milieu shows that the *Cg*Gpr1 and *Cg*Gpa1 mediated activation of *Cg*Kss1 MAPK could be involved in the regulation of *Cg*Ste6 expression required for efflux of *Cg*Yhi1 in *C. glabrata*.

Intriguingly, although *C. glabrata* is generally an asexually-reproducing yeast, it harbours the genes involved in mating and maintains opposite mating types, with each mating type expressing Ste6 as well as both the pheromone GPCR receptors promiscuously, and therefore insensitive to pheromones (Muller et al., 2008; Kumari et al., 2018; Supplementary Figure S6b). Here, we deleted *C. glabrata* pheromone biosynthesis gene *CgMFA*2 (*CAGL0C01919g*), given its significant homology to factor **a** in *S. cerevisiae*, and found that the induction of hyphal growth in *C. albicans*, by the CFS of the *cgmfa2*Δ, was reduced by ∼30 % (Supplementary Figure S7a and S7b). We surmised that *Cg*Yhi1 shared the *Cg*Ste6 transporter with *Cg*Mfa2, and that the reduced hyphal induction could be due to decreased efflux of *Cg*Yhi1 in the absence of *Cg*Mfa2. To confirm whether the efflux of *Cg*Yhi1 was facilitated by *Cg*Mfa2, we studied the intra- and extracellular levels of *Cg*Yhi1 in the *cgmfa2*Δ mutant. We found that, while the intracellular *Cg*Yhi1 level was not affected significantly, it was undetectable extracellularly in the absence of *Cg*Mfa2 function (Figure 2h). This indicated that efflux, but not the expression, of *Cg*Yhi1 was associated with *Cg*Mfa2 in *C. glabrata*. Conversely, we also performed molecular dynamics (MD) simulation study, using the predicted 3D structures of *Cg*Mfa2 and *Cg*Yhi1, to explore any molecular interaction between the two proteins. We found that the two proteins can interact with each other and form a stable complex (Figure 2i). This supports our inference that *Cg*Mfa2 likely aids in the efflux of *Cg*Yhi1 via *Cg*Ste6 in *C. glabrata*. We further observed that the majority of the amino acid residues of *Cg*Mfa2, that are involved in interaction with *Cg*Yhi1, are conserved with those of the mature factor **a** (farnesylated and carboxymethylated factor **a**) of *S. cerevisiae* (Supplementary Figure S8; Chen et al., 1997). Our *in silico* analysis showed that *C. glabrata* does carry orthologs of the genes involved in processing of factor **a** in *S. cerevisiae*. This suggests that *Cg*Mfa2 likely undergoes similar post- translational processing as that observed in case of factor **a** in *S. cerevisiae,* and that the mature *Cg*Mfa2 interacts with *Cg*Yhi1 to form a stable complex (Figure 2i; Supplementary Figure 8). The previous study highlighted that the lipid modification and extreme hydrophobicity resulted in differential migration of the mature factor **a** in *S. cerevisiae* (Chen et al., 1997). Indeed, our western blot analysis results showed hybridisation with anti-Yhi1 antibody at a higher (∼40 kDa) position, when compared with the calculated molecular weight, suggesting that *Cg*Yhi1-*Cg*Mfa2 likely forms a tetrameric complex. This needs to be investigated further using an appropriate analytical tool(s).

## A novel pentapeptide motif is required for *Cg*Yhi1 function

Our *in silico* analysis, using InterProScan and MOTIF Search online web services, to study the structure-function relationship, did not show any known domain or motif in *Cg*Yhi1. However, the BLASTn and tBLASTn analyses showed that some of the *C. glabrata* clinical isolates, particularly from Europe, harboured a variant of *Cg*Yhi1, with a length shorter by two amino acids in the N-terminus (*Cg*Yhi1^3-66^). We found that the *Cg*Yhi1^3-66^ variant induced hyphal growth in *C. albicans*, when compared with the activity of the full-length *Cg*Yhi1 (Figure 3a to 3c).Thus, due to lack of any clue, we expressed the N-terminal (Yhi1^1-32^) and C-terminal (Yhi1^33-66^) halves of the *Cg*Yhi1 protein individually, in the *cgyhi1*Δ mutant (Figure 3a). We surmised that, depending upon the presence of any cryptic domain or motif, expression of either or both halves of *Cg*Yhi1 would lose the protein function. However, to our surprise, we found that the CFS from both the strains, expressing either Yhi1^1-32^ or Yhi1^33-66^, induced hyphal growth in *C. albicans* (Figure 3b, 3c).

**Figure 3.**
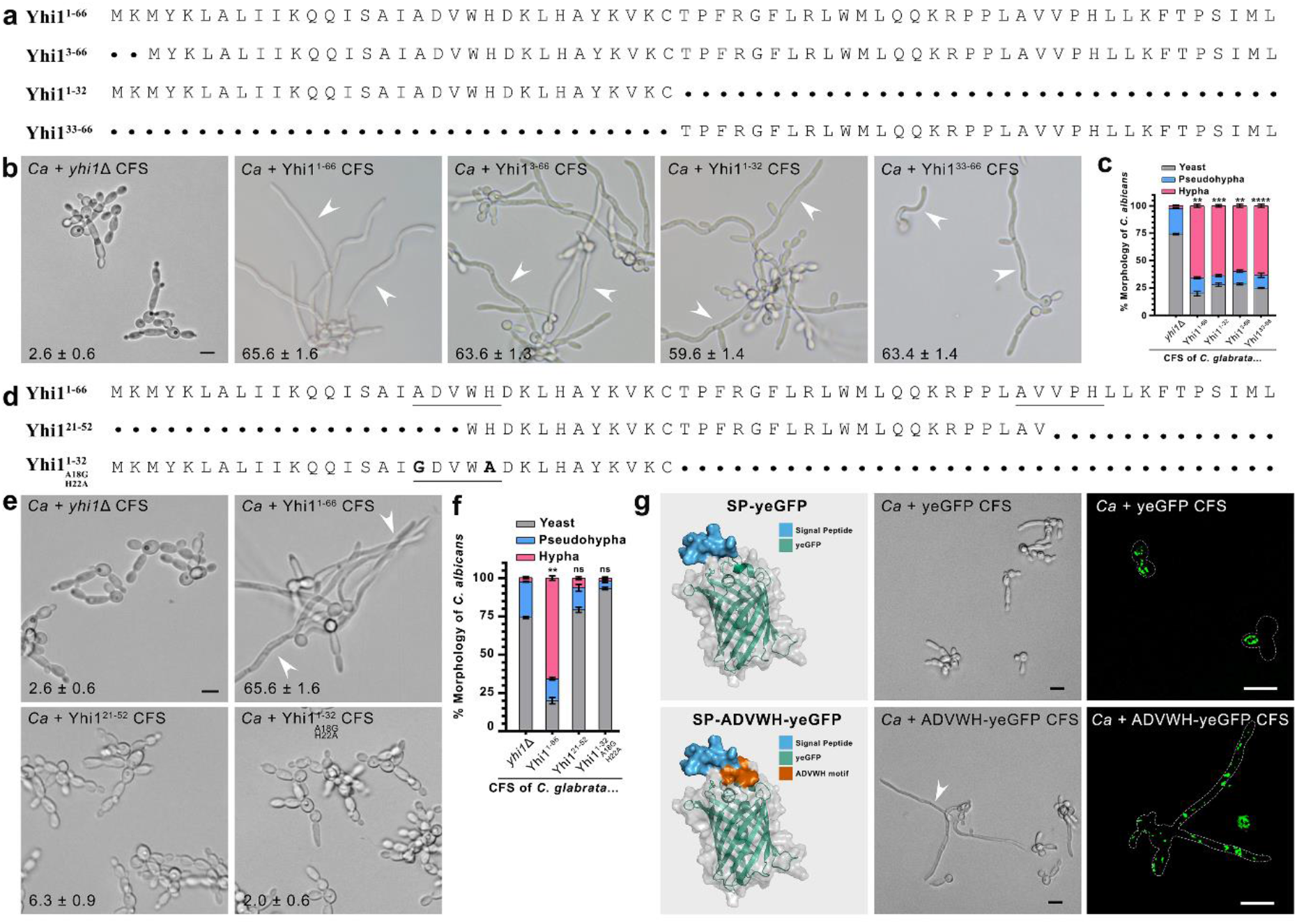
A novel pentapeptide motif in *Cg*Yhi1 is responsible for inducing hyphal growth in *C. albicans.* **(a)** Amino acid sequences of different size variants (full-length or truncated) of *Cg*Yhi1 tested for their ability to induce hyphal growth in *C. albicans*. **(b)** and **(c)** Representative brightfield micrographs and bar graph showing the hyphal growth induced in *C. albicans*, upon treatment with CFS from the *Cgyhi1*Δ *C. glabrata* transformants expressing either of the Yhi1 size variants mentioned in (a). Inset values in (b) represent mean ± S.E.M. of the percentage number of hyphae observed from three independent experiments. n = 300 each. White arrowheads mark the true hyphae developed by *C. albicans*. Scale bar, 5 µm. Statistical analysis was applied relative to *C. glabrata yhi1*Δ. *P* > 0.05 (ns); (*P* < 0.01 (**); two-way ANOVA with Tukey’s multiple comparisons. *P* < 0.01 (**); *P* < 0.001 (***); *P* < 0.0001 (****); two-way ANOVA with Tukey’s multiple comparisons. **(d)** Amino acid sequences of full-length or truncated Yhi1 variants, with or without an intact pentapeptide AXVXH motif, that were tested for their ability to induce hyphal growth in *C. albicans*. **(e)** and **(f)** Representative brightfield micrographs and a bar graph showing the response of *C. albicans* to the CFS from *Cgyhi1*Δ *C. glabrata* transformants expressing either of the Yhi1 size variants mentioned in (c). Inset values in (e) represent the percentage number of *C. albicans* hyphae observed upon treatment with the specified CFS. The data represents mean ± S.E.M from three independent experiments. n = 300 each. Scale bar, 5 µm. Statistical analysis was applied relative to *C. glabrata yhi1*Δ. *P* > 0.05 (ns); *P* < 0.01 (**); two-way ANOVA with Tukey’s multiple comparison. **(g)** The predicted structure of recombinant secretory yeast-enhanced Green Fluorescent Protein (ye-GFP), with or without the ADVWH pentapeptide motif (RoseTTAFold), and the corresponding response of *C. albicans* (brightfield and confocal micrographs) towards CFS harvested from the *cgyhi1*Δ *C. glabrata* strain expressing either of the two ye-GFPs. The predicted structures of both the ye-GFP versions have been slightly modified, without altering the structural characteristics, using the WinCoot software for representation purposes. Note that the brightfield and fluorescence images are from two separate experiments and therefore from different fields. A dashed grey outline marks the hyphal and yeast cells. Scale bar, 10 µm.

We re-examined the protein sequence to determine if both the halves of *Cg*Yhi1 had any feature in common – a likely reason underlying the unaltered hyphal growth inducing activity. We noticed that a putative motif-like pentapeptide sequence, AXVXH, was present in both the halves of *Cg*Yhi1 (underlined; Figure 3d). We performed *in silico* analysis to find if this motif-like sequence (ADVWH or AVVPH) was present in any other known proteins. We found that while ADVWH sequence was present in 19 protein sequences, the AVVPH sequence was present in 846 protein sequences from different organisms (Supplementary Table 3). However, AXVXH pentapeptide sequence has not been hitherto characterised or reported. Thus, to assess whether this pentapeptide sequence was associated with the function of *Cg*Yhi1, we (i) mutated A18G and H22A in Yhi^1-32^ to modify the ADVWH sequence, and (ii) expressed Yhi1^21-52^ – a truncated peptide sequence between the two AXVXH sequences of *Cg*Yhi1 (Figure 3d). Importantly, we observed that the *C. glabrata cgyhi1*Δ mutant expressing either of the two *Cg*Yhi1 variants failed to induce hyphal growth in *C. albicans* (Figure 3e and 3f). Our results strongly indicate that the pentapeptide AXVXH sequence is crucial for *Cg*Yhi1 function. Further studies, using synthetic peptide derivatives (5 to 12 amino acid residues) of *Cg*Yhi1, showed that more than 12 amino acid residues are required for the AXVXH pentapeptide sequence to be functional (Supplementary Figure S9). However, this needs to be verified by assessing the recombinantly-expressed *Cg*Yhi1 peptide derivatives.

In a converse experimental approach, to confirm the functional role of the AXVXH sequence, we incorporated the pentapeptide sequence ADVWH, along with a secretion signal peptide (SP), to the yeast-enhanced Green Fluorescent Protein (yeGFP; Figure 3g), to check if the protein behaved like *Cg*Yhi1. Indeed, the *cgyhi1*Δ mutant expressing the yeGFP containing the pentapeptide ADVWH (SP-ADVWH- yeGFP) induced the characteristic hyphal growth in *C. albicans*, when compared with the yeGFP without the pentapeptide (SP-yeGFP; Figure 3g). The secreted ADVWH-yeGFP localized on the surface of most yeast and hyphal cells, whereas that without the pentapeptide (yeGFP) localized only to a few yeast cells (Figure 3g). Our findings show that the AXVXH sequence is a novel functional pentapeptide motif in the unique *Cg*Yhi1 protein of *C. glabrata* that induces a pathogenesis-related morphological change in *C. albicans*.

## *Cg*Yhi1 has potential clinical applications

We serendipitously observed that one of the aforementioned synthetic peptide derivatives (Yhi1^2-13^) rather demonstrated antifungal activity in a dose-dependent manner. The Yhi1^2-13^ synthetic peptide derivative not only blocked the hyphal growth in *C. albicans*, even in the presence of a strong inducer (5 % serum), but also led to significantly crumpled growth in both *C. albicans* and *C. glabrata*, within 3 hpi (Supplementary Figure S10). The accidental finding here highlights the potential of *Cg*Yhi1 in developing a novel peptide antifungal molecule.

*C. albicans* and *C. glabrata* are well-studied opportunistic human commensals, each equipped with distinct arsenals that aid them in countering a plethora of immune responses. Recent studies have highlighted the increasing incidences of invasive *Candida* infections as alarming and challenging, mainly due to imprecise diagnosis, especially in multimodal invasive candidiasis, without a positive blood culture (Denning, 2024). Here, *CgYHI1* – a novel and unique gene – can serve as a highly precise biomarker for rapidly diagnosing *C. glabrata* in clinical samples (Supplementary Figure S11). Given its inherent resistance to first-line antifungal drugs, identification of *C. glabrata* would enable clinicians to opt for a tailored course of antifungals to effectively treat candidiasis.

Altogether, we unravel that *C. glabrata* utilizes its mating signalling pathway, to express and efflux a novel and unique small protein *Cg*Yhi1, that induces hyphal growth in *C. albicans* (Figure 4). We propose that *C. glabrata*, which otherwise cannot invade host tissue alone due to its inability to undergo filamentation, has evolved with a smarter strategy, to utilize its mating signalling and interact with *C. albicans*, so that it can piggyback on the induced hyphae of the latter, during invasive candidiasis. Basidiomycete fungal pathogens *Cryptococcus neoformans* and *Ustilago maydis* undergo filamentous (hyphal) growth during their sexual life cycle, which is critical for the formation of infectious structures (Kahmann et al., 2000; Bölker, 2001; Hull and Heitman, 2002; McClelland et al., 2004). In both these fungal pathogens, the mating MAPK signalling pathway is often triggered by environmental cues such as nutrient limitation or stress (Hartmann et al., 1999; Bahn et al., 2007; Xue et al., 2007; Ko et al., 2009). The yeast form of both the *Candida* species is thought to be optimal for colonization, whereas the filamentous form (hyphal growth) of *C. albicans* is considered critical for pathogenesis, yet unfavourable for gut colonization (Liang et al., 2024). This prompts the obvious question as to how *C. albicans* manages to utilise its morphological plasticity during commensalism and pathogenic lifestyle. It is worth studying whether *C. albicans* and/or host have any control over *Cg*Yhi1-mediated morphological switchover in *C. albicans*, especially during commensalism in a healthy host environment.

**Figure 4.**
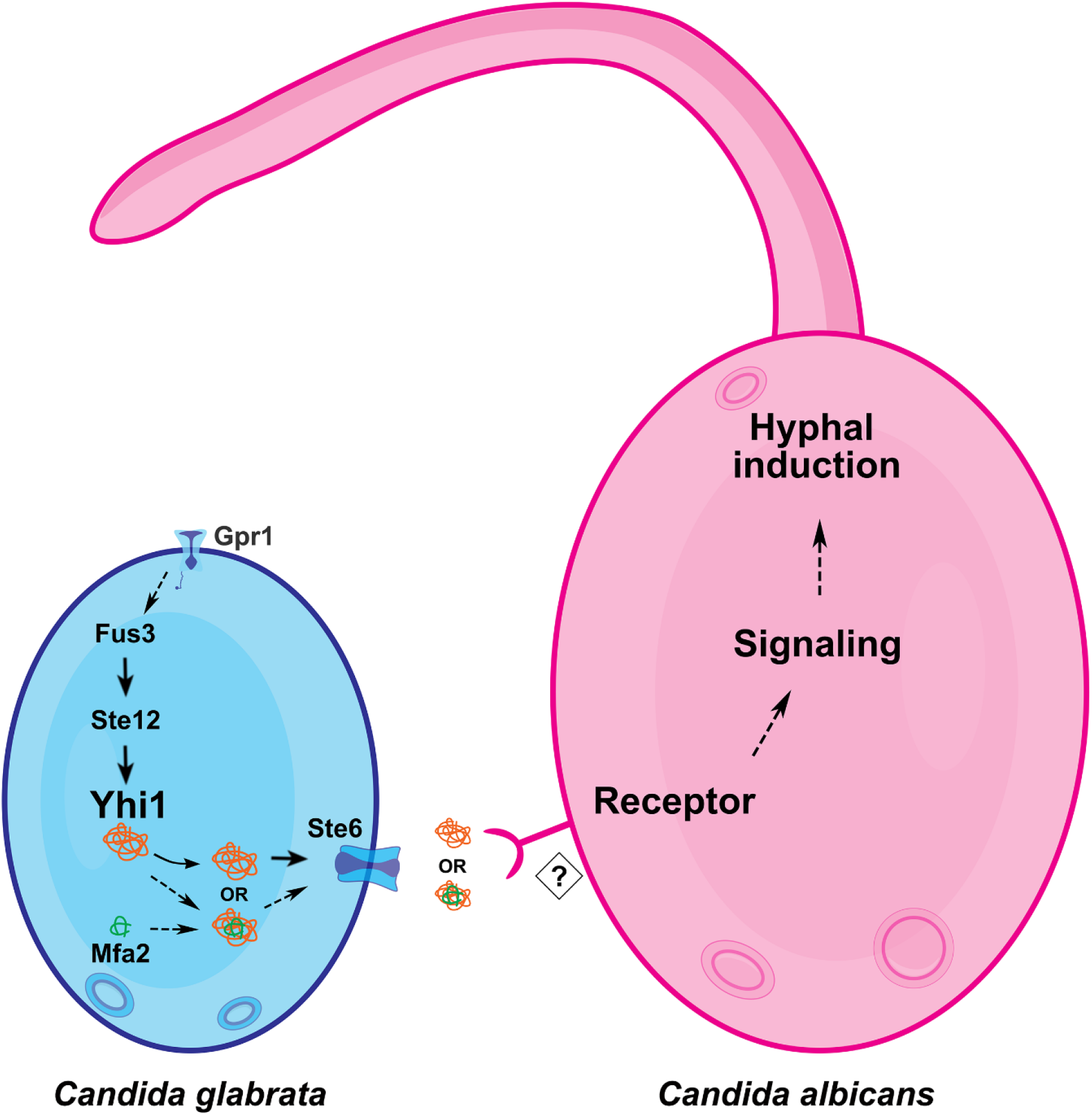
*C. glabrata* uses its mating-related MAPK signalling pathway to interact with *C. albicans*. A schematic representation of the mating signalling pathway in *C. glabrata* that leads to the expression of the novel small protein *Cg*Yhi1, which induces hyphal growth in *C. albicans*. *Cg*Yhi1 is effluxed, via non-classical secretion, through the pheromone transporter *Cg*Ste6, and the mating pheromone factor **a**2 (*Cg*Mfa2) likely aids in its efflux. In the extracellular milieu, the small protein *Cg*Yhi1 is likely recognized by a yet-unknown receptor, triggering the signalling pathway that induces hyphal growth in *C. albicans*. Overall, the findings from this study suggest that *C. glabrata*, which cannot form hyphae – a morphological characteristic essential for tissue invasion – has likely evolved to use its mating-related signalling pathway, to produce a novel secretory protein Yhi1, not to mate with the opposite mating type but rather to interact with *C. albicans*, enabling it to undergo hyphal growth and then piggyback on it to go deeper into the host tissue.

## Materials and Methods

All reagents, buffers, and media used in this study are listed in Supplementary Table S5.

## Yeast strains and culture media

*C. glabrata* WT strain included the auxotrophic strain CBS138 HTL (Schwarzmüller et al., 2014) and the parental strain CBS138. *C. albicans* WT strain included SC5314. The WT *S. cerevisiae* strains used included *S. cerevisiae* BY4741 (***MAT a***), *S. cerevisiae* BY4742 (***MAT α***), and the parental strain *S. cerevisiae* S288C. The *C. glabrata* deletion mutants with their details are listed in Supplementary Table S1. Other fungal strains used in this study are listed in Supplementary Table S4. While the *Candida* cultures were grown in YPD (1 % yeast extract, 2 % peptone, 2 % dextrose) at 37 °C, that of *S. cerevisiae* was grown at 30 °C. Cultures were washed in sterile PBS and adjusted to the required cell density. All strains were maintained on YPD plates (10 g/L yeast extract, 20 g/L protease peptone, 20 g/L dextrose, 20 g/L HiMedia^TM^ agar) and sub-cultured weekly. Glucose Minimal Media (2 % glucose, 6.79 g/L yeast nitrogen base, with ammonium sulphate, without amino acid) was used for all the assays.

## Generation of recombinant *E. coli* strains

A fusion PCR strategy was used to sequentially amplify the ‘TrxA-6xHis-TCS’ and the *CgYHI1* ORF, to generate a single cassette. *CgYHI1* ORF was cloned in frame with TrxA+6xHis+Thrombin Cleavage Site (TCS) into pET32b(+) plasmid between *XbaI* and *BamHI* sites, removing the enterokinase site of pET32b(+), as well as relocating the thrombin cleavage site to reduce the number of linker residues. The recombinant plasmid (pRPL130) was transferred to *E. coli* BL21 (DE3) pLysS strain for better expression of the recombinant protein. The *E. coli* strains and plasmids generated in this study are listed in Supplementary Table S4. The oligonucleotide primers (Sigma Aldrich) used are listed in Supplementary Table S4.

## Generation of yeast strains

*CgMFA2* deletion was performed by generation of a deletion cassette, where the ∼500 bp homologous flanking regions of *CgMFA2* were amplified by PCR (Schwarzmüller et al., 2014) using primers listed in Supplementary Table S4, and cloned into pBEVY-L (Miller III, 1998). Primers *CgMFA2*-3’UTR-F and *CgMFA2*- 3’UTR-R were used to amplify the 3’UTR of *CgMFA2*, which was cloned into pBEVY-L between *KpnI* and *SacI* sites. Primers *CgMFA2*-5’UTR-F and *CgMFA2*-5’UTR-R were used to amplify the 5’UTR of *CgMFA2,* which was cloned into the intermediate plasmid containing the 3’UTR of *CgMFA2* between *XbaI* and *BamHI* sites. *C. glabrata* CBS138 HTL was transformed as per the protocol described earlier (Gietz & Schiestl, 2007) using the *CgMFA2* deletion cassette – generated by digesting the final recombinant plasmid pRPL092 with *BamHI* and *KpnI* – yielding the *Cgmfa2*Δ deletion strain. The deletion was confirmed by PCR, and at least two independent transformants were confirmed and used for further experiments. Primers used to clone and construct the *MFA2* gene-deletion cassette, as well as for PCR confirmation, are listed in Supplementary Table S4. The corresponding strains are also listed in Supplementary Table S4.

*S. cerevisiae* strains expressing *Cg*Yhi1 were generated by cloning the ORF of *CgYHI1* into p426GPD (Mumberg et al., 1995). Primers *CgYHI1*-F and *CgYHI1*-R1 were used to amplify the ORF of *CgYHI1*, which was cloned into p426GPD between *BamHI* and *EcoRI* sites. The plasmid (pRPL033) was confirmed by restriction endonuclease digestion, and subsequently transferred to *S. cerevisiae* BY4741 and BY4742 strains using the protocol described earlier (Gietz & Schiestl, 2007). Transformants were confirmed by PCR. Primers used to construct the recombinant plasmid as well as for PCR confirmation of the transformants are listed in Supplementary Table S4. Plasmid and strains are also listed in Supplementary Table S4.

*C. glabrata yhi1*Δ strain expressing *Cg*Yhi1 or its variants was generated by cloning the complete ORF of *CgYHI1* or only the sequence of the variants into pBEVY-L (Miller III, 1998). Primers *CgYHI1*-F and *CgYHI1*-R2 were used to amplify the complete ORF of *CgYHI1* which was cloned into pBEVY-L between *BamHI* and *SalI* sites. Primers Yhi1^3-66^-F and *CgYHI1*-R2 were used to amplify the sequence of *CgYHI1,* without the first two amino acid residues, which was cloned into pBEVY-L between *BamHI* and *SalI* sites. Primers Yhi1^1-32^-F and Yhi1^1-32^-R were used to amplify the N-terminal half of *CgYHI1*, which was cloned into pBEVY-L between *BamHI* and *SalI* sites. Primers Yhi1^1-32^-F and Yhi1^1-32^-R were additionally used to amplify the N-terminal half of mutated *CgYHI1* (Yhi1^1-32(A18G, H22A)^ from a synthetic oligonucleotide sequence (ggatccATGAAGATGTATAAGCTAGCCTTGATAATAAAGCAGCAAATATCAGCAATCGGTGATGTTTGGGCTGACAAATTACATGCCTACAAAGTGAAATGCTGAgtcgac), which was cloned into pBEVY-L between *BamHI* and *SalI* sites. Primers Yhi1^33-66^-F and Yhi1^33-66^-R were used to amplify the C-terminal half of *CgYHI1*, which was cloned into pBEVY-L between *BamHI* and *SalI* sites. Primers Yhi1^21-52^-F and Yhi1^21- 52^-R were used to amplify the sequence between the 21^st^ and 52^nd^ amino acids of *CgYHI1*, which was cloned into pBEVY-L between *BamHI* and *SalI* sites.

The recombinant DNA strategy, to incorporate the pentapeptide motif along with a secretion signal peptide, was designed in such a way that the ADVWH pentapeptide sequence, which was added five amino acid residues after the secretion signal peptide and in the loop region of the GFP, would be accessible without altering the function of GFP. Primers SP-yeGFP-ADVWH-F and SP-yeGFP-ADVWH-R were used to amplify yeGFP from pKT209 (Sheff & Thorn, 2004), which was cloned into pBEVY-L between *BamHI* and *SalI*. Similarly, primers SP-yeGFP-F and SP-yeGFP- R were used to amplify yeGFP from pKT209, with a signal peptide sequence at the N-terminus (20 amino acids), which was cloned into pBEVY-L between *BamHI* and *SalI*. All the plasmids were confirmed by restriction endonuclease digestion as well as by PCR. Plasmids pRPL124, pRPL126, pRPL127 and pRPL128 were further confirmed by sequencing. Following confirmation, competent *C. glabrata yhi1*Δ cells were transformed with the recombinant plasmid pRPL034, pRPL035, pRPL100, pRPL101, pRPL124, pRPL126, pRPL127 or pRPL128. Transformants were confirmed by PCR. Primers used to construct the recombinant plasmid as well as for PCR confirmation of the transformants are listed in Supplementary Table S4. The corresponding plasmids and strains are also listed in Supplementary Table S4.

## Cell-free supernatant and hyphal induction assay

The CFS was harvested by removing 10 ml of medium from shake flasks (containing 200 ml YPD medium) at 3 h intervals over the course of 24 hpi. The cells were separated by centrifugation at 5000 *g* for 5 min. The supernatant was filtered through a 0.22 μm PES membrane filter (Millipore) to remove any residual biomass, and the filtrate was used for further experiments.

Overnight grown cells of *C. albicans* were collected by centrifugation at 5000 *g* for 5 min and washed with GMM (Xu et al., 2008). Forty-five μl each of harvested CFS (from different *C. glabrata* strains) and fresh GMM, and 10 μl of 5 x 10^5^ *C. albicans* cells were added per well in a 96-well plate (Thermo Scientific). The plate was incubated at 37 °C for 3 h and imaged. The assays were performed with technical duplicates and three independent biological replicates, and number of yeast, pseudohyphal and hyphal cells were counted and analysed for each experiment.

Hyphal auto-induction (without any trigger and for an unknown reason) in *C. albicans* can be observed occasionally. Therefore, we recommend use of freshly revived strains of *C. albicans* and *C. glabrata*; particularly, use of a *C. albicans SC5314* colony grown on GMM agar to inoculate a fresh GMM broth for the overnight growth.

## Hyphal induction assay using agar medium

*C. albicans*, *S. cerevisiae,* and *C. glabrata* strains were grown overnight in YPD/GMM broth and diluted to O.D. (A_600_) of 0.5. Five µl each of appropriately diluted cultures was spotted on YPD agar (2 cm apart) or GMM agar (1.5 cm apart) plates. The plates were incubated at 30 °C (for assay involving *S. cerevisiae*) or at 37 °C (for assay involving *Candida* species) for 10 days, and photographed using a DSLR camera. The *C. glabrata* mutants from the gene-deletion library (a kind gift from B. P. Cormack, Johns Hopkins School of Medicine) were similarly screened.

## Hyphal Induction Assay with Transwell Membrane Barrier

Nunc Carrier Plates with Inserts (Thermo Scientific) were used to visualize *C. albicans* hyphal induction in the presence or absence of *C. glabrata* or *S. cerevisiae*.

*C. albicans* cells grown overnight at 37 °C were collected by centrifugation at 5000 *g* for 5 min and washed with sterile 1X PBS. *C. albicans* cells were diluted to O.D. (A_600_) of 0.1 in 1 ml of GMM and added to the abluminal compartment. *C. glabrata* or *S. cerevisiae* cells were washed with 1X PBS and diluted by suspension in GMM to O.D. (A_600_) of 5, of which 750 μl was added to the luminal well set at medium height.

Similarly, for non-albicans *Candida* species, overnight grown cells of *C. tropicalis*, *C. dubliniensis*, or either of the two strains of *C. parapsilosis* were washed with sterile 1X PBS, diluted to O.D. (A_600_) of 0.1 in 1 ml of GMM, and added to the abluminal compartment. *C. glabrata*, *S. cerevisiae* or *C. albicans* were grown overnight, washed with 1X PBS, diluted to O.D. (A_600_) of 5 in 750 μl GMM, and added into the luminal well set at medium height. The plates were incubated at 37 °C for 3 h, and cells were observed and imaged as described.

## Organic and Aqueous Solvent Extraction of CFS

CFS of *C. glabrata* WT, grown in YPD medium for 18 h at 37 °C, was harvested by removing the cells by centrifugation at 5000 *g* for 5 min, followed by filtration (0.22 μm) of the supernatant to remove any residual biomass. The filtered CFS (100 µl) was then treated with equal volume (100 µl) of either acetonitrile, absolute ethanol, or ultrapure water, and incubated on a cell mixer with constant agitation for 1 h. The mixtures were dried by vacuum centrifugation (Concentrator Plus; Eppendorf) at ambient conditions, followed by resuspension in sterile water. The treated CFS was then used for *C. albicans* hyphal induction assay.

## RNA isolation and Reverse-Transcriptase PCR analysis

RNA was extracted as per the manufacturer’s protocol with minor modifications. Briefly, *C. glabrata* cells, grown in YPD media for 18 h at 37 °C, were collected by centrifugation at 10,000 *g* for 1 min at 4 °C, and the cell pellet was resuspended in 750 µL TRIzol reagent (Invitrogen). The 0.2 g acid-washed glass beads (425 to 600 µm diameter) were added to the pellet and the samples were vortexed vigorously (30 s, 1 min interval on ice, 5 cycles), followed by the addition of 200 µL chloroform and incubation for 5 min at room temperature (RT; 25 °C). Lysates were clarified by centrifugation at 15,000 g for 10 min at 4 °C and the aqueous phase was collected into a fresh tube. The RNA was precipitated by incubating with 500 µL isopropyl alcohol for 1 h at -20 °C. After precipitation, the samples were centrifuged at 12,000 *g* for 10 mins at 4 °C; the precipitated pellets were washed with 1 ml of 75 % chilled ethanol, and then resuspended in DEPC-treated water. It was then treated with DNaseI (Thermo Scientific, EN0521) at 37 °C for 15 min to ensure complete removal of any trace DNA contamination. The samples were re-treated with TRIzol and all the steps until the RNA precipitation were repeated. The precipitated pellets were resuspended in DEPC-treated water with 0.5 U/µL Protector RNase inhibitor (Roche, 3335399001). RNA integrity was confirmed by running the sample on 1 % agarose gel. cDNA was synthesized using RNA (∼500 ng) and iScript cDNA Synthesis Kit (BioRad, 1708891). cDNA samples were used for further PCR. Primers (CgYHI1-qPCR-F and CgYHI1-qPCR-R for *YHI1*, CgARP2-F and CgARP2-R for *ARP2*; Supplementary Table 3) were used at a final concentration of 500 nM. PCR amplifications were performed using an Applied Biosciences SimpliAmp thermocycler. The products were resolved on 2 % agarose gel and visualized using UVITEC Cambridge Essential gel documentation system.

## Yhi1 expression, purification, and assays

The *E. coli* BL21 (DE3) pLysS strain expressing 6xHis-tagged Yhi1 (His-Yhi1; RPL- ECS-04; Supplementary Table S3) was grown overnight in LB broth containing ampicillin at 37 °C. One ml of this overnight grown culture was added to 99 ml fresh LB broth and incubated at 37 °C for 4 h, followed by additional incubation at 37 °C for 4 h in the presence of 100 µM IPTG. Cells were collected by centrifugation at 13,000 *g* for 1 min and washed twice with 1X sterile PBS. Using a probe ultrasonicator (VibraCell, Sonics), the cells were sonicated at 40 % amplitude for 90 sec (2 sec pulses), with a 5 min interval on ice, for 5 cycles. The samples were then centrifuged at 8000 *g* for 10 min at 4 °C. The supernatant was collected and 1X protease inhibitor cocktail (Sigma Aldrich) was added to it. Nickel-NTA purification of the His-Yhi1 was performed as per the manufacturer’s (Merck) protocol. The purity of the eluted protein was checked by standard SDS-PAGE. The eluted protein was then dialysed using 5 mM Tris-Cl buffer (pH 7), and protein concentration was estimated using Bradford reagent (Sigma Aldrich).

For hyphal growth assay, overnight grown *C. albicans* cells were collected by centrifugation at 5000 *g* for 5 min and washed with GMM. In a 96-well plate (Thermo Scientific), 10 μl of 5 x 10^5^ *C. albicans* cells were added to 45 μl of fresh GMM in each well. Different dilutions of dialyzed His-Yhi1 (prepared in sterile GMM at 50 µM, 100 µM, 200 µM, 300 µM, 400 µM, or 500 µM) were added to these wells to make the total volume to 100 µl per well. The plate was incubated at 37 °C for 3 h and imaged using brightfield microscopy.

## Western blot analysis

For intracellular samples, *C. glabrata* WT and mutant cells were grown in YPD medium (10 ml) at 37 °C for 18 h. The CFS was transferred to fresh tubes, and the cells were washed twice with chilled 1X TE buffer (10 mM Tris-HCl, 1 mM EDTA; pH 8). The cells were then lysed using 425-600 μm acid-washed glass beads in 1x TE buffer containing protease inhibitors (Sigma-Aldrich) and a Neuation iRupt 24P tissue homogenizer at a speed setting of 4000 rpm with 30 sec vortex cycle (5 times) with 15 sec pauses in between vortex cycle, in multiple 2 ml microcentrifuge tubes.

The total cell lysate was then clarified by centrifugation in a refrigerated microfuge at 4 °C for 10 min at 10000 *g*. To bring the protein concentration in a range suitable for a western blot analysis, the harvested CFS was filtered (0.22 μm) to remove any residual biomass, and then subjected to concentration (20-fold) using 3 kDa molecular weight cut-off centrifugal filters (Amicon, Merck) at 5000 *g* for 45 min at RT.

Protein concentrations in the cell lysate and CFS were determined using Bradford reagent (Sigma Aldrich). Total protein (∼10 μg) was separated on a 12 % SDS–PA gel before transferring to a polyvinylidene difluoride (PVDF) membrane (GE Healthcare, pore size 0.2 μm), and blocked in 2.5 % Bovine Serum Albumin (BSA), dissolved in 1X Tris-buffered saline solution with 0.1 % Tween 20 (TBST). A rabbit polyclonal antiserum directed against Yhi1, which was produced by using synthetic Yhi1 protein (BiotechDesk, India), was used as the primary antibody. All antibody dilutions were prepared in 2.5 % BSA dissolved in 1X TBST. Membrane was incubated with the primary antibody (α-Yhi1; 1:1,000) for 3 h at RT, and washed thrice with 1X TBST prior to incubation with the secondary antibody (Goat anti-Rabbit IgG (H+L) Cross-Adsorbed Secondary Antibody, HRP-conjugated; 1:10,000) for 3 h at RT. The membrane was washed thrice with 1X TBST and developed using WESTAR ANTARES ECL substrate (Cyanagen), then imaged using iBright FL1500 Imaging System (Invitrogen) or Amersham ImageQuant 500 CCD imaging system (Cytiva).

## Molecular docking and MD simulation

Protein sequences of *Cg*Yhi1 (*CAGL0D06666g*) and *Cg*Mfa2 (*CAGL0C01919g*) were retrieved from *Candida* genome database (CGD) and used for 3D structure prediction by RoseTTAFold modelling method (Baek et al., 2021) in Robetta online server (https://robetta.bakerlab.org/). The predicted structures were then evaluated using Ramachandran plot.

To understand the molecular interaction between *Cg*Yhi1 and *Cg*Mfa2, molecular dynamics (MD) simulation studies were performed. Molecular docking was performed using LightDock online server (Jiménez-García, 2018), wherein *Cg*Yhi1 was used as protein and *Cg*Mfa2 was used as ligand. Both the protein and ligand backbone were kept flexible for the docking studies. Based on the docking score, the top 10 predicted docked models were used for MD simulation studies. The docked structures were prepared using the solution builder module in CHARMM-GUI v3.7 (Jo et al., 2008). MD simulations were performed using GROMACS 2019.2 with CHARMM36m force field. The simulation system was prepared with the help of a CHARMM-GUI (Jo et al., 2008) server, where protein-ligand complexes were placed in a rectangular box of water molecules. Physiological ionic strength of 0.15 moles/L was achieved using Na^+^ or Cl^-^ as counter-ions, and the temperature was set to 303.15 kelvin. The system was subjected to equilibration, including NVT and NPT ensembles. During equilibration, the temperature was controlled using the v-rescale thermostat, and the pressure was controlled using the Berendsen pressure coupling. After equilibration, the production MD simulations were carried out for 100 ns simulation time, where atomic coordinates were captured every 0.1 nanosecond (ns). Root-mean-square deviation (RMSD) graphs were plotted using QT-Grace software v026. Visualization of the structures was done using PyMOL 2.5.2.

## Multiple sequence alignment

Multiple sequence alignment of amino acid sequences of Ste6 from *C. glabrata* (*CAGL0K00363g*), *S. cerevisiae* (YKL209C), *C. albicans* (orf19.7440), and *Schizosaccharomyces pombe* (SPBC25B2.02c), was performed using the T-Coffee web server (Notredame et al., 2000), and the resulting alignment was rendered through ESPript (Robert & Gouet, 2014). Multiple sequence alignment of *Cg*Mfa2 (*CAGL0C01919g*) with *Sc*Mfa1 (*YDR461W*) and *Sc*Mfa2 (*YNL145W*) was performed using Clustal Omega (Sievers et al., 2011).

### *In silico* Motif Search

The amino acid sequence of *Cg*Yhi1 was submitted to the InterProScan web service for scanning against InterPro protein signature databases (Jones et al., 2014; Blum et al., 2021). Additionally, *Cg*Yhi1 sequence was used as query in the MOTIF Search (https://www.genome.jp/tools/motif/MOTIF) hosted by the GenomeNet online database and computational services (Bioinformatics Center, Kyoto University, Japan). Databases used, for finding out sequence motifs in the query sequence, were PROSITE pattern & profile (Sigrist et al., 2012), NCBI-CDD (Marchler-Bauer et al., 2012) and Pfam (Finn et al., 2014). The pentapeptide sequence pattern, A-D-V-W-H and/or A-V-V-P-H, was searched against protein sequence libraries using the Motif search service on the GenomeNet online server (https://www.genome.jp/tools/motif/MOTIF2). The target sequence libraries included GenBank, UniProt, RefSeq, and PDBSTR.

## Synthetic Yhi1 peptide derivatives handling and assays

All five short Yhi1-derived peptides were synthesized commercially (GenetoProtein, India). Stock solutions (1 mM) were prepared in ultrapure water, aliquoted, and stored at -20 °C. Working solutions were prepared in GMM at the final concentration indicated in the assay.

For hyphal growth assay, overnight grown *C. albicans* cells were collected by centrifugation at 5000 g for 5 min and washed with GMM. In a 96-well plate (Thermo Scientific), 10 μl of 5 x 10^5^ *C. albicans* cells were added to 45 μl of fresh GMM in each well. Different dilutions of each synthetic peptide (prepared in sterile GMM at 25 µM, 50 µM, 75 µM, 100 µM, or 250 µM) were added to these wells to make the total volume to 100 µl per well. The plate was incubated at 37 °C for 3 h and imaged using brightfield microscopy.

## Assay using ADVWH-motif tagged yeGFP

The CFS from *C. glabrata yhi1*Δ strains expressing SP-ADVWH-yeGFP or SP- yeGFP was collected and filtered before using in the hyphal induction assay. The same CFS was further concentrated, as mentioned earlier, for live-cell imaging, where 10 µl of 20x concentrated CFS and 8 µl of fresh medium were inoculated with 2 µl of *C. albicans* (O.D. (A_600_) of 0.05) on a sterile 35 mm glass-bottom Petri dish (SPL Life Sciences), and incubated at 37 °C for 3 h. Following the incubation, the medium was gently aspirated, and the cells were washed twice with sterile 1X PBS to remove residual medium, followed by resuspension in 1X PBS for microscopy.

The localization of secreted ADVWH-yeGFP or yeGFP on *C. albicans* was imaged as described.

## Microscopy and Imaging

Hyphal induction assays using plates with or without transwell membrane barrier, were observed and imaged using bright-field microscopy on a Nikon Eclipse Ti2 inverted microscope with a 20x/0.50NA objective or an extra-long working distance (ELWD) 40x/0.60NA objective, with a DIC filter set.

For time-lapse live-cell imaging, *C. glabrata* or *S. cerevisiae* cells (O.D. (A_600_) of 5 in 750 μl GMM) were added into the luminal well set at medium height, and *C. albicans* cells (O.D. (A_600_) of 0.1 in 1 ml of GMM) were added to the abluminal compartment of the transwell plate. A constant temperature of 37°C was maintained for 3 h and images were taken every 10 min using a Nikon Eclipse Ti2 inverted microscope with an extra-long working distance (ELWD) 40x/0.60NA objective with a DIC filter set.

Hyphal induction assay on agar medium was observed and photographed using a Nikon D90 Digital SLR camera and a 105 mm, *f* 2.8 macro lens.

To visualize localisation of secreted ADVWH-yeGFP or yeGFP on *C. albicans*, z- stack images (with 0.3-micron steps) were captured by laser scanning microscopy using a PlanApo λ 100x oil immersion objective (SR HP Apo TIRF 100XAC Oil/1.49 NA) on a Nikon Eclipse Ti2 inverted microscope mounted with an AX confocal system (Nikon). GFP excitation and emission were performed using the FITC channel (Ex. 405-488 nm; Em. 505-547 nm). Maximum intensity projection of best z-slices was used to create the final image. The outline of the cells was marked by a dashed line using Adobe Photoshop CC 2019 software.

The bright-field microscopy images were acquired using the NIS-Elements AR acquisition software (5.40.00, Build 1637; Nikon) with the DS-Fi3 CMOS camera (Nikon). The fluorescence microscopy images were acquired and processed using the NIS-Elements AR acquisition software (5.42.06, Build 1821; Nikon) with an AX scan head. Images and videos were processed using ImageJ (https://imagej.net/software/imagej/), NIS-Elements AR, and Adobe Photoshop CC 2019.

## Statistical analyses

Statistical data involving analyses of percentage hyphal induction was evaluated using two-way ANOVA with Tukey’s multiple comparison test, and percentage hyphal induction of *C. glabrata* CFS and *S. cerevisiae* CFS was evaluated using an unpaired two-tailed *t*-test with Welch’s correction. Mean values, and corresponding standard error of mean, for quantitative data on percentage hyphal induction, were calculated using GraphPad Prism software 8.0.1.

## Data Availability Statement

All data supporting the findings of this study are available within the manuscript and its Supplementary Information files. Sequences for all genes mentioned in this study are available on Candida Genome Database (http://www.candidagenome.org/), Saccharomyces Genome Database (https://www.yeastgenome.org/), and NCBI (https://www.ncbi.nlm.nih.gov/nucleotide/). Source data files will be shared upon request.

## Acknowledgement

We thank P. J. Bhat and N. S. Punekar (Indian Institute of Technology Bombay) for sharing plasmids. We thank B.P. Cormack (Johns Hopkins School of Medicine) for the kind gift of the *C. glabrata* mutant library, and J. Berman (Tel Aviv University) and K. Kuchler (Medical University of Vienna) for sharing yeast strains. We also thank all members of the Fungal Pathobiology and Host-Pathogen Interaction group for helpful discussions and suggestions. We thank Industrial Research and Consultancy Centre (IRCC), IIT Bombay, for use of the Super-Resolution Confocal Microscopy facility. We thank SpaceTime High Performance Computing Complex (ST HPCC), Computer Centre, IIT Bombay, for providing computational infrastructure for MD studies.

## Author Contributions

V.A.P. conceived, designed, and performed the experiments, analysed the data, and co-wrote the manuscript. P.M.T. contributed experimental data by generating initial recombinant plasmids, contributed *in silico* data, and performed analysis, and contributed to the reviewing and editing of the manuscript. H.P. contributed experimental data by generating a deletion mutant and analysed the data. R.N.P. conceived and supervised the project, provided advice on protocols, experimental design and implementation, data interpretation, and provided reagents, materials, and analysis tools, and led the drafting of the manuscript.

