## Supplementary Figures & Legends for "A novel *Candida glabrata* protein regulated by mating signalling pathway shapes inter-species interaction"

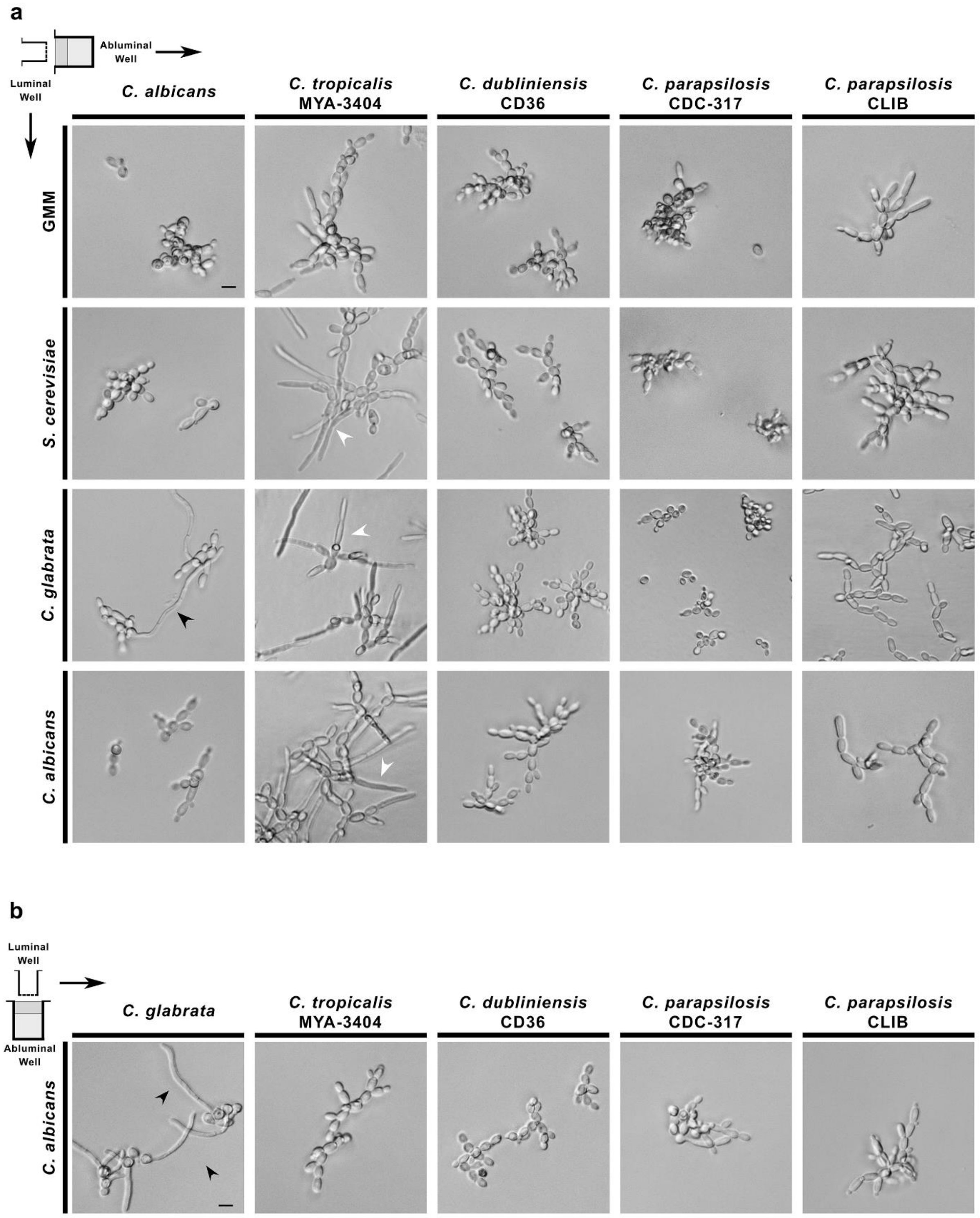


**Supplementary Figure S1. Hyphal growth is induced specifically during interaction between *C. glabrata* and *C. albicans*. (a)** Representative brightfield micrographs showing the response of various *Candida* species, cultured in an abluminal well of a Transwell co-culture setup, to *C. glabrata* or *C. albicans* cultured in the luminal well. GMM and *S. cerevisiae* culture in the luminal well were used as the control conditions. While black arrowheads mark the hyphae developed by *C. albicans* in response to *C. glabrata*, white arrowheads denote the rather non-specifically induced hyphal growth of *C. tropicalis* in response to any species. Scale bar, 5 μm. **(b)** Representative brightfield micrographs showing the response of *C. albicans*, grown in an abluminal well of a Transwell co-culture setup, to various non-albicans candida species cultured in the luminal well. White arrowheads mark true hyphae. Scale bar, 5 μm.

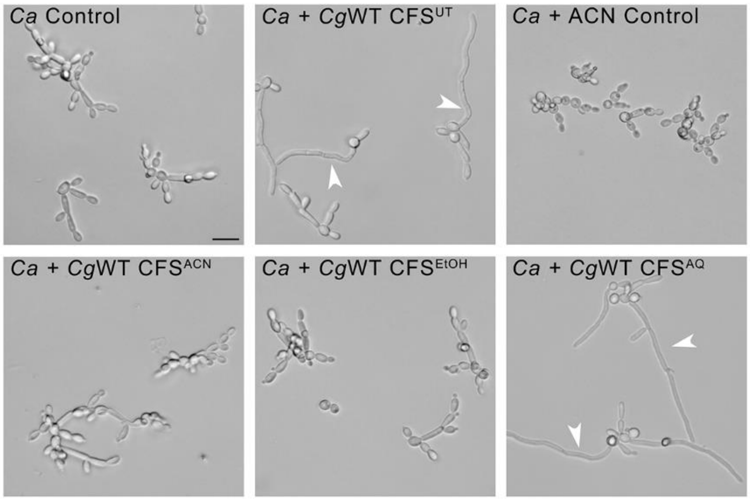


**Supplementary Figure S2. The inducer of hyphal growth in *C. albicans* is a secretory protein, rather than a small molecule, of *C. glabrata*.** Representative brightfield micrographs showing the response of *C. albicans* to treatment with CFS of WT *C. glabrata* extracted with an organic solvent [Acetonitrile (ACN) or Ethanol (EtOH)] or water (AQ). White arrowheads denote true hyphae developed by *C. albicans*. CFS$^{UP}$, unprocessed CFS; ACN control, residual acetonitrile solvent control. Scale bar, 5 µm.

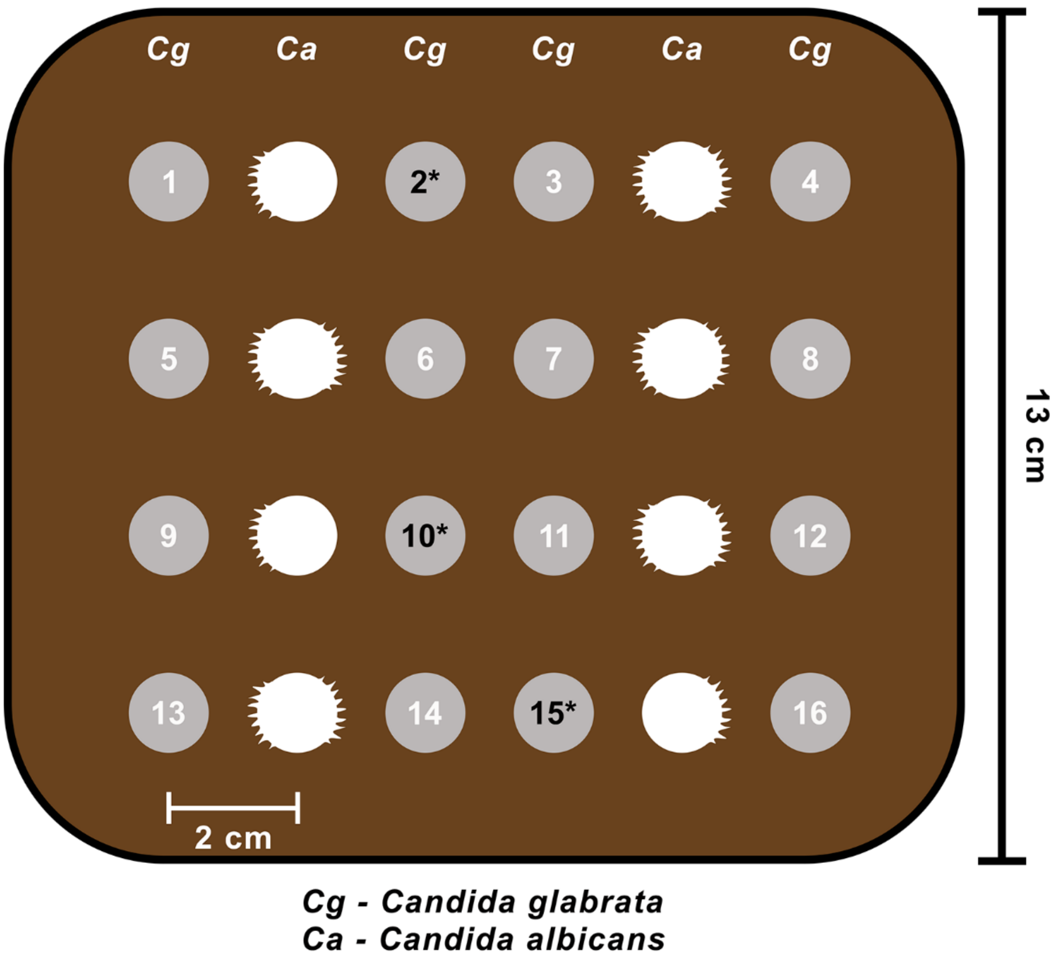


**Supplementary Figure S3. A schematic depicting the strategy used to screen the *C. glabrata* gene-deletion library to identify the genes involved in inducing the hyphal growth in *C. albicans*.** Five µl each of cultures (0.5 O.D.$_{600nm}$) of individual *C. glabrata* mutant and WT *C. albicans* was inoculated, at a distance of ~2 cm from each other, on YPD agar medium in large petri dishes, and incubated at 37$^{o}$C for 10 days post inoculation (dpi). Numbers (in white and black font) denote colonies of individual gene-deletion mutants of *C. glabrata*. Asterisks mark the colonies of the representative *C. glabrata* mutants that were impaired in inducing hyphal growth in *C. albicans*.

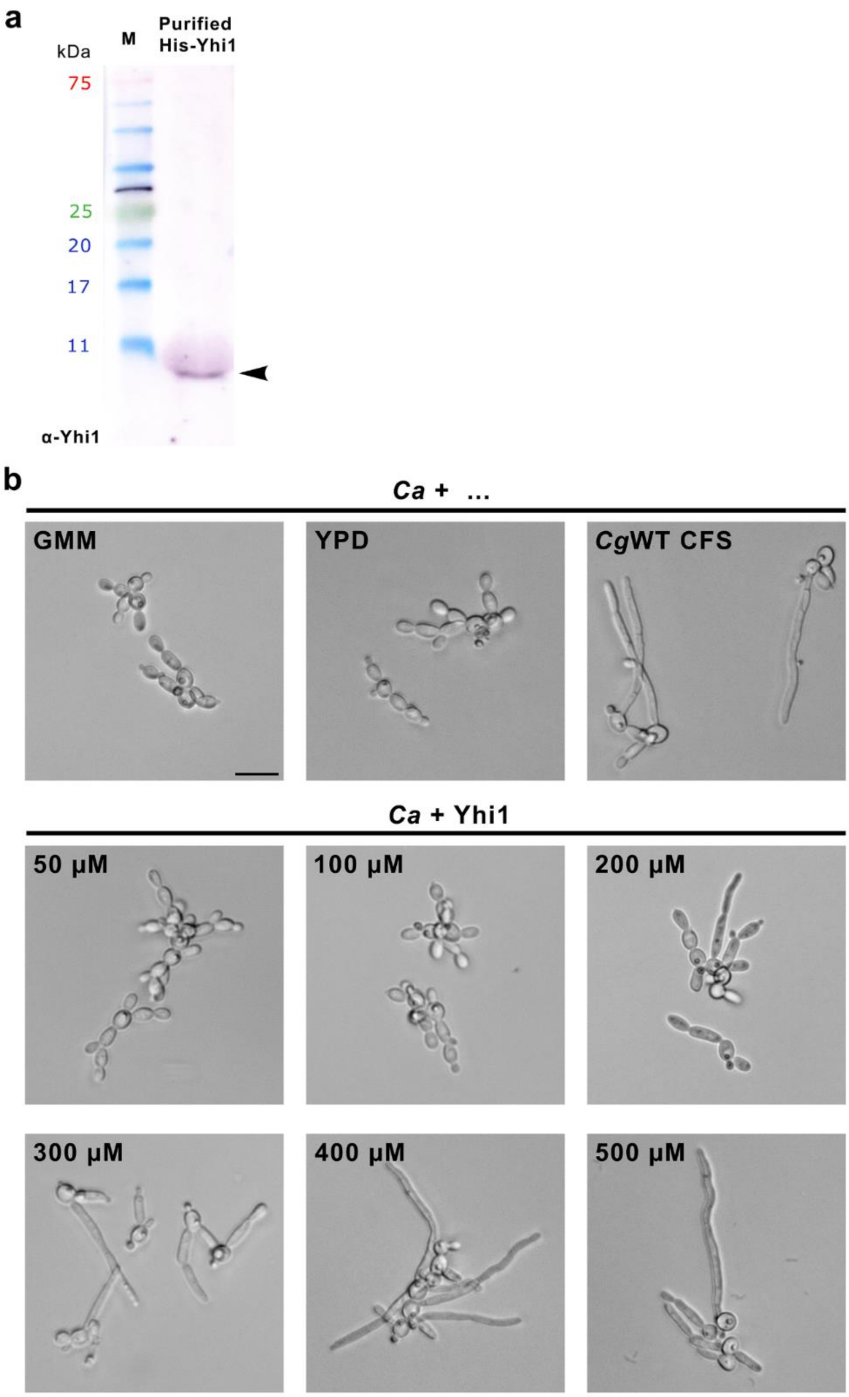


**Supplementary Figure S4. Minimum concentration of purified Yhi1 required to induce hyphal growth in *C. albicans*. (a)** Immunoblot analysis, using a polyclonal α-Yhi1 antibody, of purified Yhi1. Black arrowhead marks the position of the purified monomeric Yhi1 (~11 kDa) including the linker peptide. Numbers on the left denote the molecular weights (kDa) of corresponding proteins from the molecular marker

(M). **(b)** Representative brightfield micrographs showing the response of *C. albicans* when treated with different concentrations of heterologously expressed (in *E. coli*) and purified Yhi1. Scale bar, 10 μm.

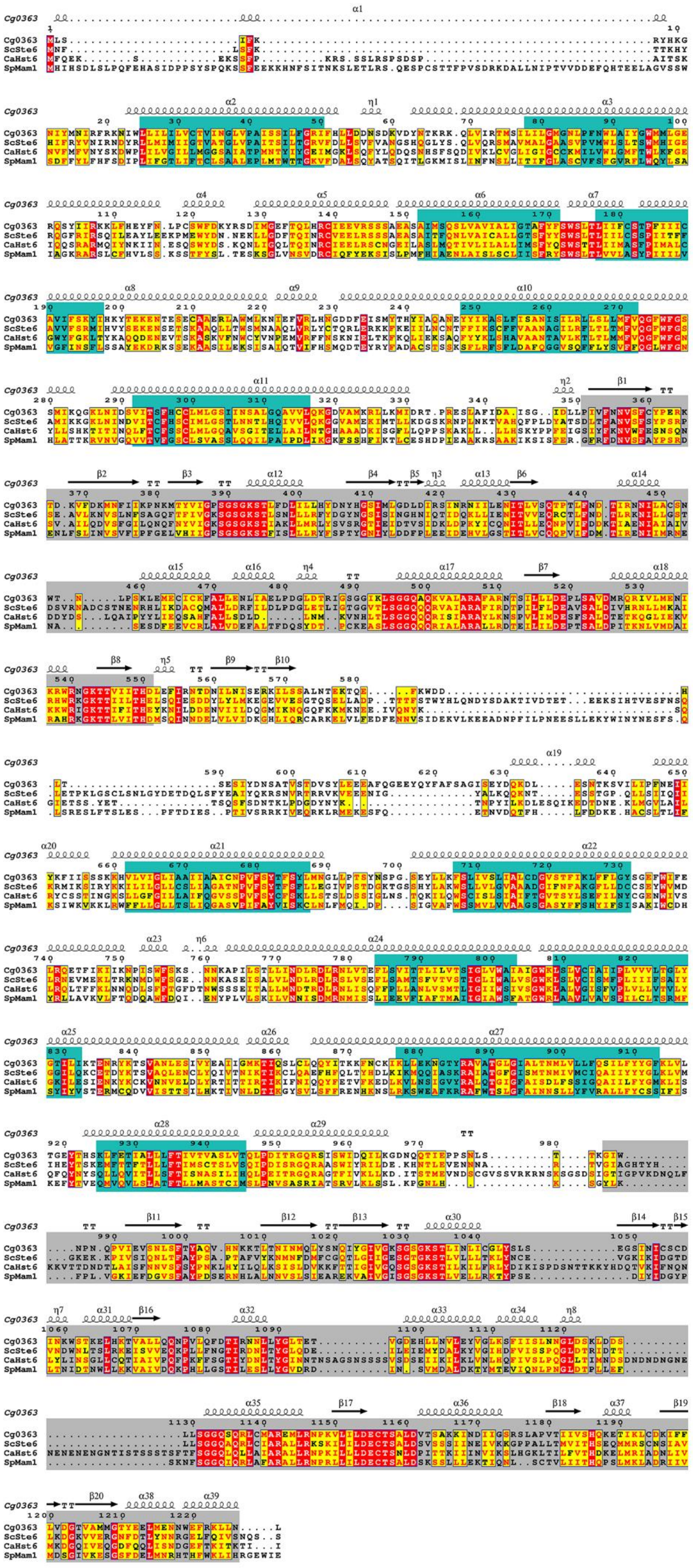
Cg0363
ScSte6
CaHst6
SpMam1

**Supplementary Figure S5. Multiple sequence alignment of pheromone transporter Ste6 from different ascomycetes yeast species.** Multiple sequence alignment of *C. glabrata* Ste6 (*CAGL0K00363g*; Cg0363) with Ste6 from *S. cerevisiae* (YKL209C; ScSte6), *C. albicans* (orf19.7440; CaHst6), and *Schizosaccharomyces pombe* (SPBC25B2.02c; SpMam1) performed using the T-Coffee Web-Server and the resulting alignment visualized through ESPript. The amino acid sequence identity values of *Cg*Ste6 with *Sc*Ste6, *Ca*Hst6, and *Sp*Mam1, are 42%, 31%, and 23%, respectively. The secondary structural features for *Cg*Ste6 are indicated above the sequence alignment. The transmembrane region (teal) and P-loop containing nucleoside triphosphate hydrolases (grey) are highlighted.

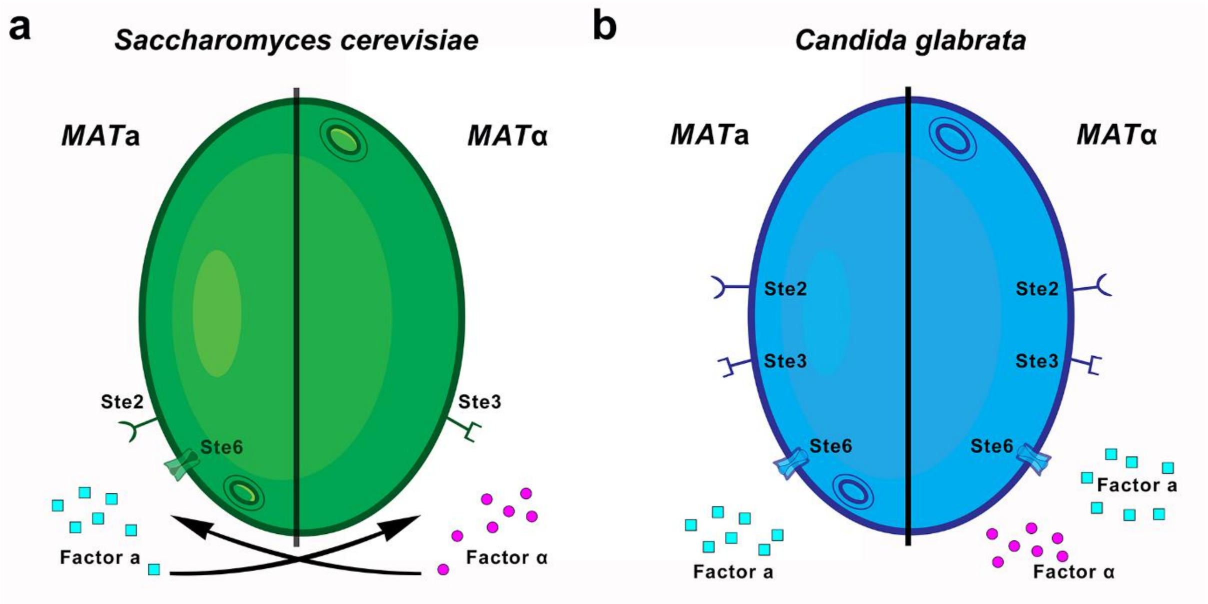


**Supplementary Figure S6. A comparative sketch of pheromone production and perception in *S. cerevisiae* and *C. glabrata*. (a)** A schematic representation of the mating signalling pathway, involving the mating-type-specific expression, transport, and perception of pheromones factor a and α, in *S. cerevisiae*. Mating-type **a** cells (***MAT*a**) express the pheromone factor **a** (filled circles) and its transporter Ste6, along with the receptor Ste2 for the opposite mating pheromone. Similarly, mating-type **α** cells (***MAT*α**) express the pheromone factor **α** (filled squares) and the receptor Ste3 for factor **a**. **(b)** A schematic representation of the promiscuous mating machinery in *C. glabrata* (inferred from Muller et al., 2008; Kumari et al., 2018).

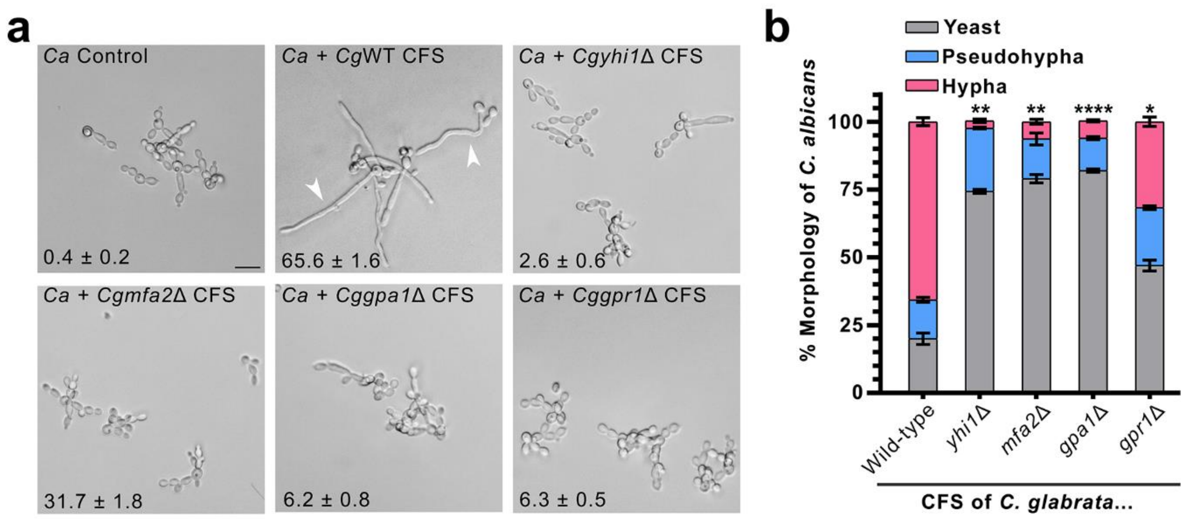


**Supplementary Figure S7**. ***Cg*Yhi1 likely requires mating pheromone a for its transport via *Cg*Ste6.** Representative brightfield micrographs **(a)** and bar graph **(b)** showing the response of *C. albicans* to treatment with CFS from the wild type or specified gene-deletion mutant of *C. glabrata*. White arrowheads mark true hyphae developed by *C. albicans*. Scale bar, 5 µm. Inset values in **(a)** represent the percentage number of *C. albicans* hyphae observed under the specified condition. The data represent mean ± S.E.M from three independent experiments. n = 300 each. Statistical analysis was applied relative to *C. glabrata* WT. $P < 0.05$ (*); $P < 0.01$ (**); $P < 0.0001$ (****); two-way ANOVA with Tukey's multiple comparisons.

**a**

```
CgMfa2   MQP----TIEATQKDNTQEKRDNYIVKGFFWSPDCVIA   34
ScMfa1   MQPST--ATAAPKEKTSSEKKDNYIIKGVFWDPACVIA   36
ScMfa2   MQPITTASTQATQKDKSSEKKDNYIIKGLFWDPACVIA   38
         ***    :  * ::..:.**:****:**.**.* ****
```

**b**

| | ResName_A | ResID_A | Chain_A | Atom_A | ResName_B | ResID_B | Chain_B | Atom_B | Distance |
|---|---|---|---|---|---|---|---|---|---|
| TRP_ILE | TRP | 21 | A | CZ2 | ILE | 33 | B | 3HG2 | 1.802386999 |
| LEU_PHE | LEU | 6 | A | 2HD1 | PHE | 26 | B | HN | 2.111351013 |
| LEU_ILE | LEU | 6 | A | 1HD2 | ILE | 21 | B | CG1 | 2.116555691 |
| LEU_CYS | LEU | 39 | A | HG | CYS | 31 | B | O | 2.221213818 |
| ILE_CYS | ILE | 17 | A | 2HG2 | CYS | 31 | B | HN | 2.614689112 |
| MET_ILE | MET | 1 | A | CE | ILE | 21 | B | HA | 2.738320112 |
| TRP_PHE | TRP | 42 | A | HE1 | PHE | 26 | B | HZ | 2.795815468 |
| GLN_PHE | GLN | 46 | A | HG2 | PHE | 25 | B | CE2 | 2.828815699 |
| ILE_ILE | ILE | 17 | A | HA | ILE | 33 | B | HD1 | 2.871984005 |
| TRP_CYS | TRP | 42 | A | CH2 | CYS | 31 | B | SG | 2.90960598 |
| VAL_ILE | VAL | 20 | A | HB | ILE | 33 | B | 1HG1 | 3.305887938 |
| LYS_GLU | LYS | 5 | A | HE2 | GLU | 6 | B | HB1 | 3.618135929 |
| PHE_ASP | PHE | 38 | A | CG | ASP | 30 | B | OD1 | 2.443645954 |
| PHE_CYS | PHE | 35 | A | CG | CYS | 31 | B | HN | 2.463675022 |
| GLN_THR | GLN | 13 | A | OE1 | THR | 4 | B | HG1 | 2.467816353 |
| TRP_TRP | TRP | 42 | A | CZ2 | TRP | 27 | B | O | 2.730384588 |
| ILE_THR | ILE | 9 | A | CD | THR | 4 | B | HG1 | 2.762933969 |
| ALA_ILE | ALA | 16 | A | HN | ILE | 33 | B | 2HG2 | 3.746504545 |
| PHE_SER | PHE | 38 | A | CG | SER | 28 | B | HA | 3.996723652 |

**c**

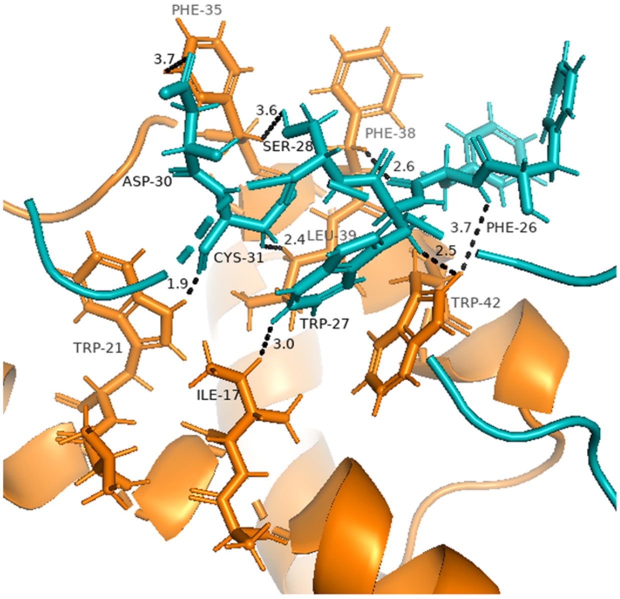


**Supplementary Figure S8**. ***Cg*Yhi1 likely interacts with mating pheromone *Cg*Mfa2. (a)** Multiple sequence alignment of *Cg*Mfa2 (*CAGL0C01919g*) with *Sc*Mfa1 (*YDR461W*) and *Sc*Mfa2 (*YNL145W*), with the residues comprising the functional mature mating pheromones of *S. cerevisiae* highlighted in bold (grey shade). The

*Cg*Mfa2 residues involved in interaction with *Cg*Yhi1 are highlighted in blue. **(b)** List of all likely stably interacting amino acid residues of *Cg*Yhi1 and *Cg*Mfa2 obtained after 100 nanosecond MD simulation runs. **(c)** Structural representation of interacting residues of *Cg*Yhi1-*Cg*Mfa2 protein complex in one of the states. Amino acid residues of *Cg*Yhi1 are shown as orange sticks, and those of *Cg*Mfa2 are shown as cyan sticks.

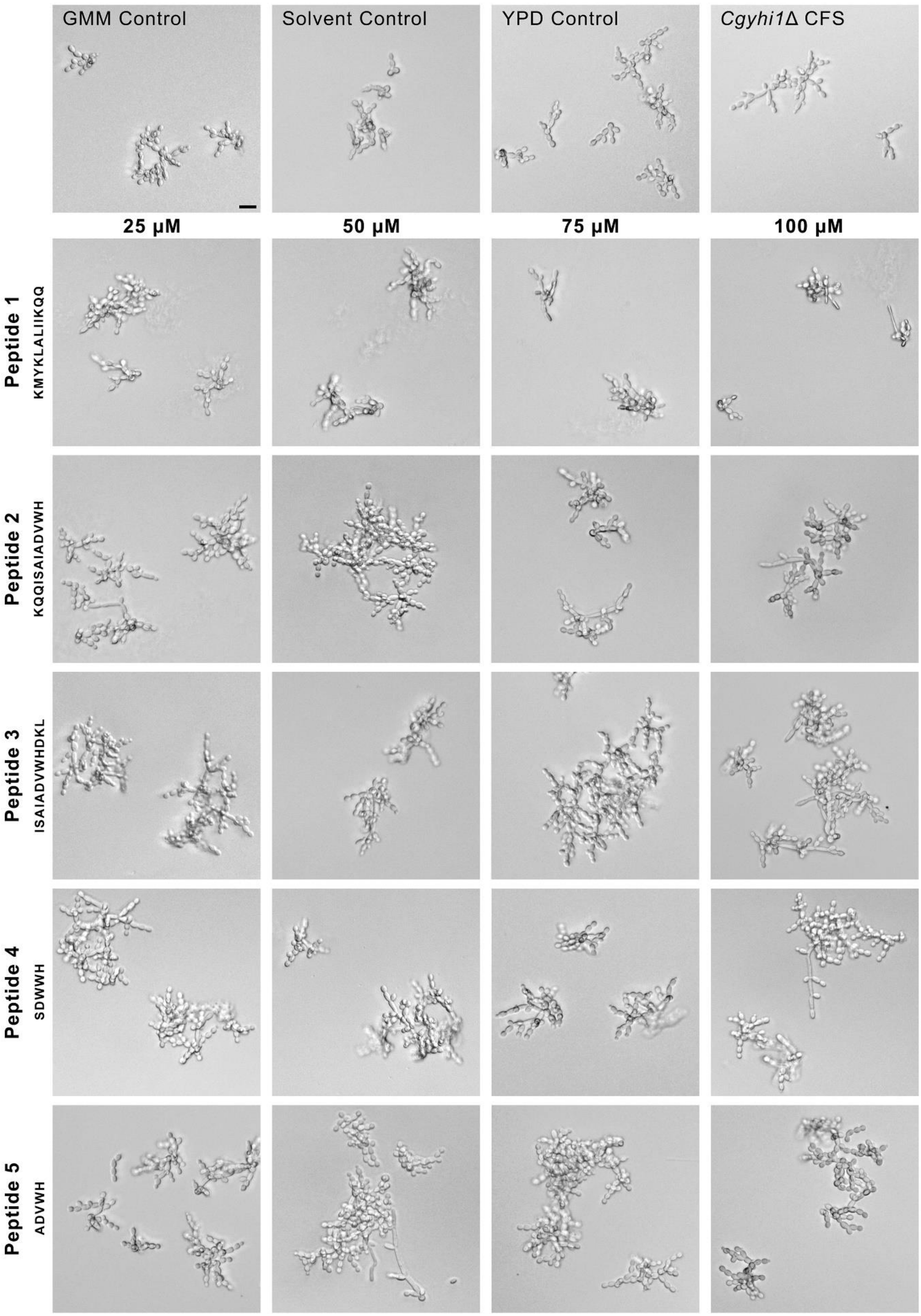
GMM Control
Solvent Control
YPD Control
*Cgyhi1*Δ CFS
25 µM
50 µM
75 µM
100 µM
Peptide 1
KMYKLALIIKQQ
Peptide 2
KQQISAIADVWH
Peptide 3
ISAIADVWHDKL
Peptide 4
SDVWH
Peptide 5
ADVWH

**Supplementary Figure S9. Effect of synthetic peptides, representing different truncated versions of Yhi1, on the morphology of *C. albicans*.** Representative brightfield micrographs showing the response of *C. albicans* when treated with various synthetic peptides representing the specified truncated Yhi1 at different concentrations. Peptide 1, Yhi1$^{2-13}$ (without ADVWH pentapeptide); Peptide 2, Yhi1$^{11-22}$ (includes ADVWH pentapeptide); Peptide 3, Yhi1$^{14-25}$ (includes ADVWH pentapeptide); Peptide 4, modified Yhi1$^{18-22}$ (modified ADVWH pentapeptide); Peptide 5, Yhi1$^{18-22}$ (ADVWH pentapeptide alone). Scale bar, 10 μm.

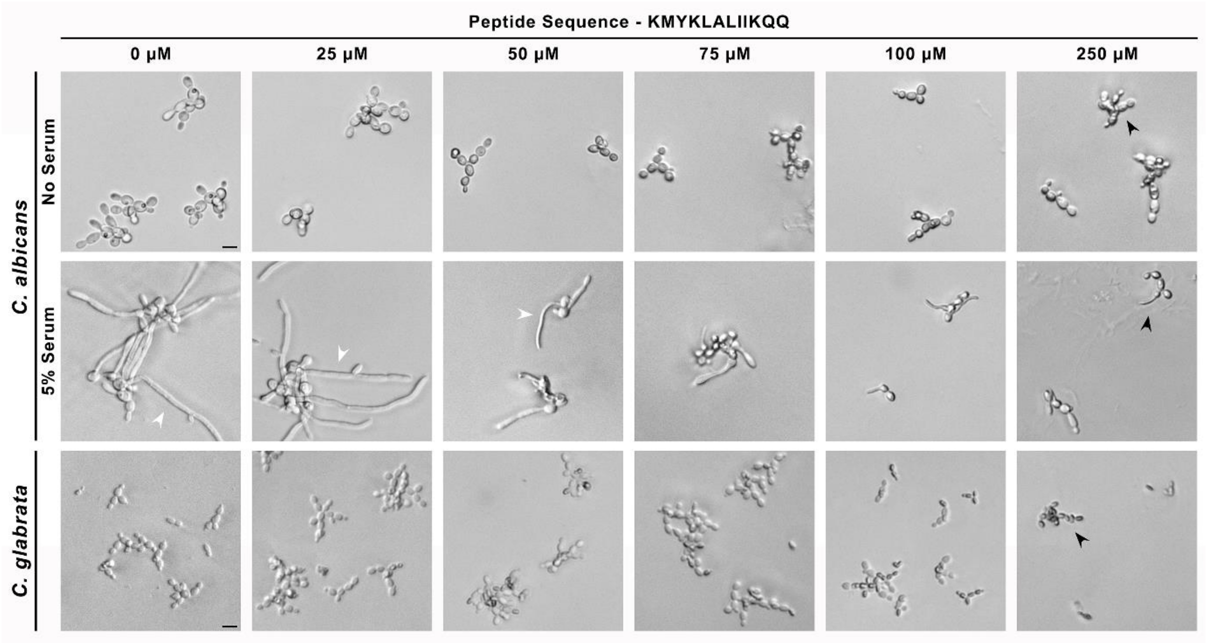


**Supplementary Figure S10. A short synthetic peptide derived from the *Cg*Yhi1 N-terminus exhibits antifungal activity.** Representative brightfield micrographs showing the response of *C. albicans* (upper panels) when treated with different concentrations of a short synthetic peptide (Yhi1$^{2-13}$ - Peptide 1 from the previous experiment – Supplementary Figure S9) representing the N-terminus of Yhi1, in the presence or absence of serum. Lower panel with representative brightfield micrographs depicting the effect of Yhi1$^{2-13}$ on *C. glabrata*. White arrowheads depict hyphal growth in *C. albicans*; whereas, the black arrowheads mark the impaired growth of *C. albicans* and *C. glabrata*. Scale bar, 10 μm.

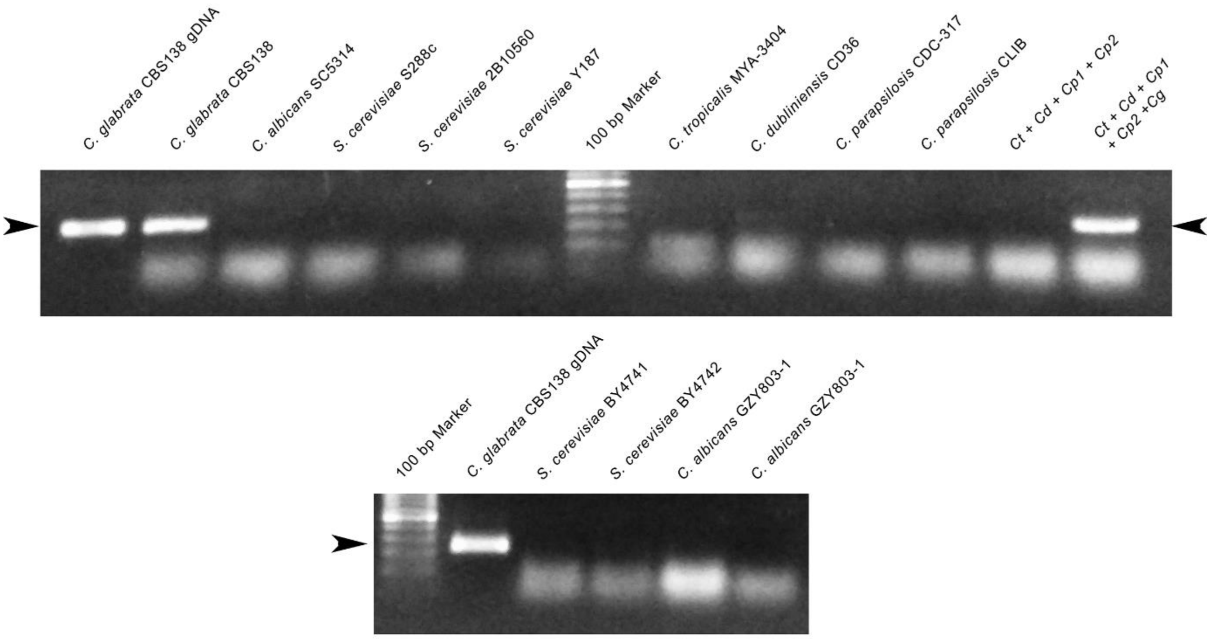


**Supplementary Figure S11. *CgYHI1* is a potential biomarker for a quick and accurate diagnosis of mixed-species candidiasis involving *C. glabrata*.** Agarose gel electrophoresis showing results of a colony PCR using *CgYHI1*-specific oligonucleotide primers, performed with various yeast strains, either alone or in combination, with or without *C. glabrata* in the combination. Ct, *C. tropicalis*; Cd, *C. dubliniensis*; Cp1, *C. parapsilosis* CDC-317; Cp2, *C. parapsilosis* CLIB; Cg, *C. glabrata*. White arrowheads mark the position of the amplified PCR product using *CgYHI1*-specific primers (~220 bp).

**Supplementary Movies:**

**Supplementary Movie 1.** *C. glabrata* induces hyphal growth in *C. albicans* in an in vitro assay using a Transwell co-culture setup.

**Supplementary Movie 2.** Response of *C. albicans* towards *S. cerevisiae* in an in vitro assay using a Transwell co-culture setup.

**Supplementary Tables:**

**Supplementary Table S1.** List of *C. glabrata* gene-deletion mutants used in this study.

**Supplementary Table S2.** List of *C. glabrata* gene-deletion mutants impaired in inducing hyphal growth in *C. albicans*.

**Supplementary Table S3.** List of proteins, with their source, containing either ADVWH or AVVPH pentapeptide sequence pattern.

**Supplementary Table S4.** List of *E. coli* and yeast strains, plasmids, and oligonucleotide primers used in this study.

**Supplementary Table S5.** List of reagents, growth media, and buffers used in this study.
