## Supplementary Table S1 for "A novel *Candida glabrata* protein regulated by mating signalling pathway shapes inter-species interaction"

List of *C. glabrata* deletion strains (Schwarzmüller et al., 2014; Purohit & Gajjar, 2022) screened in this study

| Well | Systematic Name | Deletion Status | Standard Name | Description | Feature Type | *S. cerevisiae* Ortholog | Link to CGD | Reference |
|---|---|---|---|---|---|---|---|---|
| 1A1 | CAGL0A00451g | Verified | *PDR1* | Zinc finger transcription factor, activator of drug resistance genes via pleiotropic drug response elements (PDRE); regulates drug efflux pumps and controls multi-drug resistance; gene upregulated and/or mutated in azole-resistant strains | ORF, Verified | *PDR1* | CAGL0A00451g | Schwarzmüller et al., 2014 |
| 1A2 | CAGL0A00517g | No Deletion Verified | | Ortholog(s) have calcium transmembrane transporter activity, phosphorylative mechanism activity and role in calcium ion transport, cellular calcium ion homeostasis, transmembrane transport | ORF, Uncharacterized | *PMC1* | CAGL0A00517g | Schwarzmüller et al., 2014 |
| 1A3 | CAGL0A01133g | Verified | | Ortholog(s) have role in cellular response to hydrogen peroxide and mitochondrial inner membrane, vacuolar membrane localization | ORF, Uncharacterized | *MDL2* | CAGL0A01133g | Schwarzmüller et al., 2014 |
| 1A4 | CAGL0A03586g | Verified | | Ortholog(s) have role in response to pheromone triggering conjugation with cellular fusion and fungal-type cell wall localization | ORF, Uncharacterized | *AFB1* | CAGL0A03586g | Schwarzmüller et al., 2014 |
| 1A5 | CAGL0A04081g | Verified | | Predicted GPI-linked cell wall protein | ORF, Uncharacterized | *NCW2* | CAGL0A04081g | Schwarzmüller et al., 2014 |
| 1A6 | CAGL0A04455g | Verified | *SEF1* | Putative RNA polymerase II transcription factor, involved in regulation of iron acquisition genes; required for growth under iron depletion | ORF, Verified | *SEF1* | CAGL0A04455g | Schwarzmüller et al., 2014 |
| 1A7 | CAGL0B00858g | Verified | *STE50* | Ortholog(s) have SAM domain binding, protein kinase regulator activity | ORF, Uncharacterized | *STE50* | CAGL0B00858g | Schwarzmüller et al., 2014 |
| 1A8 | CAGL0B00968g | Verified | | Ortholog(s) have role in cytogamy, maintenance of protein location in cell cortex, regulation of termination of mating projection growth and integral component of plasma membrane, mating projection tip localization | ORF, Uncharacterized | *FUS1* | CAGL0B00968g | Schwarzmüller et al., 2014 |
| 1A9 | CAGL0B02079g | Wrong Strain | | Ortholog(s) have azole transmembrane transporter activity, role in asperfuranone biosynthetic process, azole transmembrane transport and plasma membrane localization | ORF, Uncharacterized | *AZR1* | CAGL0B02079g | Schwarzmüller et al., 2014 |
| 1A10 | CAGL0B02211g | Verified | *CCH1* | Putative calcium transporter; subunit of a plasma membrane gated channel involved in Ca2+ uptake; N-glycosylated integral membrane protein; required for viability upon prolonged fluconazole stress | ORF, Verified | *CCH1* | CAGL0B02211g | Schwarzmüller et al., 2014 |
| 1A11 | CAGL0B02739g | No Deletion Verified | *STE11* | Mitogen activated protein kinase involved in control of hypertonic stress response, filamentous growth, and virulence | ORF, Verified | *STE11* | CAGL0B02739g | Schwarzmüller et al., 2014 |
| 1A12 | CAGL0B03421g | Verified | | Ortholog(s) have DNA-binding transcription activator activity, RNA polymerase II-specific, RNA polymerase II cis-regulatory region sequence-specific DNA binding activity | ORF, Uncharacterized | *HAP1* | CAGL0B03421g | Schwarzmüller et al., 2014 |
| 1B1 | CAGL0B03487g | No Deletion Verified | *TEF3* | Translation elongation factor eEF3 | ORF, Uncharacterized | *YEF3* | CAGL0B03487g | Schwarzmüller et al., 2014 |
| 1B2 | CAGL0B04521g | Verified | *RAS1* | Ortholog(s) have CTP binding, GDP binding, GTP binding, GTPase activity, UTP binding activity | ORF, Uncharacterized | *RAS1* | CAGL0B04521g | Schwarzmüller et al., 2014 |
| 1B3 | CAGL0C00110g | No Deletion Verified | *EPA6* | Sub-telomerically encoded adhesin with a role in cell adhesion; binds to ligands containing a terminal galactose residue; expressed during murine urinary tract infection, biofilm-upregulated; belongs to adhesin cluster I | ORF, Verified | *FLO1* | CAGL0C00110g | Schwarzmüller et al., 2014 |
| 1B4 | CAGL0C01199g | No Deletion Verified | *UPC2A* | Putative Zn(2)-Cys(6) binuclear cluster transcription factor, involved in transcriptional regulation of ergosterol biosynthesis and uptake | ORF, Verified | *ECM22* | CAGL0C01199g | Schwarzmüller et al., 2014 |
| 1B5 | CAGL0C01463g | Verified | | rtholog(s) have role in fungal-type cell wall organization, glycerol metabolic process, regulation of cell growth, response to salt stress | ORF, Uncharacterized | *TCO89* | CAGL0C01463g | Schwarzmüller et al., 2014 |
| 1B6 | CAGL0C02211g | Verified | *UTR2* | Putative glycoside hydrolase of the Crh family; predicted GPI-anchor | ORF, Verified | *UTR2* | CAGL0C02211g | Schwarzmüller et al., 2014 |
| 1B7 | CAGL0C02277g | Verified | *GLN3* | Putative zinc finger transcription factor with a predicted role in nitrogen catabolite repression | ORF, Verified | *GLN3* | CAGL0C02277g | Schwarzmüller et al., 2014 |
| 1B8 | CAGL0C02343g | Verified | *ARB1* | Ortholog(s) have ATPase activity, role in ribosomal small subunit export from nucleus, ribosome biogenesis and nucleus localization | ORF, Verified | *ARB1* | CAGL0C02343g | Schwarzmüller et al., 2014 |
| 1B9 | CAGL0C03289g | No Deletion Verified | *YBT1* | Putative ABC transporter involved in bile acid transport; gene is upregulated in azole-resistant strain | ORF, Uncharacterized | *YBT1* | CAGL0C03289g | Schwarzmüller et al., 2014 |
| 1B10 | CAGL0C03872g | Verified | *TIR3* | Putative GPI-linked cell wall protein involved in sterol uptake | ORF, Verified | *TIR3* | CAGL0C03872g | Schwarzmüller et al., 2014 |
| 1B11 | CAGL0D01936g | Verified | *SNF7* | Ortholog(s) have role in ATP export, ESCRT III complex assembly, cellular response to anoxia, intralumenal vesicle formation and late endosome to vacuole transport, | ORF, Uncharacterized | *SNF7* | CAGL0D01936g | Schwarzmüller et al., 2014 |
| 1B12 | CAGL0D02530g | Verified | *EGT2* | Putative endoglucanase; glycoside hydrolase; predicted GPI-anchor | ORF, Verified | *EGT2* | CAGL0D02530g | Schwarzmüller et al., 2014 |
| 1C1 | CAGL0D02882g | Verified | *SSK1* | Predicted phosphorelay response regulator activity and role in phosphorelay signal transduction system | ORF, Uncharacterized | *SSK1* | CAGL0D02882g | Schwarzmüller et al., 2014 |
| 1C2 | CAGL0D02904g | Verified | | Ortholog(s) have DNA-binding transcription factor activity, sequence-specific DNA binding activity and role in purine nucleobase catabolic process, regulation of gene expression, regulation of transcription, DNA-templated | ORF, Uncharacterized | *PPR1* | CAGL0D02904g | Schwarzmüller et al., 2014 |
| 1C3 | CAGL0D03674g | Verified | | Ortholog(s) have ATP binding, ATPase activity, ribosome binding, translation termination factor activity and role in poly(A)+ mRNA export from nucleus, regulation of translational termination, ribosomal small subunit biogenesis | ORF, Uncharacterized | *NEW1* | CAGL0D03674g | Schwarzmüller et al., 2014 |
| 1C4 | CAGL0D03850g | Verified | | Ortholog(s) have DNA translocase activity and role in double-strand break repair via nonhomologous end joining, nucleosome disassembly, regulation of transcription, DNA-templated, transcription elongation from RNA polymerase II promoter | ORF, Uncharacterized | *RSC30* | CAGL0D03850g | Schwarzmüller et al., 2014 |
| 1C5 | CAGL0D05258g | No Deletion Verified | | Ortholog(s) have ATP binding, ATPase activity, DNA secondary structure binding, DNA/DNA annealing activity, chromatin binding, double-stranded DNA binding and minor groove of adenine-thymine-rich DNA binding | ORF, Uncharacterized | *SMC2* | CAGL0D05258g | Schwarzmüller et al., 2014 |
| 1C6 | CAGL0D06028g | No Deletion Verified | | Ortholog(s) have GDP-dissociation inhibitor activity, GTPase activator activity, role in mitotic spindle orientation checkpoint, negative regulation of exit from mitosis and Bfa1-Bub2 complex, spindle pole body localization | ORF, Uncharacterized | *BFA1* | CAGL0D06028g | Schwarzmüller et al., 2014 |
| 1C7 | CAGL0D06204g | Verified | | Ortholog(s) have role in cellular response to pH, invasive growth in response to glucose limitation, meiotic cell cycle and penicillin biosynthetic process | ORF, Uncharacterized | *RIM8* | CAGL0D06204g | Schwarzmüller et al., 2014 |
| 1C8 | CAGL0D06512g | Verified | | Putative membrane bound guanine nucleotide exchange factor; gene is upregulated in azole-resistant strain | ORF, Uncharacterized | *CDC25* | CAGL0D06512g | Schwarzmüller et al., 2014 |
| 1C9 | CAGL0E00385g | Verified | | Ortholog(s) have mitochondrion localization | ORF, Uncharacterized | *MDL1* | CAGL0E00385g | Schwarzmüller et al., 2014 |
| 1C10 | CAGL0E01595g | No Deletion Verified | | Putative glycoside hydrolase of the Gas/Phr family; predicted GPI-anchor | ORF, Uncharacterized | *GAS4* | CAGL0E01595g | Schwarzmüller et al., 2014 |
| 1C11 | CAGL0E01771g | Verified | *YPS5* | Putative aspartic protease; predicted GPI-anchor; member of a YPS gene cluster that is required for virulence in mice; gene is downregulated in azole-resistant strain | ORF, Verified | *MCK7* | CAGL0E01771g | Schwarzmüller et al., 2014 |
| 1C12 | CAGL0E03355g | Verified | | Has domain(s) with predicted ABC-type transporter activity, ATP binding, ATPase activity, role in transmembrane transport and membrane localization | ORF, Uncharacterized | *BPT1* | CAGL0E03355g | Schwarzmüller et al., 2014 |
| 1D1 | CAGL0E03476g | Verified | | Ortholog(s) have guanyl-nucleotide exchange factor activity and role in Ras protein signal transduction, regulation of cell cycle, regulation of secretion, traversing start control point of mitotic cell cycle | ORF, Uncharacterized | *CDC25* | CAGL0E03476g | Schwarzmüller et al., 2014 |
| 1D2 | CAGL0E03762g | Verified | | Ortholog(s) have DNA-binding transcription repressor activity, RNA polymerase II-specific activity | ORF, Uncharacterized | *RIM101* | CAGL0E03762g | Schwarzmüller et al., 2014 |
| 1D3 | CAGL0E03982g | Verified | | Ortholog(s) have role in response to metal ion, xenobiotic detoxification by transmembrane export across the plasma membrane and fungal-type vacuole membrane localization | ORF, Uncharacterized | *VMR1* | CAGL0E03982g | Schwarzmüller et al., 2014 |
| 1D4 | CAGL0E04620g | Verified | *PST1* | Ecm33-family protein with a predicted role in cell wall biogenesis and organization; predicted GPI-anchor | ORF, Verified | *PST1* | CAGL0E04620g | Schwarzmüller et al., 2014 |
| 1D5 | CAGL0E05434g | No Deletion Verified | *ZCF8* | Ortholog(s) have DNA binding activity, role in transcription elongation by RNA polymerase II and cell cortex of cell tip localization | ORF, Uncharacterized | *TEA1* | CAGL0E05434g | Schwarzmüller et al., 2014 |
| 1D6 | CAGL0E06248g | Verified | *RCN1* | Regulator of calcineurin; required for cell growth and activation of the calcineurin-Crz1 pathway in the presence of micafungin; activates calcineurin-dependent signaling | ORF, Verified | *RCN1* | CAGL0E06248g | Schwarzmüller et al., 2014 |
| 1D7 | CAGL0E06666g | Verified | *EPA2* | Epithelial adhesion protein; predicted GPI-anchor; belongs to adhesin cluster I | ORF, Verified | *FLO11* | CAGL0E06666g | Schwarzmüller et al., 2014 |
| 1D8 | CAGL0F00495g | No Deletion Verified | | Ortholog(s) have protein kinase activity, protein serine/threonine kinase activity | ORF, Uncharacterized | *TOR1* | CAGL0F00495g | Schwarzmüller et al., 2014 |
| 1D9 | CAGL0F01419g | Verified | *AUS1* | ATP-binding cassette transporter involved in sterol uptake | ORF, Verified | *AUS1* | CAGL0F01419g | Schwarzmüller et al., 2014 |
| 1D10 | CAGL0F01463g | Verified | *TIR1* | Putative GPI-linked cell wall mannoprotein of the Srp1p/Tip1p family | ORF, Verified | *TIR1* | CAGL0F01463g | Schwarzmüller et al., 2014 |
| 1D11 | CAGL0F02079g | Verified | | Ortholog(s) have DNA secondary structure binding, double-stranded DNA binding, minor groove of adenine-thymine-rich DNA binding, topological DNA co-entrapment activity | ORF, Uncharacterized | *SMC1* | CAGL0F02079g | Schwarzmüller et al., 2014 |
| 1D12 | CAGL0F02519g | Verified | *ZCF9* | Has domain(s) with predicted DNA binding, DNA-binding transcription factor activity, RNA polymerase II-specific, zinc ion binding activity and role in DNA-templated transcription, regulation of DNA-templated transcription | ORF, Uncharacterized | *YJL206C* | CAGL0F02519g | Schwarzmüller et al., 2014 |
| 1E1 | CAGL0F02717g | Verified | *PDH1* | Multidrug transporter, predicted plasma membrane ATP-binding cassette (ABC) transporter; regulated by Pdr1p; involved in fluconazole resistance | ORF, Verified | *PDR15* | CAGL0F02717g | Schwarzmüller et al., 2014 |
| 1E2 | CAGL0F03025g | Verified | *ZCF10* | Ortholog(s) have DNA-binding transcription factor activity, RNA polymerase II-specific activity, role in aromatic amino acid family catabolic process, positive regulation of transcription by RNA polymerase II and nucleus localization | ORF, Uncharacterized | *ARO80* | CAGL0F03025g | Schwarzmüller et al., 2014 |
| 1E3 | CAGL0F03883g | Verified | *GAS4* | Putative glycoside hydrolase of the Gas/Phr family; predicted GPI-anchor | ORF, Verified | *GAS3* | CAGL0F03883g | Schwarzmüller et al., 2014 |
| 1E4 | CAGL0F04081g | Verified | *TEC2* | Has domain(s) with predicted DNA-binding transcription factor activity and role in regulation of DNA-templated transcription | ORF, Uncharacterized | *TEC1* | CAGL0F04081g | Schwarzmüller et al., 2014 |
| 1E5 | CAGL0F05357g | No Deletion Verified | *ZCF11* | Ortholog(s) have DNA-binding transcription factor binding, DNA-binding transcription repressor activity, RNA polymerase II-specific, RNA polymerase II cis-regulatory region sequence-specific DNA binding activity | ORF, Uncharacterized | *UME6* | CAGL0F05357g | Schwarzmüller et al., 2014 |
| 1E6 | CAGL0F05995g | Verified | *MSN2* | Putative transcription factor similar to S. cerevisiae Msn2p; involved in response to oxidative stress | ORF, Verified | *MSN2* | CAGL0F05995g | Schwarzmüller et al., 2014 |
| 1E7 | CAGL0F06545g | Verified | | Ortholog(s) have role in ascospore formation and cell division site, endocytic vesicle, exocytic vesicle, fungal-type vacuole lumen, growing cell tip, plasma membrane, plasma membrane of cell tip localization | ORF, Uncharacterized | *RIM9* | CAGL0F06545g | Schwarzmüller et al., 2014 |
| 1E8 | CAGL0F06655g | Verified | | Ortholog(s) have role in establishment or maintenance of actin cytoskeleton polarity, regulation of cell growth and TORC2 complex, plasma membrane localization | ORF, Uncharacterized | *AVO2* | CAGL0F06655g | Schwarzmüller et al., 2014 |
| 1E9 | CAGL0F06677g | Verified | *GPA1* | Ortholog(s) have G-protein beta-subunit binding, GTPase activity | ORF, Uncharacterized | *GPA1* | CAGL0F06677g | Schwarzmüller et al., 2014 |
| 1E10 | CAGL0F07579g | Verified | *CWP1.2* | Serine-type protease of the cell wall; GPI-linked cell wall protein that appears to be involved in virulence | ORF, Verified | *CWP1* | CAGL0F07579g | Schwarzmüller et al., 2014 |
| 1E11 | CAGL0F07601g | Verified | *CWP1.1* | GPI-linked cell wall protein | ORF, Verified | *CPW2* | CAGL0F07601g | Schwarzmüller et al., 2014 |
| 1E12 | CAGL0F07755g | Verified | *CEP3* | Centromere binding factor 3b; inner kinetochore protein | ORF, Verified | *CEP3* | CAGL0F07755g | Schwarzmüller et al., 2014 |
| 1F1 | CAGL0F07865g | Verified | *UPC2B* | Putative Zn(2)-Cys(6) binuclear cluster transcriptional regulator of ergosterol biosynthesis | ORF, Uncharacterized | *UPC2* | CAGL0F07865g | Schwarzmüller et al., 2014 |
| 1F2 | CAGL0F07909g | Verified | *ZFC15* | Putative zinc finger transcription factor; contains predicted zinc-binding and DNA-binding domains; point mutants identified among echinocandin-resistant clinical isolates | ORF, Verified | *TBS2* | CAGL0F07909g | Schwarzmüller et al., 2014 |
| 1F3 | CAGL0F08833g | Verified | | Putative adhesin-like protein | ORF, Verified | *MSB2* | CAGL0F08833g | Schwarzmüller et al., 2014 |
| 1F4 | CAGL0F09053g | Verified | | Ortholog(s) have carbohydrate binding, mannosyl-oligosaccharide 1,2-alpha-mannosidase activity | ORF, Uncharacterized | *MNL1* | CAGL0F09053g | Schwarzmüller et al., 2014 |
| 1F5 | CAGL0F09229g | Verified | *TOG1* | RNA polymerase II transcription factor, involved in oxidative stress resistance and virulence | ORF, Verified | *TOG1* | CAGL0F09229g | Schwarzmüller et al., 2014 |
| 1F6 | CAGL0G00242g | Verified | *YOR1* | Putative ABC transporter involved in multidrug efflux; mutation suppresses TOR and calcineurin signaling; gene is upregulated in azole-resistant strain | ORF, Uncharacterized | *YOR1* | CAGL0G00242g | Schwarzmüller et al., 2014 |
| 1F7 | CAGL0G01056g | Verified | *GAS3* | Putative glycoside hydrolase of the Gas/Phr family; predicted GPI-anchor | ORF, Uncharacterized | *GAS2* | CAGL0G01056g | Schwarzmüller et al., 2014 |
| 1F8 | CAGL0G01320g | Verified | | Ortholog(s) have protein tyrosine/serine/threonine phosphatase activity | ORF, Uncharacterized | *MSG5* | CAGL0G01320g | Schwarzmüller et al., 2014 |
| 1F9 | CAGL0G01386g | Verified | | Ortholog(s) have role in TOR signaling, actin cytoskeleton organization, actin filament bundle assembly, eisosome assembly, endosomal transport and establishment or maintenance of actin cytoskeleton polarity, protein localization to plasma membrane regulation of cell growth | ORF, Uncharacterized | *SLM2* | CAGL0G01386g | Schwarzmüller et al., 2014 |
| 1F10 | CAGL0G02409g | Verified | *SRP40* | Ortholog(s) have nucleolus localization | ORF, Uncharacterized | *SRP40* | CAGL0G02409g | Schwarzmüller et al., 2014 |
| 1F11 | CAGL0G02827g | Verified | | Ortholog(s) have phosphatidylinositol-4,5-bisphosphate binding, sphingolipid binding activity | ORF, Uncharacterized | *SLM1* | CAGL0G02827g | Schwarzmüller et al., 2014 |
| 1F12 | CAGL0G03597g | Verified | *SHO1* | S. cerevisiae ortholog SHO1 has role in establishment of cell polarity, osmosensory signaling pathway, signal transduction involved in filamentous growth, cellular response to heat; osmosensitivity of CBS138 strain not seen in other strains | ORF, Uncharacterized | *SHO1* | CAGL0G03597g | Schwarzmüller et al., 2014 |
| 1G1 | CAGL0G04125g | Verified | | Protein with similarity to S. cerevisiae Sag1 agglutinin, involved in cell adhesion; predicted GPI-anchor | ORF, Uncharacterized | *SAG1* | CAGL0G04125g | Schwarzmüller et al., 2014 |
| 1G2 | CAGL0G05896g | Verified | | Putative adhesin-like protein | ORF, Uncharacterized | *DSE2* | CAGL0G05896g | Schwarzmüller et al., 2014 |
| 1G3 | CAGL0G06314g | No Deletion Verified | | Ortholog(s) have role in cellular response to alkaline pH, cellular response to salt stress, post-translational protein targeting to membrane, translocation and endoplasmic reticulum localization | ORF, Uncharacterized | *FRT2* | CAGL0G06314g | Schwarzmüller et al., 2014 |
| 1G4 | CAGL0G07667g | Verified | | Ortholog(s) have cholesterol binding, fatty acid binding, magnesium ion binding, sterol binding activity, role in fatty acid transport, sterol transport and extracellular region localization | ORF, Verified | *PRY1* | CAGL0G07667g | Schwarzmüller et al., 2014 |

| | | | | | | | | |
|---|---|---|---|---|---|---|---|---|
| **1G5** | CAGL0G08041g | No Deletion Verified | | Ortholog(s) have iron ion binding activity and role in positive regulation of translation, ribosomal large subunit biogenesis, ribosomal subunit export from nucleus, ribosome disassembly, translational initiation, translational termination | ORF, Uncharacterized | *RLI1* | CAGL0G08041g | Schwarzmüller et al., 2014 |
| **1G6** | CAGL0G08844g | Verified | *ZCF17* | Has domain(s) with predicted DNA binding, DNA-binding transcription factor activity, RNA polymerase II-specific, zinc ion binding activity and role in DNA-templated transcription, regulation of DNA-templated transcription | ORF, Uncharacterized | *ASG1* | CAGL0G08844g | Schwarzmüller et al., 2014 |
| **1G8** | CAGL0G09449g | Verified | *CRH1* | Putative glycoside hydrolase; predicted GPI-anchor | ORF, Verified | *CRH1* | CAGL0G09449g | Schwarzmüller et al., 2014 |
| **1G9** | CAGL0G09757g | Verified | *ZCF18* | Has domain(s) with predicted DNA binding, DNA-binding transcription factor activity, RNA polymerase II-specific, zinc ion binding activity and role in DNA-templated transcription, regulation of DNA-templated transcription | ORF, Uncharacterized | *YLR278C* | CAGL0G09757g | Schwarzmüller et al., 2014 |
| **1G10** | CAGL0H00220g | No Deletion Verified | | Has domain(s) with predicted role in attachment of GPI anchor to protein and GPI-anchor transamidase complex localization | ORF, Uncharacterized | *GAB1* | CAGL0H00220g | Schwarzmüller et al., 2014 |
| **1G11** | CAGL0H00396g | Verified | *ZCF19* | Ortholog(s) have DNA-binding transcription activator activity, RNA polymerase II-specific, DNA-binding transcription repressor activity, RNA polymerase II-specific activity | ORF, Uncharacterized | *LEU3* | CAGL0H00396g | Schwarzmüller et al., 2014 |
| **1G12** | CAGL0H01507g | No Deletion Verified | *ZCF20* | Ortholog(s) have role in chromatin remodeling, nucleosome disassembly, regulation of DNA-templated transcription, regulation of nuclear cell cycle DNA replication, transcription elongation by RNA polymerase II | ORF, Uncharacterized | *RSC3* | CAGL0H01507g | Schwarzmüller et al., 2014 |
| **1H1** | CAGL0H01661g | No Deletion Verified | | Ecm33-family protein with a predicted role in cell wall biogenesis and organization; predicted GPI-anchor | ORF, Uncharacterized | *SPS2* | CAGL0H01661g | Schwarzmüller et al., 2014 |
| **1H2** | CAGL0H01683g | Verified | *ZCF21* | Has domain(s) with predicted DNA-binding transcription factor activity, RNA polymerase II-specific, zinc ion binding activity and role in regulation of DNA-templated transcription | ORF, Uncharacterized | *URC2* | CAGL0H01683g | Schwarzmüller et al., 2014 |
| **1H3** | CAGL0H02563g | Verified | | Predicted GPI-linked protein | ORF, Uncharacterized | *HOR7* | CAGL0H02563g | Schwarzmüller et al., 2014 |
| **1H4** | CAGL0H03553g | No Deletion Verified | | Ortholog(s) have protein kinase activity, protein serine/threonine kinase activity, protein tyrosine kinase activity and role in homologous chromosome segregation, regulation of DNA repair, regulation of linear element assembly | ORF, Uncharacterized | *HRR25* | CAGL0H03553g | Schwarzmüller et al., 2014 |
| **1H5** | CAGL0H03861g | Verified | | Ortholog(s) have chromatin binding, mRNA binding, single-stranded telomeric DNA binding activity and role in nuclear mRNA surveillance, poly(A)+ mRNA export from nucleus, telomere maintenance | ORF, Uncharacterized | *GBP2* | CAGL0H03861g | Schwarzmüller et al., 2014 |
| **1H6** | CAGL0H04367g | Verified | *WAR1* | Ortholog(s) have DNA-binding transcription factor activity, sequence-specific DNA binding activity, role in positive regulation of transcription by RNA polymerase II and nucleus localization | ORF, Uncharacterized | *WAR1* | CAGL0H04367g | Schwarzmüller et al., 2014 |
| **1H7** | CAGL0H05621g | Verified | *RLM1* | Putative transcription factor with a predicted role in cell wall integrity | ORF, Uncharacterized | *RLM1* | CAGL0H05621g | Schwarzmüller et al., 2014 |
| **1H8** | CAGL0H06413g | Verified | *SSR1* | Putative GPI-linked cell wall protein | ORF, Verified | *CCW14* | CAGL0H06413g | Schwarzmüller et al., 2014 |
| **1H9** | CAGL0H06875g | Verified | | Ortholog(s) have transcription coregulator activity, role in regulation of arginine metabolic process and nucleus localization | ORF, Uncharacterized | *ARG81* | CAGL0H06875g | Schwarzmüller et al., 2014 |
| **1H10** | CAGL0H08129g | No Deletion Verified | | Ortholog(s) have role in protein processing, pseudohyphal growth and plasma membrane localization | ORF, Uncharacterized | *DFG16* | CAGL0H08129g | Schwarzmüller et al., 2014 |
| **1H11** | CAGL0H08624g | Verified | *MCM16* | Ortholog(s) have role in chromosome segregation, establishment of mitotic sister chromatid cohesion and kinetochore localization | ORF, Uncharacterized | *MCM16* | CAGL0H08624g | Schwarzmüller et al., 2014 |
| **2A1** | CAGL0H09592g | No Deletion Verified | | Putative GPI-linked cell wall protein | ORF, Uncharacterized | *TIR1* | CAGL0H09592g | Schwarzmüller et al., 2014 |
| **2A2** | CAGL0I02552g | No Deletion Verified | | Predicted sequence-specific DNA binding transcription factor, negative regulator of azole resistance; acts as transcriptional repressor of ATP-binding Cassette (ABC) transporter genes | ORF, Verified | *STB5* | CAGL0I02552g | Schwarzmüller et al., 2014 |
| **2A3** | CAGL0I03498g | Verified | | Ortholog(s) have MAP kinase kinase activity | ORF, Uncharacterized | *STE7* | CAGL0I03498g | Schwarzmüller et al., 2014 |
| **2A4** | CAGL0I04092g | Verified | | Ortholog(s) have role in ascospore wall assembly | ORF, Uncharacterized | *SHE10* | CAGL0I04092g | Schwarzmüller et al., 2014 |
| **2A5** | CAGL0I04862g | Verified | *SNQ2* | Predicted plasma membrane ATP-binding cassette (ABC) transporter, putative transporter involved in multidrug resistance; involved in Pdr1p-mediated azole resistance | ORF, Verified | *SNQ2* | CAGL0I04862g | Schwarzmüller et al., 2014 |
| **2A6** | CAGL0I04950g | No Deletion Verified | | Ortholog(s) have molecular adaptor activity, protein-membrane adaptor activity | ORF, Uncharacterized | *TSC11* | CAGL0I04950g | Schwarzmüller et al., 2014 |
| **2A7** | CAGL0I05236g | Verified | *BCY1* | cAMP dependent protein kinase, regulatory subunit | ORF, Uncharacterized | *BCY1* | CAGL0I05236g | Schwarzmüller et al., 2014 |
| **2A8** | CAGL0I05896g | No Deletion Verified | *YAK1* | Putative serine-threonine protein kinase, involved in biofilm formation, required for expression of adhesin genes EPA6 and EPA7 | ORF, Verified | *YAK1* | CAGL0I05896g | Schwarzmüller et al., 2014 |
| **2A9** | CAGL0I06138g | Verified | | Ortholog(s) have cyclin-dependent protein serine/threonine kinase inhibitor activity and role in chemotropism, maintenance of protein location in nucleus, mitotic cell cycle G1 arrest in response to pheromone | ORF, Uncharacterized | *FAR1* | CAGL0I06138g | Schwarzmüller et al., 2014 |
| **2A10** | CAGL0I07293g | No Deletion Verified | | Adhesin-like cell wall protein; predicted GPI-anchor; belongs to adhesin cluster V | ORF, Uncharacterized | | CAGL0I07293g | Schwarzmüller et al., 2014 |
| **2A11** | CAGL0I07755g | Verified | *HAL9* | Ortholog(s) have role in positive regulation of transcription by RNA polymerase II, response to salt stress | ORF, Uncharacterized | *HAL9* | CAGL0I07755g | Schwarzmüller et al., 2014 |
| **2A12** | CAGL0I08019g | No Deletion Verified | *ROA1* | Ortholog(s) have fungal-type vacuole membrane localization | ORF, Uncharacterized | *YOL075C* | CAGL0I08019g | Schwarzmüller et al., 2014 |
| **2B1** | CAGL0I08195g | Verified | | Ortholog(s) have GTP binding, adenylate cyclase activator activity | ORF, Uncharacterized | *GPA2* | CAGL0I08195g | Schwarzmüller et al., 2014 |
| **2B2** | CAGL0I08503g | Verified | | Ortholog(s) have phosphoadenylyl-sulfate reductase (thioredoxin) activity and role in sulfate assimilation, phosphoadenylyl sulfate reduction by phosphoadenylyl-sulfate reductase (thioredoxin) | ORF, Uncharacterized | *MET16* | CAGL0I08503g | Schwarzmüller et al., 2014 |
| **2B3** | CAGL0I09416g | Verified | | Ortholog(s) have TORC2 complex localization | ORF, Uncharacterized | *BIT2* | CAGL0I09416g | Schwarzmüller et al., 2014 |
| **2B4** | CAGL0J02530g | No Deletion Verified | | Putative adhesion protein; predicted GPI-anchor; belongs to adhesin cluster VI | ORF, Uncharacterized | | CAGL0J02530g | Schwarzmüller et al., 2014 |
| **2B5** | CAGL0J04290g | Verified | | Ortholog(s) have MAP kinase activity, role in pheromone response MAPK cascade and nucleus localization | ORF, Uncharacterized | *FUS3* | CAGL0J04290g | Schwarzmüller et al., 2014 |
| **2B6** | CAGL0J04752g | No Deletion Verified | | Ortholog(s) have guanyl-nucleotide exchange factor activity and role in fungal-type cell wall organization, positive regulation of mitotic actomyosin contractile ring assembly, signal transduction | ORF, Verified | *TUS1* | CAGL0J04752g | Schwarzmüller et al., 2014 |
| **2B7** | CAGL0J07150g | No Deletion Verified | | Ortholog(s) have DNA-binding transcription activator activity, RNA polymerase II-specific, RNA polymerase II cis-regulatory region sequence-specific DNA binding activity | ORF, Uncharacterized | *PIP2* | CAGL0J07150g | Schwarzmüller et al., 2014 |
| **2B8** | CAGL0J08206g | Verified | | Ortholog(s) have F-bar domain binding, actin filament binding, profilin binding activity | ORF, Uncharacterized | *BNI1* | CAGL0J08206g | Schwarzmüller et al., 2014 |
| **2B9** | CAGL0J08613g | Verified | | Ortholog(s) have calcium channel activity, calcium-activated cation channel activity, potassium channel activity, sodium channel activity, voltage-gated monoatomic ion channel activity | ORF, Uncharacterized | *YVC1* | CAGL0J08613g | Schwarzmüller et al., 2014 |
| **2B10** | CAGL0J11308g | Verified | | Ortholog(s) have protein serine/threonine kinase activity, role in negative regulation of endocytosis, regulation of sphingolipid biosynthetic process and Golgi apparatus, cytoplasm, plasma membrane localization | ORF, Uncharacterized | *NPR1* | CAGL0J11308g | Schwarzmüller et al., 2014 |
| **2B11** | CAGL0J11462g | Wrong Strain | | Predicted GPI-linked cell wall protein | ORF, Uncharacterized | *YNL190W* | CAGL0J11462g | Schwarzmüller et al., 2014 |
| **2B12** | CAGL0J11748g | Verified | *PLB2* | Putative phospholipase B; predicted GPI-anchor | ORF, Verified | *PLB2* | CAGL0J11748g | Schwarzmüller et al., 2014 |
| **2C1** | CAGL0J11770g | Verified | *PLB1* | Putative phospholipase B; predicted GPI-anchor | ORF, Verified | *PLB1* | CAGL0J11770g | Schwarzmüller et al., 2014 |
| **2C2** | CAGL0K00363g | Verified | | Ortholog(s) have ABC-type oligopeptide transporter activity, ABC-type peptide transporter activity and role in conjugation with cellular fusion, peptide pheromone export, peptide pheromone export by transmembrane transport | ORF, Uncharacterized | *STE6* | CAGL0K00363g | Schwarzmüller et al., 2014 |
| **2C3** | CAGL0K01331g | Verified | | Ortholog(s) have phosphoprotein phosphatase activity, protein serine/threonine phosphatase activity | ORF, Uncharacterized | *SIT4* | CAGL0K01331g | Schwarzmüller et al., 2014 |
| **2C4** | CAGL0K01507g | Verified | | Ortholog(s) have G protein-coupled receptor activity, glucose binding activity | ORF, Uncharacterized | *GPR1* | CAGL0K01507g | Schwarzmüller et al., 2014 |
| **2C5** | CAGL0K02673g | Verified | *STE20* | Putative signal transducing kinase of the PAK (p21-activated kinase) family; involved in maintaining cell wall integrity, osmotic stress response, and virulence | ORF, Verified | *STE20* | CAGL0K02673g | Schwarzmüller et al., 2014 |
| **2C6** | CAGL0K05423g | Verified | | Ortholog(s) have role in negative regulation of TORC1 signaling | ORF, Uncharacterized | *TIP41* | CAGL0K05423g | Schwarzmüller et al., 2014 |
| **2C7** | CAGL0K06479g | Verified | | Ortholog(s) have role in cell wall integrity MAPK cascade | ORF, Uncharacterized | *PKH3* | CAGL0K06479g | Schwarzmüller et al., 2014 |
| **2C8** | CAGL0K06985g | Verified | | Ortholog(s) have DNA-binding transcription repressor activity, RNA polymerase II-specific, sequence-specific DNA binding activity | ORF, Uncharacterized | *ERT1* | CAGL0K06985g | Schwarzmüller et al., 2014 |
| **2C9** | CAGL0K09790g | Wrong Strain | | Ortholog(s) have actin binding, adenylate cyclase binding activity | ORF, Uncharacterized | *SRV2* | CAGL0K09790g | Schwarzmüller et al., 2014 |
| **2C10** | CAGL0K10164g | Verified | | Predicted GPI-linked protein; putative adhesin-like protein | ORF, Uncharacterized | *SED1* | CAGL0K10164g | Schwarzmüller et al., 2014 |
| **2C11** | CAGL0K10472g | No Deletion Verified | | Ortholog(s) have role in protein unfolding, regulation of translational elongation and cytosolic ribosome localization | ORF, Uncharacterized | *GCN20* | CAGL0K10472g | Schwarzmüller et al., 2014 |
| **2C12** | CAGL0K10868g | Verified | *CTA1* | Putative catalase A; gene is downregulated in azole-resistant strain; regulated by oxidative stress and glucose starvation; protein abundance increased in ace2 mutant cells | ORF, Verified | *CTA1* | CAGL0K10868g | Schwarzmüller et al., 2014 |
| **2D1** | CAGL0K11902g | Verified | | Ortholog(s) have RNA polymerase II cis-regulatory region sequence-specific DNA binding activity | ORF, Uncharacterized | *LYS14* | CAGL0K11902g | Schwarzmüller et al., 2014 |
| **2D2** | CAGL0K12078g | Verified | | Ortholog(s) have DNA-binding transcription repressor activity, RNA polymerase II-specific, RNA polymerase II cis-regulatory region sequence-specific DNA binding, RNA polymerase II-specific DNA-binding transcription factor binding activity | ORF, Uncharacterized | *NRG1* | CAGL0K12078g | Schwarzmüller et al., 2014 |
| **2D3** | CAGL0K12430g | Verified | *STE2* | Ortholog(s) have mating-type P-factor pheromone receptor activity, mating-type alpha-factor pheromone receptor activity | ORF, Uncharacterized | *STE2* | CAGL0K12430g | Schwarzmüller et al., 2014 |
| **2D4** | CAGL0K12562g | Verified | *RIM15* | Ortholog(s) have protein kinase activity, protein serine/threonine kinase activity | ORF, Uncharacterized | *RIM15* | CAGL0K12562g | Schwarzmüller et al., 2014 |
| **2D5** | CAGL0L00605g | Verified | *CNB1* | Regulatory subunit of calcineurin, calcium/calmodulin-dependent Ser/Thr-specific protein phosphatase; regulates stress-responding transcription factor Crz1p; involved in thermotolerance, response to ER stress, cell wall integrity, virulence | ORF, Verified | *CNB1* | CAGL0L00605g | Schwarzmüller et al., 2014 |
| **2D6** | CAGL0L01727g | Verified | *DCW1* | Putative glycoside hydrolase with GPI anchoring; involved in cell wall biogenesis; null mutant cannot grow in the presence of calcineurin inhibitors and shows increased expression of RCN2 | ORF, Verified | *DCW1* | CAGL0L01727g | Schwarzmüller et al., 2014 |
| **2D7** | CAGL0L01771g | Verified | *EFG2* | Ortholog(s) have DNA-binding transcription factor activity, role in positive regulation of pseudohyphal growth, positive regulation of transcription by RNA polymerase II and nucleus localization | ORF, Uncharacterized | *PHD1* | CAGL0L01771g | Schwarzmüller et al., 2014 |
| **2D8** | CAGL0L01903g | Verified | *RGT1* | Ortholog(s) have DNA binding, DNA-binding transcription activator activity, RNA polymerase II-specific, RNA polymerase II cis-regulatory region sequence-specific DNA binding activity | ORF, Uncharacterized | *RGT1* | CAGL0L01903g | Schwarzmüller et al., 2014 |
| **2D9** | CAGL0L02761g | Verified | | Ortholog(s) have G-protein alpha-subunit binding, G-protein gamma-subunit binding, protein kinase binding, scaffold protein binding, small GTPase binding activity | ORF, Uncharacterized | *STE4* | CAGL0L02761g | Schwarzmüller et al., 2014 |
| **2D10** | CAGL0L03377g | Verified | | Ortholog(s) have DNA-binding transcription activator activity, RNA polymerase II-specific, RNA polymerase II cis-regulatory region sequence-specific DNA binding activity | ORF, Uncharacterized | *SIP4* | CAGL0L03377g | Schwarzmüller et al., 2014 |
| **2D11** | CAGL0L03674g | Verified | | Ortholog(s) have DNA-binding transcription factor activity and role in regulation of transcription by RNA polymerase II | ORF, Uncharacterized | *GSM1* | CAGL0L03674g | Schwarzmüller et al., 2014 |
| **2D12** | CAGL0L04400g | Verified | | Ortholog(s) have DNA-binding transcription factor activity, RNA polymerase II-specific, RNA polymerase II cis-regulatory region sequence-specific DNA binding activity | ORF, Uncharacterized | *YRR1* | CAGL0L04400g | Schwarzmüller et al., 2014 |
| **2E1** | CAGL0L04576g | Verified | | Ortholog(s) have DNA-binding transcription factor activity, RNA polymerase II-specific activity and role in regulation of transcription by RNA polymerase II | ORF, Uncharacterized | *YRM1* | CAGL0L04576g | Schwarzmüller et al., 2014 |
| **2E3** | CAGL0L06402g | No Deletion Verified | *YCF1* | Ortholog(s) have ABC-type glutathione S-conjugate transporter activity, ABC-type phytochelatin transporter activity, bilirubin transmembrane transporter activity | ORF, Uncharacterized | *YCF1* | CAGL0L06402g | Schwarzmüller et al., 2014 |
| **2E4** | CAGL0L07656g | Verified | *GLO1* | Ortholog(s) have lactoylglutathione lyase activity, zinc ion binding activity and role in glutathione metabolic process, methylglyoxal catabolic process to D-lactate via S-lactoyl-glutathione | ORF, Uncharacterized | *GLO1* | CAGL0L07656g | Schwarzmüller et al., 2014 |
| **2E5** | CAGL0L07744g | Verified | *ADP1* | Has domain(s) with predicted ABC-type transporter activity, ATP binding, ATPase activity and membrane localization | ORF, Uncharacterized | *ADP1* | CAGL0L07744g | Schwarzmüller et al., 2014 |
| **2E6** | CAGL0L09691g | Verified | | Ortholog(s) have DNA-binding transcription activator activity, RNA polymerase II-specific, RNA polymerase II cis-regulatory region sequence-specific DNA binding activity | ORF, Uncharacterized | *PUT3* | CAGL0L09691g | Schwarzmüller et al., 2014 |
| **2E7** | CAGL0L10956g | Verified | | Ortholog(s) have role in invasive growth in response to glucose limitation, protein processing, sporulation resulting in formation of a cellular spore and peroxisome localization | ORF, Uncharacterized | *RIM20* | CAGL0L10956g | Schwarzmüller et al., 2014 |
| **2E8** | CAGL0L11110g | Verified | *CNA1* | Catalytic subunit of calcineurin, calcium/calmodulin-dependent Ser/Thr-specific protein phosphatase; regulates stress-responding transcription factor Crz1p; involved in thermotolerance, response to ER stress, cell wall integrity, virulence | ORF, Verified | *CMP2* | CAGL0L11110g | Schwarzmüller et al., 2014 |
| **2E9** | CAGL0L12056g | Verified | *BMH1(A)* | 14-3-3 protein | ORF, Verified | *BMH1* | CAGL0L12056g | Schwarzmüller et al., 2014 |
| **2E10** | CAGL0L12782g | Verified | *DIG1* | Ortholog(s) have transcription corepressor activity and role in negative regulation of invasive growth in response to glucose limitation, negative regulation of pseudohyphal growth, negative regulation of transcription by RNA polymerase II | ORF, Uncharacterized | *DIG1* | CAGL0L12782g | Schwarzmüller et al., 2014 |
| **2E11** | CAGL0M01254g | No Deletion Verified | *STE12* | Putative transcription factor, required for filamentous growth induced by nitrogen starvation and for virulence; functionally complements S. cerevisiae ste12 mutant | ORF, Verified | *STE12* | CAGL0M01254g | Schwarzmüller et al., 2014 |
| **2E12** | CAGL0M01760g | Verified | *CDR1* | Multidrug transporter of ATP-binding cassette (ABC) superfamily, involved in resistance to azoles; expression regulated by Pdr1p; increased abundance in azole resistant strains; expression increased by loss of the mitochondrial genome | ORF, Verified | *PDR5* | CAGL0M01760g | Schwarzmüller et al., 2014 |
| **2F1** | CAGL0M01826g | Verified | *ECM33* | Ecm33-family protein with a predicted role in cell wall biogenesis and organization; predicted GPI-anchor | ORF, Verified | *ECM33* | CAGL0M01826g | Schwarzmüller et al., 2014 |

| | | | | | | | | |
|---|---|---|---|---|---|---|---|---|
| 2F2 | CAGL0M02387g | Verified | PXA1 | Ortholog(s) have ATPase-coupled transmembrane transporter activity, role in ATP transport, fatty acid transport and peroxisomal membrane, peroxisome localization | ORF, Uncharacterized | PXA1 | CAGL0M02387g | Schwarzmüller et al., 2014 |
| 2F3 | CAGL0M02651g | Verified | | Ortholog(s) have DNA-binding transcription activator activity, RNA polymerase II-specific, RNA polymerase II cis-regulatory region sequence-specific DNA binding activity | ORF, Uncharacterized | RDS2 | CAGL0M02651g | Schwarzmüller et al., 2014 |
| 2F4 | CAGL0M03025g | Verified | | Ortholog(s) have DNA-binding transcription activator activity, RNA polymerase II-specific, RNA polymerase II cis-regulatory region sequence-specific DNA binding activity | ORF, Uncharacterized | CAT8 | CAGL0M03025g | Schwarzmüller et al., 2014 |
| 2F5 | CAGL0M03597g | Verified | MID1 | Putative calcium transporter; putative regulatory subunit of a plasma membrane gated channel involved in Ca2+ uptake; required for viability upon prolonged fluconazole stress | ORF, Verified | MID1 | CAGL0M03597g | Schwarzmüller et al., 2014 |
| 2F6 | CAGL0M03663g | Verified | | Ortholog(s) have role in ascospore formation, cellular response to alkaline pH, fungal-type cell wall biogenesis, invasive growth in response to glucose limitation, regulation of vacuole organization and plasma membrane localization | ORF, Uncharacterized | RIM21 | CAGL0M03663g | Schwarzmüller et al., 2014 |
| 2F7 | CAGL0M03773g | Verified | | Predicted GPI-linked adhesin-like protein | ORF, Uncharacterized | TOS6 | CAGL0M03773g | Schwarzmüller et al., 2014 |
| 2F8 | CAGL0M04169g | Verified | KRE1 | Putative cell wall protein with similarity to S. cerevisiae Kre1p; predicted role in cell wall biogenesis and organization; predicted GPI-anchor | ORF, Uncharacterized | KRE1 | CAGL0M04169g | Schwarzmüller et al., 2014 |
| 2F9 | CAGL0M04191g | Verified | YPS1 | Yapsin family aspartic protease; predicted GPI-anchor; complements cell wall defect phenotypes of S. cerevisiae yps1 mutant; regulation of pH homeostasis under acid conditions; induced by high temperature, Slt1- and Crz1p-dependent | ORF, Verified | YPS1 | CAGL0M04191g | Schwarzmüller et al., 2014 |
| 2F10 | CAGL0M04565g | Verified | | Ortholog(s) have glucan endo-1,3-beta-D-glucosidase activity | ORF, Uncharacterized | ACF2 | CAGL0M04565g | Schwarzmüller et al., 2014 |
| 2F11 | CAGL0M05049g | No Deletion Verified | | Putative glycoside hydrolase; predicted GPI-anchor | ORF, Verified | DFG5 | CAGL0M05049g | Schwarzmüller et al., 2014 |
| 2F12 | CAGL0M05357g | No Deletion Verified | | Ortholog(s) have molecular adaptor activity, phosphatidylinositol-3-phosphate binding, protein-macromolecule adaptor activity | ORF, Uncharacterized | BEM1 | CAGL0M05357g | Schwarzmüller et al., 2014 |
| 2G1 | CAGL0M05907g | Verified | | Ortholog(s) have role in negative regulation of transcription by RNA polymerase II | ORF, Uncharacterized | OAF3 | CAGL0M05907g | Schwarzmüller et al., 2014 |
| 2G2 | CAGL0M06831g | Verified | CRZ1 | Transcription factor; downstream component of the calcineurin signaling pathway | ORF, Verified | CRZ1 | CAGL0M06831g | Schwarzmüller et al., 2014 |
| 2G3 | CAGL0M07425g | No Deletion Verified | | Has domain(s) with predicted role in regulation of signal transduction | ORF, Uncharacterized | TAP42 | CAGL0M07425g | Schwarzmüller et al., 2014 |
| 2G4 | CAGL0M07634g | No Deletion Verified | EFG1 | Transcription factor involved in control of biofilm formation | ORF, Verified | SOK2 | CAGL0M07634g | Schwarzmüller et al., 2014 |
| 2G5 | CAGL0M08184g | Verified | STE3 | Putative a-factor pheromone receptor | ORF, Uncharacterized | STE3 | CAGL0M08184g | Schwarzmüller et al., 2014 |
| 2G6 | CAGL0M08404g | No Deletion Verified | | Ortholog(s) have cAMP-dependent protein kinase activity | ORF, Uncharacterized | TPK3 | CAGL0M08404g | Schwarzmüller et al., 2014 |
| 2G7 | CAGL0M08756g | No Deletion Verified | | Putative exo-1,3-beta-glucanase; predicted GPI-anchor | ORF, Verified | EXG2 | CAGL0M08756g | Schwarzmüller et al., 2014 |
| 2G8 | CAGL0M09207g | Verified | | Ortholog(s) have G-protein beta-subunit binding activity and role in pheromone-dependent signal transduction involved in conjugation with cellular fusion | ORF, Uncharacterized | STE18 | CAGL0M09207g | Schwarzmüller et al., 2014 |
| 2G9 | CAGL0M09669g | Verified | | Ortholog(s) have cysteine-type endopeptidase activity and role in protein processing | ORF, Uncharacterized | RIM13 | CAGL0M09669g | Schwarzmüller et al., 2014 |
| 2G10 | CAGL0M10791g | Verified | No Information | | | | | Schwarzmüller et al., 2014 |
| 2G11 | CAGL0M11440g | Verified | | Ortholog(s) have DNA-binding transcription factor activity and role in amino acid catabolic process, positive regulation of DNA-templated transcription | ORF, Uncharacterized | CHA4 | CAGL0M11440g | Schwarzmüller et al., 2014 |
| 2G12 | CAGL0M11726g | Verified | | Putative GPI-linked cell wall adhesin-like protein | ORF, Uncharacterized | CCW12 | CAGL0M11726g | Schwarzmüller et al., 2014 |
| 2H1 | CAGL0M11748g | No Deletion Verified | HOG1 | Ortholog(s) have MAP kinase activity, RNA polymerase II CTD heptapeptide repeat kinase activity, calmodulin binding, chromatin binding activity | ORF, Verified | HOG1 | CAGL0M11748g | Schwarzmüller et al., 2014 |
| 2H2 | CAGL0M12298g | Verified | | Ortholog(s) have DNA-binding transcription activator activity, RNA polymerase II-specific, RNA polymerase II cis-regulatory region sequence-specific DNA binding activity | ORF, Uncharacterized | OAF1 | CAGL0M12298g | Schwarzmüller et al., 2014 |
| 2H3 | CAGL0M13739g | Verified | ATM1 | Ortholog(s) have ATPase-coupled transmembrane transporter activity, role in iron-sulfur cluster assembly, iron-sulfur cluster transmembrane transport and mitochondrial inner membrane, mitochondrion localization | ORF, Uncharacterized | ATM1 | CAGL0M13739g | Schwarzmüller et al., 2014 |
| 2H4 | CAGL0M13849g | Verified | GAS2 | Putative glycoside hydrolase of the Gas/Phr family; predicted GPI-anchor | ORF, Verified | GAS1 | CAGL0M13849g | Schwarzmüller et al., 2014 |
| 2H5 | CAGL0I10945g | Verified | YCA1 | Putative metacaspase, cysteine protease involved in apoptosis in response to stresses | ORF, Uncharacterized | MCA1 | CAGL0I10945g | Schwarzmüller et al., 2014 |
| 2H6 | CAGL0L06490g | No Deletion Verified | | Ortholog(s) have role in mitochondrion inheritance, negative regulation of proteolysis, protein folding and mitochondrial inner membrane, mitochondrial membrane, mitochondrion localization | ORF, Uncharacterized | PHB2 | CAGL0L06490g | Schwarzmüller et al., 2014 |
| 2H7 | CAGL0L04092g | No Deletion Verified | | Ortholog(s) have serine-type endopeptidase activity, serine-type peptidase activity | ORF, Uncharacterized | NMA111 | CAGL0L04092g | Schwarzmüller et al., 2014 |
| 2H8 | CAGL0F08371g | Verified | TNA1 | High-affinity nicotinic acid transporter; strongly induced under niacin-limiting conditions | ORF, Verified | TNA1 | CAGL0F08371g | Schwarzmüller et al., 2014 |
| 2H9 | CAGL0J02948g | No Deletion Verified | FCY2 | Purine-cytosine transporter | ORF, Uncharacterized | FCY22 | CAGL0J02948g | Schwarzmüller et al., 2014 |
| 2H10 | CAGL0L00671g | No Deletion Verified | FCY21 | Purine-cytosine transporter | ORF, Uncharacterized | FCY21 | CAGL0L00671g | Schwarzmüller et al., 2014 |
| 2H11 | CAGL0L06138g | Verified | TPN1 | Ortholog(s) have vitamin transmembrane transporter activity, role in vitamin transport and plasma membrane localization | ORF, Uncharacterized | TPN1 | CAGL0L06138g | Schwarzmüller et al., 2014 |
| 3A4 | CAGL0G04873g | Verified | ROM2 | Ortholog(s) have guanyl-nucleotide exchange factor activity, phosphatidylinositol-4,5-bisphosphate binding activity | ORF, Uncharacterized | ROM2 | CAGL0G04873g | Schwarzmüller et al., 2014 |
| 3A5 | CAGL0L03520g | Verified | BCK1 | Ortholog(s) have MAP kinase kinase kinase activity | ORF, Uncharacterized | BCK1 | CAGL0L03520g | Schwarzmüller et al., 2014 |
| 3A6 | CAGL0J03828g | Verified | MKK1 | Ortholog(s) have MAP kinase kinase activity and role in cell wall integrity MAPK cascade, pexophagy, positive regulation of calcium-mediated signaling, regulation of fungal-type cell wall organization, signal transduction | ORF, Uncharacterized | MKK1 | CAGL0J03828g | Schwarzmüller et al., 2014 |
| 3A8 | CAGL0J11506g | Verified | | Ortholog(s) have chitin synthase activity, role in ascospore wall chitin biosynthetic process, septum digestion after cytokinesis and chitosome, plasma membrane localization | ORF, Uncharacterized | CHS1 | CAGL0J11506g | Schwarzmüller et al., 2014 |
| 3A9 | CAGL0G00858g | Verified | | Ortholog(s) have transmembrane signaling receptor activity, role in IRE1-mediated unfolded protein response, fungal-type cell wall organization, pexophagy, response to acidic pH, response to osmotic stress and plasma membrane localization | ORF, Uncharacterized | MID2 | CAGL0G00858g | Schwarzmüller et al., 2014 |
| 3A11 | CAGL0L06336g | Verified | | Ortholog(s) have G-protein beta-subunit binding, MAP-kinase scaffold activity, phosphatidylinositol-4,5-bisphosphate b | ORF, Uncharacterized | STE5 | CAGL0L06336g | Schwarzmüller et al., 2014 |
| 3A12 | CAGL0J00253g | Verified | | Putative adhesin-like protein | ORF, Uncharacterized | MTL1 | CAGL0J00253g | Schwarzmüller et al., 2014 |
| 3B1 | CAGL0J00539g | Verified | SLT2 | Mitogen-activated protein kinase with a role in cell wall integrity | ORF, Verified | SLT2 | CAGL0J00539g | Schwarzmüller et al., 2014 |
| 3B3 | CAGL0L05632g | Verified | PBS2 | Ortholog(s) have MAP kinase kinase activity, MAP-kinase scaffold activity | ORF, Uncharacterized | PBS2 | CAGL0L05632g | Schwarzmüller et al., 2014 |
| 3B6 | CAGL0M13827g | Verified | FKS3 | Putative 1,3-beta-d-glucan synthase component | ORF, Uncharacterized | FKS3 | CAGL0M13827g | Schwarzmüller et al., 2014 |
| 3B7 | CAGL0K04037g | Verified | FKS2 | Putative 1,3-beta-glucan synthase component; functionally redundant with Fks1p; "hot spot" mutations in FKS2 confer resistance to echinocandins | ORF, Verified | GSC2 | CAGL0K04037g | Schwarzmüller et al., 2014 |
| 3B8 | CAGL0D02794g | Verified | | Protein of unknown function; Ortholog(s) are endoplasmic reticulum (ER) membrane protein; involved in the transloca | ORF, Uncharacterized | WSC4 | CAGL0D02794g | Schwarzmüller et al., 2014 |
| 3B9 | CAGL0F01507g | Verified | SLG1 | Putative sensor of stress-activated signaling; gene is downregulated in azole-resistant strain | ORF, Uncharacterized | SLG1 | CAGL0F01507g | Schwarzmüller et al., 2014 |
| 3B10 | CAGL0G02497g | Verified | | Ortholog(s) have promoter-terminator loop anchoring activity, ribonucleoprotein complex binding activity | ORF, Uncharacterized | MLP1 | CAGL0G02497g | Schwarzmüller et al., 2014 |
| 3B12 | CAGL0B04389g | Verified | CHS3 | Putative class IV chitin synthase; mutants display a significantly thickened cell wall mannoprotein layer; mutants are delayed in cell wall formation during protoplast regeneration | ORF, Verified | CHS3 | CAGL0B04389g | Schwarzmüller et al., 2014 |
| 3E6 | CAGL0L00627g | Wrong Strain | | Ortholog(s) have GTPase activating protein binding, cAMP-dependent protein kinase inhibitor activity | ORF, Uncharacterized | GPB2 | CAGL0L00627g | Schwarzmüller et al., 2014 |
| 3E8 | CAGL0J07282g | Verified | | Ortholog(s) have GTPase activating protein binding, cAMP-dependent protein kinase inhibitor activity | ORF, Uncharacterized | GPB1 | CAGL0J07282g | Schwarzmüller et al., 2014 |
| 3E11 | CAGL0G09020g | Verified | | Ortholog(s) have cAMP-dependent protein kinase activity, protein serine/threonine kinase activity | ORF, Uncharacterized | TPK2 | CAGL0G09020g | Schwarzmüller et al., 2014 |
| 3E12 | CAGL0K09944g | Verified | PDE2 | Ortholog(s) have 3',5'-cyclic-AMP phosphodiesterase activity | ORF, Uncharacterized | PDE2 | CAGL0K09944g | Schwarzmüller et al., 2014 |
| 3H12 | CAGL0K04169g | Verified | | Ortholog(s) have MAP kinase activity | ORF, Uncharacterized | KSS1 | CAGL0K04169g | Schwarzmüller et al., 2014 |
| 4A1 | CAGL0D05434g | Verified | ROX1 | Protein involved in regulation of ergosterol biosynthesis; mutations suppress fluconazole sensitivity of upc2a mutants | ORF, Verified | ROX1 | CAGL0D05434g | Schwarzmüller et al., 2014 |
| 4A2 | CAGL0K05841g | Verified | HAP1 | Has domain(s) with predicted DNA binding, DNA-binding transcription activity, DNA-binding transcription factor activity, RNA polymerase II-specific, zinc ion binding activity | ORF, Uncharacterized | HAP1 | CAGL0K05841g | Schwarzmüller et al., 2014 |
| 4A3 | CAGL0K03003g | Verified | | Ortholog(s) have DNA-binding transcription factor activity, RNA polymerase II-specific, RNA polymerase II cis-regulatory region sequence-specific DNA binding, RNA polymerase II-specific DNA-binding transcription factor binding activity | ORF, Uncharacterized | MOT3 | CAGL0K03003g | Schwarzmüller et al., 2014 |
| 4A4 | CAGL0B01441g | Verified | RPD3 | Ortholog(s) have histone H3K14 deacetylase activity, histone H3K9 deacetylase activity, histone H4K16 deacetylase activity, protein lysine deacetylase activity and role in epigenetic regulation of gene expression, heterochromatin formation | ORF, Uncharacterized | RPD3 | CAGL0B01441g | Schwarzmüller et al., 2014 |
| 4A5 | CAGL0L00583g | Verified | BCR1 | Ortholog(s) have role in carbon catabolite activation of transcription from RNA polymerase II promoter, regulation of transcription by RNA polymerase II and nucleus localization | ORF, Uncharacterized | USV1 | CAGL0L00583g | Schwarzmüller et al., 2014 |
| 4A6 | CAGL0E01859g | Verified | YPS10 | Putative aspartic protease; predicted GPI-anchor; member of a YPS gene cluster that is required for virulence in mice; induced in response to low pH and high temperature | ORF, Verified | YPS1 | CAGL0E01859g | Schwarzmüller et al., 2014 |
| 4A7 | CAGL0E01837g | Wrong Strain | YPS9 | Putative aspartic protease; predicted GPI-anchor; member of a YPS gene cluster that is required for virulence in mice; expression induced at high temperature | ORF, Verified | MKC7 | CAGL0E01837g | Schwarzmüller et al., 2014 |
| 4A8 | CAGL0L06424g | Verified | | Predicted GPI-linked adhesin-like protein | ORF, Verified | CCW12 | CAGL0L06424g | Schwarzmüller et al., 2014 |
| 4A9 | CAGL0G00286g | No Deletion Verified | GAS1 | Glycoside hydrolase of the Gas/Phr family, involved in cell wall maintenance; confers resistance to azole drugs; predicted GPI-anchor | ORF, Verified | GAS1 | CAGL0G00286g | Schwarzmüller et al., 2014 |
| 4A10 | CAGL0E02321g | Verified | | Putative phospholipase B; predicted GPI-anchor | ORF, Verified | PLB3 | CAGL0E02321g | Schwarzmüller et al., 2014 |
| 4A11 | CAGL0E01419g | Verified | YPS2 | Putative aspartic protease; predicted GPI-anchor; member of a YPS gene cluster that is required for virulence in mice; induced in response to low pH and high temperature | ORF, Verified | MKC7 | CAGL0E01419g | Schwarzmüller et al., 2014 |
| 4A12 | CAGL0J08712g | Verified | | Ortholog(s) have nicotinamide riboside transmembrane transporter activity, nucleobase transmembrane transporter activity, nucleoside transmembrane transporter activity | ORF, Uncharacterized | FUN26 | CAGL0J08712g | Schwarzmüller et al., 2014 |
| 4B1 | CAGL0I05478g | No Deletion Verified | | Ortholog(s) have ubiquitin-protein transferase activity, role in protein monoubiquitination, protein polyubiquitination, ubiquitin-dependent ERAD pathway and endoplasmic reticulum membrane localization | ORF, Uncharacterized | UBC6 | CAGL0I05478g | Schwarzmüller et al., 2014 |
| 4B2 | CAGL0M08250g | Verified | | Ortholog(s) have zinc ion transmembrane transporter activity, role in intracellular zinc ion homeostasis, zinc ion transport and fungal-type vacuole membrane localization | ORF, Uncharacterized | ZRT3 | CAGL0M08250g | Schwarzmüller et al., 2014 |
| 4B3 | CAGL0C00693g | Verified | CCC1 | Putative vacuolar iron transporter | ORF, Uncharacterized | CCC1 | CAGL0C00693g | Schwarzmüller et al., 2014 |
| 4B4 | CAGL0K05489g | No Deletion Verified | | Ortholog(s) have role in endoplasmic reticulum membrane fusion, retrograde vesicle-mediated transport, Golgi to endoplasmic reticulum and SNARE complex, endoplasmic reticulum localization | ORF, Uncharacterized | UFE1 | CAGL0K05489g | Schwarzmüller et al., 2014 |
| 4B5 | CAGL0M05533g | Verified | DUR1,2 | Has domain(s) with predicted ATP binding activity | ORF, Verified | DUR12 | CAGL0M05533g | Schwarzmüller et al., 2014 |
| 4B6 | CAGL0J08998g | No Deletion Verified | | Ortholog(s) have structural molecule activity and role in COPII-coated vesicle budding, endoplasmic reticulum to Golgi vesicle-mediated transport, intracellular protein transport, nuclear envelope organization | ORF, Uncharacterized | SEC31 | CAGL0J08998g | Schwarzmüller et al., 2014 |
| 4B7 | CAGL0L03828g | No Deletion Verified | | Ortholog(s) have electron transfer activity, role in sterol biosynthetic process and endoplasmic reticulum membrane localization | ORF, Uncharacterized | CYB5 | CAGL0L03828g | Schwarzmüller et al., 2014 |
| 4B9 | CAGL0M04807g | No Deletion Verified | SNF2 | Component of the chromatin remodelling Swi/Snf complex; involved in regulation of biofilm formation | ORF, Verified | SNF2 | CAGL0M04807g | Schwarzmüller et al., 2014 |
| 4B10 | CAGL0E03718g | Verified | SNF6 | Component of the chromatin remodelling Swi/Snf complex; involved in regulation of biofilm formation | ORF, Verified | SNF6 | CAGL0E03718g | Schwarzmüller et al., 2014 |
| 4B11 | CAGL0F09097g | Verified | SKN7 | Predicted transcription factor, involved in oxidative stress response; required for induction of TRX2, TRR1 and TSA1 transcription under oxidative stress | ORF, Verified | SKN7 | CAGL0F09097g | Schwarzmüller et al., 2014 |
| 4B12 | CAGL0I05896g | Wrong Strain | YAK1 | Putative serine-threonine protein kinase, involved in biofilm formation, required for expression of adhesin genes EPA6 and EPA7 | ORF, Verified | YAK1 | CAGL0I05896g | Schwarzmüller et al., 2014 |
| 4C1 | CAGL0I04818g | No Deletion Verified | | Ortholog(s) have chitin synthase activity, role in chitin biosynthetic process, mitotic actomyosin contractile ring contraction and cellular bud neck localization | ORF, Uncharacterized | CHS2 | CAGL0I04818g | Schwarzmüller et al., 2014 |
| 4C2 | CAGL0K06193g | No Deletion Verified | ADA2 | Putative transcription coactivator, component of the Spt-Ada-Gcn5 acetyltransferase (SAGA) complex; involved in drug resistance and virulence | ORF, Verified | ADA2 | CAGL0K06193g | Schwarzmüller et al., 2014 |
| 4C3 | CAGL0A01452g | Verified | CWH41 | Putative glucosidase I; glycoside hydrolase; predicted GPI-anchor | ORF, Uncharacterized | CWH41 | CAGL0A01452g | Schwarzmüller et al., 2014 |
| 4C4 | CAGL0A01584g | Verified | | Ortholog(s) have cell adhesion molecule binding activity, role in agglutination involved in conjugation with cellular fusion and fungal-type cell wall localization | ORF, Uncharacterized | AGA2 | CAGL0A01584g | Schwarzmüller et al., 2014 |
| 4C5 | CAGL0A02486g | Verified | | Ortholog(s) have role in fungal-type cell wall organization and Golgi apparatus localization | ORF, Uncharacterized | SBE2 | CAGL0A02486g | Schwarzmüller et al., 2014 |
| 4C6 | CAGL0A04411g | Verified | SKT5 | Ortholog(s) have enzyme activator activity and role in fungal-type cell wall beta-glucan biosynthetic process, fungal-type cell wall chitin biosynthetic process, mitotic division septum assembly | ORF, Uncharacterized | SKT5 | CAGL0A04411g | Schwarzmüller et al., 2014 |
| 4C7 | CAGL0B00528g | Verified | | Ortholog(s) have protein kinase inhibitor activity, role in fungal-type cell wall organization and cellular bud neck localization | ORF, Uncharacterized | LRE1 | CAGL0B00528g | Schwarzmüller et al., 2014 |
| 4C8 | CAGL0B01969g | Verified | | Putative protein; gene is downregulated in azole-resistant strain | ORF, Uncharacterized | ECM18 | CAGL0B01969g | Schwarzmüller et al., 2014 |

| | | | | | | | | |
|---|---|---|---|---|---|---|---|---|
| **4C10** | CAGL0B04565g | Verified | | Ortholog(s) have alpha-1,2-mannosyltransferase activity, role in protein N-linked glycosylation, protein O-linked glycosylation and Golgi apparatus localization | ORF, Uncharactarized | *KTR1* | CAGL0B04565g | Schwarzmüller et al., 2014 |
| **4C11** | CAGL0B04631g | Verified | | Putative polyphosphatidylinositol phosphatase; null mutant does not show dependence on CRZ1 in response to a cell wall stressor | ORF, Uncharactarized | *INP53* | CAGL0B04631g | Schwarzmüller et al., 2014 |
| **4C12** | CAGL0C00209g | No Deletion Verified | *AWP7* | Putative adhesin-like cell wall protein; belongs to adhesin cluster IV; predicted GPI-anchor | ORF, Verified | *DAN4* | CAGL0C00209g | Schwarzmüller et al., 2014 |
| **4D1** | CAGL0C00363g | No Deletion Verified | *KRE9* | Protein involved in cell wall beta-1,6-glucan synthesis | ORF, Verified | *KRE9* | CAGL0C00363g | Schwarzmüller et al., 2014 |
| **4D2** | CAGL0C04367g | No Deletion Verified | | Non-essential protein of unknown function; likely exists as tetramer, may be regulated by the binding of small-molecule ligands (possibly sulfate ions), may have a role in yeast cell-wall biogenesis | ORF, Uncharactarized | *ECM15* | CAGL0C04367g | Schwarzmüller et al., 2014 |
| **4D3** | CAGL0C04961g | No Deletion Verified | | Ortholog(s) have role in intracellular calcium ion homeostasis, regulation of G0 to G1 transition | ORF, Uncharactarized | *ECM27* | CAGL0C04961g | Schwarzmüller et al., 2014 |
| **4D4** | CAGL0C05599g | Verified | | Ortholog(s) have GTPase activator activity, role in regulation of actin cytoskeleton organization, regulation of cell wall organization or biogenesis and cell cortex of cell tip, division septum localization | ORF, Uncharactarized | *LRG1* | CAGL0C05599g | Schwarzmüller et al., 2014 |
| **4D5** | CAGL0D00220g | Verified | | Ortholog(s) have role in ribosomal large subunit export from nucleus and nucleolus, nucleus, preribosome, large subunit precursor localization | ORF, Uncharactarized | *ECM1* | CAGL0D00220g | Schwarzmüller et al., 2014 |
| **4D6** | CAGL0D01034g | Verified | *VIG9* | GDP-mannose pyrophosphorylase involved in the synthesis of GDP-mannose for protein glycosylation | ORF, Verified | *PSA1* | CAGL0D01034g | Schwarzmüller et al., 2014 |
| **4D7** | CAGL0D03256g | Verified | | Ortholog(s) have sterol esterase activity, role in cell wall mannoprotein biosynthetic process, sterol metabolic process and membrane, plasma membrane localization | ORF, Uncharactarized | *YEH2* | CAGL0D03256g | Schwarzmüller et al., 2014 |
| **4D8** | CAGL0D06446g | Verified | *STT4* | Ortholog(s) have 1-phosphatidylinositol 4-kinase activity and role in autophagosome-lysosome fusion, autophagy of mitochondrion, macroautophagy, microlipophagy, phosphatidylinositol phosphate biosynthetic process | ORF, Uncharactarized | *STT4* | CAGL0D06446g | Schwarzmüller et al., 2014 |
| **4D9** | CAGL0D06622g | Verified | | Ortholog(s) have MAP-kinase scaffold activity | ORF, Uncharactarized | *SPA2* | CAGL0D06622g | Schwarzmüller et al., 2014 |
| **4D10** | CAGL0E01353g | Verified | | Putative high-affinity zinc transporter; gene is downregulated in azole-resistant strain | ORF, Uncharactarized | *ZRT2* | CAGL0E01353g | Schwarzmüller et al., 2014 |
| **4D11** | CAGL0E02255g | No Deletion Verified | | Ortholog(s) have role in fungal-type cell wall organization and plasma membrane localization | ORF, Verified | *ZEO1* | CAGL0E02255g | Schwarzmüller et al., 2014 |
| **4D12** | CAGL0E02761g | Verified | | Ortholog(s) have role in chromatin remodeling, nucleosome disassembly, transcription elongation by RNA polymerase II and RSC-type complex localization | ORF, Uncharactarized | *LDB7* | CAGL0E02761g | Schwarzmüller et al., 2014 |
| **4E1** | CAGL0E02783g | Verified | | Ortholog(s) have cargo adaptor activity, ubiquitin binding activity and role in actin cortical patch assembly, endocytosis, negative regulation of Arp2/3 complex-mediated actin nucleation | ORF, Uncharactarized | *SLA1* | CAGL0E02783g | Schwarzmüller et al., 2014 |
| **4E2** | CAGL0E02915g | Verified | | Putative adhesin-like protein | ORF, Verified | *SCW11* | CAGL0E02915g | Schwarzmüller et al., 2014 |
| **4E3** | CAGL0E03564g | No Deletion Verified | | Ortholog(s) have GTP binding, phosphatidylinositol-4-phosphate binding, phosphatidylinositol-5-phosphate binding, structural constituent of cytoskeleton activity | ORF, Uncharactarized | *CDC3* | CAGL0E03564g | Schwarzmüller et al., 2014 |
| **4E4** | CAGL0E04840g | No Deletion Verified | | Ortholog(s) have role in ascospore formation and ascospore wall, meiotic spindle, prospore membrane, septin complex, spindle microtubule localization | ORF, Uncharactarized | *SPR28* | CAGL0E04840g | Schwarzmüller et al., 2014 |
| **4E5** | CAGL0F00275g | No Deletion Verified | | Ortholog(s) have GTP binding, phosphatidylinositol-4-phosphate binding, phosphatidylinositol-5-phosphate binding, structural constituent of cytoskeleton activity | ORF, Uncharactarized | *CDC11* | CAGL0F00275g | Schwarzmüller et al., 2014 |
| **4E6** | CAGL0F00297g | No Deletion Verified | | Ortholog(s) have alpha-1,6-mannosyltransferase activity and mannan polymerase complex localization | ORF, Uncharactarized | *HOC1* | CAGL0F00297g | Schwarzmüller et al., 2014 |
| **4E7** | CAGL0F01287g | No Deletion Verified | | Putative transglycosidase with a predicted role in the elongation of 1,3-beta-glucan | ORF, Verified | *GAS5* | CAGL0F01287g | Schwarzmüller et al., 2014 |
| **4E8** | CAGL0F02453g | Verified | | Ortholog(s) have actinin binding activity and role in regulation of cellular response to stress, regulation of unidimensional cell growth | ORF, Uncharactarized | *ECM25* | CAGL0F02453g | Schwarzmüller et al., 2014 |
| **4E9** | CAGL0F03003g | Verified | | Ortholog(s) have osmosensor activity and role in (1->3)-beta-D-glucan biosynthetic process, cellular bud site selection, fungal-type cell wall organization, hyperosmotic response, osmosensory signaling pathway via Sho1 osmosensor | ORF, Verified | *HKR1* | CAGL0F03003g | Schwarzmüller et al., 2014 |
| **4E10** | CAGL0F03487g | No Deletion Verified | | Protein of unknown function; Ortholog(s) are component of the EKC/KEOPS protein complex | ORF, Uncharactarized | *GON7* | CAGL0F03487g | Schwarzmüller et al., 2014 |
| **4E11** | CAGL0F03597g | No Deletion Verified | | Ortholog(s) have phosphatidylinositol N-acetylglucosaminyltransferase activity and endoplasmic reticulum localization | ORF, Uncharactarized | *SPT14* | CAGL0F03597g | Schwarzmüller et al., 2014 |
| **4E12** | CAGL0F04521g | Verified | | Ortholog of *S. cerevisiae* : ECM13 and *Saccharomyces cerevisiae S288C* : YBL043W | ORF, Uncharactarized | *ECM13* | CAGL0F04521g | Schwarzmüller et al., 2014 |
| **4F1** | CAGL0F04873g | Verified | | Ortholog(s) have glucosidase activity and role in (1->6)-beta-D-glucan biosynthetic process, fungal-type cell wall organization | ORF, Verified | *KRE6* | CAGL0F04873g | Schwarzmüller et al., 2014 |
| **4F2** | CAGL0F06501g | Verified | | Ortholog(s) have acetyl-CoA:L-glutamate N-acetyltransferase activity, role in ornithine biosynthetic process and mitochondrial matrix localization | ORF, Uncharactarized | *ARG7* | CAGL0F06501g | Schwarzmüller et al., 2014 |
| **4F3** | CAGL0G00220g | No Deletion Verified | | Ortholog(s) have glucan endo-1,3-beta-D-glucosidase activity, role in fungal-type cell wall organization and fungal-type cell wall localization | ORF, Verified | *BGL2* | CAGL0G00220g | Schwarzmüller et al., 2014 |
| **4F4** | CAGL0G00308g | No Deletion Verified | *SCW4* | Putative transglycosidase with a predicted role in the modification of 1,3-beta-glucan | ORF, Verified | *SCW4* | CAGL0G00308g | Schwarzmüller et al., 2014 |
| **4F5** | CAGL0G00814g | Verified | *CHS5* | Ortholog(s) have small GTPase binding activity | ORF, Uncharactarized | *CHS5* | CAGL0G00814g | Schwarzmüller et al., 2014 |
| **4F6** | CAGL0G02101g | Verified | *ECM4* | Putative omega class glutathione transferase; gene is downregulated in azole-resistant strain | ORF, Verified | *ECM4* | CAGL0G02101g | Schwarzmüller et al., 2014 |
| **4F7** | CAGL0G02343g | Verified | | Ortholog(s) have role in mitotic spindle elongation, vesicle-mediated transport and Golgi membrane localization | ORF, Uncharactarized | *TVP38* | CAGL0G02343g | Schwarzmüller et al., 2014 |
| **4F8** | CAGL0G05566g | Verified | | Ortholog(s) have role in ascospore formation, fungal-type cell wall organization and cell cortex localization | ORF, Uncharactarized | *FMP45* | CAGL0G05566g | Schwarzmüller et al., 2014 |
| **4F9** | CAGL0G05918g | Verified | | Ortholog(s) have unfolded protein binding activity and role in endoplasmic reticulum to Golgi vesicle-mediated transport, fungal-type cell wall chitin biosynthetic process, protein folding | ORF, Uncharactarized | *CHS7* | CAGL0G05918g | Schwarzmüller et al., 2014 |
| **4F10** | CAGL0G06072g | Verified | | Has domain(s) with predicted metallocarboxypeptidase activity, zinc ion binding activity and role in proteolysis | ORF, Uncharactarized | *ECM14* | CAGL0G06072g | Schwarzmüller et al., 2014 |
| **4F11** | CAGL0G06754g | Verified | | Ortholog(s) have 1-phosphatidylinositol binding, GTPase activity, molecular adaptor activity, phosphatidylinositol-4-phosphate binding, phosphatidylinositol-5-phosphate binding, structural constituent of cytoskeleton activity | ORF, Uncharactarized | *CDC10* | CAGL0G06754g | Schwarzmüller et al., 2014 |
| **4F12** | CAGL0H00550g | No Deletion Verified | | Ortholog(s) have role in mitochondrial genome maintenance | ORF, Uncharactarized | *ILM1* | CAGL0H00550g | Schwarzmüller et al., 2014 |
| **4G1** | CAGL0H00847g | Verified | | Ortholog(s) have UDP-galactose transmembrane transporter activity and role in UDP-galactose transmembrane transport, UDP-glucose transmembrane transport, UDP-glucose transmembrane transport into endoplasmic reticulum | ORF, Uncharactarized | *HUT1* | CAGL0H00847g | Schwarzmüller et al., 2014 |
| **4G2** | CAGL0H01287g | Verified | *SSD1* | Putative mRNA-binding protein involved in negative regulation of translation; regulates echinocandin resistance genes FKS1 and FKS2 | ORF, Uncharactarized | *SSD1* | CAGL0H01287g | Schwarzmüller et al., 2014 |
| **4G3** | CAGL0H05005g | Verified | | Ortholog(s) have phosphatidylinositol transfer activity | ORF, Uncharactarized | *CSR1* | CAGL0H05005g | Schwarzmüller et al., 2014 |
| **4G4** | CAGL0H06435g | Verified | | Ortholog of *S. cerevisiae* : ECM19 and *Saccharomyces cerevisiae S288C* : YLR390W | ORF, Uncharactarized | *ECM19* | CAGL0H06435g | Schwarzmüller et al., 2014 |
| **4G5** | CAGL0H06545g | Verified | | Ortholog(s) have role in autophagy of mitochondrion, mitochondria-nucleus signaling pathway and mitochondrial outer membrane, mitochondrion localization | ORF, Uncharactarized | *ATG32* | CAGL0H06545g | Schwarzmüller et al., 2014 |
| **4G6** | CAGL0H07403g | Verified | *KRE2* | Ortholog(s) have alpha-1,2-mannosyltransferase activity and role in N-glycan processing, cell wall mannoprotein biosynthetic process, chain elongation of O-linked mannose residue, protein O-linked glycosylation | ORF, Uncharactarized | *KRE2* | CAGL0H07403g | Schwarzmüller et al., 2014 |
| **4G7** | CAGL0M07381g | Verified | | Ortholog(s) have mannosylphosphate transferase activity, role in endoplasmic reticulum to Golgi vesicle-mediated transport, fungal-type cell wall organization, protein N-linked glycosylation and endoplasmic reticulum, membrane localization | ORF, Uncharactarized | *KTR6* | CAGL0M07381g | Schwarzmüller et al., 2014 |
| **4G8** | CAGL0H07997g | No Deletion Verified | *KNH1* | Protein involved in cell wall beta 1,6-glucan synthesis, similar to Kre9p | ORF, Verified | *KRE9* | CAGL0H07997g | Schwarzmüller et al., 2014 |
| **4G9** | CAGL0I00484g | Verified | | Ortholog(s) have glucan endo-1,6-beta-glucosidase activity and role in fungal-type cell wall beta-glucan metabolic process, fungal-type cell wall disassembly involved in conjugation with cellular fusion, glucan metabolic process | ORF, Verified | *EXG1* | CAGL0I00484g | Schwarzmüller et al., 2014 |
| **4G10** | CAGL0I01188g | Verified | | Ortholog(s) have GTP binding, GTPase activity, molecular adaptor activity, phosphatidylinositol-4-phosphate binding, phosphatidylinositol-5-phosphate binding, structural constituent of cytoskeleton activity | ORF, Uncharactarized | *CDC12* | CAGL0I01188g | Schwarzmüller et al., 2014 |
| **4G11** | CAGL0I06160g | Verified | *PIR4* | Pir protein family member, putative cell wall component | ORF, Verified | *CIS3* | CAGL0I06160g | Schwarzmüller et al., 2014 |
| **4G12** | CAGL0I06182g | No Deletion Verified | *PIR2* | Pir protein family member, putative cell wall component | ORF, Verified | *HSP150* | CAGL0I06182g | Schwarzmüller et al., 2014 |
| **4H1** | CAGL0I06204g | No Deletion Verified | *PIR1* | Pir protein family member, putative cell wall component | ORF, Verified | *PIR5* | CAGL0I06204g | Schwarzmüller et al., 2014 |
| **4H2** | CAGL0I06512g | No Deletion Verified | *BEM2* | Ortholog(s) have GTPase activator activity and role in actin cytoskeleton organization, mitotic morphogenesis checkpoint signaling, negative regulation of Rho protein signal transduction | ORF, Uncharactarized | *BEM2* | CAGL0I06512g | Schwarzmüller et al., 2014 |
| **4H3** | CAGL0I07513g | Verified | | Ortholog(s) have protein kinase activity, protein serine/threonine kinase activity | ORF, Uncharactarized | *PKH2* | CAGL0I07513g | Schwarzmüller et al., 2014 |
| **4H4** | CAGL0G04609g | Verified | *PKH2* | Ortholog(s) have protein serine/threonine kinase activity | ORF, Uncharactarized | *PKH1* | CAGL0G04609g | Schwarzmüller et al., 2014 |
| **4H5** | CAGL0I08459g | No Deletion Verified | *RHO1* | Ortholog(s) have G-protein beta-subunit binding, GDP binding, GTPase activity, enzyme activator activity, signaling adaptor activity | ORF, Verified | *RHO1* | CAGL0I08459g | Schwarzmüller et al., 2014 |
| **4H6** | CAGL0I08855g | Verified | | Ortholog(s) have guanyl-nucleotide exchange factor activity, role in intra-Golgi vesicle-mediated transport, protein-containing complex assembly and TRAPPII protein complex, trans-Golgi network localization | ORF, Uncharactarized | *TRS65* | CAGL0I08855g | Schwarzmüller et al., 2014 |
| **4H7** | CAGL0I10054g | No Deletion Verified | | Ortholog(s) have glucosidase activity, role in (1->6)-beta-D-glucan biosynthetic process, fungal-type cell wall organization, negative regulation of autophagy, sphingolipid biosynthetic process and membrane localization | ORF, Uncharactarized | *SKN1* | CAGL0I10054g | Schwarzmüller et al., 2014 |
| **4H9** | CAGL0J00803g | Verified | | Ortholog(s) have structural constituent of cytoskeleton activity, role in fungal-type cell wall organization, negative regulation of microtubule depolymerization and microtubule localization | ORF, Uncharactarized | *MHP1* | CAGL0J00803g | Schwarzmüller et al., 2014 |
| **4H10** | CAGL0J02134g | Verified | | Ortholog(s) have phosphatidylinositol-4,5-bisphosphate 5-phosphatase activity, role in phosphatidylinositol dephosphorylation and cytoplasm, membrane localization | ORF, Uncharactarized | *INP51* | CAGL0J02134g | Schwarzmüller et al., 2014 |
| **4H11** | CAGL0J02244g | No Deletion Verified | | Has domain(s) with predicted mannose-6-phosphate isomerase activity, zinc ion binding activity and role in GDP-mannose biosynthetic process, carbohydrate metabolic process | ORF, Uncharactarized | *PMI40* | CAGL0J02244g | Schwarzmüller et al., 2014 |
| **4H12** | CAGL0J03366g | Verified | *GET2* | Ortholog(s) have protein transmembrane transporter activity, protein-membrane adaptor activity | ORF, Uncharactarized | *GET2* | CAGL0J03366g | Schwarzmüller et al., 2014 |
| **5A2** | CAGL0J05236g | Verified | *LAS21* | Protein predicted to be involved in GPI anchor biosynthesis in the ER; null mutant cannot grow in the presence of calcineurin inhibitors | ORF, Uncharactarized | *LAS21* | CAGL0J05236g | Schwarzmüller et al., 2014 |
| **5A3** | CAGL0J05544g | Wrong Strain | | Ortholog(s) have protein-macromolecule adaptor activity, role in ascospore wall assembly, endocytosis, reticulophagy and actin cytoskeleton-regulatory complex localization | ORF, Uncharactarized | *END3* | CAGL0J05544g | Schwarzmüller et al., 2014 |
| **5A4** | CAGL0J06072g | Verified | *CBK1* | Ortholog(s) have protein serine/threonine kinase activity | ORF, Uncharactarized | *CBK1* | CAGL0J06072g | Schwarzmüller et al., 2014 |
| **5A5** | CAGL0J07656g | Verified | *SLA2* | Ortholog(s) have role in actin cortical patch assembly, actin filament organization, endocytosis, negative regulation of Arp2/3 complex-mediated actin nucleation and actin cortical patch, incipient cellular bud site localization | ORF, Uncharactarized | *SLA2* | CAGL0J07656g | Schwarzmüller et al., 2014 |
| **5A6** | CAGL0J08910g | Verified | | Has domain(s) with predicted hydrolase activity, hydrolyzing O-glycosyl compounds activity and role in carbohydrate metabolic process | ORF, Uncharactarized | *CRR1* | CAGL0J08910g | Schwarzmüller et al., 2014 |
| **5A7** | CAGL0J09416g | Verified | | Ortholog(s) have fungal-type vacuole lumen, fungal-type vacuole membrane localization | ORF, Uncharactarized | *SNA4* | CAGL0J09416g | Schwarzmüller et al., 2014 |
| **5A8** | CAGL0K02277g | Verified | | Ortholog(s) have G-protein beta/gamma-subunit complex binding activity | ORF, Uncharactarized | *DSE1* | CAGL0K02277g | Schwarzmüller et al., 2014 |
| **5A9** | CAGL0K03487g | No Deletion Verified | | Ortholog(s) have Arp2/3 complex binding, actin filament binding, microfilament motor activity | ORF, Uncharactarized | *MYO5* | CAGL0K03487g | Schwarzmüller et al., 2014 |
| **5A10** | CAGL0K04455g | Verified | | Ortholog(s) have role in ascospore formation and ascospore wall, prospore membrane, septin complex localization | ORF, Uncharactarized | *SPR3* | CAGL0K04455g | Schwarzmüller et al., 2014 |
| **5A11** | CAGL0K05247g | No Deletion Verified | | Ortholog(s) have ubiquitin protein ligase binding activity, role in fungal-type cell wall organization, regulation of transcription by RNA polymerase II and nucleus localization | ORF, Uncharactarized | *CSR2* | CAGL0K05247g | Schwarzmüller et al., 2014 |
| **5A12** | CAGL0K06127g | Wrong Strain | | Ortholog(s) have role in reciprocal meiotic recombination, replication-born double-strand break repair via sister chromatid exchange, synaptonemal complex organization and nucleus, synaptonemal complex localization | ORF, Uncharactarized | *ECM11* | CAGL0K06127g | Schwarzmüller et al., 2014 |
| **5B1** | CAGL0K06281g | No Deletion Verified | | Ortholog(s) have guanylate kinase activity and role in GDP biosynthetic process | ORF, Uncharactarized | *GUK1* | CAGL0K06281g | Schwarzmüller et al., 2014 |
| **5B2** | CAGL0K06963g | Verified | *ROT2* | Ortholog(s) have Glc2Man9GlcNAc2 oligosaccharide glucosidase activity, glucan 1,3-alpha-glucosidase activity and role in N-glycan processing, endoplasmic reticulum mannose trimming, fungal-type cell wall beta-glucan biosynthetic process | ORF, Uncharactarized | *ROT2* | CAGL0K06963g | Schwarzmüller et al., 2014 |
| **5B3** | CAGL0K08228g | Verified | | Ortholog(s) have role in protein targeting to vacuolar membrane, vacuole organization and vacuolar membrane localization | ORF, Uncharactarized | *HFL1* | CAGL0K08228g | Schwarzmüller et al., 2014 |
| **5B4** | CAGL0K08316g | No Deletion Verified | *RHO4* | Ortholog(s) have GTPase activity and role in maintenance of cell polarity, positive regulation of formin-nucleated actin cable assembly, regulation of formin-nucleated actin cable assembly | ORF, Uncharactarized | *RHO4* | CAGL0K08316g | Schwarzmüller et al., 2014 |
| **5B5** | CAGL0K08558g | No Deletion Verified | | Ortholog(s) have glutamine-fructose-6-phosphate transaminase (isomerizing) activity and role in fungal-type cell wall chitin biosynthetic process | ORF, Uncharactarized | *GFA1* | CAGL0K08558g | Schwarzmüller et al., 2014 |

| | | | | | | | | |
|---|---|---|---|---|---|---|---|---|
| **5B6** | CAGL0K11231g | Verified | *MNN10* | Ortholog(s) have alpha-1,6-mannosyltransferase activity, role in cell wall mannoprotein biosynthetic process, division septum assembly, protein N-linked glycosylation and Golgi apparatus, mannan polymerase complex localization | ORF, Uncharacterized | *MNN10* | CAGL0K11231g | Schwarzmüller et al., 2014 |
| **5B7** | CAGL0H09614g | No Deletion Verified | | Putative GPI-linked cell wall protein | ORF, Verified | *TIR1* | CAGL0H09614g | Schwarzmüller et al., 2014 |
| **5B8** | CAGL0L02255g | No Deletion Verified | | Protein of unknown function | ORF, Uncharacterized | *ECM9* | CAGL0L02255g | Schwarzmüller et al., 2014 |
| **5B9** | CAGL0L03113g | No Deletion Verified | | Ortholog(s) have role in vesicle-mediated transport and Golgi membrane localization | ORF, Uncharacterized | *GMH1* | CAGL0L03113g | Schwarzmüller et al., 2014 |
| **5B10** | CAGL0L03608g | Verified | | Ortholog(s) have role in Golgi to plasma membrane transport, fungal-type cell wall chitin biosynthetic process and exomer complex, trans-Golgi network transport vesicle localization | ORF, Uncharacterized | *CHS6* | CAGL0L03608g | Schwarzmüller et al., 2014 |
| **5B11** | CAGL0L03696g | Verified | | Has domain(s) with predicted role in transmembrane transport and membrane localization | ORF, Uncharacterized | *ECM3* | CAGL0L03696g | Schwarzmüller et al., 2014 |
| **5B12** | CAGL0L05962g | No Deletion Verified | | Ortholog(s) have mRNA binding, protein-macromolecule adaptor activity | ORF, Uncharacterized | *MPT5* | CAGL0L05962g | Schwarzmüller et al., 2014 |
| **5C1** | CAGL0L06534g | No Deletion Verified | | Ortholog(s) have role in regulation of fungal-type cell wall biogenesis, regulation of mitotic cell cycle and cellular bud neck, incipient cellular bud site localization | ORF, Uncharacterized | *SMI1* | CAGL0L06534g | Schwarzmüller et al., 2014 |
| **5C2** | CAGL0L07458g | No Deletion Verified | | Ortholog(s) have U6 snRNA binding activity, role in mRNA splicing, via spliceosome and Prp19 complex, post-mRNA release spliceosomal complex, spliceosomal complex localization | ORF, Uncharacterized | *ECM2* | CAGL0L07458g | Schwarzmüller et al., 2014 |
| **5C3** | CAGL0L07854g | Verified | *CWH43* | Ortholog(s) have role in GPI anchor biosynthetic process, GPI anchor metabolic process, fungal-type cell wall organization | ORF, Uncharacterized | *CWH43* | CAGL0L07854g | Schwarzmüller et al., 2014 |
| **5C4** | CAGL0L08140g | Verified | | Ortholog(s) have small GTPase binding activity, role in endocytosis, fungal-type cell wall organization, intracellular protein transport, protein lipidation, response to pH and cytosol, late endosome localization | ORF, Uncharacterized | *BPH1* | CAGL0L08140g | Schwarzmüller et al., 2014 |
| **5C5** | CAGL0L10670g | No Deletion Verified | | Predicted GPI-linked cell wall protein | ORF, Uncharacterized | *ROT1* | CAGL0L10670g | Schwarzmüller et al., 2014 |
| **5C6** | CAGL0L11528g | Verified | *BIG1* | Ortholog(s) have role in fungal-type cell wall biogenesis and endoplasmic reticulum membrane localization | ORF, Uncharacterized | *BIG1* | CAGL0L11528g | Schwarzmüller et al., 2014 |
| **5C7** | CAGL0L11550g | No Deletion Verified | | Ortholog(s) have protein kinase activity, role in budding cell apical bud growth, fungal-type cell wall organization and cellular bud, incipient cellular bud site, mating projection tip localization | ORF, Uncharacterized | *KIC1* | CAGL0L11550g | Schwarzmüller et al., 2014 |
| **5C8** | CAGL0L11572g | Verified | | Ortholog(s) have role in fungal-type cell wall organization | ORF, Uncharacterized | *SBE22* | CAGL0L11572g | Schwarzmüller et al., 2014 |
| **5C9** | CAGL0L11880g | Verified | *RPH1* | Ortholog(s) have DNA-binding transcription repressor activity, RNA polymerase II-specific, histone H3K36 demethylase activity, histone H3K9 demethylase activity, sequence-specific DNA binding activity | ORF, Uncharacterized | *RPH1* | CAGL0L11880g | Schwarzmüller et al., 2014 |
| **5C10** | CAGL0M00374g | No Deletion Verified | | Ortholog(s) have sulfite reductase (NADPH) activity, role in sulfate assimilation, sulfur amino acid biosynthetic process and sulfite reductase complex (NADPH) localization | ORF, Uncharacterized | *MET5* | CAGL0M00374g | Schwarzmüller et al., 2014 |
| **5C11** | CAGL0M00748g | Verified | *ECM7* | Putative integral membrane protein required for high-affinity Ca2+ influx | ORF, Verified | *ECM7* | CAGL0M00748g | Schwarzmüller et al., 2014 |
| **5C12** | CAGL0M00924g | No Deletion Verified | | Protein of unknown function; may play a role in cell wall biosynthesis | ORF, Uncharacterized | *ECM30* | CAGL0M00924g | Schwarzmüller et al., 2014 |
| **5D1** | CAGL0M01628g | Verified | *SAC7* | Ortholog(s) have GTPase activator activity | ORF, Verified | *SAC7* | CAGL0M01628g | Schwarzmüller et al., 2014 |
| **5D2** | CAGL0M01892g | No Deletion Verified | | Protein of unknown function | ORF, Uncharacterized | *ECM8* | CAGL0M01892g | Schwarzmüller et al., 2014 |
| **5D3** | CAGL0M03157g | Verified | | Ortholog(s) have role in autophagy, fungal-type cell wall organization, rDNA heterochromatin formation and phagophore assembly site localization | ORF, Uncharacterized | *IRS4* | CAGL0M03157g | Schwarzmüller et al., 2014 |
| **5D4** | CAGL0M05489g | Verified | | Has domain(s) with predicted mannosyltransferase activity, role in protein glycosylation and membrane localization | ORF, Uncharacterized | *KTR3* | CAGL0M05489g | Schwarzmüller et al., 2014 |
| **5D5** | CAGL0M05841g | Verified | | Ortholog(s) have mannosyltransferase activity, role in cell wall mannoprotein biosynthetic process, protein N-linked glycosylation and Golgi apparatus localization | ORF, Uncharacterized | *KTR2* | CAGL0M05841g | Schwarzmüller et al., 2014 |
| **5D6** | CAGL0M06787g | No Deletion Verified | | Has domain(s) with predicted mannosyltransferase activity, role in protein glycosylation and membrane localization | ORF, Uncharacterized | *KTR5* | CAGL0M06787g | Schwarzmüller et al., 2014 |
| **5D7** | CAGL0M08492g | Verified | *PIR3* | Pir protein family member, putative cell wall component | ORF, Verified | *PIR1* | CAGL0M08492g | Schwarzmüller et al., 2014 |
| **5D8** | CAGL0M09449g | Verified | | Ortholog(s) have ubiquitin protein ligase binding activity and role in ubiquitin-dependent endocytosis | ORF, Uncharacterized | *ECM21* | CAGL0M09449g | Schwarzmüller et al., 2014 |
| **5D9** | CAGL0M09779g | No Deletion Verified | *CTS1* | Putative endochitinase with a predicted role in cell separation | ORF, Verified | *CTS1* | CAGL0M09779g | Schwarzmüller et al., 2014 |
| **5D10** | CAGL0M13805g | Verified | *MP65* | 65 kDa mannoprotein | ORF, Verified | *SCW10* | CAGL0M13805g | Schwarzmüller et al., 2014 |
| **5D11** | CAGL0A00143g | Verified | *No Information* | | | | | Schwarzmüller et al., 2014 |
| **5D12** | CAGL0A00649g | No Deletion Verified | | Protein of unknown function | ORF, Uncharacterized | | CAGL0A00649g | Schwarzmüller et al., 2014 |
| **5E1** | CAGL0A00715g | No Deletion Verified | | Protein of unknown function | ORF, Uncharacterized | | CAGL0A00715g | Schwarzmüller et al., 2014 |
| **5E2** | CAGL0A01892g | Verified | | Protein of unknown function | ORF, Uncharacterized | | CAGL0A01892g | Schwarzmüller et al., 2014 |
| **5E3** | CAGL0A02299g | Verified | | Protein of unknown function | ORF, Verified | | CAGL0A02299g | Schwarzmüller et al., 2014 |
| **5E4** | CAGL0A02343g | No Deletion Verified | | Protein of unknown function | ORF, Uncharacterized | | CAGL0A02343g | Schwarzmüller et al., 2014 |
| **5E5** | CAGL0A03410g | Verified | | Protein of unknown function | ORF, Uncharacterized | | CAGL0A03410g | Schwarzmüller et al., 2014 |
| **5E6** | CAGL0A04609g | Verified | *No Information* | | | | | Schwarzmüller et al., 2014 |
| **5E7** | CAGL0A04763g | Verified | *No Information* | | | | | Schwarzmüller et al., 2014 |
| **5E8** | CAGL0A04851g | No Deletion Verified | *AWP14* | Adhesin protein associated with high-biofilm-forming strains | ORF, Verified | | CAGL0A04851g | Schwarzmüller et al., 2014 |
| **5E9** | CAGL0B00572g | Verified | | Protein of unknown function | ORF, Uncharacterized | | CAGL0B00572g | Schwarzmüller et al., 2014 |
| **5E10** | CAGL0B03025g | No Deletion Verified | *No Information* | | | | | Schwarzmüller et al., 2014 |
| **5E11** | CAGL0B03223g | Verified | | Protein of unknown function | ORF, Uncharacterized | | CAGL0B03223g | Schwarzmüller et al., 2014 |
| **5E12** | CAGL0C00781g | Verified | | Protein of unknown function | ORF, Uncharacterized | | CAGL0C00781g | Schwarzmüller et al., 2014 |
| **5F1** | CAGL0C00803g | Verified | | Homolog of CAGL0C00968g (Adhesin-like protein with a predicted role in cell adhesion; belongs to adhesin cluster VII; predicted GPI-anchor) | ORF, Uncharacterized | | CAGL0C00803g | Schwarzmüller et al., 2014 |
| **5F2** | CAGL0C00869g | Verified | | Homolog of CAGL0C00781g (Protein of unknown function) | ORF, Uncharacterized | | CAGL0C00869g | Schwarzmüller et al., 2014 |
| **5F3** | CAGL0C01111g | Verified | *No Information* | | | | | Schwarzmüller et al., 2014 |
| **5F4** | CAGL0C01617g | No Deletion Verified | | Protein of unknown function | ORF, Uncharacterized | | CAGL0C01617g | Schwarzmüller et al., 2014 |
| **5F5** | CAGL0C02057g | No Deletion Verified | | Protein of unknown function | ORF, Uncharacterized | | CAGL0C02057g | Schwarzmüller et al., 2014 |
| **5F6** | CAGL0C02365g | Verified | | Protein of unknown function | ORF, Uncharacterized | | CAGL0C02365g | Schwarzmüller et al., 2014 |
| **5F7** | CAGL0C02915g | Verified | *No Information* | | | | | Schwarzmüller et al., 2014 |
| **5F8** | CAGL0C03311g | Verified | | Protein of unknown function | ORF, Uncharacterized | | CAGL0C03311g | Schwarzmüller et al., 2014 |
| **5F9** | CAGL0C04158g | No Deletion Verified | *No Information* | | | | | Schwarzmüller et al., 2014 |
| **5F10** | CAGL0C05401g | Verified | | Protein of unknown function | ORF, Uncharacterized | | CAGL0C05401g | Schwarzmüller et al., 2014 |
| **5F11** | CAGL0D00396g | Verified | | Protein of unknown function | ORF, Uncharacterized | | CAGL0D00396g | Schwarzmüller et al., 2014 |
| **5F12** | CAGL0D01254g | No Deletion Verified | | Protein of unknown function | ORF, Uncharacterized | | CAGL0D01254g | Schwarzmüller et al., 2014 |
| **5G1** | CAGL0D02750g | Verified | | Protein of unknown function | ORF, Deleted, Uncharacterized | | CAGL0D02750g | Schwarzmüller et al., 2014 |
| **5G2** | CAGL0D04840g | Verified | | Ortholog(s) have role in Group I intron splicing, mitochondrial mRNA processing | ORF, Uncharacterized | *MSS18* | CAGL0D04840g | Schwarzmüller et al., 2014 |
| **5G3** | CAGL0D05104g | Verified | *Novel* | Protein of unknown function | ORF, Uncharacterized | | CAGL0D05104g | Schwarzmüller et al., 2014 |
| **5G4** | CAGL0D05456g | No Deletion Verified | | Protein of unknown function | ORF, Uncharacterized | | CAGL0D05456g | Schwarzmüller et al., 2014 |
| **5G5** | CAGL0D05654g | Verified | | Protein of unknown function | ORF, Uncharacterized | | CAGL0D05654g | Schwarzmüller et al., 2014 |
| **5G6** | CAGL0D06380g | No Deletion Verified | | Protein of unknown function | ORF, Uncharacterized | | CAGL0D06380g | Schwarzmüller et al., 2014 |
| **5G7** | CAGL0D06534g | Verified | | Protein of unknown function | ORF, Uncharacterized | | CAGL0D06534g | Schwarzmüller et al., 2014 |
| **5G8** | CAGL0D06666g | Verified | | Protein of unknown function | ORF, Uncharacterized | | CAGL0D06666g | Schwarzmüller et al., 2014 |
| **5G9** | CAGL0D06710g | No Deletion Verified | | Protein of unknown function | ORF, Uncharacterized | | CAGL0D06710g | Schwarzmüller et al., 2014 |
| **5G10** | CAGL0E00165g | Verified | | Putative adhesin-like protein; belongs to adhesin cluster III; predicted GPI anchor | ORF, Merged/Split, Uncharacterized | | CAGL0E00165g | Schwarzmüller et al., 2014 |
| **5G11** | CAGL0E01375g | Verified | *No Information* | | | | | Schwarzmüller et al., 2014 |
| **5G12** | CAGL0E02211g | Verified | | Protein of unknown function | ORF, Uncharacterized | | CAGL0E02211g | Schwarzmüller et al., 2014 |
| **5H1** | CAGL0E03069g | Verified | | Ortholog(s) have role in maturation of SSU-rRNA from tricistronic rRNA transcript (SSU-rRNA, 5.8S rRNA, LSU-rRNA), ribosomal small subunit biogenesis and nucleolus, small-subunit processome localization | ORF, Uncharacterized | *ENP2* | CAGL0E03069g | Schwarzmüller et al., 2014 |
| **5H2** | CAGL0E03135g | Verified | *No Information* | | | | | Schwarzmüller et al., 2014 |
| **5H3** | CAGL0E03498g | Verified | | Protein of unknown function | ORF, Uncharacterized | | CAGL0E03498g | Schwarzmüller et al., 2014 |
| **5H4** | CAGL0E04202g | Verified | *No Information* | | | | | Schwarzmüller et al., 2014 |
| **5H5** | CAGL0E04466g | No Deletion Verified | | Ortholog(s) have role in meiotic sister chromatid cohesion, positive regulation of sister chromatid cohesion, protein localization to chromosome, centromeric region | ORF, Uncharacterized | *SPO13* | CAGL0E04466g | Schwarzmüller et al., 2014 |
| **5H6** | CAGL0E04554g | No Deletion Verified | | Putative protein; gene is upregulated in azole-resistant strain | ORF, Uncharacterized | | CAGL0E04554g | Schwarzmüller et al., 2014 |
| **5H7** | CAGL0E06094g | Verified | | Protein of unknown function | ORF, Uncharacterized | | CAGL0E06094g | Schwarzmüller et al., 2014 |
| **5H8** | CAGL0E06578g | Verified | *No Information* | | | | | Schwarzmüller et al., 2014 |
| **5H9** | CAGL0E06622g | Verified | *No Information* | | | | | Schwarzmüller et al., 2014 |
| **5H10** | CAGL0F00341g | Verified | | Protein of unknown function | ORF, Uncharacterized | | CAGL0F00341g | Schwarzmüller et al., 2014 |
| **5H11** | CAGL0F00781g | Verified | *No Information* | | | | | Schwarzmüller et al., 2014 |
| **5H12** | CAGL0F01441g | Verified | *No Information* | | | | | Schwarzmüller et al., 2014 |
| **6A1** | CAGL0M07293g | No Deletion Verified | *PDR12* | Putative ABC transporter of weak organic acids; gene is downregulated in azole-resistant strain | ORF, Uncharacterized | *PDR12* | CAGL0M07293g | Schwarzmüller et al., 2014 |
| **6A2** | CAGL0K02145g | Verified | | Ortholog(s) have DNA-binding transcription factor activity and role in cellular response to sulfur dioxide, regulation of cellular response to stress | ORF, Uncharacterized | *COM2* | CAGL0K02145g | Schwarzmüller et al., 2014 |
| **6A3** | CAGL0A00583g | Verified | | Has domain(s) with predicted DNA binding, DNA-binding transcription factor activity, RNA polymerase II-specific, zinc ion binding activity and role in DNA-templated transcription, regulation of DNA-templated transcription | ORF, Uncharacterized | *PDR8* | CAGL0A00583g | Schwarzmüller et al., 2014 |
| **6A4** | CAGL0A01111g | No Deletion Verified | | Ortholog(s) have ATPase, proton-transporting ATP synthase activity, rotational mechanism activity and role in proton motive force-driven ATP synthesis | ORF, Uncharacterized | *ATP15* | CAGL0A01111g | Schwarzmüller et al., 2014 |
| **6A5** | CAGL0B00440g | Verified | | Protein of unknown function | ORF, Uncharacterized | | CAGL0B00440g | Schwarzmüller et al., 2014 |
| **6A6** | CAGL0B02651g | Verified | | Ortholog(s) have DNA-binding transcription factor activity, RNA polymerase II-specific, RNA polymerase II-specific DNA-binding transcription factor binding, cis-regulatory region sequence-specific DNA binding activity | ORF, Uncharacterized | *MET32* | CAGL0B02651g | Schwarzmüller et al., 2014 |
| **6A7** | CAGL0B03355g | Verified | | Putative DNA polymerase II subunit; gene is upregulated in azole-resistant strain | ORF, Uncharacterized | *DPB3* | CAGL0B03355g | Schwarzmüller et al., 2014 |
| **6A8** | CAGL0B04895g | No Deletion Verified | | Ortholog(s) have RNA polymerase II cis-regulatory region sequence-specific DNA binding activity and role in negative regulation of transcription by RNA polymerase II, positive regulation of transcription by RNA polymerase II | ORF, Uncharacterized | *RFX1* | CAGL0B04895g | Schwarzmüller et al., 2014 |
| **6A9** | CAGL0C00583g | Verified | | Ortholog(s) have mRNA binding, sequence-specific mRNA binding activity, role in endoplasmic reticulum inheritance, intracellular mRNA localization, mating type switching and cellular bud tip localization | ORF, Uncharacterized | *SHE3* | CAGL0C00583g | Schwarzmüller et al., 2014 |
| **6A10** | CAGL0C00715g | No Deletion Verified | | Ortholog(s) have role in ribosomal large subunit assembly and preribosome, large subunit precursor localization | ORF, Uncharacterized | *RSA3* | CAGL0C00715g | Schwarzmüller et al., 2014 |
| **6A11** | CAGL0C02739g | Verified | | Ortholog(s) have role in mitochondrion organization, phospholipid homeostasis and mitochondrial outer membrane localization | ORF, Uncharacterized | *FUN14* | CAGL0C02739g | Schwarzmüller et al., 2014 |
| **6A12** | CAGL0D00726g | No Deletion Verified | | Ortholog(s) have translation regulator activity, role in mitochondrial respiratory chain complex III assembly, positive regulation of mitochondrial translation and mitochondrial inner membrane, mitochondrial ribosome localization | ORF, Uncharacterized | *CBS1* | CAGL0D00726g | Schwarzmüller et al., 2014 |
| **6B1** | CAGL0D01210g | Verified | | Has domain(s) with predicted amino-acid racemase activity, racemase activity, acting on amino acids and derivatives, racemase and epimerase activity, acting on amino acids and derivatives activity | ORF, Uncharacterized | | CAGL0D01210g | Schwarzmüller et al., 2014 |
| **6B2** | CAGL0D01364g | No Deletion Verified | *CYC8* | Protein of unknown function | ORF, Uncharacterized | *CYC8* | CAGL0D01364g | Schwarzmüller et al., 2014 |
| **6B3** | CAGL0D05368g | No Deletion Verified | | Has domain(s) with predicted transcription coregulator activity, role in regulation of transcription by RNA polymerase II and mediator complex localization | ORF, Uncharacterized | *SRB6* | CAGL0D05368g | Schwarzmüller et al., 2014 |
| **6B4** | CAGL0D05632g | No Deletion Verified | | Ortholog(s) have copper chaperone activity, role in mitochondrial cytochrome c oxidase assembly, protein maturation by copper ion transfer and cytosol, mitochondrial intermembrane space localization | ORF, Uncharacterized | *COX17* | CAGL0D05632g | Schwarzmüller et al., 2014 |

| | | | | | | | | |
|---|---|---|---|---|---|---|---|---|
| **6B5** | CAGL0E04312g | No Deletion Verified | | Ortholog(s) have DNA-binding transcription activator activity, RNA polymerase II-specific, RNA polymerase II cis-regulatory region sequence-specific DNA binding activity and role in positive regulation of transcription by RNA polymerase II | ORF, Uncharacterized | STP2 | CAGL0E04312g | Schwarzmüller et al., 2014 |
| **6B6** | CAGL0F00209g | Verified | | Has domain(s) with predicted transmembrane transporter activity and role in transmembrane transport | ORF, Uncharacterized | TNA1 | CAGL0F00209g | Schwarzmüller et al., 2014 |
| **6B7** | CAGL0F03619g | Verified | | Ortholog(s) have role in establishment of mitotic spindle orientation and dynactin complex, spindle pole body localization | ORF, Uncharacterized | NIP100 | CAGL0F03619g | Schwarzmüller et al., 2014 |
| **6B8** | CAGL0F06237g | No Deletion Verified | | Ortholog(s) have DNA replication origin binding, DNA-binding transcription activator activity, RNA polymerase II-specific, DNA-binding transcription factor activity and RNA polymerase II-specific | ORF, Uncharacterized | MCM1 | CAGL0F06237g | Schwarzmüller et al., 2014 |
| **6B9** | CAGL0F06787g | Verified | | Ortholog(s) have ubiquitin ligase inhibitor activity | ORF, Uncharacterized | MND2 | CAGL0F06787g | Schwarzmüller et al., 2014 |
| **6B10** | CAGL0F07161g | No Deletion Verified | | Ortholog(s) have role in ubiquinone biosynthetic process, ubiquinone biosynthetic process via 3,4-dihydroxy-5-polyprenylbenzoate and mitochondrion localization | ORF, Merged/Split, Uncharacterized | | CAGL0F07161g | Schwarzmüller et al., 2014 |
| **6B11** | CAGL0F08195g | No Deletion Verified | | Ortholog(s) have role in pseudohyphal growth and chromatin localization | ORF, Uncharacterized | MGA1 | CAGL0F08195g | Schwarzmüller et al., 2014 |
| **6B12** | CAGL0G01012g | No Deletion Verified | | Ortholog(s) have role in ascospore formation, ascospore wall assembly, positive regulation of ascospore-type prospore membrane formation and cytoplasm localization | ORF, Uncharacterized | SPO77 | CAGL0G01012g | Schwarzmüller et al., 2014 |
| **6C1** | CAGL0H06347g | Verified | *DEP1* | Ortholog(s) have role in negative regulation of rDNA heterochromatin formation and negative regulation of silent mating-type cassette heterochromatin formation, negative regulation of transcription by RNA polymerase II, positive regulation of transcription by RNA polymerase II, regulation of DNA-templated DNA replication initiation, regulation of transcription by RNA polymerase II | ORF, Uncharacterized | DEP1 | CAGL0H06347g | Schwarzmüller et al., 2014 |
| **6C2** | CAGL0I03432g | Verified | | Ortholog(s) have role in DNA damage response and FANCM-MHF complex localization | ORF, Uncharacterized | MHF2 | CAGL0I03432g | Schwarzmüller et al., 2014 |
| **6C3** | CAGL0I06226g | No Deletion Verified | | Protein of unknown function, ortholog(s) contain a J-domain, which is a region with homology to the *E. coli* DnaJ protein | ORF, Uncharacterized | JJJ2 | CAGL0I06226g | Schwarzmüller et al., 2014 |
| **6C4** | CAGL0I10538g | Verified | | Ortholog(s) have role in mRNA splicing, via spliceosome | ORF, Uncharacterized | SYF2 | CAGL0I10538g | Schwarzmüller et al., 2014 |
| **6C5** | CAGL0I10769g | Verified | *MCM1* | Has domain(s) with predicted DNA binding, DNA-binding transcription factor activity, RNA polymerase II-specific, cis-regulatory region sequence-specific DNA binding, protein dimerization activity | ORF, Uncharacterized | MCM1 | CAGL0I10769g | Schwarzmüller et al., 2014 |
| **6C6** | CAGL0J06490g | Verified | | Has domain(s) with predicted ubiquitin binding activity | ORF, Uncharacterized | CUE4 | CAGL0J06490g | Schwarzmüller et al., 2014 |
| **6C7** | CAGL0J10582g | Verified | | Has domain(s) with predicted nucleic acid binding activity | ORF, Uncharacterized | PSP2 | CAGL0J10582g | Schwarzmüller et al., 2014 |
| **6C8** | CAGL0K02585g | No Deletion Verified | *YAP3* | bZIP domain-containing protein | ORF, Uncharacterized | YAP3 | CAGL0K02585g | Schwarzmüller et al., 2014 |
| **6C9** | CAGL0K03245g | Verified | | Protein of unknown function | ORF, Uncharacterized | | CAGL0K03245g | Schwarzmüller et al., 2014 |
| **6C10** | CAGL0K08668g | No Deletion Verified | *MET28* | bZIP domain-containing protein | ORF, Uncharacterized | MET28 | CAGL0K08668g | Schwarzmüller et al., 2014 |
| **6C11** | CAGL0K08690g | Verified | | Protein of unknown function, ortholog(s) expression directly regulated by the metabolic and meiotic transcriptional regulator Ume6p; overexpression causes a cell cycle delay or arrest | ORF, Uncharacterized | YIR016W | CAGL0K08690g | Schwarzmüller et al., 2014 |
| **6C12** | CAGL0K12320g | Verified | | Ortholog(s) have role in actin cortical patch assembly, barbed-end actin filament capping and actin cortical patch, membrane raft localization | ORF, Uncharacterized | AIM3 | CAGL0K12320g | Schwarzmüller et al., 2014 |
| **6D1** | CAGL0L01067g | No Deletion Verified | | Ortholog(s) have cytoplasm localization | ORF, Uncharacterized | PAR32 | CAGL0L01067g | Schwarzmüller et al., 2014 |
| **6D2** | CAGL0L03916g | No Deletion Verified | | Protein of unknown function | ORF, Uncharacterized | AZF1 | CAGL0L03916g | Schwarzmüller et al., 2014 |
| **6D3** | CAGL0L06226g | Verified | | Has domain(s) with predicted RNA binding, nucleic acid binding activity | ORF, Uncharacterized | HEK2 | CAGL0L06226g | Schwarzmüller et al., 2014 |
| **6D4** | CAGL0M08778g | Verified | *SWM1* | Ortholog(s) have ubiquitin protein ligase activity, role in anaphase-promoting complex assembly, ascospore wall assembly, regulation of mitotic metaphase/anaphase transition and anaphase-promoting complex, nucleus localization | ORF, Uncharacterized | SWM1 | CAGL0M08778g | Schwarzmüller et al., 2014 |
| **6D5** | CAGL0M12507g | Verified | *VHR1* | Transcriptional activator of genes involved in biotin metabolism; required for survival and proliferation in macrophages | ORF, Verified | VHR1 | CAGL0M12507g | Schwarzmüller et al., 2014 |
| **6D6** | CAGL0A03696g | No Deletion Verified | *GAL83* | Ortholog(s) have AMP-activated protein kinase activity, enzyme-substrate adaptor activity | ORF, Uncharacterized | GAL83 | CAGL0A03696g | Schwarzmüller et al., 2014 |
| **6D7** | CAGL0J04202g | No Deletion Verified | *HSP12* | Heat shock protein; gene is upregulated in azole-resistant strain; expression upregulated in biofilm vs planktonic cell culture | ORF, Verified | HSP12 | CAGL0J04202g | Schwarzmüller et al., 2014 |
| **6D8** | CAGL0M10659g | No Deletion Verified | | Ortholog(s) have structural constituent of chromatin activity, role in chromatin remodeling, positive regulation of transcription by RNA polymerase II and SWI/SNF complex, chromatin, cytosol, nucleus localization | ORF, Uncharacterized | SNF12 | CAGL0M10659g | Schwarzmüller et al., 2014 |
| **6D9** | CAGL0D01232g | No Deletion Verified | | Ortholog(s) have signal sequence binding activity, role in vacuolar transport and late endosome localization | ORF, Uncharacterized | MRL1 | CAGL0D01232g | Schwarzmüller et al., 2014 |
| **6D10** | CAGL0J04312g | Verified | | Ortholog(s) have signal sequence binding activity and role in Golgi to endosome transport, Golgi to vacuole transport, mitophagy, pexophagy, protein targeting to vacuole, vacuolar transport | ORF, Uncharacterized | PEP1 | CAGL0J04312g | Schwarzmüller et al., 2014 |
| **6D11** | CAGL0J06886g | No Deletion Verified | *VPS30* | Ortholog(s) have role in autophagy, cytoplasm to vacuole transport by the Cvt pathway, late endosome to vacuole transport, macroautophagy, pexophagy and phosphatidylinositol biosynthetic process, piecemeal microautophagy of the nucleus, retrograde transport, endosome to Golgi | ORF, Uncharacterized | VPS30 | CAGL0J06886g | Schwarzmüller et al., 2014 |
| **6D12** | CAGL0G09977g | No Deletion Verified | *GDB1* | Ortholog(s) have 4-alpha-glucanotransferase activity, amylo-alpha-1,6-glucosidase activity and role in glycogen catabolic process | ORF, Uncharacterized | GDB1 | CAGL0G09977g | Schwarzmüller et al., 2014 |
| **6E1** | CAGL0E05412g | No Deletion Verified | *KRE5* | UDP-glucose:glycoprotein glucosyltransferase that synthesizes 1,6-beta-D-glucan; required for proper cell wall morphology; null mutants are inviable; repressible mutants show ER stress, increased cell wall chitins, and abnormal cell shape | ORF, Verified | KRE5 | CAGL0E05412g | Schwarzmüller et al., 2014 |
| **6E2** | CAGL0G02717g | Verified | | Ortholog(s) have glucan 1,4-alpha-glucosidase activity, role in glycogen catabolic process and fungal-type vacuole localization | ORF, Uncharacterized | SGA1 | CAGL0G02717g | Schwarzmüller et al., 2014 |
| **6E3** | CAGL0J09922g | No Deletion Verified | | Ortholog(s) have cellular bud scar, fungal-type cell wall localization | ORF, Verified | SUN4 | CAGL0J09922g | Schwarzmüller et al., 2014 |
| **6E4** | CAGL0J02288g | Verified | *BAR1* | Ortholog(s) have aspartic-type endopeptidase activity, role in cellular response to pheromone, peptide catabolic process and extracellular region, fungal-type cell wall localization | ORF, Uncharacterized | BAR1 | CAGL0J02288g | Schwarzmüller et al., 2014 |
| **6E5** | CAGL0A02431g | Verified | *YPS7* | Putative aspartic protease; predicted GPI-anchor; expression induced at high temperature | ORF, Verified | YPS7 | CAGL0A02431g | Schwarzmüller et al., 2014 |
| **6E6** | CAGL0E06556g | Verified | | Ortholog(s) have inositol phosphosphingolipid phospholipase activity, mannosyl-inositol phosphorylceramide phospholipase activity and cell periphery, fungal-type vacuole, membrane localization | ORF, Uncharacterized | ISC1 | CAGL0E06556g | Schwarzmüller et al., 2014 |
| **6E7** | CAGL0L11154g | No Deletion Verified | | Ortholog(s) have lysophospholipase activity, role in phosphatidylcholine catabolic process, regulation of phospholipid biosynthetic process and endoplasmic reticulum localization | ORF, Uncharacterized | NTE1 | CAGL0L11154g | Schwarzmüller et al., 2014 |
| **6E8** | CAGL0A01177g | No Deletion Verified | | Ortholog(s) have phosphatidylinositol phospholipase C activity and role in inositol phosphate biosynthetic process, phospholipid catabolic process, protein localization to kinetochore, signal transduction involved in filamentous growth | ORF, Uncharacterized | PLC1 | CAGL0A01177g | Schwarzmüller et al., 2014 |
| **6E9** | CAGL0H03575g | Verified | | Ortholog(s) have role in ascospore-type prospore membrane formation, spindle pole body duplication, spore membrane bending pathway and endoplasmic reticulum, nucleus, prospore membrane localization | ORF, Uncharacterized | SPO1 | CAGL0H03575g | Schwarzmüller et al., 2014 |
| **6E10** | CAGL0L03135g | Verified | | Putative phospholipase D; gene is upregulated in azole-resistant strain | ORF, Uncharacterized | SPO14 | CAGL0L03135g | Schwarzmüller et al., 2014 |
| **6E11** | CAGL0M12815g | No Deletion Verified | | Ortholog(s) have role in positive regulation of protein sumoylation, regulation of synaptonemal complex assembly and condensed nuclear chromosome localization | ORF, Uncharacterized | SPO22 | CAGL0M12815g | Schwarzmüller et al., 2014 |
| **6E12** | CAGL0E04510g | Verified | | Ortholog(s) have phospholipase activity, role in cardiolipin metabolic process, phosphatidylethanolamine metabolic process, phospholipid metabolic process and mitochondrial matrix localization | ORF, Uncharacterized | DDL1 | CAGL0E04510g | Schwarzmüller et al., 2014 |
| **6F1** | CAGL0L08910g | Verified | | Ortholog(s) have role in mRNA metabolic process, mitochondrial translational initiation | ORF, Uncharacterized | AEP3 | CAGL0L08910g | Schwarzmüller et al., 2014 |
| **6F2** | CAGL0G03267g | Verified | | Ortholog(s) have role in protein targeting to membrane | ORF, Uncharacterized | AST2 | CAGL0G03267g | Schwarzmüller et al., 2014 |
| **6F3** | CAGL0H08558g | No Deletion Verified | *ATG11* | Protein with a predicted role in pexophagy | ORF, Uncharacterized | ATG11 | CAGL0H08558g | Schwarzmüller et al., 2014 |
| **6F4** | CAGL0G09999g | Verified | | Ortholog(s) have armadillo repeat domain binding, protein kinase regulator activity | ORF, Uncharacterized | ATG13 | CAGL0G09999g | Schwarzmüller et al., 2014 |
| **6F5** | CAGL0J07634g | No Deletion Verified | *SPO72* | Ortholog(s) have phosphatidylinositol-3-phosphate binding activity | ORF, Uncharacterized | ATG2 | CAGL0J07634g | Schwarzmüller et al., 2014 |
| **6F6** | CAGL0M00968g | No Deletion Verified | | Ortholog(s) have role in cytoplasm to vacuole transport by the Cvt pathway, nucleophagy, positive regulation of macroautophagy, protein localization to phagophore assembly site and phagophore assembly site localization | ORF, Uncharacterized | ATG23 | CAGL0M00968g | Schwarzmüller et al., 2014 |
| **6F7** | CAGL0D02464g | No Deletion Verified | | Has domain(s) with predicted phosphatase activity, phosphatidylinositol-3,5-bisphosphate 5-phosphatase activity and role in phosphatidylinositol dephosphorylation | ORF, Uncharacterized | FIG4 | CAGL0D02464g | Schwarzmüller et al., 2014 |
| **6F8** | CAGL0M02431g | Verified | *KES1* | Protein with a predicted role in ergosterol biosynthesis; protein abundance increased in ace2 mutant cells | ORF, Uncharacterized | KES1 | CAGL0M02431g | Schwarzmüller et al., 2014 |
| **6F9** | CAGL0M03707g | Verified | | Ortholog(s) have guanyl-nucleotide exchange factor activity, role in endocytosis, protein targeting to vacuole, retrograde transport, endosome to Golgi and early endosome membrane, endosome, trans-Golgi network localization | ORF, Uncharacterized | MON2 | CAGL0M03707g | Schwarzmüller et al., 2014 |
| **6F10** | CAGL0I09130g | Verified | | Ortholog(s) have role in response to amino acid and plasma membrane localization | ORF, Uncharacterized | PTR3 | CAGL0I09130g | Schwarzmüller et al., 2014 |
| **6F11** | CAGL0M08008g | No Deletion Verified | | Ortholog(s) have protein-containing complex binding activity and role in early endosome to late endosome transport, regulation of protein-containing complex assembly, vacuolar acidification | ORF, Uncharacterized | RAV1 | CAGL0M08008g | Schwarzmüller et al., 2014 |
| **6F12** | CAGL0I06116g | Verified | | Ortholog(s) have serine-type endopeptidase activity, role in protein autoprocessing, protein processing, response to amino acid and plasma membrane localization | ORF, Uncharacterized | SSY5 | CAGL0I06116g | Schwarzmüller et al., 2014 |
| **6G1** | CAGL0K12254g | Verified | | Ortholog(s) have role in negative regulation of gluconeogenesis, proteasome-mediated ubiquitin-dependent protein catabolic process, protein catabolic process in the vacuole, protein targeting to vacuole | ORF, Uncharacterized | VID24 | CAGL0K12254g | Schwarzmüller et al., 2014 |
| **6G2** | CAGL0H03993g | Verified | *CIT1* | Ortholog(s) have citrate (Si)-synthase activity, role in acetyl-CoA catabolic process, citrate metabolic process, tricarboxylic acid cycle and mitochondrion localization | ORF, Uncharacterized | CIT1 | CAGL0H03993g | Schwarzmüller et al., 2014 |
| **6G3** | CAGL0L06798g | Verified | | Ortholog(s) have L-malate dehydrogenase activity, mRNA binding activity, role in NADH regeneration, fatty acid beta-oxidation and peroxisomal matrix, peroxisome localization | ORF, Uncharacterized | MDH3 | CAGL0L06798g | Schwarzmüller et al., 2014 |
| **6G4** | CAGL0J03058g | Verified | | Predicted isocitrate lyase that converts isocitrate to glyoxylate and succinate in glyoxalate cycle; required for growth on acetate, ethanol or oleic acid; required for virulence in mouse | ORF, Uncharacterized | ICL1 | CAGL0J03058g | Schwarzmüller et al., 2014 |
| **6G5** | CAGL0F02431g | Verified | *ACO2* | Ortholog(s) have role in mitochondrial translation, tricarboxylic acid cycle and cytoplasm, mitochondrion, nucleus localization | ORF, Uncharacterized | ACO2 | CAGL0F02431g | Schwarzmüller et al., 2014 |
| **6G6** | CAGL0D06424g | Verified | *ACO1* | Putative aconitate hydratase | ORF, Verified | ACO1 | CAGL0D06424g | Schwarzmüller et al., 2014 |
| **6G7** | CAGL0L03982g | No Deletion Verified | *MLS1* | Ortholog(s) have malate synthase activity, role in glyoxylate cycle and cytosol, peroxisomal matrix, peroxisome localization | ORF, Uncharacterized | MLS1 | CAGL0L03982g | Schwarzmüller et al., 2014 |
| **6G8** | CAGL0E01705g | Verified | | Ortholog(s) have L-malate dehydrogenase activity, role in gluconeogenesis, protein import into peroxisome matrix and cytosol, nuclear periphery localization | ORF, Uncharacterized | MDH2 | CAGL0E01705g | Schwarzmüller et al., 2014 |
| **6G9** | CAGL0F08041g | Verified | *PFK1* | Putative phosphofructokinase, alpha subunit; increased protein abundance in azole resistant strain | ORF, Uncharacterized | PFK1 | CAGL0F08041g | Schwarzmüller et al., 2014 |
| **6G10** | CAGL0I02486g | No Deletion Verified | *ENO1* | Putative enolase I; protein abundance increased in azole resistant strain and in ace2 mutant cells | ORF, Verified | ENO2 | CAGL0I02486g | Schwarzmüller et al., 2014 |
| **6G11** | CAGL0H06633g | Verified | *PCK1* | Putative phosphoenolpyruvate carboxykinase; gene is downregulated in azole-resistant strain | ORF, Verified | PCK1 | CAGL0H06633g | Schwarzmüller et al., 2014 |
| **6G12** | CAGL0H04939g | No Deletion Verified | *FBP1* | Ortholog(s) have fructose 1,6-bisphosphate 1-phosphatase activity, role in gluconeogenesis, reactive oxygen species metabolic process and cytosol, periplasmic space localization | ORF, Uncharacterized | FBP1 | CAGL0H04939g | Schwarzmüller et al., 2014 |
| **6H1** | CAGL0L10758g | No Deletion Verified | *PFK2* | Putative 6-phosphofructokinase, beta subunit; protein abundance increased in ace2 mutant cells | ORF, Uncharacterized | PFK2 | CAGL0L10758g | Schwarzmüller et al., 2014 |
| **6H2** | CAGL0A03740g | No Deletion Verified | | Has domain(s) with predicted FAD binding, acyl-CoA oxidase activity, flavin adenine dinucleotide binding, oxidoreductase activity, acting on the CH-CH group of donors activity | ORF, Uncharacterized | POX1 | CAGL0A03740g | Schwarzmüller et al., 2014 |
| **6H3** | CAGL0L02167g | No Deletion Verified | | Ortholog(s) have 3-hydroxyacyl-CoA dehydrogenase activity, enoyl-CoA hydratase activity, role in fatty acid beta-oxidation and peroxisome localization | ORF, Uncharacterized | FOX2 | CAGL0L02167g | Schwarzmüller et al., 2014 |
| **6H4** | CAGL0H09460g | Verified | | Ortholog(s) have long-chain fatty acid-CoA ligase activity, medium-chain fatty acid-CoA ligase activity, very long-chain fatty acid-CoA ligase activity, role in long-chain fatty acid metabolic process and peroxisome localization | ORF, Uncharacterized | FAA2 | CAGL0H09460g | Schwarzmüller et al., 2014 |
| **6H5** | CAGL0H08437g | Verified | *VPS15* | Ortholog(s) have protein serine/threonine kinase activity, ubiquitin binding activity | ORF, Verified | VPS15 | CAGL0H08437g | Schwarzmüller et al., 2014 |
| **6H6** | CAGL0M08910g | Verified | *SNF1* | Putative serine/threonine protein kinase required for trehalose utilization | ORF, Verified | SNF1 | CAGL0M08910g | Schwarzmüller et al., 2014 |
| **6H7** | CAGL0B04147g | Verified | | Ortholog(s) have protein serine/threonine kinase activity, role in division septum assembly, exit from mitosis, mitotic cytokinesis, regulation of mRNA catabolic process and cellular bud neck, spindle pole body localization | ORF, Uncharacterized | DBF20 | CAGL0B04147g | Schwarzmüller et al., 2014 |
| **6H8** | CAGL0F03245g | Verified | *IRE1* | Putative protein kinase and endonuclease required for response to ER stress independently of Hac1p; involved in nonspecific degradation of ER-localized mRNAs but not in unfolded protein response via activation of Hac1p | ORF, Verified | IRE1 | CAGL0F03245g | Schwarzmüller et al., 2014 |
| **7A1** | CAGL0F09075g | No Deletion Verified | | Ortholog(s) have protein serine/threonine kinase activity | ORF, Uncharacterized | SCH9 | CAGL0F09075g | Schwarzmüller et al., 2014 |
| **7A2** | CAGL0G02607g | Verified | | Ortholog(s) have protein serine/threonine kinase activity | ORF, Uncharacterized | PRK1 | CAGL0G02607g | Schwarzmüller et al., 2014 |
| **7A3** | CAGL0M13167g | No Deletion Verified | | Ortholog(s) have protein serine/threonine kinase activity | ORF, Uncharacterized | ELM1 | CAGL0M13167g | Schwarzmüller et al., 2014 |

| | | | | | | | | |
|---|---|---|---|---|---|---|---|---|
| 7A4 | CAGL0K03399g | Verified | YPK2 | Ortholog(s) have protein serine/threonine kinase activity | ORF, Uncharacterized | YPK2 | CAGL0K03399g | Schwarzmüller et al., 2014 |
| 7A5 | CAGL0M03729g | No Deletion Verified | | Ortholog(s) have kinase activity, protein serine/threonine kinase activity | ORF, Uncharacterized | CLA4 | CAGL0M03729g | Schwarzmüller et al., 2014 |
| 7A6 | CAGL0F00913g | Verified | | Ortholog(s) have protein serine/threonine kinase activity and role in negative regulation of cytoplasmic translation, negative regulation of glycogen biosynthetic process, regulation of (1->6)-beta-D-glucan biosynthetic process | ORF, Uncharacterized | PSK2 | CAGL0F00913g | Schwarzmüller et al., 2014 |
| 7A7 | CAGL0D02244g | Verified | | Ortholog(s) have protein serine/threonine kinase activity | ORF, Uncharacterized | MEK1 | CAGL0D02244g | Schwarzmüller et al., 2014 |
| 7A8 | CAGL0K00693g | Verified | | Ortholog(s) have protein tyrosine kinase activity | ORF, Uncharacterized | SWE1 | CAGL0K00693g | Schwarzmüller et al., 2014 |
| 7A9 | CAGL0D01694g | Verified | | Ortholog(s) have MAP kinase activity, role in ascospore wall assembly, negative regulation of sporulation resulting in formation of a cellular spore and prospore membrane leading edge localization | ORF, Uncharacterized | SMK1 | CAGL0D01694g | Schwarzmüller et al., 2014 |
| 7A10 | CAGL0E01683g | Verified | YGK3 | Ortholog(s) have protein serine/threonine kinase activity and role in proteolysis | ORF, Uncharacterized | YGK3 | CAGL0E01683g | Schwarzmüller et al., 2014 |
| 7A11 | CAGL0I05192g | Verified | | Ortholog(s) have protein serine/threonine kinase activity and role in DNA damage response, donor selection, regulation of transcription by RNA polymerase I, regulation of transcription by RNA polymerase III | ORF, Uncharacterized | CKA1 | CAGL0I05192g | Schwarzmüller et al., 2014 |
| 7A12 | CAGL0G02035g | Verified | CKA2 | Catalytic subunit of casein kinase 2 (CK2), involved in regulation of sphingolipid biosynthesis | ORF, Uncharacterized | CKA2 | CAGL0G02035g | Schwarzmüller et al., 2014 |
| 7B1 | CAGL0A00275g | No Deletion Verified | | Ortholog(s) have protein kinase regulator activity, protein serine/threonine kinase inhibitor activity and role in DNA damage response, regulation of transcription by RNA polymerase I, regulation of transcription by RNA polymerase III | ORF, Uncharacterized | CKB1 | CAGL0A00275g | Schwarzmüller et al., 2014 |
| 7B2 | CAGL0I00946g | Verified | CKB2 | Ortholog(s) have protein kinase regulator activity, protein serine/threonine kinase activator activity, protein serine/threonine kinase inhibitor activity | ORF, Uncharacterized | CKB2 | CAGL0I00946g | Schwarzmüller et al., 2014 |
| 7B3 | CAGL0F03047g | No Deletion Verified | | Ortholog(s) have AMP-activated protein kinase activity, enzyme-substrate adaptor activity and role in regulation of protein-containing complex assembly, signal transduction | ORF, Uncharacterized | SIP1 | CAGL0F03047g | Schwarzmüller et al., 2014 |
| 7B4 | CAGL0L07326g | Verified | DUN1 | Ortholog(s) have protein kinase activity, protein serine/threonine kinase activity and role in DNA damage checkpoint signaling, double-strand break repair via nonhomologous end joining, replication fork processing | ORF, Uncharacterized | DUN1 | CAGL0L07326g | Schwarzmüller et al., 2014 |
| 7B5 | CAGL0K12496g | No Deletion Verified | | Ortholog(s) have cyclin-dependent protein kinase activating kinase activity, role in G2/M transition of mitotic cell cycle, meiotic cell cycle, negative regulation of transcription initiation by RNA polymerase II and cytoplasm localization | ORF, Uncharacterized | CAK1 | CAGL0K12496g | Schwarzmüller et al., 2014 |
| 7B6 | CAGL0H07535g | No Deletion Verified | | Ortholog(s) have RNA polymerase II complex binding, cyclin-dependent protein kinase activity, cyclin-dependent protein serine/threonine kinase activity, histone binding, protein kinase activity, protein serine/threonine kinase activity | ORF, Uncharacterized | CDC28 | CAGL0H07535g | Schwarzmüller et al., 2014 |
| 7B7 | CAGL0L12474g | No Deletion Verified | PHO85 | Ortholog(s) have cyclin-dependent protein serine/threonine kinase activity | ORF, Uncharacterized | PHO85 | CAGL0L12474g | Schwarzmüller et al., 2014 |
| 7B8 | CAGL0K10604g | Verified | | Ortholog(s) have calmodulin-dependent protein kinase activity, role in signal transduction and cytoplasm localization | ORF, Uncharacterized | CMK1 | CAGL0K10604g | Schwarzmüller et al., 2014 |
| 7B9 | CAGL0F04741g | Verified | | Ortholog(s) have calmodulin binding, calmodulin-dependent protein kinase activity, protein kinase activity | ORF, Uncharacterized | CMK2 | CAGL0F04741g | Schwarzmüller et al., 2014 |
| 7B10 | CAGL0K12342g | No Deletion Verified | | Ortholog(s) have beta-1,4-mannosyltransferase activity, role in oligosaccharide-lipid intermediate biosynthetic process, protein N-linked glycosylation and endoplasmic reticulum localization | ORF, Uncharacterized | ALG1 | CAGL0K12342g | Schwarzmüller et al., 2014 |
| 7B11 | CAGL0D01122g | No Deletion Verified | | Ortholog(s) have GDP-Man:Man3GlcNAc2-PP-Dol alpha-1,2-mannosyltransferase activity, alpha-1,2-mannosyltransferase activity | ORF, Uncharacterized | ALG11 | CAGL0D01122g | Schwarzmüller et al., 2014 |
| 7B12 | CAGL0M05731g | No Deletion Verified | | Ortholog(s) have GDP-Man:Man1GlcNAc2-PP-Dol alpha-1,3-mannosyltransferase activity, glycolipid 1,6-alpha-mannosyltransferase activity and role in oligosaccharide-lipid intermediate biosynthetic process | ORF, Verified | ALG2 | CAGL0M05731g | Schwarzmüller et al., 2014 |
| 7C1 | CAGL0E06028g | Verified | ALG5 | Putative glucosyltransferase involved in N-linked glycosylation and vesicular trafficking; null mutant shows mild sensitivity to calcineurin inhibitors | ORF, Uncharacterized | ALG5 | CAGL0E06028g | Schwarzmüller et al., 2014 |
| 7C2 | CAGL0E02629g | Verified | ALG6 | Putative glucosyltransferase involved in N-linked glycosylation and vesicular trafficking; null mutant shows mild sensitivity to calcineurin inhibitors and elevated expression of RCN2 | ORF, Uncharacterized | ALG6 | CAGL0E02629g | Schwarzmüller et al., 2014 |
| 7C3 | CAGL0C01727g | No Deletion Verified | | Ortholog(s) have UDP-N-acetylglucosamine-dolichyl-phosphate N-acetylglucosaminephosphotransferase activity and role in aerobic respiration, dolichol-linked oligosaccharide biosynthetic process, protein N-linked glycosylation | ORF, Uncharacterized | ALG7 | CAGL0C01727g | Schwarzmüller et al., 2014 |
| 7C4 | CAGL0L06556g | No Deletion Verified | | Ortholog(s) have dolichyl pyrophosphate Glc2Man9GlcNAc2 alpha-1,2-glucosyltransferase activity, role in protein N-linked glycosylation and endoplasmic reticulum membrane localization | ORF, Uncharacterized | DIE2 | CAGL0L06556g | Schwarzmüller et al., 2014 |
| 7C5 | CAGL0G09955g | No Deletion Verified | | Ortholog(s) have dolichyl-phosphate beta-D-mannosyltransferase activity and role in GPI anchor biosynthetic process, dolichol-linked oligosaccharide biosynthetic process | ORF, Uncharacterized | DPM1 | CAGL0G09955g | Schwarzmüller et al., 2014 |
| 7C6 | CAGL0D05742g | No Deletion Verified | | Has domain(s) with predicted role in protein glycosylation and membrane localization | ORF, Uncharacterized | OST1 | CAGL0D05742g | Schwarzmüller et al., 2014 |
| 7C7 | CAGL0B04499g | No Deletion Verified | | Ortholog(s) have role in protein N-linked glycosylation and oligosaccharyltransferase complex localization | ORF, Uncharacterized | OST2 | CAGL0B04499g | Schwarzmüller et al., 2014 |
| 7C8 | CAGL0J08569g | Verified | | Ortholog(s) have dolichyl-diphosphooligosaccharide-protein glycotransferase activity, protein-disulfide reductase activity | ORF, Uncharacterized | OST3 | CAGL0J08569g | Schwarzmüller et al., 2014 |
| 7C9 | CAGL0J08294g | No Deletion Verified | | Ortholog(s) have borate efflux transmembrane transporter activity, role in borate transport, protein targeting to vacuole and fungal-type vacuole, plasma membrane localization | ORF, Uncharacterized | BOR1 | CAGL0J08294g | Schwarzmüller et al., 2014 |
| 7C10 | CAGL0J00627g | No Deletion Verified | | Ortholog(s) have role in vacuolar acidification and membrane localization | ORF, Uncharacterized | VMA16 | CAGL0J00627g | Schwarzmüller et al., 2014 |
| 7C11 | CAGL0G07040g | Verified | | Ortholog(s) have protein-disulfide reductase activity, role in protein N-linked glycosylation, protein-containing complex assembly and oligosaccharyltransferase complex localization | ORF, Uncharacterized | OST6 | CAGL0G07040g | Schwarzmüller et al., 2014 |
| 7C12 | CAGL0A04587g | Verified | | Ortholog(s) have dol-P-Man:Man(5)GlcNAc(2)-PP-Dol alpha-1,3-mannosyltransferase activity | ORF, Uncharacterized | ALG3 | CAGL0A04587g | Schwarzmüller et al., 2014 |
| 7D1 | CAGL0A00209g | No Deletion Verified | | Ortholog(s) have dolichyl-diphosphooligosaccharide-protein glycotransferase activity, role in protein N-linked glycosylation and oligosaccharyltransferase complex localization | ORF, Uncharacterized | STT3 | CAGL0A00209g | Schwarzmüller et al., 2014 |
| 7D2 | CAGL0H09196g | No Deletion Verified | VRG4 | GDP-mannose transporter involved in glycosylation in the Golgi | ORF, Verified | VRG4 | CAGL0H09196g | Schwarzmüller et al., 2014 |
| 7D3 | CAGL0E04180g | No Deletion Verified | | Has domain(s) with predicted role in dolichol-linked oligosaccharide biosynthetic process | ORF, Uncharacterized | ALG14 | CAGL0E04180g | Schwarzmüller et al., 2014 |
| 7D4 | CAGL0D06270g | No Deletion Verified | | Ortholog(s) have N-acetylglucosaminyldiphosphodolichol N-acetylglucosaminyltransferase activity, role in dolichol-linked oligosaccharide biosynthetic process and UDP-N-acetylglucosamine transferase complex, cytoplasm, cytosol localization | ORF, Uncharacterized | ALG13 | CAGL0D06270g | Schwarzmüller et al., 2014 |
| 7D5 | CAGL0L01331g | Verified | ANP1 | Alpha-1,6-mannosyltransferase with a role in protein N-linked glycosylation in Golgi | ORF, Verified | ANP1 | CAGL0L01331g | Schwarzmüller et al., 2014 |
| 7D6 | CAGL0K04103g | No Deletion Verified | | Ortholog(s) have dolichyldiphosphatase activity, role in lipid biosynthetic process, protein N-linked glycosylation and endoplasmic reticulum membrane localization | ORF, Uncharacterized | CAX4 | CAGL0K04103g | Schwarzmüller et al., 2014 |
| 7D7 | CAGL0M10769g | No Deletion Verified | | Ortholog(s) have alpha-1,6-mannosyltransferase activity and role in dolichol-linked oligosaccharide biosynthetic process, protein glycosylation | ORF, Uncharacterized | ALG12 | CAGL0M10769g | Schwarzmüller et al., 2014 |
| 7D8 | CAGL0I09922g | Verified | | Ortholog(s) have acetylglucosaminyltransferase activity, role in protein N-linked glycosylation and Golgi medial cisterna localization | ORF, Uncharacterized | GNT1 | CAGL0I09922g | Schwarzmüller et al., 2014 |
| 7D9 | CAGL0H09130g | Verified | | Ortholog(s) have enzyme activator activity, role in protein N-linked glycosylation, protein O-linked glycosylation and Golgi apparatus localization | ORF, Uncharacterized | MNN4 | CAGL0H09130g | Schwarzmüller et al., 2014 |
| 7D10 | CAGL0L12804g | No Deletion Verified | | Ortholog(s) have alpha-1,6-mannosyltransferase activity, mannosyltransferase activity, role in protein N-linked glycosylation and Golgi apparatus, cis-Golgi network, mannan polymerase II complex, mannan polymerase complex localization | ORF, Uncharacterized | MNN9 | CAGL0L12804g | Schwarzmüller et al., 2014 |
| 7D11 | CAGL0M00528g | Verified | | Ortholog(s) have mannosyl-oligosaccharide 1,2-alpha-mannosidase activity and role in protein deglycosylation involved in glycoprotein catabolic process, ubiquitin-dependent ERAD pathway, ubiquitin-dependent glycoprotein ERAD pathway | ORF, Uncharacterized | MNS1 | CAGL0M00528g | Schwarzmüller et al., 2014 |
| 7D12 | CAGL0J04378g | No Deletion Verified | | Ortholog(s) have role in glycolipid translocation and endoplasmic reticulum membrane localization | ORF, Uncharacterized | RFT1 | CAGL0J04378g | Schwarzmüller et al., 2014 |
| 7E1 | CAGL0J08778g | Verified | | Ortholog(s) have structural molecule activity and role in COPII-coated vesicle budding, nuclear pore localization, positive regulation of DNA-templated transcription, positive regulation of TORC1 signaling, ubiquitin-dependent ERAD pathway | ORF, Uncharacterized | SEC13 | CAGL0J08778g | Schwarzmüller et al., 2014 |
| 7E2 | CAGL0M07700g | No Deletion Verified | | Ortholog(s) have dolichol kinase activity, role in dolichyl monophosphate biosynthetic process and endoplasmic reticulum membrane localization | ORF, Uncharacterized | SEC59 | CAGL0M07700g | Schwarzmüller et al., 2014 |
| 7E3 | CAGL0G07887g | Verified | | Ortholog(s) have role in protein-containing complex assembly and endoplasmic reticulum membrane localization | ORF, Verified | VOA1 | CAGL0G07887g | Schwarzmüller et al., 2014 |
| 7E4 | CAGL0I00704g | No Deletion Verified | | Ortholog(s) have structural molecule activity, role in protein N-linked glycosylation and endoplasmic reticulum membrane, oligosaccharyltransferase complex localization | ORF, Uncharacterized | SWP1 | CAGL0I00704g | Schwarzmüller et al., 2014 |
| 7E5 | CAGL0I08063g | No Deletion Verified | | Ortholog(s) have RNA binding, double-stranded DNA binding, protein-containing complex binding activity | ORF, Uncharacterized | THP1 | CAGL0I08063g | Schwarzmüller et al., 2014 |
| 7E6 | CAGL0L08646g | No Deletion Verified | | Ortholog(s) have cysteine-type peptidase activity, deSUMOylase activity, protein-containing complex binding activity, role in G2/M transition of mitotic cell cycle, protein desumoylation and nuclear envelope, nucleolus localization | ORF, Uncharacterized | ULP1 | CAGL0L08646g | Schwarzmüller et al., 2014 |
| 7E7 | CAGL0B02321g | No Deletion Verified | | Ortholog(s) have alpha-1,6-mannosyltransferase activity and role in cell wall mannoprotein biosynthetic process, fungal-type cell wall organization, mannose biosynthetic process, protein N-linked glycosylation | ORF, Uncharacterized | VAN1 | CAGL0B02321g | Schwarzmüller et al., 2014 |
| 7E8 | CAGL0L02365g | No Deletion Verified | | Ortholog(s) have glycosyltransferase activity, oligosaccharyl transferase activity and role in protein N-linked glycosylation | ORF, Uncharacterized | WBP1 | CAGL0L02365g | Schwarzmüller et al., 2014 |
| 7E9 | CAGL0C04048g | Verified | | Has domain(s) with predicted glycosyltransferase activity and role in protein glycosylation | ORF, Uncharacterized | MNT3 | CAGL0C04048g | Schwarzmüller et al., 2014 |
| 7E10 | CAGL0L07216g | Verified | | Ortholog(s) have dolichyl-phosphate-mannose-protein mannosyltransferase activity | ORF, Uncharacterized | PMT1 | CAGL0L07216g | Schwarzmüller et al., 2014 |
| 7E11 | CAGL0J08734g | Verified | PMT2 | Ortholog(s) have dolichyl-phosphate-mannose-protein mannosyltransferase activity | ORF, Uncharacterized | PMT2 | CAGL0J08734g | Schwarzmüller et al., 2014 |
| 7E12 | CAGL0M00220g | Verified | | Ortholog(s) have dolichyl-phosphate-mannose-protein mannosyltransferase activity | ORF, Uncharacterized | PMT4 | CAGL0M00220g | Schwarzmüller et al., 2014 |
| 7F1 | CAGL0K00979g | No Deletion Verified | | Ortholog(s) have role in protein O-linked glycosylation, protein O-linked mannosylation | ORF, Uncharacterized | PMT6 | CAGL0K00979g | Schwarzmüller et al., 2014 |
| 7F2 | CAGL0A00429g | No Deletion Verified | ERG4 | Putative C24 sterol reductase | ORF, Uncharacterized | ERG4 | CAGL0A00429g | Schwarzmüller et al., 2014 |
| 7F3 | CAGL0F01793g | No Deletion Verified | ERG3 | Delta 5,6 sterol desaturase; C-5 sterol desaturase; predicted transmembrane domain and endolasmic reticulum (ER) binding motif; gene used for molecular typing of *C. glabrata* strain isolates | ORF, Uncharacterized | ERG3 | CAGL0F01793g | Schwarzmüller et al., 2014 |
| 7F4 | CAGL0L11506g | No Deletion Verified | HMG1 | Putative hydroxymethylglutaryl-CoA reductase; orthologs catalyze conversion of HMG-CoA to mevalonate, an early step in the ergosterol biosynthetic pathway | ORF, Verified | HMG1 | CAGL0L11506g | Schwarzmüller et al., 2014 |
| 7F5 | CAGL0H04653g | No Deletion Verified | ERG6 | C24 sterol methyltransferase; mutation confers resistance to amphotericin B and nystatin and increased sensitivity to antifungal and cell wall-affecting drugs | ORF, Verified | ERG6 | CAGL0H04653g | Schwarzmüller et al., 2014 |
| 7F6 | CAGL0M07656g | Verified | ERG5 | Putative C22 sterol desaturase | ORF, Verified | ERG5 | CAGL0M07656g | Schwarzmüller et al., 2014 |
| 7F7 | CAGL0L10714g | No Deletion Verified | ERG2 | C-8 sterol isomerase | ORF, Uncharacterized | ERG2 | CAGL0L10714g | Schwarzmüller et al., 2014 |
| 7F8 | CAGL0I02970g | No Deletion Verified | ERG24 | Ortholog(s) have delta14-sterol reductase activity and role in ergosterol biosynthetic process | ORF, Uncharacterized | ERG24 | CAGL0I02970g | Schwarzmüller et al., 2014 |
| 7F9 | CAGL0G09515g | Verified | | Ortholog(s) have glucan exo-1,3-beta-glucosidase activity, role in ascospore formation and ascospore wall, fungal-type cell wall localization | ORF, Verified | SPR1 | CAGL0G09515g | Schwarzmüller et al., 2014 |
| 7F10 | CAGL0M13189g | No Deletion Verified | MSN4 | Putative transcription factor similar to S. cerevisiae Msn4p; involved in response to oxidative stress | ORF, Verified | MSN4 | CAGL0M13189g | Schwarzmüller et al., 2014 |
| 7F11 | CAGL0H03487g | No Deletion Verified | AFT1 | Putative RNA polymerase II transcription factor, involved in regulation of iron acquisition genes; required for growth under iron depletion | ORF, Verified | AFT1 | CAGL0H03487g | Schwarzmüller et al., 2014 |
| 7F12 | CAGL0G09042g | Verified | | Ortholog(s) have DNA-binding transcription factor activity, RNA polymerase II-specific, RNA polymerase II cis-regulatory region sequence-specific DNA binding, iron-sulfur cluster binding activity | ORF, Uncharacterized | AFT2 | CAGL0G09042g | Schwarzmüller et al., 2014 |
| 7G1 | CAGL0E04092g | No Deletion Verified | SIT1 | Putative siderophore-iron transporter with 14 transmembrane domains; required for iron-dependent survival in macrophages; mRNA levels elevated under iron deficiency conditions; plasma membrane localized | ORF, Verified | ARN1 | CAGL0E04092g | Schwarzmüller et al., 2014 |
| 7G2 | CAGL0F06413g | No Deletion Verified | FET3 | Putative copper ferroxidase involved in iron uptake | ORF, Verified | FET3 | CAGL0F06413g | Schwarzmüller et al., 2014 |
| 7G3 | CAGL0F00187g | No Deletion Verified | FET4 | Has domain(s) with predicted role in transmembrane transport | ORF, Uncharacterized | FET4 | CAGL0F00187g | Schwarzmüller et al., 2014 |
| 7G4 | CAGL0K12738g | Verified | | Ortholog(s) have ferroxidase activity, role in iron ion transport and fungal-type vacuole membrane, membrane raft localization | ORF, Uncharacterized | FET5 | CAGL0K12738g | Schwarzmüller et al., 2014 |
| 7G5 | CAGL0C03333g | Verified | | Ortholog(s) have ferric-chelate reductase activity, role in cellular response to iron ion starvation, copper ion import, reductive iron assimilation and fungal-type vacuole membrane localization | ORF, Uncharacterized | FRE6 | CAGL0C03333g | Schwarzmüller et al., 2014 |
| 7G6 | CAGL0M07942g | Verified | | Ortholog(s) have oxidoreductase activity, acting on metal ions activity and role in intracellular monoatomic cation homeostasis | ORF, Uncharacterized | FRE8 | CAGL0M07942g | Schwarzmüller et al., 2014 |
| 7G7 | CAGL0K05863g | No Deletion Verified | NOX1 | Putative superoxide-generating NAD(P)H oxidase; null mutant is hypersensitive to oxidative stress and fails to induce human transglutaminase 2 activity or reactive oxygen species in hepatocytes | ORF, Verified | AIM14 | CAGL0K05863g | Schwarzmüller et al., 2014 |
| 7G8 | CAGL0M06281g | No Deletion Verified | DTR1 | Acetate exporter in the plasma membrane, required for virulence in Galleria mellonella model | ORF, Verified | DTR1 | CAGL0M06281g | Schwarzmüller et al., 2014 |
| 7G10 | CAGL0L10868g | Verified | | Ortholog(s) have role in cellular response to anoxia | ORF, Uncharacterized | FSF1 | CAGL0L10868g | Schwarzmüller et al., 2014 |

| | | | | | | | | |
|---|---|---|---|---|---|---|---|---|
| **7G11** | CAGL0M05511g | Verified | *FTH1* | Has domain(s) with predicted iron ion transmembrane transporter activity, role in iron ion transmembrane transport and high-affinity iron permease complex localization | ORF, Uncharacterized | *FTH1* | CAGL0M05511g | Schwarzmüller et al., 2014 |
| **7G12** | CAGL0I06743g | No Deletion Verified | *FTR1* | Putative ferrous iron transmembrane transporter involved in iron uptake | ORF, Uncharacterized | *FTR1* | CAGL0I06743g | Schwarzmüller et al., 2014 |
| **7H1** | CAGL0G03905g | No Deletion Verified | | Ortholog(s) have 2 iron, 2 sulfur cluster binding, iron ion binding, iron-sulfur cluster binding, iron-sulfur transferase activity | ORF, Uncharacterized | *ISA1* | CAGL0G03905g | Schwarzmüller et al., 2014 |
| **7H2** | CAGL0D01496g | Verified | | Ortholog(s) have iron ion binding activity and role in biotin biosynthetic process, iron-sulfur cluster assembly, protein maturation by [4Fe-4S] cluster transfer, protein maturation by iron-sulfur cluster transfer | ORF, Uncharacterized | *ISA2* | CAGL0D01496g | Schwarzmüller et al., 2014 |
| **7H3** | CAGL0H08822g | No Deletion Verified | *MMT2* | Ortholog(s) have role in intracellular iron ion homeostasis and mitochondrion localization | ORF, Uncharacterized | *MMT1* | CAGL0H08822g | Schwarzmüller et al., 2014 |
| **7H4** | CAGL0E06006g | Verified | | Ortholog(s) have role in intracellular iron ion homeostasis and mitochondrion localization | ORF, Uncharacterized | *MMT2* | CAGL0E06006g | Schwarzmüller et al., 2014 |
| **7H9** | CAGL0A03476g | Verified | *SMF3* | Ortholog(s) have role in intracellular iron ion homeostasis, iron ion transport and fungal-type vacuole membrane localization | ORF, Uncharacterized | *SMF3* | CAGL0A03476g | Schwarzmüller et al., 2014 |
| **7H10** | CAGL0J08481g | Verified | | Ortholog(s) have role in meiotic cell cycle, synaptonemal complex organization | ORF, Uncharacterized | *GMC1* | CAGL0J08481g | Schwarzmüller et al., 2014 |
| **7H11** | CAGL0M05643g | Verified | *YFH1* | Ortholog(s) have mitochondrion localization | ORF, Uncharacterized | *YFH1* | CAGL0M05643g | Schwarzmüller et al., 2014 |
| **8A1** | CAGL0A01738g | No Deletion Verified | *OCH1* | Putative alpha-1,6-mannosyltransferase of cis-Golgi apparatus, involved in cell wall maintenance; gene is upregulated in azole-resistant strain | ORF, Verified | *OCH1* | CAGL0A01738g | Schwarzmüller et al., 2014 |
| **8A2** | CAGL0G08822g | No Deletion Verified | | Ortholog(s) have role in RAM/MOR signaling pathway, budding cell apical bud growth, cell budding, cell morphogenesis, regulation of establishment or maintenance of bipolar cell polarity regulating cell shape | ORF, Uncharacterized | *TAO3* | CAGL0G08822g | Schwarzmüller et al., 2014 |
| **8A3** | CAGL0K05005g | No Deletion Verified | | Ortholog(s) have dol-P-Man:Man(6)GlcNAc(2)-PP-Dol alpha-1,2-mannosyltransferase activity, dol-P-Man:Man(8)GlcNAc(2)-PP-Dol alpha-1,2-mannosyltransferase activity, mannosyltransferase activity | ORF, Uncharacterized | *ALG9* | CAGL0K05005g | Schwarzmüller et al., 2014 |
| **8A4** | CAGL0I04774g | No Deletion Verified | | Ortholog(s) have calcium-release channel activity, enzyme regulator activity | ORF, Uncharacterized | *CSG2* | CAGL0I04774g | Schwarzmüller et al., 2014 |
| **8A5** | CAGL0M05621g | No Deletion Verified | | Ortholog(s) have inositol phosphorylceramide mannosyltransferase activity and role in glycosphingolipid biosynthetic process, mannosyl-inositol phosphorylceramide biosynthetic process, sphingolipid biosynthetic process | ORF, Uncharacterized | *CSH1* | CAGL0M05621g | Schwarzmüller et al., 2014 |
| **8A6** | CAGL0B02233g | No Deletion Verified | | Ortholog(s) have phosphatidylinositol N-acetylglucosaminyltransferase activity and role in GPI anchor biosynthetic process | ORF, Uncharacterized | *GPI1* | CAGL0B02233g | Schwarzmüller et al., 2014 |
| **8A7** | CAGL0F07843g | No Deletion Verified | | Ortholog(s) have alpha-1,2-mannosyltransferase activity and role in GPI anchor biosynthetic process | ORF, Uncharacterized | *GPI10* | CAGL0F07843g | Schwarzmüller et al., 2014 |
| **8A8** | CAGL0H01485g | No Deletion Verified | | Ortholog(s) have mannose-ethanolamine phosphotransferase activity and role in GPI anchor biosynthetic process | ORF, Uncharacterized | *GPI11* | CAGL0H01485g | Schwarzmüller et al., 2014 |
| **8A9** | CAGL0M03047g | No Deletion Verified | | Ortholog(s) have N-acetylglucosaminylphosphatidylinositol deacetylase activity | ORF, Uncharacterized | *GPI12* | CAGL0M03047g | Schwarzmüller et al., 2014 |
| **8A10** | CAGL0G04015g | No Deletion Verified | | Ortholog(s) have mannose-ethanolamine phosphotransferase activity, transferase activity, transferring phosphorus-containing groups activity and role in GPI anchor biosynthetic process | ORF, Uncharacterized | *GPI13* | CAGL0G04015g | Schwarzmüller et al., 2014 |
| **8A11** | CAGL0B03905g | No Deletion Verified | | Ortholog(s) have mannosyltransferase activity, role in GPI anchor biosynthetic process, fungal-type cell wall organization and glycosylphosphatidylinositol-mannosyltransferase I complex localization | ORF, Uncharacterized | *GPI14* | CAGL0B03905g | Schwarzmüller et al., 2014 |
| **8A12** | CAGL0J01848g | No Deletion Verified | | Ortholog(s) have RNA polymerase I activity, role in nucleolar large rRNA transcription by RNA polymerase I, transcription by RNA polymerase I and RNA polymerase I complex localization | ORF, Uncharacterized | *RPA135* | CAGL0J01848g | Schwarzmüller et al., 2014 |
| **8B1** | CAGL0M13453g | No Deletion Verified | | Ortholog(s) have role in attachment of GPI anchor to protein and GPI-anchor transamidase complex, endoplasmic reticulum membrane localization | ORF, Uncharacterized | *GPI16* | CAGL0M13453g | Schwarzmüller et al., 2014 |
| **8B2** | CAGL0F03179g | No Deletion Verified | | Ortholog(s) have role in attachment of GPI anchor to protein and GPI-anchor transamidase complex, endoplasmic reticulum, membrane, nuclear inner membrane localization | ORF, Uncharacterized | *GPI17* | CAGL0F03179g | Schwarzmüller et al., 2014 |
| **8B3** | CAGL0C04235g | No Deletion Verified | | Ortholog(s) have alpha-1,6-mannosyltransferase activity, mannosyltransferase activity, role in GPI anchor biosynthetic process and endoplasmic reticulum, endoplasmic reticulum membrane, mannosyltransferase complex localization | ORF, Uncharacterized | *GPI18* | CAGL0C04235g | Schwarzmüller et al., 2014 |
| **8B4** | CAGL0J10604g | No Deletion Verified | | Ortholog(s) have UDP-glycosyltransferase activity, role in GPI anchor biosynthetic process and endoplasmic reticulum membrane, glycosylphosphatidylinositol-N-acetylglucosaminyltransferase (GPI-GnT) complex localization | ORF, Uncharacterized | *GPI19* | CAGL0J10604g | Schwarzmüller et al., 2014 |
| **8B5** | CAGL0H05401g | No Deletion Verified | | Ortholog(s) have role in GPI anchor biosynthetic process and glycosylphosphatidylinositol-N-acetylglucosaminyltransferase (GPI-GnT) complex localization | ORF, Uncharacterized | *GPI2* | CAGL0H05401g | Schwarzmüller et al., 2014 |
| **8B6** | CAGL0M01298g | No Deletion Verified | | Ortholog(s) have GPI-anchor transamidase activity, role in attachment of GPI anchor to protein and GPI-anchor transamidase complex, endoplasmic reticulum membrane localization | ORF, Uncharacterized | *GPI8* | CAGL0M01298g | Schwarzmüller et al., 2014 |
| **8B7** | CAGL0G07491g | No Deletion Verified | *MNN11* | Alpha-1,6-mannosyltransferase with a role in protein glycosylation | ORF, Verified | *MNN11* | CAGL0G07491g | Schwarzmüller et al., 2014 |
| **8B8** | CAGL0K06897g | No Deletion Verified | | Protein of unknown function | ORF, Uncharacterized | *YBR225W* | CAGL0K06897g | Schwarzmüller et al., 2014 |
| **8B9** | CAGL0M11572g | No Deletion Verified | | Protein of unknown function, mutant in ortholog(s) is deficient in cell wall mannosylphosphate and has long chronological lifespan | ORF, Uncharacterized | *LCL2* | CAGL0M11572g | Schwarzmüller et al., 2014 |
| **8B10** | CAGL0M04323g | No Deletion Verified | *ACE2* | Putative transcription factor; null mutation results in hypervirulence in immunocompromised mice | ORF, Verified | *ACE2* | CAGL0M04323g | Schwarzmüller et al., 2014 |
| **8B11** | CAGL0E01331g | No Deletion Verified | *SWI5* | Transcription factor; mutants display increased fungal burdens in mouse lungs and brain | ORF, Uncharacterized | *SWI5* | CAGL0E01331g | Schwarzmüller et al., 2014 |
| **8B12** | CAGL0G05423g | No Deletion Verified | *HO* | Putative endonuclease with a predicted role in mating-type switching | ORF, Uncharacterized | *HO* | CAGL0G05423g | Schwarzmüller et al., 2014 |
| **8C1** | CAGL0E06600g | Verified | | Putative adhesin-like protein; belongs to adhesin cluster V | ORF, Uncharacterized | *FLO5* | CAGL0E06600g | Schwarzmüller et al., 2014 |
| **8C2** | CAGL0E06644g | Verified | *EPA1* | Sub-telomerically encoded adhesin with a role in cell adhesion; GPI-anchored cell wall protein; N-terminal ligand binding domain binds to ligands containing a terminal galactose residue; belongs to adhesin cluster I | ORF, Verified | | CAGL0E06644g | Schwarzmüller et al., 2014 |
| **8C3** | CAGL0E06688g | No Deletion Verified | *EPA3* | Epithelial adhesion protein, involved in biofilm formation and azole drug resistance; belongs to adhesin cluster I; GPI-anchored | ORF, Verified | | CAGL0E06688g | Schwarzmüller et al., 2014 |
| **8C4** | CAGL0I00220g | No Deletion Verified | *EPA23* | Predicted GPI-linked adhesin-like protein; belongs to adhesin cluster I | ORF, Uncharacterized | | CAGL0I00220g | Schwarzmüller et al., 2014 |
| **8C5** | CAGL0L12980g | Verified | *SET1* | Histone methyltransferase (H3-K4 specific), subunit of the COMPASS (Set1C) complex; involved in regulation of drug resistance genes | ORF, Verified | *SET1* | CAGL0L12980g | Schwarzmüller et al., 2014 |
| **8C7** | CAGL0C00297g | Verified | *SET2* | Ortholog(s) have RNA binding, histone H3K36 methyltransferase activity | ORF, Uncharacterized | *SET2* | CAGL0C00297g | Schwarzmüller et al., 2014 |
| **8C8** | CAGL0L03091g | Verified | | Ortholog(s) have methylated histone binding activity, role in negative regulation of meiotic nuclear division, protein methylation, regulation of DNA-templated transcription and Set3 complex localization | ORF, Uncharacterized | *SET3* | CAGL0L03091g | Schwarzmüller et al., 2014 |
| **8C9** | CAGL0C05511g | No Deletion Verified | | Ortholog(s) have alkaline phosphatase activity, phosphatase activity, phosphoglycolate phosphatase activity, phosphoprotein phosphatase activity and role in carbohydrate metabolic process | ORF, Uncharacterized | *PHO13* | CAGL0C05511g | Schwarzmüller et al., 2014 |
| **8C10** | CAGL0G06556g | Verified | *PHO23* | Ortholog(s) have methylated histone binding activity | ORF, Uncharacterized | *PHO23* | CAGL0G06556g | Schwarzmüller et al., 2014 |
| **8C11** | CAGL0D01430g | No Deletion Verified | | Ortholog(s) have protein lysine deacetylase activity, role in positive regulation of transcription by RNA polymerase II and histone deacetylase complex localization | ORF, Uncharacterized | *HOS1* | CAGL0D01430g | Schwarzmüller et al., 2014 |
| **8C12** | CAGL0A03322g | No Deletion Verified | *HOS2* | Putative NAD-dependent histone deacetylase; subunit of the Set3 and Rpd3L complexes | ORF, Uncharacterized | *HOS2* | CAGL0A03322g | Schwarzmüller et al., 2014 |
| **8D1** | CAGL0J06974g | No Deletion Verified | | Ortholog(s) have histone deacetylase activity and role in positive regulation of transcription by RNA polymerase II, regulation of transcription by RNA polymerase II | ORF, Uncharacterized | *HOS3* | CAGL0J06974g | Schwarzmüller et al., 2014 |
| **8D2** | CAGL0C05357g | Verified | *HST1* | Histone deacetylase; ortholog of *S. cerevisiae* SIR2; sensor of niacin limitation; regulates gene expression under niacin-limiting conditions | ORF, Verified | *HST1* | CAGL0C05357g | Schwarzmüller et al., 2014 |
| **8D3** | CAGL0K01463g | Verified | *SIR2* | Putative NAD+ dependent histone deacetylase of the Sirtuin family, involved in subtelomeric silencing | ORF, Verified | *SIR2* | CAGL0K01463g | Schwarzmüller et al., 2014 |
| **8D4** | CAGL0M00770g | Verified | *SIR3* | Protein involved in subtelomeric silencing; mutants display increased colonization of the mouse kidney relative to the wild-type strain | ORF, Verified | *SIR3* | CAGL0M00770g | Schwarzmüller et al., 2014 |
| **8D9** | CAGL0K11396g | No Deletion Verified | *SIR4* | Protein involved in subtelomeric silencing and regulation of biofilm formation | ORF, Verified | *SIR4* | CAGL0K11396g | Schwarzmüller et al., 2014 |
| **8D10** | CAGL0L09042g | No Deletion Verified | | Ortholog(s) have chromatin binding, histone H4K12 acetyltransferase activity, histone H4K5 acetyltransferase activity, histone H4K8 acetyltransferase activity, histone acetyltransferase activity | ORF, Uncharacterized | *HAT1* | CAGL0L09042g | Schwarzmüller et al., 2014 |
| **8D11** | CAGL0E00693g | No Deletion Verified | *ADA3* | Putative transcription coactivator, component of the Spt-Ada-Gcn5 acetyltransferase (SAGA) complex; involved in drug resistance and virulence; null mutant grows slowly but shows increased invasive growth and increased virulence | ORF, Verified | *NGG1* | CAGL0E00693g | Schwarzmüller et al., 2014 |
| **8D12** | CAGL0K02981g | No Deletion Verified | | Ortholog(s) have histone H2A acetyltransferase activity, histone H4 acetyltransferase activity and role in heterochromatin formation | ORF, Uncharacterized | *NAT4* | CAGL0K02981g | Schwarzmüller et al., 2014 |
| **8E1** | CAGL0E00561g | No Deletion Verified | *TUP11* | General repressor of transcription; paralog of TUP1; acts as a stronger repressor than Tup1p and more closely related to ScTUP1 | ORF, Verified | *TUP1* | CAGL0E00561g | Schwarzmüller et al., 2014 |
| **8E2** | CAGL0J07348g | No Deletion Verified | | Ortholog(s) have structural constituent of cytoskeleton activity, role in mitotic spindle organization, spindle pole body organization and outer plaque of mitotic spindle pole body, outer plaque of spindle pole body localization | ORF, Uncharacterized | *CNM67* | CAGL0J07348g | Schwarzmüller et al., 2014 |
| **8E3** | CAGL0D06490g | No Deletion Verified | | Ortholog(s) have chitin deacetylase activity and role in ascospore wall assembly | ORF, Uncharacterized | *CDA2* | CAGL0D06490g | Schwarzmüller et al., 2014 |
| **8E4** | CAGL0F02651g | No Deletion Verified | | Ortholog(s) have catalytic activity, role in ascospore wall assembly and cytosol localization | ORF, Uncharacterized | *DIT1* | CAGL0F02651g | Schwarzmüller et al., 2014 |
| **8E5** | CAGL0F02607g | Verified | | Ortholog(s) have NADPH dehydrogenase activity, role in ascospore wall assembly and endoplasmic reticulum localization | ORF, Uncharacterized | *DIT2* | CAGL0F02607g | Schwarzmüller et al., 2014 |
| **8E6** | CAGL0L09933g | Verified | | Ortholog(s) have K48-linked polyubiquitin modification-dependent protein binding, K63-linked polyubiquitin modification-dependent protein binding, protein-macromolecule adaptor activity, ubiquitin binding activity | ORF, Uncharacterized | *CUE5* | CAGL0L09933g | Schwarzmüller et al., 2014 |
| **8E8** | CAGL0K08118g | No Deletion Verified | | Ortholog(s) have phosphatidylinositol binding activity | ORF, Uncharacterized | *LEC1* | CAGL0K08118g | Schwarzmüller et al., 2014 |
| **8E9** | CAGL0E03454g | Verified | *IMH1* | Protein with a role in vesicular transport; protein abundance increased in ace2 mutant cells | ORF, Uncharacterized | *IMH1* | CAGL0E03454g | Schwarzmüller et al., 2014 |
| **8E10** | CAGL0K08998g | Verified | | Ortholog(s) have role in ascospore formation, ascospore wall assembly and prospore membrane localization | ORF, Uncharacterized | *OSW1* | CAGL0K08998g | Schwarzmüller et al., 2014 |
| **8E11** | CAGL0F01859g | No Deletion Verified | | Ortholog(s) have role in ascospore wall assembly and cytoplasm, prospore membrane localization | ORF, Uncharacterized | *OSW2* | CAGL0F01859g | Schwarzmüller et al., 2014 |
| **8E12** | CAGL0M13387g | No Deletion Verified | | Ortholog(s) have role in ascospore wall assembly | ORF, Uncharacterized | *PFS1* | CAGL0M13387g | Schwarzmüller et al., 2014 |
| **8F1** | CAGL0M13519g | No Deletion Verified | | Ortholog(s) have ATP-dependent activity, acting on RNA, RNA binding, RNA helicase activity | ORF, Uncharacterized | *HAS1* | CAGL0M13519g | Schwarzmüller et al., 2014 |
| **8F2** | CAGL0L11286g | No Deletion Verified | | Ortholog(s) have role in ascospore-type prospore membrane formation, spore membrane bending pathway and cytoplasm, prospore membrane localization | ORF, Uncharacterized | *SMA2* | CAGL0L11286g | Schwarzmüller et al., 2014 |
| **8F3** | CAGL0A01672g | No Deletion Verified | | Protein of unknown function | ORF, Verified | *SEC9* | CAGL0A01672g | Schwarzmüller et al., 2014 |
| **8F4** | CAGL0M05313g | No Deletion Verified | | Protein of unknown function | ORF, Uncharacterized | *YBR197C* | CAGL0M05313g | Schwarzmüller et al., 2014 |
| **8F5** | CAGL0M07183g | Verified | | Ortholog(s) have role in intracellular monoatomic ion homeostasis, mitochondrion inheritance, mitochondrion organization, regulation of cardiolipin metabolic process and mitochondrial inner membrane, mitochondrion localization | ORF, Uncharacterized | *MDM31* | CAGL0M07183g | Schwarzmüller et al., 2014 |
| **8F6** | CAGL0D05720g | No Deletion Verified | | Ortholog(s) have role in ascospore formation, ascospore wall assembly | ORF, Uncharacterized | *SPO75* | CAGL0D05720g | Schwarzmüller et al., 2014 |
| **8F7** | CAGL0H01639g | No Deletion Verified | | Ortholog(s) have protein serine/threonine kinase activity | ORF, Uncharacterized | *SPS1* | CAGL0H01639g | Schwarzmüller et al., 2014 |
| **8F8** | CAGL0J06050g | No Deletion Verified | | Ortholog(s) have role in cell wall assembly and extracellular region localization | ORF, Verified | *YGP1* | CAGL0J06050g | Schwarzmüller et al., 2014 |
| **8F9** | CAGL0J03740g | No Deletion Verified | | Ortholog(s) have protein kinase activator activity, role in ascospore formation, ascospore wall assembly, positive regulation of protein autophosphorylation and ascospore wall localization | ORF, Uncharacterized | *SSP2* | CAGL0J03740g | Schwarzmüller et al., 2014 |
| **8F10** | CAGL0B01991g | Verified | | Ortholog(s) have palmitoyltransferase activity and role in cortical actin cytoskeleton organization, establishment of cell polarity, protein palmitoylation, regulation of exocytosis, vacuole fusion, non-autophagic | ORF, Uncharacterized | *SWF1* | CAGL0B01991g | Schwarzmüller et al., 2014 |
| **8F11** | CAGL0M03465g | Verified | | Ortholog(s) have ammonium transmembrane transporter activity, role in ammonium transmembrane transport, nitrogen utilization and plasma membrane localization | ORF, Uncharacterized | *ATO2* | CAGL0M03465g | Schwarzmüller et al., 2014 |
| **8F12** | CAGL0G08316g | No Deletion Verified | *ARV1* | Lipid transporter involved in sterol trafficking and transport of glycosylphosphatidylinositol and sphingolipid precursors | ORF, Uncharacterized | *ARV1* | CAGL0G08316g | Schwarzmüller et al., 2014 |
| **8G1** | CAGL0J05874g | Verified | | Ortholog(s) have role in cellular response to amino acid stimulus, proteasome-mediated ubiquitin-dependent protein catabolic process, ubiquitin-dependent protein catabolic process and Asi complex, nuclear inner membrane localization | ORF, Uncharacterized | *ASI2* | CAGL0J05874g | Schwarzmüller et al., 2014 |
| **8G2** | CAGL0J03542g | No Deletion Verified | | Ortholog(s) have phospholipase activity | ORF, Uncharacterized | *ATG15* | CAGL0J03542g | Schwarzmüller et al., 2014 |
| **8G3** | CAGL0I03652g | No Deletion Verified | | Ortholog(s) have phospholipid scramblase activity | ORF, Uncharacterized | *ATG9* | CAGL0I03652g | Schwarzmüller et al., 2014 |
| **8G4** | CAGL0A03212g | No Deletion Verified | | Ortholog(s) have ammonium transmembrane transporter activity, role in ammonium transmembrane transport, nitrogen utilization, transmembrane transport and plasma membrane localization | ORF, Uncharacterized | *ATO3* | CAGL0A03212g | Schwarzmüller et al., 2014 |
| **8G5** | CAGL0D04686g | No Deletion Verified | | Ortholog(s) have metalloendopeptidase activity and role in axial cellular bud site selection, cytogamy, peptide mating pheromone maturation involved in positive regulation of conjugation with cellular fusion | ORF, Uncharacterized | *AXL1* | CAGL0D04686g | Schwarzmüller et al., 2014 |
| **8G6** | CAGL0L07546g | No Deletion Verified | | Ortholog(s) have amino acid transmembrane transporter activity and role in amino acid transport, transmembrane transport | ORF, Uncharacterized | *BAP3* | CAGL0L07546g | Schwarzmüller et al., 2014 |

| | | | | | | | | |
|---|---|---|---|---|---|---|---|---|
| 8G7 | CAGL0H03399g | No Deletion Verified | | Ortholog(s) have (R)-carnitine transmembrane transporter activity, choline transmembrane transporter activity, ethanolamine transmembrane transporter activity | ORF, Uncharacterized | HNM1 | CAGL0H03399g | Schwarzmüller et al., 2014 |
| 8G8 | CAGL0C05445g | No Deletion Verified | | Ortholog(s) have role in endocytosis | ORF, Uncharacterized | BRE4 | CAGL0C05445g | Schwarzmüller et al., 2014 |
| 8G9 | CAGL0E01507g | No Deletion Verified | | Has domain(s) with predicted transmembrane transporter activity and role in transmembrane transport | ORF, Uncharacterized | BSC6 | CAGL0E01507g | Schwarzmüller et al., 2014 |
| 8G10 | CAGL0K12408g | No Deletion Verified | | Ortholog(s) have phosphatidylinositol deacylase activity | ORF, Uncharacterized | BST1 | CAGL0K12408g | Schwarzmüller et al., 2014 |
| 8G11 | CAGL0K12474g | No Deletion Verified | | Ortholog(s) have role in regulation of transcription by RNA polymerase II and CCR4-NOT complex localization | ORF, Uncharacterized | CAF16 | CAGL0K12474g | Schwarzmüller et al., 2014 |
| 8G12 | CAGL0K10956g | No Deletion Verified | | Protein of unknown function, Ortholog(s) found to be mitochondrial cytochrome c oxidase (complex IV) assembly factor; also involved in translational regulation of Cox1p and prevention of Cox1p aggregation before assembly; associates with complex IV assembly intermediates and complex III/complex IV supercomplexes; located in the mitochondrial membrane | ORF, Uncharacterized | COX14 | CAGL0K10956g | Schwarzmüller et al., 2014 |
| 8H1 | CAGL0I01562g | No Deletion Verified | | Has domain(s) with predicted proton transmembrane transporter activity, proton-transporting ATPase activity, rotational mechanism activity and role in proton transmembrane transport | ORF, Uncharacterized | VMA3 | CAGL0I01562g | Schwarzmüller et al., 2014 |
| 8H2 | CAGL0L02827g | Verified | | Ortholog(s) have protein tyrosine phosphatase activity | ORF, Uncharacterized | PTP2 | CAGL0L02827g | Schwarzmüller et al., 2014 |
| 8H3 | CAGL0F02827g | No Deletion Verified | | Ortholog(s) have protein folding chaperone, protein-containing complex binding activity | ORF, Uncharacterized | DFM1 | CAGL0F02827g | Schwarzmüller et al., 2014 |
| 8H4 | CAGL0K08426g | Verified | | Protein of unknown function, Ortholog(s) are component of the p24 complex; have a role in misfolded protein quality control; binds to GPI anchor proteins and mediates their efficient transport from the ER to the Golgi; integral membrane protein that associates with endoplasmic reticulum-derived COPII-coated vesicles | ORF, Uncharacterized | EMP24 | CAGL0K08426g | Schwarzmüller et al., 2014 |
| 8H5 | CAGL0L13244g | Verified | | Ortholog(s) have carbohydrate binding activity, role in endoplasmic reticulum to Golgi vesicle-mediated transport and COPII-coated ER to Golgi transport vesicle, Golgi membrane localization | ORF, Uncharacterized | EMP46 | CAGL0L13244g | Schwarzmüller et al., 2014 |
| 8H6 | CAGL0B01683g | Verified | TMN2 | Ortholog(s) have role in intracellular copper ion homeostasis, invasive growth in response to glucose limitation, pseudohyphal growth, vacuolar transport and fungal-type vacuole membrane localization | ORF, Uncharacterized | TMN2 | CAGL0B01683g | Schwarzmüller et al., 2014 |
| 8H7 | CAGL0K12034g | No Deletion Verified | ENA1 | Na(+)-ATPase with broad substrate specificity; plays a role in sodium detoxification; highly upregulated by increased osmotic pressure or sodium concentration | ORF, Verified | ENA1 | CAGL0K12034g | Schwarzmüller et al., 2014 |
| 8H8 | CAGL0J01870g | No Deletion Verified | | Putative Ca2+ ATPase | ORF, Uncharacterized | PMR1 | CAGL0J01870g | Schwarzmüller et al., 2014 |
| 8H9 | CAGL0K01793g | No Deletion Verified | | Ortholog(s) have similarity to Emp24p and Erv25p; member of the p24 family involved in ER to Golgi transport | ORF, Uncharacterized | ERP3 | CAGL0K01793g | Schwarzmüller et al., 2014 |
| 8H10 | CAGL0C02761g | Verified | | Ortholog(s) have role in endoplasmic reticulum to Golgi vesicle-mediated transport, protein retention in ER lumen and COPII-coated ER to Golgi transport vesicle localization | ORF, Uncharacterized | ERP2 | CAGL0C02761g | Schwarzmüller et al., 2014 |
| 8H11 | CAGL0I01232g | Verified | | Ortholog(s) have similarity to Emp24p and Erv25p; member of the p24 family involved in ER to Golgi transport | ORF, Uncharacterized | ERP5 | CAGL0I01232g | Schwarzmüller et al., 2014 |
| 8H12 | CAGL0G00682g | No Deletion Verified | | Ortholog(s) have role in endoplasmic reticulum to Golgi vesicle-mediated transport, protein retention in ER lumen and COPII-coated ER to Golgi transport vesicle localization | ORF, Uncharacterized | ERP1 | CAGL0G00682g | Schwarzmüller et al., 2014 |
| 9A1 | CAGL0M06985g | No Deletion Verified | | Ortholog(s) have L-cystine transmembrane transporter activity, role in L-cystine transport and endosome, fungal-type vacuole, plasma membrane localization | ORF, Uncharacterized | ERS1 | CAGL0M06985g | Schwarzmüller et al., 2014 |
| 9A2 | CAGL0I00440g | No Deletion Verified | | Ortholog(s) have cargo receptor activity and role in ascospore formation, axial cellular bud site selection, endoplasmic reticulum to Golgi vesicle-mediated transport | ORF, Uncharacterized | ERV14 | CAGL0I00440g | Schwarzmüller et al., 2014 |
| 9A3 | CAGL0M05929g | Verified | | Ortholog(s) have role in protein import into mitochondrial matrix and PAM complex, Tim23 associated import motor localization | ORF, Uncharacterized | PAM17 | CAGL0M05929g | Schwarzmüller et al., 2014 |
| 9A4 | CAGL0M12276g | No Deletion Verified | GEM1 | GTPase subunit of the mitochondrial ERMES complex; involved in regulation of mitochondrion organization; null mutant shows increased resistance to fluconazole, abnormal mitochondrial morphology, and decreased growth on glycerol | ORF, Verified | GEM1 | CAGL0M12276g | Schwarzmüller et al., 2014 |
| 9A5 | CAGL0E03872g | No Deletion Verified | | Ortholog(s) have SNAP receptor activity, role in Golgi vesicle transport, vesicle fusion and Golgi medial cisterna, SNARE complex localization | ORF, Uncharacterized | GOS1 | CAGL0E03872g | Schwarzmüller et al., 2014 |
| 9A6 | CAGL0E02519g | No Deletion Verified | | Ortholog(s) have role in intracellular zinc ion homeostasis, negative regulation of transcription by RNA polymerase II, response to toxic substance and plasma membrane localization | ORF, Uncharacterized | IZH2 | CAGL0E02519g | Schwarzmüller et al., 2014 |
| 9A7 | CAGL0I07491g | Verified | | Ortholog(s) have role in intracellular zinc ion homeostasis | ORF, Uncharacterized | IZH4 | CAGL0I07491g | Schwarzmüller et al., 2014 |
| 9A8 | CAGL0D02442g | Verified | LEM3 | Glycoprotein involved in the membrane translocation of phospholipids and alkylphosphocholine drugs | ORF, Verified | LEM3 | CAGL0D02442g | Schwarzmüller et al., 2014 |
| 9A9 | CAGL0I03872g | Verified | | Ortholog(s) have glucose binding activity, role in glucose mediated signaling pathway and plasma membrane localization | ORF, Uncharacterized | RGT2 | CAGL0I03872g | Schwarzmüller et al., 2014 |
| 9A10 | CAGL0I00286g | No Deletion Verified | | Has domain(s) with predicted transmembrane transporter activity, role in transmembrane transport and membrane localization | ORF, Uncharacterized | HXT2 | CAGL0I00286g | Schwarzmüller et al., 2014 |
| 9A11 | CAGL0G06864g | Verified | | Protein of unknown function; Ortholog(s) are Nuclear envelope protein; required for SPB insertion, SPB duplication, Kar5p localization near the SPB and nuclear fusion; interacts with Mps2p to tether half-bridge to core SPB; N-terminal acetylation by Eco1p regulates its role in nuclear organization; localizes to the SPB half bridge and telomeres during meiosis; required with Ndj1p and Csm4p for meiotic bouquet formation and telomere-led rapid prophase movement; member of the SUN protein family (Sad1-UNC-84 homology) | ORF, Uncharacterized | MPS3 | CAGL0G06864g | Schwarzmüller et al., 2014 |
| 9A12 | CAGL0I09240g | No Deletion Verified | | Ortholog(s) have N-terminal protein N-methyltransferase activity, role in N-terminal peptidyl-proline dimethylation, cytoplasmic translation and cytosol localization | ORF, Uncharacterized | TAE1 | CAGL0I09240g | Schwarzmüller et al., 2014 |
| 9B1 | CAGL0I10494g | Verified | | Ortholog(s) have role in eisosome assembly, plasma membrane organization, protein localization to plasma membrane and eisosome, growing cell tip, plasma membrane localization | ORF, Uncharacterized | FHN1 | CAGL0I10494g | Schwarzmüller et al., 2014 |
| 9B2 | CAGL0L08888g | No Deletion Verified | | Ortholog(s) have sterol binding activity, role in sphingolipid metabolic process, sterol transport and fungal-type vacuole membrane localization | ORF, Uncharacterized | NCR1 | CAGL0L08888g | Schwarzmüller et al., 2014 |
| 9B3 | CAGL0K02959g | No Deletion Verified | | Ortholog(s) have phosphoprotein phosphatase activity | ORF, Uncharacterized | NEM1 | CAGL0K02959g | Schwarzmüller et al., 2014 |
| 9B4 | CAGL0M08602g | No Deletion Verified | | Ortholog(s) have ATPase-coupled monoatomic cation transmembrane transporter activity, copper ion binding activity and role in copper ion export, intracellular iron ion homeostasis, transmembrane transport | ORF, Uncharacterized | CCC2 | CAGL0M08602g | Schwarzmüller et al., 2014 |
| 9B5 | CAGL0J08371g | Verified | | Ortholog(s) have role in plasma membrane fusion involved in cytogamy, regulation of plasma membrane sterol distribution and mating projection tip localization | ORF, Uncharacterized | PRM1 | CAGL0J08371g | Schwarzmüller et al., 2014 |
| 9B6 | CAGL0G04433g | Verified | | Has domain(s) with predicted transmembrane transporter activity and role in transmembrane transport | ORF, Uncharacterized | PRM10 | CAGL0G04433g | Schwarzmüller et al., 2014 |
| 9B7 | CAGL0M02167g | No Deletion Verified | | Ortholog(s) are pheromone-regulated protein proposed to be involved in mating; predicted to have 1 transmembrane segment; transcriptionally regulated by Ste12p during mating and by Cat8p during the diauxic shift | ORF, Uncharacterized | PRM4 | CAGL0M02167g | Schwarzmüller et al., 2014 |
| 9B8 | CAGL0J10076g | No Deletion Verified | | Protein of unknown function | ORF, Uncharacterized | YNL058C | CAGL0J10076g | Schwarzmüller et al., 2014 |
| 9B9 | CAGL0L01463g | Verified | | Ortholog(s) have potassium ion transmembrane transporter activity, role in potassium ion transmembrane transport and cellular bud tip, mating projection tip, plasma membrane localization | ORF, Uncharacterized | PRM6 | CAGL0L01463g | Schwarzmüller et al., 2014 |
| 9B10 | CAGL0K07876g | No Deletion Verified | | Ortholog(s) have role in intracellular distribution of mitochondria, mitochondrial fission, mitochondrion inheritance, mitochondrion organization and cytoplasm, mitochondrion localization | ORF, Uncharacterized | MDM36 | CAGL0K07876g | Schwarzmüller et al., 2014 |
| 9B11 | CAGL0D00242g | Verified | | Ortholog(s) have unfolded protein binding activity, role in protein folding, ubiquitin-dependent ERAD pathway and endoplasmic reticulum membrane localization | ORF, Uncharacterized | CNE1 | CAGL0D00242g | Schwarzmüller et al., 2014 |
| 9B12 | CAGL0D00352g | No Deletion Verified | PXA2 | Ortholog(s) have ATPase-coupled transmembrane transporter activity, role in ATP transport, fatty acid transmembrane transport and peroxisomal membrane, peroxisome localization | ORF, Uncharacterized | PXA2 | CAGL0D00352g | Schwarzmüller et al., 2014 |
| 9C1 | CAGL0L10142g | No Deletion Verified | RSB1 | Putative sphingolipid flippase; gene is upregulated in azole-resistant strain | ORF, Uncharacterized | RSB1 | CAGL0L10142g | Schwarzmüller et al., 2014 |
| 9C2 | CAGL0K00715g | Verified | RTA1 | Putative protein involved in 7-aminocholesterol resistance; gene is upregulated in azole-resistant strain | ORF, Uncharacterized | RTA1 | CAGL0K00715g | Schwarzmüller et al., 2014 |
| 9C3 | CAGL0G03663g | No Deletion Verified | | Ortholog(s) have FFAT motif binding, phosphatidylinositol binding, protein-membrane adaptor activity | ORF, Uncharacterized | SCS2 | CAGL0G03663g | Schwarzmüller et al., 2014 |
| 9C4 | CAGL0C03179g | No Deletion Verified | | Ortholog(s) have SNAP receptor activity and role in endoplasmic reticulum to Golgi vesicle-mediated transport, retrograde vesicle-mediated transport, Golgi to endoplasmic reticulum, vesicle fusion, vesicle fusion with Golgi apparatus | ORF, Uncharacterized | SEC22 | CAGL0C03179g | Schwarzmüller et al., 2014 |
| 9C5 | CAGL0C02717g | Verified | | Ortholog(s) have phosphoprotein phosphatase activity and role in negative regulation of phospholipid biosynthetic process, nuclear envelope organization, regulation of phospholipid biosynthetic process, reticulophagy | ORF, Uncharacterized | SPO7 | CAGL0C02717g | Schwarzmüller et al., 2014 |
| 9C7 | CAGL0H06457g | No Deletion Verified | | Ortholog(s) have metalloendopeptidase activity, role in mitochondrial protein processing, peptide mating pheromone maturation involved in positive regulation of conjugation with cellular fusion and mitochondrial matrix localization | ORF, Uncharacterized | STE23 | CAGL0H06457g | Schwarzmüller et al., 2014 |
| 9C8 | CAGL0L01551g | Verified | | Ortholog(s) have role in ascospore formation, endocytosis, protein localization to eisosome filament and cell cortex, eisosome, plasma membrane localization | ORF, Uncharacterized | SUR7 | CAGL0L01551g | Schwarzmüller et al., 2014 |
| 9C9 | CAGL0E06204g | No Deletion Verified | | Has domain(s) with predicted proton transmembrane transporter activity, proton-transporting ATPase activity, rotational mechanism activity and role in proton transmembrane transport | ORF, Uncharacterized | VMA11 | CAGL0E06204g | Schwarzmüller et al., 2014 |
| 9C10 | CAGL0L01947g | No Deletion Verified | | Ortholog(s) have ubiquitin protein ligase activity, ubiquitin-protein transferase activity | ORF, Uncharacterized | TUL1 | CAGL0L01947g | Schwarzmüller et al., 2014 |
| 9C11 | CAGL0G01342g | No Deletion Verified | | Ortholog(s) have enzyme activator activity, role in positive regulation of phosphatidylinositol biosynthetic process, protein localization to vacuolar membrane and PAS complex, fungal-type vacuole membrane, membrane localization | ORF, Uncharacterized | VAC7 | CAGL0G01342g | Schwarzmüller et al., 2014 |
| 9C12 | CAGL0M13255g | No Deletion Verified | | Ortholog(s) have endoplasmic reticulum localization | ORF, Uncharacterized | YET1 | CAGL0M13255g | Schwarzmüller et al., 2014 |
| 9D1 | CAGL0C03047g | No Deletion Verified | | Ortholog(s) have histone chaperone activity, role in constitutive heterochromatin formation, nucleosome organization and FACT complex, nucleus localization | ORF, Uncharacterized | SPT16 | CAGL0C03047g | Schwarzmüller et al., 2014 |
| 9D2 | CAGL0H07513g | No Deletion Verified | IFA38 | Predicted ketoreductase involved in production of very long chain fatty acid for sphingolipid biosynthesis; mutants show reduced sensitivity to caspofungin and increased sensitivity to micafungin | ORF, Verified | IFA38 | CAGL0H07513g | Schwarzmüller et al., 2014 |
| 9D3 | CAGL0G05093g | Verified | | Has domain(s) with predicted ATP binding, ATPase activity; Ortholog(s) have similarity to ABC transporter family members; lacks predicted membrane-spanning regions; transcriptionally activated by Yrm1p along with genes involved in multidrug resistance | ORF, Uncharacterized | YDR061W | CAGL0G05093g | Schwarzmüller et al., 2014 |
| 9D4 | CAGL0G08019g | Verified | | Ortholog(s) have plasma membrane localization | ORF, Uncharacterized | ILT1 | CAGL0G08019g | Schwarzmüller et al., 2014 |
| 9D9 | CAGL0F03641g | Verified | | Protein of unknown function | ORF, Uncharacterized | YML018C | CAGL0F03641g | Schwarzmüller et al., 2014 |
| 9D10 | CAGL0K12584g | No Deletion Verified | | Ortholog(s) have AP-1 adaptor complex binding activity, role in clathrin-coated vesicle cargo loading and membrane localization | ORF, Uncharacterized | MIL1 | CAGL0K12584g | Schwarzmüller et al., 2014 |
| 9D11 | CAGL0C03267g | No Deletion Verified | FPS1 | Glycerol transporter; involved in flucytosine resistance; double fps1/fps2 mutant accumulates glycerol, has constitutive cell wall stress, is hypersensitive to caspofungin in vitro and in vivo | ORF, Verified | FPS1 | CAGL0C03267g | Schwarzmüller et al., 2014 |
| 9D12 | CAGL0E05940g | Verified | | Ortholog(s) have FAD transmembrane transporter activity, calcium channel activity | ORF, Uncharacterized | FLC1 | CAGL0E05940g | Schwarzmüller et al., 2014 |
| 9E1 | CAGL0E02981g | No Deletion Verified | | Ortholog(s) have glycero-3-phosphocholine acyltransferase activity and role in phosphatidylcholine acyl-chain remodeling, phosphatidylcholine biosynthesis from sn-glycero-3-phosphocholine, phosphatidylcholine biosynthetic process | ORF, Uncharacterized | GPC1 | CAGL0E02981g | Schwarzmüller et al., 2014 |
| 9E2 | CAGL0F08481g | No Deletion Verified | | Protein of unknown function; Ortholog(s) predicted to contain a single transmembrane domain; localized to both the mitochondrial outer membrane and the plasma membrane | ORF, Uncharacterized | YGR266W | CAGL0F08481g | Schwarzmüller et al., 2014 |
| 9E3 | CAGL0J00363g | No Deletion Verified | | Putative protein of major facilitator superfamily; gene is downregulated in azole-resistant strain | ORF, Uncharacterized | YHK8 | CAGL0J00363g | Schwarzmüller et al., 2014 |
| 9E4 | CAGL0G05962g | No Deletion Verified | | Ortholog(s) have endoplasmic reticulum localization | ORF, Uncharacterized | YHR140W | CAGL0G05962g | Schwarzmüller et al., 2014 |
| 9E5 | CAGL0G04081g | No Deletion Verified | | Ortholog(s) have endoplasmic reticulum localization | ORF, Uncharacterized | THI73 | CAGL0G04081g | Schwarzmüller et al., 2014 |
| 9E6 | CAGL0M09933g | No Deletion Verified | | Ortholog(s) have role in lipid droplet formation, lipid droplet organization, negative regulation of sphingolipid biosynthetic process, positive regulation of lipid biosynthetic process, protein localization | ORF, Uncharacterized | SEI1 | CAGL0M09933g | Schwarzmüller et al., 2014 |
| 9E8 | CAGL0I00726g | Verified | | Ortholog(s) have protein-containing complex binding activity, role in ascospore wall assembly, lipid droplet organization and lipid droplet localization | ORF, Uncharacterized | LDO16 | CAGL0I00726g | Schwarzmüller et al., 2014 |
| 9E9 | CAGL0K04279g | Verified | SCM4 | Mitochondrial outer membrane protein of unknown function; Ortholog(s) predicted to have 4 transmembrane segments; import is mediated by Tom70p and Mim1p | ORF, Uncharacterized | SCM4 | CAGL0K04279g | Schwarzmüller et al., 2014 |
| 9E10 | CAGL0H02519g | Verified | | Ortholog(s) are putative membrane protein, involved in salt tolerance; cytoplasmic localization in a punctate pattern | ORF, Uncharacterized | YMR253C | CAGL0H02519g | Schwarzmüller et al., 2014 |
| 9E11 | CAGL0I04664g | Verified | | Ortholog(s) have role in cell morphogenesis involved in conjugation with cellular fusion, cytogamy and fungal-type cell wall, mating projection tip localization | ORF, Uncharacterized | FIG1 | CAGL0I04664g | Schwarzmüller et al., 2014 |
| 9E12 | CAGL0C00539g | No Deletion Verified | | Ortholog(s) have role in positive regulation of (R)-carnitine transmembrane transport, positive regulation of polyamine transmembrane transport and endoplasmic reticulum membrane, fungal-type vacuole membrane, plasma membrane localization | ORF, Uncharacterized | AGP2 | CAGL0C00539g | Schwarzmüller et al., 2014 |
| 9F1 | CAGL0F05929g | No Deletion Verified | | Ortholog(s) have role in negative regulation of cytosolic calcium ion concentration and cellular bud neck, plasma membrane localization | ORF, Uncharacterized | RCH1 | CAGL0F05929g | Schwarzmüller et al., 2014 |

| | | | | | | | | |
|---|---|---|---|---|---|---|---|---|
| **9F2** | CAGL0L08294g | Verified | | Ortholog(s) have role in axial cellular bud site selection and cell division site, cellular bud neck, cellular bud neck septin ring, incipient cellular bud site, plasma membrane localization | ORF, Verified | *AXL2* | CAGL0L08294g | Schwarzmüller et al., 2014 |
| **9F3** | CAGL0E00429g | No Deletion Verified | | Ortholog(s) have aminophospholipid flippase activity and role in actin cortical patch localization, endocytosis, intracellular protein transport, phospholipid translocation, retrograde transport, endosome to Golgi | ORF, Uncharactarized | *CDC50* | CAGL0E00429g | Schwarzmüller et al., 2014 |
| **9F4** | CAGL0I02508g | Verified | | Ortholog(s) have copper ion transmembrane transporter activity and role in copper ion import, intracellular copper ion homeostasis, protein maturation by copper ion transfer | ORF, Uncharactarized | *CTR2* | CAGL0I02508g | Schwarzmüller et al., 2014 |
| **9F5** | CAGL0K02827g | Verified | *DID4* | Ortholog(s) have role in late endosome to vacuole transport, protein transport to vacuole involved in ubiquitin-dependent protein catabolic process via the multivesicular body sorting pathway | ORF, Uncharactarized | *DID4* | CAGL0K02827g | Schwarzmüller et al., 2014 |
| **9F6** | CAGL0G06270g | No Deletion Verified | | Ortholog(s) have phosphatidylcholine flippase activity, phosphatidylethanolamine flippase activity, phosphatidylserine flippase activity | ORF, Uncharactarized | *DRS2* | CAGL0G06270g | Schwarzmüller et al., 2014 |
| **9F7** | CAGL0K12914g | Verified | | Ortholog(s) have carbohydrate derivative binding activity, role in endoplasmic reticulum to Golgi vesicle-mediated transport and COPII-coated ER to Golgi transport vesicle, Golgi membrane, endoplasmic reticulum membrane localization | ORF, Uncharactarized | *EMP47* | CAGL0K12914g | Schwarzmüller et al., 2014 |
| **9F8** | CAGL0G08217g | No Deletion Verified | | Ortholog(s) have palmitoyltransferase activity, protein-cysteine S-palmitoyltransferase activity and role in protein palmitoylation, protein targeting to membrane | ORF, Uncharactarized | *ERF2* | CAGL0G08217g | Schwarzmüller et al., 2014 |
| **9F9** | CAGL0L11308g | Verified | | Ortholog(s) have role in endoplasmic reticulum to Golgi vesicle-mediated transport and COPII-coated ER to Golgi transport vesicle, Golgi membrane, endoplasmic reticulum membrane localization | ORF, Uncharactarized | *ERV41* | CAGL0L11308g | Schwarzmüller et al., 2014 |
| **9F10** | CAGL0M11946g | No Deletion Verified | | Ortholog(s) have role in endoplasmic reticulum to Golgi vesicle-mediated transport and COPII-coated ER to Golgi transport vesicle, Golgi membrane, endoplasmic reticulum membrane localization | ORF, Uncharactarized | *ERV46* | CAGL0M11946g | Schwarzmüller et al., 2014 |
| **9F11** | CAGL0H06017g | No Deletion Verified | *FLR1* | Multidrug transporter of the major facilitator superfamily involved in 5-flucytosine resistance; gene is downregulated in azole-resistant strain | ORF, Verified | *FLR1* | CAGL0H06017g | Schwarzmüller et al., 2014 |
| **9F12** | CAGL0L03267g | No Deletion Verified | *GAP1* | Ortholog(s) have L-proline transmembrane transporter activity, amino acid transmembrane transporter activity, beta-alanine transmembrane transporter activity, polyamine transmembrane transporter activity | ORF, Uncharactarized | *GAP1* | CAGL0L03267g | Schwarzmüller et al., 2014 |
| **9G1** | CAGL0L01485g | Verified | | Putative protein of the ER membrane involved in hexose transporter secretion; gene is upregulated in azole-resistant strain | ORF, Uncharactarized | *GSF2* | CAGL0L01485g | Schwarzmüller et al., 2014 |
| **9G2** | CAGL0H07271g | No Deletion Verified | | Ortholog(s) have inositol pentakisphosphate 2-kinase activity, role in inositol phosphate biosynthetic process, nuclear-transcribed mRNA catabolic process, non-stop decay and nucleus localization | ORF, Uncharactarized | *IPK1* | CAGL0H07271g | Schwarzmüller et al., 2014 |
| **9G3** | CAGL0C03938g | No Deletion Verified | | Has domain(s) with predicted glycosyltransferase activity and role in protein glycosylation | ORF, Uncharactarized | *MNT3* | CAGL0C03938g | Schwarzmüller et al., 2014 |
| **9G4** | CAGL0K04367g | Verified | | Ortholog(s) have L-methionine secondary active transmembrane transporter activity, role in methionine import across plasma membrane, sulfur amino acid transport and plasma membrane localization | ORF, Uncharactarized | *MUP1* | CAGL0K04367g | Schwarzmüller et al., 2014 |
| **9G5** | CAGL0G00352g | No Deletion Verified | | Ortholog(s) have role in coenzyme A biosynthetic process and CoA-synthesizing protein complex localization | ORF, Uncharactarized | *CAB4* | CAGL0G00352g | Schwarzmüller et al., 2014 |
| **9G6** | CAGL0K08272g | Verified | | Ortholog(s) have sphingosine-1-phosphate phosphatase activity, role in phospholipid dephosphorylation, sphingolipid biosynthetic process and endoplasmic reticulum localization | ORF, Uncharactarized | *YSR3* | CAGL0K08272g | Schwarzmüller et al., 2014 |
| **9G7** | CAGL0I01012g | Verified | *PEP12* | Ortholog(s) have SNAP receptor activity and role in Golgi to vacuole transport, cytoplasm to vacuole transport by the Cvt pathway, macroautophagy, vacuole inheritance | ORF, Uncharactarized | *PEP12* | CAGL0I01012g | Schwarzmüller et al., 2014 |
| **9G8** | CAGL0B02475g | Verified | *PHO84* | Ortholog(s) have inorganic phosphate transmembrane transporter activity, manganese ion transmembrane transporter activity, selenite:proton symporter activity | ORF, Uncharactarized | *PHO84* | CAGL0B02475g | Schwarzmüller et al., 2014 |
| **9G9** | CAGL0L04378g | No Deletion Verified | | Has domain(s) with predicted transmembrane transporter activity and role in transmembrane transport | ORF, Uncharactarized | *PNS1* | CAGL0L04378g | Schwarzmüller et al., 2014 |
| **9G10** | CAGL0G08624g | Verified | *QDR2* | Drug:H+ antiporter of the Major Facilitator Superfamily, confers imidazole drug resistance, involved in quinidine/multidrug efflux; gene is activated by Pdr1p; upregulated in azole-resistant strain | ORF, Verified | *QDR1* | CAGL0G08624g | Schwarzmüller et al., 2014 |
| **9G11** | CAGL0F03443g | No Deletion Verified | | Ortholog(s) have metalloendopeptidase activity, role in CAAX-box protein processing, peptide mating pheromone maturation involved in positive regulation of conjugation with cellular fusion and endoplasmic reticulum membrane localization | ORF, Uncharactarized | *RCE1* | CAGL0F03443g | Schwarzmüller et al., 2014 |
| **9G12** | CAGL0K00297g | Verified | | Ortholog(s) have phosphatidylinositol-3,5-bisphosphate 3-phosphatase activity, phosphatidylinositol-3-phosphate phosphatase activity, phosphatidylinositol-4-phosphate phosphatase activity | ORF, Uncharactarized | *SAC1* | CAGL0K00297g | Schwarzmüller et al., 2014 |
| **9H1** | CAGL0M06127g | No Deletion Verified | | Ortholog(s) have protein transmembrane transporter activity and role in filamentous growth, post-translational protein targeting to membrane, translocation | ORF, Uncharactarized | *SEC66* | CAGL0M06127g | Schwarzmüller et al., 2014 |
| **9H2** | CAGL0K00649g | Verified | | Ortholog(s) have role in endoplasmic reticulum to Golgi vesicle-mediated transport and endoplasmic reticulum, membrane localization | ORF, Uncharactarized | *SOP4* | CAGL0K00649g | Schwarzmüller et al., 2014 |
| **9H3** | CAGL0E06160g | No Deletion Verified | | Ortholog(s) have SNAP receptor activity, phosphatidic acid binding activity and role in ascospore formation, ascospore-type prospore assembly, ascospore-type prospore membrane formation | ORF, Uncharactarized | *SSO2* | CAGL0E06160g | Schwarzmüller et al., 2014 |
| **9H4** | CAGL0L02651g | Verified | | Ortholog(s) have aminopeptidase activity, role in peptide pheromone maturation and trans-Golgi network localization | ORF, Uncharactarized | *STE13* | CAGL0L02651g | Schwarzmüller et al., 2014 |
| **9H5** | CAGL0M13321g | Verified | | Ortholog(s) have COPII receptor activity and role in COPII-coated vesicle cargo loading, endoplasmic reticulum to Golgi vesicle-mediated transport, fungal-type cell wall organization, protein retention in Golgi apparatus | ORF, Uncharactarized | *SVP26* | CAGL0M13321g | Schwarzmüller et al., 2014 |
| **9H6** | CAGL0D03146g | Verified | | Ortholog(s) have role in Golgi to endosome transport, Golgi to plasma membrane protein transport, protein localization to Golgi apparatus and Golgi membrane, trans-Golgi network localization | ORF, Uncharactarized | *SYS1* | CAGL0D03146g | Schwarzmüller et al., 2014 |
| **9H7** | CAGL0L06292g | No Deletion Verified | | Ortholog(s) have role in vesicle-mediated transport and Golgi membrane localization | ORF, Uncharactarized | *TVP15* | CAGL0L06292g | Schwarzmüller et al., 2014 |
| **9H8** | CAGL0K03025g | No Deletion Verified | | Ortholog(s) have role in vesicle-mediated transport and Golgi membrane localization | ORF, Uncharactarized | *TVP18* | CAGL0K03025g | Schwarzmüller et al., 2014 |
| **9H9** | CAGL0K09988g | Verified | | Ortholog(s) have role in vesicle-mediated transport and Golgi membrane localization | ORF, Uncharactarized | *TVP23* | CAGL0K09988g | Schwarzmüller et al., 2014 |
| **9H10** | CAGL0E02079g | Verified | | Ortholog(s) have sequence-specific DNA binding activity, role in positive regulation of transcription by RNA polymerase II and nucleus localization | ORF, Uncharactarized | *MSN1* | CAGL0E02079g | Schwarzmüller et al., 2014 |
| **9H11** | CAGL0J07876g | No Deletion Verified | | Protein of unknown function | ORF, Uncharactarized | *RTC4* | CAGL0J07876g | Schwarzmüller et al., 2014 |
| **9H12** | CAGL0M02585g | Verified | *SPP1* | Predicted transcriptional activator, subunit of Set1/COMPASS complex with histone methyltransferase activity (H3-K4 specific) | ORF, Uncharactarized | *SPP1* | CAGL0M02585g | Schwarzmüller et al., 2014 |
| **10A1** | CAGL0K00517g | No Deletion Verified | | Ortholog(s) have protein serine/threonine kinase activity | ORF, Uncharactarized | *TOR2* | CAGL0K00517g | Schwarzmüller et al., 2014 |
| **10A2** | CAGL0I07887g | No Deletion Verified | | Ortholog(s) have protein-macromolecule adaptor activity, role in protein-containing complex localization, ubiquitin-dependent ERAD pathway, vesicle organization and Cvt complex, cytoplasm, phagophore assembly site localization | ORF, Uncharactarized | *ATG19* | CAGL0I07887g | Schwarzmüller et al., 2014 |
| **10A3** | CAGL0A01265g | No Deletion Verified | *EPA10* | Putative adhesin-like protein; belongs to adhesin cluster I | ORF, Uncharactarized | | CAGL0A01265g | Schwarzmüller et al., 2014 |
| **10A4** | CAGL0D01892g | Verified | | Ortholog(s) have L-aspartate:2-oxoglutarate aminotransferase activity, ribosomal large subunit binding activity and role in aspartate biosynthetic process, negative regulation of translation in response to oxidative stress | ORF, Uncharactarized | *AAT2* | CAGL0D01892g | Schwarzmüller et al., 2014 |
| **10A5** | CAGL0E04884g | No Deletion Verified | | Ortholog(s) have DNA-binding transcription factor activity, RNA polymerase II-specific and RNA polymerase II cis-regulatory region sequence-specific DNA binding, RNA polymerase II-specific DNA-binding transcription factor binding, TFIIB-class transcription factor binding, TFIID-class transcription factor complex binding, transcription coactivator activity | ORF, Uncharactarized | *ADR1* | CAGL0E04884g | Schwarzmüller et al., 2014 |
| **10A6** | CAGL0D04774g | No Deletion Verified | | Ortholog(s) have adenine nucleotide transmembrane transporter activity, role in ATP transport, fatty acid beta-oxidation, peroxisome organization and peroxisomal membrane localization | ORF, Uncharactarized | *ANT1* | CAGL0D04774g | Schwarzmüller et al., 2014 |
| **10A7** | CAGL0K05577g | Verified | | Ortholog(s) have manganese ion transmembrane transporter activity, role in intracellular manganese ion homeostasis and Golgi membrane, late endosome, trans-Golgi network localization | ORF, Uncharactarized | *ATX2* | CAGL0K05577g | Schwarzmüller et al., 2014 |
| **10A8** | CAGL0J11836g | No Deletion Verified | | Ortholog(s) have carnitine O-acetyltransferase activity, role in carnitine metabolic process and mitochondrion, peroxisome localization | ORF, Uncharactarized | *CAT2* | CAGL0J11836g | Schwarzmüller et al., 2014 |
| **10A9** | CAGL0F05071g | No Deletion Verified | | Ortholog(s) have delta(3)-delta(2)-enoyl-CoA isomerase activity, role in fatty acid beta-oxidation and peroxisome localization | ORF, Uncharactarized | *ECI1* | CAGL0F05071g | Schwarzmüller et al., 2014 |
| **10A10** | CAGL0J01936g | No Deletion Verified | | Ortholog(s) have role in protein import into peroxisome matrix, protein insertion into mitochondrial outer membrane, protein targeting to mitochondrion and cytosol localization | ORF, Uncharactarized | *DJP1* | CAGL0J01936g | Schwarzmüller et al., 2014 |
| **10A11** | CAGL0I04642g | No Deletion Verified | | Ortholog(s) have long-chain fatty acid transporter activity, very long-chain fatty acid-CoA ligase activity | ORF, Uncharactarized | *FAT1* | CAGL0I04642g | Schwarzmüller et al., 2014 |
| **10A12** | CAGL0B04917g | No Deletion Verified | | S-adenosylmethionine synthetase | ORF, Verified | *IDP2* | CAGL0B04917g | Schwarzmüller et al., 2014 |
| **10B1** | CAGL0F06875g | No Deletion Verified | | Ortholog(s) have mRNA binding, saccharopine dehydrogenase (NAD+, L-lysine-forming) activity and role in lysine biosynthetic process, lysine biosynthetic process via aminoadipic acid, protein import into peroxisome matrix | ORF, Uncharactarized | *LYS1* | CAGL0F06875g | Schwarzmüller et al., 2014 |
| **10B2** | CAGL0K10978g | No Deletion Verified | | Ortholog(s) have role in lysine biosynthetic process, lysine biosynthetic process via aminoadipic acid and mitochondrion localization | ORF, Uncharactarized | *LYS4* | CAGL0K10978g | Schwarzmüller et al., 2014 |
| **10B3** | CAGL0M05687g | No Deletion Verified | | Ortholog(s) have NAD+ diphosphatase activity, role in NAD-cap decapping, NADH metabolic process, RNA decapping and peroxisome localization | ORF, Uncharactarized | *NPY1* | CAGL0M05687g | Schwarzmüller et al., 2014 |
| **10B4** | CAGL0K10846g | Verified | | Ortholog(s) have 8-oxo-7,8-dihydroguanosine triphosphate pyrophosphatase activity, coenzyme A diphosphatase activity, role in DNA repair and peroxisome localization | ORF, Uncharactarized | *PCD1* | CAGL0K10846g | Schwarzmüller et al., 2014 |
| **10B5** | CAGL0K06853g | No Deletion Verified | | Ortholog(s) have mRNA binding, oxalate-CoA ligase activity, role in oxalate catabolic process and peroxisomal matrix, peroxisomal membrane localization | ORF, Uncharactarized | *PCS60* | CAGL0K06853g | Schwarzmüller et al., 2014 |
| **10B6** | CAGL0H09174g | No Deletion Verified | *PEX1* | Ortholog(s) have ATPase activity, role in protein import into peroxisome matrix, receptor recycling, protein unfolding and peroxisomal membrane localization | ORF, Uncharactarized | *PEX1* | CAGL0H09174g | Schwarzmüller et al., 2014 |
| **10B7** | CAGL0M08690g | No Deletion Verified | *PEX10* | Ortholog(s) have ubiquitin protein ligase activity | ORF, Uncharactarized | *PEX10* | CAGL0M08690g | Schwarzmüller et al., 2014 |
| **10B8** | CAGL0D02618g | No Deletion Verified | *PEX11* | Ortholog(s) have role in fatty acid oxidation, peroxisome fission and endoplasmic reticulum, peroxisomal importomer complex, peroxisomal membrane localization | ORF, Uncharactarized | *PEX11* | CAGL0D02618g | Schwarzmüller et al., 2014 |
| **10B9** | CAGL0M07469g | No Deletion Verified | *PEX12* | Ortholog(s) have ubiquitin ligase activator activity, ubiquitin protein ligase activity and role in proteasome-mediated ubiquitin-dependent protein catabolic process, protein import into peroxisome matrix, protein polyubiquitination | ORF, Uncharactarized | *PEX12* | CAGL0M07469g | Schwarzmüller et al., 2014 |
| **10B10** | CAGL0A04147g | No Deletion Verified | *PEX13* | Ortholog(s) have protein-macromolecule adaptor activity, role in protein import into peroxisome matrix, docking and peroxisomal importomer complex, peroxisomal membrane localization | ORF, Uncharactarized | *PEX13* | CAGL0A04147g | Schwarzmüller et al., 2014 |
| **10B11** | CAGL0E05082g | No Deletion Verified | *PEX14* | Ortholog(s) have protein-macromolecule adaptor activity, role in protein import into peroxisome matrix, docking and peroxisomal importomer complex, peroxisomal membrane, peroxisome localization | ORF, Uncharactarized | *PEX14* | CAGL0E05082g | Schwarzmüller et al., 2014 |
| **10B12** | CAGL0F00935g | No Deletion Verified | *PEX15* | Ortholog(s) have protein-membrane adaptor activity, role in protein import into peroxisome matrix, receptor recycling and peroxisomal membrane localization | ORF, Uncharactarized | *PEX15* | CAGL0F00935g | Schwarzmüller et al., 2014 |
| **10C1** | CAGL0K04851g | No Deletion Verified | *PEX17* | Ortholog(s) have role in protein import into peroxisome matrix, docking and peroxisomal importomer complex, peroxisomal membrane localization | ORF, Uncharactarized | *PEX17* | CAGL0K04851g | Schwarzmüller et al., 2014 |
| **10C2** | CAGL0F08019g | Verified | *PEX21* | Ortholog(s) have role in positive regulation of binding, protein import into peroxisome matrix and cytosol, peroxisome localization | ORF, Uncharactarized | *PEX21* | CAGL0F08019g | Schwarzmüller et al., 2014 |
| **10C3** | CAGL0D00792g | No Deletion Verified | *PEX19* | Ortholog(s) have peroxisome membrane targeting sequence binding activity | ORF, Uncharactarized | *PEX19* | CAGL0D00792g | Schwarzmüller et al., 2014 |
| **10C4** | CAGL0I06292g | No Deletion Verified | *PEX2* | Has domain(s) with predicted zinc ion binding activity, role in protein import into peroxisome matrix and peroxisomal membrane localization | ORF, Uncharactarized | *PEX2* | CAGL0I06292g | Schwarzmüller et al., 2014 |
| **10C5** | CAGL0J07194g | No Deletion Verified | *PEX22* | Protein of unknown function, Ortholog(s) are putative peroxisomal membrane protein; required for import of peroxisomal proteins | ORF, Uncharactarized | *PEX22* | CAGL0J07194g | Schwarzmüller et al., 2014 |
| **10C7** | CAGL0I10879g | No Deletion Verified | *PEX25* | Ortholog(s) have role in peroxisome fission, peroxisome organization and peroxisomal membrane localization | ORF, Uncharactarized | *PEX27* | CAGL0I10879g | Schwarzmüller et al., 2014 |
| **10C8** | CAGL0I01870g | No Deletion Verified | *PEX24* | Ortholog(s) have role in peroxisome organization and peroxisomal membrane localization | ORF, Uncharactarized | *PEX28* | CAGL0I01870g | Schwarzmüller et al., 2014 |
| **10C9** | CAGL0M08866g | No Deletion Verified | *PEX29* | Ortholog(s) have role in ER-dependent peroxisome organization, peroxisome organization and endoplasmic reticulum, peroxisomal membrane, peroxisome localization | ORF, Uncharactarized | *PEX29* | CAGL0M08866g | Schwarzmüller et al., 2014 |
| **10C10** | CAGL0M01342g | No Deletion Verified | *PEX3* | Peroxisomal assembly protein required for peroxisome biogenesis | ORF, Uncharactarized | *PEX3* | CAGL0M01342g | Schwarzmüller et al., 2014 |
| **10C11** | CAGL0F08657g | No Deletion Verified | *PEX23* | Ortholog(s) have role in peroxisome organization, regulation of peroxisome organization and peroxisomal membrane localization | ORF, Uncharactarized | *PEX31* | CAGL0F08657g | Schwarzmüller et al., 2014 |
| **10C12** | CAGL0M06061g | No Deletion Verified | *PEX32* | Ortholog(s) have role in peroxisome organization and peroxisomal membrane localization | ORF, Uncharactarized | *PEX32* | CAGL0M06061g | Schwarzmüller et al., 2014 |
| **10D1** | CAGL0I10450g | No Deletion Verified | *PEX4* | Ortholog(s) have ubiquitin-protein transferase activity, role in peroxisome organization, protein autoubiquitination, protein import into peroxisome matrix, receptor recycling, protein monoubiquitination and peroxisome localization | ORF, Uncharactarized | *PEX4* | CAGL0I10450g | Schwarzmüller et al., 2014 |
| **10D2** | CAGL0K11209g | Verified | *PEX5* | Ortholog(s) have peroxisome matrix targeting signal-1 binding, protein carrier chaperone, protein-macromolecule adaptor activity and role in protein import into peroxisome matrix, protein import into peroxisome matrix, docking | ORF, Uncharactarized | *PEX5* | CAGL0K11209g | Schwarzmüller et al., 2014 |

| | | | | | | | | |
|---|---|---|---|---|---|---|---|---|
| **10D3** | CAGL0D02574g | No Deletion Verified | *PEX6* | Has domain(s) with predicted ATP binding, ATPase activity | ORF, Uncharactarized | *PEX6* | CAGL0D02574g | Schwarzmüller et al., 2014 |
| **10D4** | CAGL0B01529g | No Deletion Verified | *PEX7* | Ortholog(s) have peroxisome matrix targeting signal-2 binding activity, role in protein import into peroxisome matrix, docking and cytosol, peroxisome localization | ORF, Uncharactarized | *PEX7* | CAGL0B01529g | Schwarzmüller et al., 2014 |
| **10D9** | CAGL0K01221g | No Deletion Verified | *PEX8* | Ortholog(s) have protein-macromolecule adaptor activity, role in protein import into peroxisome matrix and peroxisomal importomer complex localization | ORF, Uncharactarized | *PEX8* | CAGL0K01221g | Schwarzmüller et al., 2014 |
| **10D10** | CAGL0A01716g | No Deletion Verified | | Ortholog(s) have nicotinamidase activity and role in negative regulation of DNA amplification, nicotinate nucleotide salvage, rDNA heterochromatin formation, subtelomeric heterochromatin formation | ORF, Uncharactarized | *PNC1* | CAGL0A01716g | Schwarzmüller et al., 2014 |
| **10D11** | CAGL0H06787g | No Deletion Verified | | Ortholog(s) have acetyl-CoA C-acyltransferase activity, mRNA binding activity, role in fatty acid beta-oxidation and mitochondrial intermembrane space, peroxisomal matrix, peroxisome localization | ORF, Uncharactarized | *POT1* | CAGL0H06787g | Schwarzmüller et al., 2014 |
| **10D12** | CAGL0F07315g | No Deletion Verified | | Ortholog(s) have ADP binding, AMP binding, AMP-activated protein kinase activity, ATP binding, protein serine/threonine kinase activator activity | ORF, Uncharactarized | *SNF4* | CAGL0F07315g | Schwarzmüller et al., 2014 |
| **10E1** | CAGL0H08063g | No Deletion Verified | | Ortholog(s) have 2,4-dienoyl-CoA reductase (NADPH) activity, role in ascospore formation, fatty acid catabolic process and peroxisomal matrix localization | ORF, Uncharactarized | *SPS19* | CAGL0H08063g | Schwarzmüller et al., 2014 |
| **10E2** | CAGL0L06094g | No Deletion Verified | *STR3* | Putative cystathionine beta-lyase; gene is upregulated in azole-resistant strain | ORF, Uncharactarized | *STR3* | CAGL0L06094g | Schwarzmüller et al., 2014 |
| **10E3** | CAGL0B03465g | No Deletion Verified | | Ortholog(s) have role in ethanol metabolic process and mitochondrial inner membrane localization | ORF, Uncharactarized | *SYM1* | CAGL0B03465g | Schwarzmüller et al., 2014 |
| **10E4** | CAGL0B04059g | Verified | | Ortholog(s) have acyl-CoA hydrolase activity, role in fatty acid beta-oxidation, fatty acid oxidation and peroxisome localization | ORF, Uncharactarized | *TES1* | CAGL0B04059g | Schwarzmüller et al., 2014 |
| **10E5** | CAGL0L02299g | Verified | *VPS1* | Ortholog(s) have GTPase activity, actin filament binding activity | ORF, Uncharactarized | *VPS1* | CAGL0L02299g | Schwarzmüller et al., 2014 |
| **10E6** | CAGL0L10736g | No Deletion Verified | | Putative peripheral membrane protein of peroxisomes; gene is upregulated in azole-resistant strain | ORF, Uncharactarized | *INP1* | CAGL0L10736g | Schwarzmüller et al., 2014 |
| **10E7** | CAGL0J08547g | No Deletion Verified | | Protein of unknown function; Ortholog(s) have peroxisomal matrix-localized lipase activity; required for normal peroxisome morphology; contains a peroxisomal targeting signal type 1 (PTS1) and a lipase motif; peroxisomal import requires the PTS1 receptor, Pex5p and self-interaction; transcriptionally activated by Yrm1p along with genes involved in multidrug resistance; oleic acid inducible | ORF, Uncharactarized | *LPX1* | CAGL0J08547g | Schwarzmüller et al., 2014 |
| **10E9** | CAGL0J08184g | No Deletion Verified | | Ortholog(s) have basic amino acid transmembrane transporter activity and role in basic amino acid transport | ORF, Uncharactarized | *ALP1* | CAGL0J08184g | Schwarzmüller et al., 2014 |
| **10E10** | CAGL0B04433g | No Deletion Verified | | Has domain(s) with predicted transmembrane transporter activity, role in transmembrane transport and membrane localization | ORF, Uncharactarized | *FUR4* | CAGL0B04433g | Schwarzmüller et al., 2014 |
| **10E11** | CAGL0L07788g | No Deletion Verified | | Ortholog(s) have cytoskeletal protein binding, lipid binding activity | ORF, Uncharactarized | *RVS161* | CAGL0L07788g | Schwarzmüller et al., 2014 |
| **10E12** | CAGL0J07480g | No Deletion Verified | | Ortholog(s) have role in chitin biosynthetic process, division septum assembly and cellular bud neck, cellular bud neck septin collar, incipient cellular bud site, septin ring localization | ORF, Uncharactarized | *BNI4* | CAGL0J07480g | Schwarzmüller et al., 2014 |
| **10F1** | CAGL0C03069g | No Deletion Verified | | Ortholog(s) have CDP-diacylglycerol-serine O-phosphatidyltransferase activity | ORF, Uncharactarized | *CHO1* | CAGL0C03069g | Schwarzmüller et al., 2014 |
| **10F2** | CAGL0E03201g | No Deletion Verified | | Ortholog(s) have phosphatidylethanolamine N-methyltransferase activity, role in phosphatidylcholine biosynthetic process and cortical endoplasmic reticulum, perinuclear endoplasmic reticulum localization | ORF, Uncharactarized | *CHO2* | CAGL0E03201g | Schwarzmüller et al., 2014 |
| **10F3** | CAGL0M04367g | No Deletion Verified | | Ortholog(s) have choline kinase activity, ethanolamine kinase activity and role in phosphatidylcholine biosynthetic process, phosphatidylethanolamine biosynthetic process | ORF, Uncharactarized | *CKI1* | CAGL0M04367g | Schwarzmüller et al., 2014 |
| **10F4** | CAGL0K09570g | Verified | | Ortholog(s) have diacylglycerol cholinephosphotransferase activity, role in CDP-choline pathway, mitotic nuclear membrane biogenesis, phosphatidic acid biosynthetic process and Golgi apparatus, mitochondrial outer membrane localization | ORF, Uncharactarized | *CPT1* | CAGL0K09570g | Schwarzmüller et al., 2014 |
| **10F5** | CAGL0I03784g | No Deletion Verified | | Ortholog(s) have cardiolipin synthase (CMP-forming) activity | ORF, Uncharactarized | *CRD1* | CAGL0I03784g | Schwarzmüller et al., 2014 |
| **10F6** | CAGL0L13068g | Verified | | Ortholog(s) have diacylglycerol cholinephosphotransferase activity, ethanolaminephosphotransferase activity and role in phosphatidylcholine biosynthetic process, phosphatidylethanolamine biosynthetic process | ORF, Uncharactarized | *EPT1* | CAGL0L13068g | Schwarzmüller et al., 2014 |
| **10F7** | CAGL0I10747g | No Deletion Verified | | Ortholog(s) have ammonium transmembrane transporter activity, role in ammonium transmembrane transport, nitrogen utilization and plasma membrane localization | ORF, Uncharactarized | *MEP1* | CAGL0I10747g | Schwarzmüller et al., 2014 |
| **10F8** | CAGL0L08184g | No Deletion Verified | *FEN1* | Predicted fatty acid elongase with role in sphingolipid biosynthetic process; mutants show reduced sensitivity to caspofungin and increased sensitivity to micafungin | ORF, Verified | *ELO2* | CAGL0L08184g | Schwarzmüller et al., 2014 |
| **10F9** | CAGL0K08162g | No Deletion Verified | | Ortholog(s) have glycerol-3-phosphate O-acyltransferase activity, glycerone-phosphate O-acyltransferase activity and role in phospholipid biosynthetic process, regulation of triglyceride metabolic process | ORF, Uncharactarized | *GPT2* | CAGL0K08162g | Schwarzmüller et al., 2014 |
| **10F10** | CAGL0L03432g | No Deletion Verified | *GWT1* | Inositol acyltransferase with role in early steps of GPI anchor biosynthetic process; antifungal drug target | ORF, Uncharactarized | *GWT1* | CAGL0L03432g | Schwarzmüller et al., 2014 |
| **10F11** | CAGL0B01947g | No Deletion Verified | *INO2* | Transcriptional regulator involved in de novo inositol biosynthesis; activator of INO1 gene expression; mutants unable to grow in the absence of inositol | ORF, Verified | *INO2* | CAGL0B01947g | Schwarzmüller et al., 2014 |
| **10F12** | CAGL0I07359g | No Deletion Verified | *INO4* | Transcriptional regulator involved in de novo inositol biosynthesis; activator of INO1 gene expression; mutants unable to grow in the absence of inositol | ORF, Verified | *INO4* | CAGL0I07359g | Schwarzmüller et al., 2014 |
| **10G1** | CAGL0K02739g | No Deletion Verified | | Ortholog(s) have sphingosine N-acyltransferase activity, role in ceramide biosynthetic process and acyl-CoA ceramide synthase complex localization | ORF, Uncharactarized | *LAG1* | CAGL0K02739g | Schwarzmüller et al., 2014 |
| **10G2** | CAGL0I04334g | No Deletion Verified | | Ortholog(s) have sphingosine-1-phosphate phosphatase activity, role in calcium-mediated signaling and endoplasmic reticulum localization | ORF, Uncharactarized | *LCB3* | CAGL0I04334g | Schwarzmüller et al., 2014 |
| **10G3** | CAGL0F08723g | Verified | | Ortholog(s) have ethanolamine-phosphate cytidylyltransferase activity and role in phosphatidylethanolamine biosynthetic process | ORF, Uncharactarized | *ECT1* | CAGL0F08723g | Schwarzmüller et al., 2014 |
| **10G4** | CAGL0K03267g | No Deletion Verified | *OPI1* | Putative transcriptional regulator with a role in de novo inositol biosynthesis; essential gene | ORF, Verified | *OPI1* | CAGL0K03267g | Schwarzmüller et al., 2014 |
| **10G5** | CAGL0F00363g | No Deletion Verified | *OPI3* | Ortholog(s) have phosphatidyl-N-dimethylethanolamine N-methyltransferase activity, phosphatidyl-N-methylethanolamine N-methyltransferase activity and role in phosphatidylcholine biosynthetic process | ORF, Uncharactarized | *OPI3* | CAGL0F00363g | Schwarzmüller et al., 2014 |
| **10G6** | CAGL0K01749g | No Deletion Verified | | Ortholog(s) have lipid binding, sterol transfer activity and role in ER to Golgi ceramide transport, endocytosis, exocytosis, maintenance of cell polarity, piecemeal microautophagy of the nucleus, reticulophagy, sterol transport | ORF, Uncharactarized | *OSH2* | CAGL0K01749g | Schwarzmüller et al., 2014 |
| **10G7** | CAGL0J10780g | No Deletion Verified | | Ortholog(s) have lipid binding, sterol transfer activity | ORF, Uncharactarized | *OSH3* | CAGL0J10780g | Schwarzmüller et al., 2014 |
| **10G8** | CAGL0K02849g | No Deletion Verified | | Has domain(s) with predicted lipid binding activity | ORF, Uncharactarized | *OSH7* | CAGL0K02849g | Schwarzmüller et al., 2014 |
| **10G9** | CAGL0B04015g | No Deletion Verified | | Has domain(s) with predicted catalytic activity, choline-phosphate cytidylyltransferase activity and role in CDP-choline pathway, biosynthetic process | ORF, Uncharactarized | *PCT1* | CAGL0B04015g | Schwarzmüller et al., 2014 |
| **10G10** | CAGL0J07436g | No Deletion Verified | *PDR16* | Putative ABC transporter | ORF, Uncharactarized | *PDR16* | CAGL0J07436g | Schwarzmüller et al., 2014 |
| **10G11** | CAGL0J08074g | Verified | | Ortholog(s) have phosphatidylinositol transfer activity | ORF, Uncharactarized | *PDR17* | CAGL0J08074g | Schwarzmüller et al., 2014 |
| **10G12** | CAGL0J06226g | No Deletion Verified | | Ortholog(s) have phosphatidylserine decarboxylase activity | ORF, Verified | *PSD1* | CAGL0J06226g | Schwarzmüller et al., 2014 |
| **10H1** | CAGL0I08745g | No Deletion Verified | | Ortholog(s) have phosphatidylserine decarboxylase activity, role in phosphatidylcholine biosynthetic process and endosome localization | ORF, Uncharactarized | *PSD2* | CAGL0I08745g | Schwarzmüller et al., 2014 |
| **10H2** | CAGL0E02849g | No Deletion Verified | | Ortholog(s) have glycerol-3-phosphate O-acyltransferase activity, glycerone-phosphate O-acyltransferase activity, role in phospholipid biosynthetic process and endoplasmic reticulum localization | ORF, Uncharactarized | *SCT1* | CAGL0E02849g | Schwarzmüller et al., 2014 |
| **10H3** | CAGL0M13101g | No Deletion Verified | | Protein of unknown function; Ortholog(s) regulate phospholipid asymmetry at plasma membrane; may act to generate normal levels of PI4P; may act together with or upstream of Stt4p; at least partially mediates proper localization of Stt4p to the plasma membrane | ORF, Uncharactarized | *SFK1* | CAGL0M13101g | Schwarzmüller et al., 2014 |
| **10H4** | CAGL0I04070g | No Deletion Verified | | Ortholog(s) have 1-acylglycerol-3-phosphate O-acyltransferase activity, role in glycerophospholipid biosynthetic process and lipid droplet localization | ORF, Uncharactarized | *SLC1* | CAGL0I04070g | Schwarzmüller et al., 2014 |
| **10H5** | CAGL0K00737g | No Deletion Verified | | Ortholog(s) have N-acetyltransferase activity, role in response to xenobiotic stimulus and nuclear envelope, plasma membrane localization | ORF, Uncharactarized | *SLI1* | CAGL0K00737g | Schwarzmüller et al., 2014 |
| **10H6** | CAGL0H01375g | No Deletion Verified | *SUR2* | Predicted sphinganine hydroxylase with role in sphingolipid biosynthesis; mutants show reduced sensitivity to caspofungin and increased sensitivity to micafungin | ORF, Verified | *SUR2* | CAGL0H01375g | Schwarzmüller et al., 2014 |
| **10H7** | CAGL0G04851g | Verified | *SUR4* | Predicted fatty acid elongase involved in production of very long chain fatty acids for sphingolipid biosynthesis; mutants show reduced sensitivity to caspofungin and increased sensitivity to micafungin | ORF, Verified | *ELO3* | CAGL0G04851g | Schwarzmüller et al., 2014 |
| **10H8** | CAGL0C05049g | No Deletion Verified | | Ortholog(s) have lipid binding, sterol transfer activity and role in endocytosis, exocytosis, maintenance of cell polarity, piecemeal microautophagy of the nucleus, sterol transport | ORF, Uncharactarized | *SWH1* | CAGL0C05049g | Schwarzmüller et al., 2014 |
| **10H9** | CAGL0D04972g | No Deletion Verified | | Ortholog(s) have 1-acylglycerophosphocholine O-acyltransferase activity | ORF, Uncharactarized | *TAZ1* | CAGL0D04972g | Schwarzmüller et al., 2014 |
| **10H10** | CAGL0F04433g | No Deletion Verified | *URA7* | CTP synthase | ORF, Uncharactarized | *URA7* | CAGL0F04433g | Schwarzmüller et al., 2014 |
| **10H11** | CAGL0I04620g | No Deletion Verified | | Ortholog(s) have acyltransferase activity and role in phosphatidylinositol acyl-chain remodeling | ORF, Uncharactarized | *CST26* | CAGL0I04620g | Schwarzmüller et al., 2014 |
| **10H12** | CAGL0A03806g | No Deletion Verified | | Ortholog(s) have role in ascospore wall assembly | ORF, Uncharactarized | *MUM3* | CAGL0A03806g | Purohit & Gajjar, 2022 |
| | CAGL0M01254g | Verified | *STE12-1* | Ortholog(s) have DNA-binding transcription factor activity | ORF, Verified | STE12 | CAGL0M01254g | Purohit & Gajjar, 2022 |
| | CAGL0H02145g | Verified | *STE12-2* | Putative transcription factor, required for filamentous growth induced by nitrogen starvation and for virulence; functionally complements S. cerevisiae ste12 mutant | ORF, Uncharactarized | STE12 | CAGL0H02145g | Purohit & Gajjar, 2022 |
| | CAGL0M01254g, CAGL0H02145g | Verified | *ste12* ΔΔ | Double deletion strain | | STE12 | | Purohit & Gajjar, 2022 |
| | CAGL0M01716g | Verified | *TEC1-1* | Transcription factor involved in control of biofilm formation | ORF, Verified | TEC1 | CAGL0M01716g | Purohit & Gajjar, 2022 |
| | CAGL0F04081g | Verified | *TEC1-2* | Has domain(s) with predicted DNA-binding transcription factor activity and role in regulation of DNA-templated transcription | ORF, Uncharactarized | TEC1 | CAGL0F04081g | Purohit & Gajjar, 2022 |
| | CAGL0M01716g, CAGL0F04081g | Verified | *tec1ΔΔ* | Double deletion strain | | TEC1 | | Purohit & Gajjar, 2022 |

| | | |
|---|---|---|
| ***C. glabrata* Deletion Library Source** | Schwarzmüller T, Ma B, Hiller E, Istel F, Tscherner M, Brunke S, Ames L, Firon A, Green B, Cabral V, Marcet-Houben M, Jacobsen ID, Quintin J, Seider K, Frohner I, Glaser W, Jungwirth H, Bachellier-Bassi S, Chauvel M, Zeidler U, Ferrandon D, Gabaldón T, Hube B, d'Enfert C, Rupp S, Cormack B, Haynes K, Kuchler K. Systematic phenotyping of a large-scale *Candida glabrata* deletion collection reveals novel antifungal tolerance genes. PLoS Pathog. 2014 Jun 19;10(6):e1004211. doi:10.1371/journal.ppat.1004211. PMID: 24945925; PMCID: PMC4063973. | Article Link |
| ***C. glabrata TEC1* and *STE12* Deletion Strains Source** | Purohit, D., Gajjar, D. Tec1 and Ste12 transcription factors play a role in adaptation to low pH stress and biofilm formation in the human opportunistic fungal pathogen *Candida glabrata*. *Int Microbiol* **25**, 789–802 (2022). https://doi.org/10.1007/s10123-022-00264-7 | Article Link |
