## Supplementary Table S3 for "A novel *Candida glabrata* protein regulated by mating signalling pathway shapes inter-species interaction"

**Supplementary Table 3.**

Output of sequence database search (GenBank, UniProt, RefSeq and PDBSTR) of MOTIF Search (MOTIF2) using A-D-V-W-H as query pattern.

| Sequence ID | Description | Position |
|---|---|---|
| rs:WP_132416323 | [WP_132416323] epoxide hydrolase family protein, partial [Kribbella albertanoniae]. | 159..165 |
| rs:WP_131517574 | [WP_131517574] epoxide hydrolase family protein [Kribbella capetownensis]. | 159..165 |
| rs:WP_134686050 | [WP_134686050] PP2C family protein-serine/threonine phosphatase [Brevibacillus migulae]. | 343..349 |
| rs:WP_157685074 | [WP_157685074] AMP-binding protein [Robbsia andropogonis]. | 731..737 |
| rs:WP_157685764 | [WP_157685764] AMP-binding protein [Robbsia andropogonis]. | 731..737 |
| rs:WP_144394495 | [WP_144394495] hypothetical protein [Pleionea sediminis]. | 95..101 |
| rs:WP_208722326 | [WP_208722326] BTAD domain-containing putative transcriptional regulator [Amycolatopsis thermoflava]. | 707..713 |
| rs:WP_237335480 | [WP_237335480] BTAD domain-containing putative transcriptional regulator [Amycolatopsis tucumanensis]. | 717..723 |
| rs:WP_063630395 | [WP_063630395] BTAD domain-containing putative transcriptional regulator [Amycolatopsis thermoflava]. | 707..713 |
| rs:WP_131364561 | [WP_131364561] epoxide hydrolase [Kribbella pittospori]. | 159..165 |
| rs:WP_219049454 | [WP_219049454] hypothetical protein [Streptomyces sp. JJ66]. | 67..73 |
| rs:WP_281792998 | [WP_281792998] GNAT family N-acetyltransferase [Desulforhabdus amnigena]. | 103..109 |
| **rs:XP_445695** | **[XP_445695] uncharacterized protein CAGL0D06666g [Nakaseomyces glabratus].** | **17..23** |
| tr:A0A9W4UWG5_9ARCH | [A0A9W4UWG5] SubName: Full=Uncharacterized protein {ECO:0000313\|EMBL:CAI9831111.1}; | 233..239 |
| tr:A0A928FZ48_9BACT | [A0A928FZ48] SubName: Full=GH3 auxin-responsive promoter family protein {ECO:0000313\|EMBL:MBE6198840.1}; | 223..229 |
| tr:A0A931RDG8_9BACT | [A0A931RDG8] RecName: Full=Formamidopyrimidine-DNA glycosylase {ECO:0000256\|HAMAP-Rule:MF_00103}; Short=Fapy-I | 23..29 |
| tr:A0A0F5K0V1_9BURK | [A0A0F5K0V1] RecName: Full=Carrier domain-containing protein {ECO:0000259\|PROSITE:PS50075}; | 689..695 |
| tr:A0A3N2GPQ5_9PSEU | [A0A3N2GPQ5] SubName: Full=DNA-binding SARP family transcriptional activator {ECO:0000313\|EMBL:ROS38604.1}; | 661..667 |
| tr:A0A346AVH5_9VIRU | [A0A346AVH5] SubName: Full=Major capsid protein {ECO:0000313\|EMBL:AXL15451.1}; | 513..519 |

**Supplementary Table 3.**

Output of sequence database search (GenBank, UniProt, RefSeq and PDBSTR) of MOTIF Search (MOTIF2) using A-V-V-P-H as query pattern.

| Sequence ID | Description | Position |
|---|---|---|
| rs:WP_380873707 | [WP_380873707] LysE family translocator [Streptomyces albogriseolus]. | 44..50 |
| rs:WP_376493330 | [WP_376493330] FtsW/RodA/SpoVE family cell cycle protein [Streptomyces sp. NPDC056132]. | 56..62 |
| rs:WP_379430877 | [WP_379430877] DUF5134 domain-containing protein [Nocardia sp. NPDC059228]. | 183..189 |
| rs:WP_394312767 | [WP_394312767] LysE family translocator [Streptomyces lusitanus]. | 44..50 |
| rs:WP_390924474 | [WP_390924474] LysE family transporter [Streptomyces albogriseolus]. | 44..50 |
| rs:WP_044663637 | [WP_044663637] GGDEF domain-containing protein [Sphingobium yanoikuyae]. | 75..81 |
| rs:WP_379384117 | [WP_379384117] MULTISPECIES: DUF5134 domain-containing protein [unclassified Nocardia]. | 184..190 |
| rs:WP_388786292 | [WP_388786292] FtsW/RodA/SpoVE family cell cycle protein [Streptomyces sp. NPDC006691]. | 56..62 |
| rs:WP_266548107 | [WP_266548107] LysE family translocator [Streptomyces albogriseolus]. | 44..50 |
| rs:WP_093839416 | [WP_093839416] Cas10/Cmr2 second palm domain-containing protein [Streptomyces aidingensis]. | 317..323 |
| rs:WP_386006080 | [WP_386006080] hypothetical protein [Streptomyces violascens]. | 56..62 |
| rs:WP_304551080 | [WP_304551080] MULTISPECIES: hypothetical protein [Bradyrhizobium]. | 341..347 |
| rs:WP_385100567 | [WP_385100567] FtsW/RodA/SpoVE family cell cycle protein [Streptomyces sp. NPDC059161]. | 56..62 |
| rs:WP_397828714 | [WP_397828714] LysE family translocator [Streptomyces althioticus]. | 44..50 |
| rs:WP_351674725 | [WP_351674725] MULTISPECIES: FtsW/RodA/SpoVE family cell cycle protein [unclassified Streptomyces]. | 56..62 |
| rs:WP_386004677 | [WP_386004677] FtsW/RodA/SpoVE family cell cycle protein [Streptomyces violascens]. | 56..62 |
| rs:WP_312549172 | [WP_312549172] MULTISPECIES: DUF192 domain-containing protein [Massilia]. | 67..73 |
| rs:WP_398872053 | [WP_398872053] FtsW/RodA/SpoVE family cell cycle protein [Streptomyces sp. NPDC020379]. | 35..41 |
| rs:WP_384101391 | [WP_384101391] hypothetical protein [Streptomyces sp. NPDC057555]. | 460..466 |
| rs:WP_388412602 | [WP_388412602] FtsW/RodA/SpoVE family cell cycle protein [Streptomyces sp. NPDC007172]. | 56..62 |
| rs:WP_378359937 | [WP_378359937] GNAT family N-acetyltransferase [Amycolatopsis sp. NPDC059657]. | 271..277 |
| rs:WP_396527014 | [WP_396527014] hypothetical protein [Kutzneria sp. NPDC052558]. | 28..34 |
| rs:WP_400099224 | [WP_400099224] LysE family translocator [Streptomyces sp. NPDC052092]. | 44..50 |
| rs:WP_379510389 | [WP_379510389] hypothetical protein [Novosphingobium bradum]. | 66..72 |
| rs:WP_379872544 | [WP_379872544] hypothetical protein [Marinactinospora rubrisoli]. | 436..442 |
| rs:WP_379436647 | [WP_379436647] MULTISPECIES: DUF5134 domain-containing protein [unclassified Nocardia]. | 184..190 |
| rs:WP_381618882 | [WP_381618882] DUF664 domain-containing protein [Streptomyces flavalbus]. | 93..99 |
| rs:WP_400002083 | [WP_400002083] LysE family translocator [Streptomyces sp. NPDC085937]. | 44..50 |
| rs:WP_374650736 | [WP_374650736] glutamate--tRNA ligase [Dongia sp.]. | 270..276 |
| rs:WP_383183075 | [WP_383183075] MULTISPECIES: DeoR/GlpR family DNA-binding transcription regulator [unclassified Streptomyces]. | 113..119 |
| rs:WP_400044276 | [WP_400044276] LysE family translocator [Streptomyces sp. NPDC052492]. | 44..50 |
| rs:WP_388982260 | [WP_388982260] FtsW/RodA/SpoVE family cell cycle protein [Streptomyces sp. NPDC001691]. | 56..62 |
| rs:WP_380802425 | [WP_380802425] LysE family translocator [Streptomyces albogriseolus]. | 44..50 |
| rs:WP_391612746 | [WP_391612746] LysE family translocator [Streptomyces albogriseolus]. | 44..50 |
| rs:WP_060910785 | [WP_060910785] MULTISPECIES: hypothetical protein [Bradyrhizobium]. | 341..347 |
| rs:WP_397631312 | [WP_397631312] LysE family translocator [Streptomyces albogriseolus]. | 44..50 |
| rs:WP_394215508 | [WP_394215508] Na+/H+ antiporter subunit A [Brachybacterium vulturis]. | 486..492 |
| rs:WP_385520781 | [WP_385520781] MULTISPECIES: FtsW/RodA/SpoVE family cell cycle protein [unclassified Streptomyces]. | 56..62 |
| rs:WP_379940807 | [WP_379940807] MFS transporter [Ensifer sp. P24N7]. | 187..193 |
| rs:WP_379463631 | [WP_379463631] ABC1 kinase family protein [Nocardia sp. NPDC059154]. | 463..469 |
| rs:WP_397639236 | [WP_397639236] LysE family translocator [Streptomyces albogriseolus]. | 44..50 |
| rs:WP_380648853 | [WP_380648853] sensor histidine kinase [Kitasatospora sp. NPDC056446]. | 231..237 |
| rs:WP_389375562 | [WP_389375562] FtsW/RodA/SpoVE family cell cycle protein [Streptomyces sp. NPDC002221]. | 56..62 |
| rs:WP_359602931 | [WP_359602931] MULTISPECIES: LysE family translocator [Streptomyces]. | 44..50 |
| rs:WP_396625071 | [WP_396625071] Bug family tripartite tricarboxylate transporter substrate binding protein [Luteitalea sp.]. | 106..112 |
| rs:WP_375225781 | [WP_375225781] glycosyltransferase family 87 protein [Burkholderia orbicola]. | 385..391 |
| rs:WP_391669571 | [WP_391669571] LysE family translocator [Streptomyces werraensis]. | 44..50 |
| rs:WP_385533049 | [WP_385533049] hypothetical protein [Streptomyces sp. NPDC059708]. | 14..20 |
| rs:WP_387841454 | [WP_387841454] LysE family translocator [Streptomyces althioticus]. | 44..50 |
| rs:WP_379445672 | [WP_379445672] AarF/UbiB family protein, partial [Nocardia sp. NPDC059246]. | 260..266 |
| rs:WP_383229431 | [WP_383229431] DeoR/GlpR family DNA-binding transcription regulator [Streptomyces sp. NPDC058372]. | 113..119 |
| rs:WP_384494656 | [WP_384494656] FtsW/RodA/SpoVE family cell cycle protein [Streptomyces sp. NPDC058961]. | 56..62 |
| rs:WP_234506276 | [WP_234506276] MULTISPECIES: S9 family peptidase [Thermus]. | 84..90 |
| rs:WP_199716164 | [WP_199716164] sugar ABC transporter ATP-binding protein [Aeromicrobium endophyticum]. | 101..107 |
| rs:WP_038372287 | [WP_038372287] Na+/H+ antiporter subunit A [Brachybacterium phenoliresistens]. | 489..495 |
| rs:WP_114355731 | [WP_114355731] Na+/H+ antiporter subunit A [Brachybacterium sp. JB7]. | 489..495 |
| rs:WP_100006770 | [WP_100006770] hypothetical protein [Staphylococcus pseudintermedius]. | 18..24 |
| rs:WP_183026377 | [WP_183026377] tripartite tricarboxylate transporter substrate binding protein [Variovorax sp. UMC13]. | 96..102 |
| rs:WP_302369294 | [WP_302369294] ABC transporter ATP-binding protein [Mycobacteroides abscessus]. | 259..265 |
| rs:WP_285574609 | [WP_285574609] cobyric acid synthase [Geothrix limicola]. | 229..235 |
| rs:WP_176485599 | [WP_176485599] Na+/H+ antiporter subunit A [Brachybacterium alimentarium]. | 489..495 |
| rs:WP_009482939 | [WP_009482939] ATP-dependent DNA helicase RecQ [Mobilicoccus pelagius]. | 268..274 |
| rs:WP_333985836 | [WP_333985836] glycosyltransferase family 87 protein [Burkholderia orbicola]. | 385..391 |
| rs:WP_333989876 | [WP_333989876] glycosyltransferase family 87 protein [Burkholderia orbicola]. | 385..391 |
| rs:WP_334006028 | [WP_334006028] glycosyltransferase family 87 protein [Burkholderia cepacia]. | 385..391 |
| rs:WP_334066184 | [WP_334066184] glycosyltransferase family 87 protein [Burkholderia cepacia]. | 385..391 |

| | | |
|---|---|---|
| rs:WP_202587168 | [WP_202587168] MULTISPECIES: glycosyltransferase family 87 protein [Burkholderia]. | 385..391 |
| rs:WP_212184276 | [WP_212184276] MULTISPECIES: glycosyltransferase family 87 protein [Burkholderia cepacia complex]. | 385..391 |
| rs:WP_185684322 | [WP_185684322] TetR/AcrR family transcriptional regulator [Novosphingobium aerophilum]. | 146..152 |
| rs:WP_131104156 | [WP_131104156] DUF4192 domain-containing protein [Ornithinimicrobium sufpigmenti]. | 26..32 |
| rs:WP_161272952 | [WP_161272952] MULTISPECIES: DeoR/GlpR family DNA-binding transcription regulator [Streptomyces]. | 112..118 |
| rs:WP_343491836 | [WP_343491836] glycosyltransferase family 87 protein [Burkholderia sp. GS2Y]. | 385..391 |
| rs:WP_261675413 | [WP_261675413] MULTISPECIES: LysE family translocator [Streptomyces]. | 44..50 |
| rs:WP_163087453 | [WP_163087453] LysE family translocator [Actinospica acidiphila]. | 44..50 |
| rs:WP_349643908 | [WP_349643908] hypothetical protein [Bradyrhizobium sp. LTSP885]. | 291..297 |
| rs:WP_366281550 | [WP_366281550] LysE family translocator [Streptomyces sp. NPDC093594]. | 44..50 |
| rs:WP_366334706 | [WP_366334706] hypothetical protein [Streptomyces sp. NPDC093707]. | 460..466 |
| rs:WP_366428392 | [WP_366428392] FtsW/RodA/SpoVE family cell cycle protein [Streptomyces sp. NPDC051173]. | 35..41 |
| rs:WP_046142206 | [WP_046142206] ABC transporter permease [Devosia soli]. | 5..11 |
| rs:WP_061910567 | [WP_061910567] ABC transporter permease [Devosia sp. Leaf420]. | 5..11 |
| rs:WP_062631639 | [WP_062631639] ABC transporter permease [Devosia sp. Leaf64]. | 5..11 |
| rs:WP_295412555 | [WP_295412555] ABC transporter permease [Devosia sp.]. | 5..11 |
| rs:WP_188568758 | [WP_188568758] PaaI family thioesterase [Undibacterium terreum]. | 39..45 |
| rs:WP_343711672 | [WP_343711672] EAL domain-containing protein [Kosakonia radicincitans]. | 350..356 |
| rs:WP_343873390 | [WP_343873390] Na+/H+ antiporter subunit A [Brachybacterium alimentarium]. | 489..495 |
| rs:WP_343913617 | [WP_343913617] hypothetical protein [Aquimarina litoralis]. | 24..30 |
| rs:WP_343932877 | [WP_343932877] Uma2 family endonuclease [Saccharothrix mutabilis]. | 13..19 |
| rs:WP_344832805 | [WP_344832805] hypothetical protein [Nonomuraea dietziae]. | 390..396 |
| rs:WP_094804071 | [WP_094804071] asparagine synthase (glutamine-hydrolyzing) [Bordetella genomosp. 5]. | 339..345 |
| rs:WP_094833094 | [WP_094833094] asparagine synthase (glutamine-hydrolyzing) [Bordetella genomosp. 1]. | 339..345 |
| rs:WP_274966742 | [WP_274966742] hypothetical protein [Acidibrevibacterium fodinaquatile]. | 102..108 |
| rs:WP_274966748 | [WP_274966748] hypothetical protein [Acidibrevibacterium fodinaquatile]. | 100..106 |
| rs:WP_297365348 | [WP_297365348] hypothetical protein [Acidiferrobacter sp.]. | 466..472 |
| rs:WP_297388678 | [WP_297388678] hypothetical protein [Acidiferrobacter sp.]. | 466..472 |
| rs:WP_114396359 | [WP_114396359] hypothetical protein [Marinitenerispora sediminis]. | 438..444 |
| rs:WP_312627041 | [WP_312627041] MFS transporter [Pseudofrankia sp. BMG5.37]. | 353..359 |
| rs:WP_233805132 | [WP_233805132] tetratricopeptide repeat protein [Paraburkholderia sp. HP33-1]. | 348..354 |
| rs:WP_183645682 | [WP_183645682] lipopolysaccharide biosynthesis protein [Nonomuraea dietziae]. | 447..453 |
| rs:WP_346004405 | [WP_346004405] hypothetical protein [Gluconobacter sp. OJB]. | 194..200 |
| rs:WP_136924423 | [WP_136924423] hypothetical protein [Polyangium aurulentum]. | 136..142 |
| rs:WP_346124023 | [WP_346124023] hypothetical protein [Nonomuraea roseola]. | 447..453 |
| rs:WP_346143333 | [WP_346143333] hypothetical protein [Nonomuraea recticatena]. | 447..453 |
| rs:WP_137936088 | [WP_137936088] hypothetical protein [Chitinivorax sp. B]. | 24..30 |
| rs:WP_352056125 | [WP_352056125] LysE family translocator [Streptomyces werraensis]. | 44..50 |
| rs:WP_352170069 | [WP_352170069] LysE family translocator [Streptomyces sp. NPDC096538]. | 44..50 |
| rs:WP_033949099 | [WP_033949099] MULTISPECIES: DeoR/GlpR family DNA-binding transcription regulator [Streptomyces]. | 112..118 |
| rs:WP_345052015 | [WP_345052015] hypothetical protein [Streptomyces rameus]. | 35..41 |
| rs:WP_345258590 | [WP_345258590] FtsW/RodA/SpoVE family cell cycle protein [Streptomyces hundungensis]. | 56..62 |
| rs:WP_354762318 | [WP_354762318] LysE family translocator [Streptomyces albogriseolus]. | 44..50 |
| rs:WP_354792250 | [WP_354792250] DeoR/GlpR family DNA-binding transcription regulator [Streptomyces albidoflavus]. | 112..118 |
| rs:WP_354848154 | [WP_354848154] DUF5134 domain-containing protein [Nocardia sp. NPDC004860]. | 184..190 |
| rs:WP_354922533 | [WP_354922533] hypothetical protein [Nonomuraea sp. NPDC003804]. | 419..425 |
| rs:WP_290259522 | [WP_290259522] dethiobiotin synthase [Simiduia curdlanivorans]. | 202..208 |
| rs:WP_260791898 | [WP_260791898] lipid-binding SYLF domain-containing protein [Occallatibacter riparius]. | 59..65 |
| rs:WP_094828706 | [WP_094828706] asparagine synthase (glutamine-hydrolyzing) [Bordetella genomosp. 1]. | 339..345 |
| rs:WP_183562550 | [WP_183562550] endospore germination permease [Paenibacillus endophyticus]. | 207..213 |
| rs:WP_237442700 | [WP_237442700] lipoprotein-releasing ABC transporter permease subunit [Sinobacterium norvegicum]. | 60..66 |
| rs:WP_244885692 | [WP_244885692] MULTISPECIES: GGDEF domain-containing protein [Sphingomonas]. | 75..81 |
| rs:WP_100573630 | [WP_100573630] FtsW/RodA/SpoVE family cell cycle protein [Streptomyces sp. CB01201]. | 56..62 |
| rs:WP_114040258 | [WP_114040258] FtsW/RodA/SpoVE family cell cycle protein [Streptomyces sp. SDr-06]. | 56..62 |
| rs:WP_246470374 | [WP_246470374] FtsW/RodA/SpoVE family cell cycle protein [Streptomyces olivoverticillatus]. | 79..85 |
| rs:WP_193204536 | [WP_193204536] dihydrodipicolinate synthase family protein [Microbispora sitophila]. | 154..160 |
| rs:WP_353859152 | [WP_353859152] tRNA lysidine(34) synthetase TilS [Azospirillum formosense]. | 406..412 |
| rs:WP_322037540 | [WP_322037540] glycosyltransferase family 87 protein [Burkholderia cepacia]. | 385..391 |
| rs:WP_322076707 | [WP_322076707] glycosyltransferase family 87 protein [Burkholderia cepacia]. | 385..391 |
| rs:WP_322078871 | [WP_322078871] glycosyltransferase family 87 protein [Burkholderia cepacia]. | 385..391 |
| rs:WP_102921921 | [WP_102921921] FtsW/RodA/SpoVE family cell cycle protein [Streptomyces eurocidicus]. | 35..41 |
| rs:WP_092187820 | [WP_092187820] hypothetical protein [Bradyrhizobium sp. cf659]. | 341..347 |
| rs:WP_301760728 | [WP_301760728] glycosyltransferase family 87 protein [Burkholderia orbicola]. | 385..391 |
| rs:WP_301762261 | [WP_301762261] glycosyltransferase family 87 protein [Burkholderia cepacia]. | 385..391 |
| rs:WP_301778533 | [WP_301778533] glycosyltransferase family 87 protein [Burkholderia orbicola]. | 385..391 |
| rs:WP_301824871 | [WP_301824871] glycosyltransferase family 87 protein [Burkholderia cepacia]. | 385..391 |
| rs:WP_320586126 | [WP_320586126] LysE family translocator [Streptomyces sp. CL7]. | 44..50 |
| rs:WP_213155456 | [WP_213155456] leucine-rich repeat domain-containing protein [Neochlamydia sp. AcF65]. | 648..654 |
| rs:WP_324763376 | [WP_324763376] MFS transporter [Sinorhizobium meliloti]. | 187..193 |
| rs:WP_189365549 | [WP_189365549] LysE family translocator [Streptomyces variabilis]. | 44..50 |
| rs:WP_190005294 | [WP_190005294] LysE family translocator [Streptomyces werraensis]. | 44..50 |
| rs:WP_193469397 | [WP_193469397] LysE family translocator [Streptomyces althioticus]. | 44..50 |

| | | |
|---|---|---|
| rs:WP_199205649 | [WP_199205649] MULTISPECIES: LysE family translocator [Streptomyces]. | 44..50 |
| rs:WP_199214788 | [WP_199214788] LysE family translocator [Streptomyces sp. BSE7-9]. | 44..50 |
| rs:WP_203349549 | [WP_203349549] LysE family translocator [Streptomyces sp. S-9]. | 44..50 |
| rs:WP_215213831 | [WP_215213831] MULTISPECIES: LysE family translocator [unclassified Streptomyces]. | 44..50 |
| rs:WP_216716340 | [WP_216716340] LysE family translocator [Streptomyces sp. PAM3C]. | 44..50 |
| rs:WP_225626974 | [WP_225626974] LysE family translocator [Streptomyces werraensis]. | 44..50 |
| rs:WP_225644661 | [WP_225644661] LysE family translocator [Streptomyces werraensis]. | 44..50 |
| rs:WP_225654317 | [WP_225654317] LysE family translocator [Streptomyces pseudogriseolus]. | 44..50 |
| rs:WP_242592017 | [WP_242592017] LysE family translocator [Streptomyces sp. GB4-14]. | 44..50 |
| rs:WP_244788928 | [WP_244788928] MULTISPECIES: LysE family translocator [Streptomyces]. | 44..50 |
| rs:WP_252541938 | [WP_252541938] LysE family translocator [Streptomyces sp. RO-S4]. | 44..50 |
| rs:WP_006133283 | [WP_006133283] MULTISPECIES: LysE family translocator [Streptomyces]. | 44..50 |
| rs:WP_033272626 | [WP_033272626] MULTISPECIES: LysE family translocator [Actinomycetes]. | 44..50 |
| rs:WP_048460485 | [WP_048460485] MULTISPECIES: LysE family translocator [unclassified Streptomyces]. | 44..50 |
| rs:WP_102640359 | [WP_102640359] LysE family translocator [Streptomyces sp. SMS_SU21]. | 44..50 |
| rs:WP_114874421 | [WP_114874421] MULTISPECIES: LysE family translocator [unclassified Streptomyces]. | 44..50 |
| rs:WP_121720447 | [WP_121720447] MULTISPECIES: LysE family translocator [unclassified Streptomyces]. | 44..50 |
| rs:WP_145829628 | [WP_145829628] MULTISPECIES: LysE family translocator [Streptomyces]. | 44..50 |
| rs:WP_167745134 | [WP_167745134] MULTISPECIES: LysE family translocator [Streptomyces]. | 44..50 |
| rs:WP_093765309 | [WP_093765309] LysE family translocator [Streptomyces sp. F-7]. | 44..50 |
| rs:WP_329784599 | [WP_329784599] asparagine synthase (glutamine-hydrolyzing) [Microvirga sp. CF3016]. | 353..359 |
| rs:WP_192774176 | [WP_192774176] lipopolysaccharide biosynthesis protein [Nonomuraea africana]. | 419..425 |
| rs:WP_094859128 | [WP_094859128] asparagine synthase (glutamine-hydrolyzing) [Bordetella genomosp. 5]. | 339..345 |
| rs:WP_371668138 | [WP_371668138] ABC transporter permease [Streptomyces sp. NBC_00289]. | 82..88 |
| rs:WP_071048061 | [WP_071048061] MFS transporter [Pseudofrankia sp. BMG5.36]. | 349..355 |
| rs:WP_287731759 | [WP_287731759] glycosyltransferase family 87 protein [Burkholderia sp.]. | 385..391 |
| rs:WP_287998997 | [WP_287998997] hypothetical protein [Acidiphilium sp.]. | 102..108 |
| rs:WP_288511154 | [WP_288511154] ABC transporter permease [uncultured Prevotellamassilia sp.]. | 258..264 |
| rs:WP_128416688 | [WP_128416688] asparagine synthase (glutamine-hydrolyzing) [Xanthomonas populi]. | 346..352 |
| rs:WP_205297884 | [WP_205297884] contractile injection system protein, VgrG/Pvc8 family [Pantoea sp. Cy-639]. | 82..88 |
| rs:WP_371141433 | [WP_371141433] glycosyltransferase family 87 protein [Burkholderia cepacia]. | 385..391 |
| rs:WP_295571799 | [WP_295571799] methyl-accepting chemotaxis protein [uncultured Stenotrophomonas sp.]. | 2..8 |
| rs:WP_351926641 | [WP_351926641] ArsA-related P-loop ATPase [Streptomyces sp. NPDC000983]. | 270..276 |
| rs:WP_326308470 | [WP_326308470] DUF5134 domain-containing protein [Nocardia sp. CDC153]. | 203..209 |
| rs:WP_326357397 | [WP_326357397] DUF5134 domain-containing protein [Nocardia sp. CDC160]. | 222..228 |
| rs:WP_356998257 | [WP_356998257] DUF5134 domain-containing protein [Nocardia sp. NPDC046763]. | 184..190 |
| rs:WP_055768366 | [WP_055768366] beta-galactosidase [Arthrobacter sp. Leaf234]. | 460..466 |
| rs:WP_056505245 | [WP_056505245] hypothetical protein [Sphingomonas sp. Leaf22]. | 4..10 |
| rs:WP_181869300 | [WP_181869300] Na+/H+ antiporter subunit A [Brachybacterium alimentarium]. | 489..495 |
| rs:WP_181870153 | [WP_181870153] Na+/H+ antiporter subunit A [Brachybacterium alimentarium]. | 489..495 |
| rs:WP_182869671 | [WP_182869671] hypothetical protein [Bradyrhizobium diazoefficiens]. | 341..347 |
| rs:WP_312717965 | [WP_312717965] DEAD/DEAH box helicase [Mobilicoccus sp.]. | 247..253 |
| rs:WP_313022139 | [WP_313022139] DEAD/DEAH box helicase [Mobilicoccus sp.]. | 247..253 |
| rs:WP_313566642 | [WP_313566642] DEAD/DEAH box helicase [Mobilicoccus sp.]. | 247..253 |
| rs:WP_071453930 | [WP_071453930] NAD(P)H-quinone oxidoreductase subunit 5 [Gloeomargarita lithophora]. | 529..535 |
| rs:WP_165883898 | [WP_165883898] hypothetical protein [Brevibacterium luteolum]. | 148..154 |
| rs:WP_288605808 | [WP_288605808] ABC transporter permease [uncultured Prevotellamassilia sp.]. | 258..264 |
| rs:WP_349791179 | [WP_349791179] LysE family translocator [Streptomyces sp. OP7]. | 44..50 |
| rs:WP_135809154 | [WP_135809154] 3-hydroxyacyl-CoA dehydrogenase NAD-binding domain-containing protein [Brevibacterium sp. S22]. | 420..426 |
| rs:WP_188516494 | [WP_188516494] TIGR01459 family HAD-type hydrolase [Alsobacter metallidurans]. | 67..73 |
| rs:WP_161536149 | [WP_161536149] hypothetical protein [Bradyrhizobium sp. LCT2]. | 341..347 |
| rs:WP_040810116 | [WP_040810116] DUF5134 domain-containing protein [Nocardia concava]. | 192..198 |
| rs:WP_185986995 | [WP_185986995] hypothetical protein [Leucobacter aridicollis]. | 91..97 |
| rs:WP_227785528 | [WP_227785528] hypothetical protein [Hymenobacter sp. BT770]. | 82..88 |
| rs:WP_228118799 | [WP_228118799] hypothetical protein [Gluconobacter japonicus]. | 213..219 |
| rs:WP_330880365 | [WP_330880365] Na+/H+ antiporter subunit A [Brachybacterium sp.]. | 486..492 |
| rs:WP_299607030 | [WP_299607030] hypothetical protein [uncultured Aquimarina sp.]. | 24..30 |
| rs:WP_299607160 | [WP_299607160] ATP-binding protein [uncultured Aquimarina sp.]. | 70..76 |
| rs:WP_299899139 | [WP_299899139] hypothetical protein [uncultured Aquimarina sp.]. | 24..30 |
| rs:WP_337363550 | [WP_337363550] ABC transporter permease [Prevotellamassilia timonensis]. | 258..264 |
| rs:WP_337418593 | [WP_337418593] ABC transporter permease [Prevotellamassilia timonensis]. | 258..264 |
| rs:WP_337481856 | [WP_337481856] ABC transporter permease [Prevotellamassilia timonensis]. | 258..264 |
| rs:WP_337577660 | [WP_337577660] ABC transporter permease [Prevotellamassilia timonensis]. | 258..264 |
| rs:WP_337667040 | [WP_337667040] ABC transporter permease [Prevotellamassilia timonensis]. | 258..264 |
| rs:WP_337773117 | [WP_337773117] ABC transporter permease [Prevotellamassilia timonensis]. | 258..264 |
| rs:WP_240695074 | [WP_240695074] dipeptidyl aminopeptidase [Thermus tengchongensis]. | 80..86 |
| rs:WP_240695271 | [WP_240695271] hypothetical protein [Thermus caldilimi]. | 14..20 |
| rs:WP_240695484 | [WP_240695484] dipeptidyl aminopeptidase [Thermus tengchongensis]. | 80..86 |
| rs:WP_241155524 | [WP_241155524] Na+/H+ antiporter subunit A [Brachybacterium sp. EE-P12]. | 486..492 |
| rs:WP_196286270 | [WP_196286270] sensor histidine kinase [Hymenobacter properus]. | 253..259 |
| rs:WP_106325849 | [WP_106325849] Gfo/Idh/MocA family protein [Actinoplanes italicus]. | 33..39 |
| rs:WP_106264038 | [WP_106264038] LacI family DNA-binding transcriptional regulator [Donghicola tyrosinivorans]. | 64..70 |

| | | |
|---|---|---|
| rs:WP_071122226 | [WP_071122226] ABC transporter permease [Prevotellamassilia timonensis]. | 258..264 |
| rs:WP_011349896 | [WP_011349896] glycosyltransferase family 87 protein [Burkholderia lata]. | 385..391 |
| rs:WP_011549063 | [WP_011549063] MULTISPECIES: glycosyltransferase family 87 protein [Burkholderia]. | 385..391 |
| rs:WP_012336897 | [WP_012336897] MULTISPECIES: glycosyltransferase family 87 protein [Burkholderia cepacia complex]. | 385..391 |
| rs:WP_021156628 | [WP_021156628] MULTISPECIES: glycosyltransferase family 87 protein [Burkholderia]. | 385..391 |
| rs:WP_027792730 | [WP_027792730] glycosyltransferase family 87 protein [Burkholderia cepacia]. | 385..391 |
| rs:WP_027808470 | [WP_027808470] glycosyltransferase family 87 protein [Burkholderia cenocepacia]. | 385..391 |
| rs:WP_027810921 | [WP_027810921] glycosyltransferase family 87 protein [Burkholderia cenocepacia]. | 385..391 |
| rs:WP_137773778 | [WP_137773778] AMP-binding protein [Citricoccus sp. SGAir0253]. | 229..235 |
| rs:WP_303373345 | [WP_303373345] AMP-binding protein [Ornithinimicrobium sp.]. | 226..232 |
| rs:WP_072485339 | [WP_072485339] non-ribosomal peptide synthetase [Streptomyces atratus]. | 1289..1295 |
| rs:WP_018097668 | [WP_018097668] MULTISPECIES: MFS transporter [Sinorhizobium]. | 187..193 |
| rs:WP_317184342 | [WP_317184342] ABC-2 family transporter protein [Devosia sp. BK]. | 5..11 |
| rs:WP_215071423 | [WP_215071423] ATP-binding cassette domain-containing protein [Streptomyces sp. ISL-36]. | 586..592 |
| rs:WP_354575025 | [WP_354575025] vitamin K epoxide reductase family protein [Frigoribacterium sp. UYMn621]. | 108..114 |
| rs:WP_175899552 | [WP_175899552] MULTISPECIES: glycosyltransferase family 87 protein [Burkholderia]. | 385..391 |
| rs:WP_104961162 | [WP_104961162] ATP-binding domain-containing protein [Pseudomonas sp. XWY-1]. | 390..396 |
| rs:WP_363028872 | [WP_363028872] FtsW/RodA/SpoVE family cell cycle protein [Streptomyces sp. NPDC048491]. | 56..62 |
| rs:WP_155356526 | [WP_155356526] hypothetical protein [Acrocarpospora macrocephala]. | 125..131 |
| rs:WP_168197267 | [WP_168197267] MFS transporter [Kribbella sp. ALI-6-A]. | 54..60 |
| rs:WP_115414989 | [WP_115414989] Na+/H+ antiporter subunit A [Brachybacterium saurashtrense]. | 486..492 |
| rs:WP_331734379 | [WP_331734379] amino acid adenylation domain-containing protein [Streptomyces atratus]. | 1289..1295 |
| rs:WP_331765605 | [WP_331765605] amino acid adenylation domain-containing protein [Streptomyces atratus]. | 1289..1295 |
| rs:WP_331955464 | [WP_331955464] GNAT family N-acetyltransferase [Pengzhenrongella sp.]. | 104..110 |
| rs:WP_332537619 | [WP_332537619] tripartite tricarboxylate transporter substrate binding protein [Comamonas sp.]. | 97..103 |
| rs:WP_332547646 | [WP_332547646] tripartite tricarboxylate transporter substrate binding protein [Comamonas sp.]. | 97..103 |
| rs:WP_143047224 | [WP_143047224] GNAT family N-acetyltransferase [Amycolatopsis xylanica]. | 271..277 |
| rs:WP_157899634 | [WP_157899634] tripartite tricarboxylate transporter substrate binding protein [Luteitalea pratensis]. | 204..210 |
| rs:WP_159911108 | [WP_159911108] tripartite tricarboxylate transporter substrate binding protein [Pantoea sp. 18069]. | 97..103 |
| rs:WP_213293411 | [WP_213293411] tripartite tricarboxylate transporter substrate binding protein [Acidovorax sp. CCYZU-2555]. | 97..103 |
| rs:WP_175503086 | [WP_175503086] tripartite tricarboxylate transporter substrate binding protein [Comamonas antarctica]. | 97..103 |
| rs:WP_239489531 | [WP_239489531] tripartite tricarboxylate transporter substrate binding protein [Luteitalea sp. TBR-22]. | 106..112 |
| rs:WP_052541700 | [WP_052541700] ABC transporter ATP-binding protein [Mycobacteroides abscessus]. | 259..265 |
| rs:WP_278135063 | [WP_278135063] glycosyltransferase family 87 protein [Burkholderia contaminans]. | 385..391 |
| rs:WP_278649678 | [WP_278649678] glycosyltransferase family 87 protein [Burkholderia lata]. | 385..391 |
| rs:WP_279329985 | [WP_279329985] LysE family translocator [Streptomyces sp. OS603R]. | 44..50 |
| rs:WP_204163849 | [WP_204163849] TetR/AcrR family transcriptional regulator [Nocardioides solisilvae]. | 106..112 |
| rs:WP_310324270 | [WP_310324270] MgtC/SapB family protein [Roseateles asaccharophilus]. | 145..151 |
| rs:WP_310624708 | [WP_310624708] glycosyltransferase family 87 protein [Burkholderia cenocepacia]. | 385..391 |
| rs:WP_310639972 | [WP_310639972] glycosyltransferase family 87 protein [Burkholderia cenocepacia]. | 385..391 |
| rs:WP_283407162 | [WP_283407162] GGDEF domain-containing protein [Novosphingobium panipatense]. | 75..81 |
| rs:WP_231283807 | [WP_231283807] MULTISPECIES: MFS transporter [unclassified Rhizobium]. | 215..221 |
| rs:WP_011085162 | [WP_011085162] MULTISPECIES: hypothetical protein [Bradyrhizobium]. | 341..347 |
| rs:WP_213013576 | [WP_213013576] GPP34 family phosphoprotein [Paractinoplanes toevensis]. | 214..220 |
| rs:WP_234834400 | [WP_234834400] MFS transporter, partial [Sinorhizobium meliloti]. | 130..136 |
| rs:WP_013844453 | [WP_013844453] MFS transporter [Sinorhizobium meliloti]. | 187..193 |
| rs:WP_014526759 | [WP_014526759] MFS transporter [Sinorhizobium meliloti]. | 187..193 |
| rs:WP_027994073 | [WP_027994073] MFS transporter [Sinorhizobium meliloti]. | 187..193 |
| rs:WP_051077382 | [WP_051077382] MFS transporter [Rhizobium leguminosarum]. | 169..175 |
| rs:WP_065279944 | [WP_065279944] MFS transporter [Rhizobium leguminosarum]. | 169..175 |
| rs:WP_127527462 | [WP_127527462] MFS transporter [Sinorhizobium meliloti]. | 187..193 |
| rs:WP_127541402 | [WP_127541402] MFS transporter [Sinorhizobium meliloti]. | 187..193 |
| rs:WP_127636393 | [WP_127636393] MFS transporter [Sinorhizobium meliloti]. | 187..193 |
| rs:WP_127638374 | [WP_127638374] MFS transporter [Sinorhizobium meliloti]. | 187..193 |
| rs:WP_127666086 | [WP_127666086] MFS transporter [Sinorhizobium meliloti]. | 187..193 |
| rs:WP_127712797 | [WP_127712797] MFS transporter [Sinorhizobium meliloti]. | 187..193 |
| rs:WP_128203614 | [WP_128203614] MFS transporter [Sinorhizobium meliloti]. | 187..193 |
| rs:WP_132523598 | [WP_132523598] MFS transporter [Rhizobium sp. BK376]. | 205..211 |
| rs:WP_146722441 | [WP_146722441] MFS transporter [Sinorhizobium meliloti]. | 187..193 |
| rs:WP_153313695 | [WP_153313695] MFS transporter [Sinorhizobium meliloti]. | 187..193 |
| rs:WP_157718912 | [WP_157718912] MFS transporter [Sinorhizobium meliloti]. | 187..193 |
| rs:WP_157813705 | [WP_157813705] MFS transporter [Sinorhizobium meliloti]. | 187..193 |
| rs:WP_168322101 | [WP_168322101] MFS transporter [Rhizobium leguminosarum]. | 205..211 |
| rs:WP_171602467 | [WP_171602467] MULTISPECIES: MFS transporter [Rhizobium]. | 187..193 |
| rs:WP_180695129 | [WP_180695129] MFS transporter [Rhizobium changzhiense]. | 187..193 |
| rs:WP_222364227 | [WP_222364227] MFS transporter [Rhizobium leguminosarum]. | 169..175 |
| rs:WP_245417639 | [WP_245417639] MFS transporter [Aminobacter sp. AP02]. | 169..175 |
| rs:WP_245500436 | [WP_245500436] MFS transporter [Rhizobium leguminosarum]. | 215..221 |
| rs:WP_246664996 | [WP_246664996] MFS transporter [Neorhizobium sp. P12A]. | 169..175 |
| rs:WP_246802144 | [WP_246802144] MFS transporter [Rhizobium leguminosarum]. | 215..221 |
| rs:WP_042490938 | [WP_042490938] MULTISPECIES: GGDEF domain-containing protein [Bacteria]. | 75..81 |
| rs:WP_369157352 | [WP_369157352] LysE family translocator [Streptomyces sp. R02]. | 44..50 |

| | | |
|---|---|---|
| rs:WP_369325123 | [WP_369325123] glycosyltransferase family 87 protein [Burkholderia cepacia]. | 385..391 |
| rs:WP_056892648 | [WP_056892648] MULTISPECIES: alpha-(1->3)-arabinofuranosyltransferase family protein [unclassified Nocardioides]. | 159..165 |
| rs:WP_091025798 | [WP_091025798] DUF6328 family protein [Nocardioides szechwanensis]. | 145..151 |
| rs:WP_042325584 | [WP_042325584] HAD family hydrolase [Paraburkholderia ginsengisoli]. | 212..218 |
| rs:WP_275597966 | [WP_275597966] MULTISPECIES: MFS transporter [unclassified Sinorhizobium]. | 187..193 |
| rs:WP_170273726 | [WP_170273726] hypothetical protein [Brevibacterium luteolum]. | 148..154 |
| rs:WP_332824719 | [WP_332824719] acetate--CoA ligase [Ramlibacter sp.]. | 24..30 |
| rs:WP_009491680 | [WP_009491680] asparagine synthase (glutamine-hydrolyzing) [Microvirga lotononidis]. | 353..359 |
| rs:WP_305648316 | [WP_305648316] hypothetical protein [Rhodoferax sp.]. | 192..198 |
| rs:WP_008330017 | [WP_008330017] M14-type cytosolic carboxypeptidase [Maritimibacter alkaliphilus]. | 381..387 |
| rs:WP_138423355 | [WP_138423355] M14-type cytosolic carboxypeptidase [Maritimibacter alexandrii]. | 381..387 |
| rs:WP_287436025 | [WP_287436025] M14-type cytosolic carboxypeptidase [Maritimibacter sp.]. | 381..387 |
| rs:WP_288929052 | [WP_288929052] M14-type cytosolic carboxypeptidase [uncultured Maritimibacter sp.]. | 381..387 |
| rs:WP_358874298 | [WP_358874298] LysE family translocator [Streptomyces sp. NPDC005908]. | 44..50 |
| rs:WP_211937778 | [WP_211937778] sigma-70 family RNA polymerase sigma factor [Phenylobacterium montanum]. | 13..19 |
| rs:WP_281472870 | [WP_281472870] hypothetical protein [Yinghuangia seranimata]. | 435..441 |
| rs:WP_357163935 | [WP_357163935] ABC transporter ATP-binding protein [Micromonospora sp. NPDC047548]. | 72..78 |
| rs:WP_357582649 | [WP_357582649] hypothetical protein [Micromonospora sp. NPDC049891]. | 156..162 |
| rs:WP_357597183 | [WP_357597183] hypothetical protein [Nonomuraea dietziae]. | 447..453 |
| rs:WP_358151329 | [WP_358151329] hypothetical protein [Nonomuraea dietziae]. | 447..453 |
| rs:WP_358244043 | [WP_358244043] DeoR/GlpR family DNA-binding transcription regulator [Streptomyces albidoflavus]. | 112..118 |
| rs:WP_367426218 | [WP_367426218] LysE family translocator [Streptomyces albogriseolus]. | 44..50 |
| rs:WP_362263493 | [WP_362263493] hypothetical protein [Streptomyces sp. NPDC042638]. | 35..41 |
| rs:WP_301767810 | [WP_301767810] glycosyltransferase family 87 protein [Burkholderia aenigmatica]. | 385..391 |
| rs:WP_301776182 | [WP_301776182] glycosyltransferase family 87 protein [Burkholderia sp. AU45274]. | 385..391 |
| rs:WP_301801738 | [WP_301801738] glycosyltransferase family 87 protein [Burkholderia sp. AU45251]. | 385..391 |
| rs:WP_124480244 | [WP_124480244] MULTISPECIES: glycosyltransferase family 87 protein [Burkholderia cepacia complex]. | 385..391 |
| rs:WP_212206308 | [WP_212206308] MULTISPECIES: glycosyltransferase family 87 protein [Burkholderia cepacia complex]. | 385..391 |
| rs:WP_363919055 | [WP_363919055] hypothetical protein [Micromonospora sp. NPDC049374]. | 121..127 |
| rs:WP_363933930 | [WP_363933930] LysE family translocator [Streptomyces werraensis]. | 44..50 |
| rs:WP_363991299 | [WP_363991299] LysE family translocator [Streptomyces werraensis]. | 44..50 |
| rs:WP_364043792 | [WP_364043792] LysE family translocator [Streptomyces werraensis]. | 44..50 |
| rs:WP_364143160 | [WP_364143160] LysE family translocator [Streptomyces werraensis]. | 44..50 |
| rs:WP_364206342 | [WP_364206342] hypothetical protein [Amycolatopsis sp. NPDC089917]. | 28..34 |
| rs:WP_259799746 | [WP_259799746] hypothetical protein [Brevibacterium luteolum]. | 148..154 |
| rs:WP_259835505 | [WP_259835505] hypothetical protein [Brevibacterium luteolum]. | 148..154 |
| rs:WP_259845580 | [WP_259845580] hypothetical protein [Brevibacterium luteolum]. | 148..154 |
| rs:WP_259855924 | [WP_259855924] hypothetical protein [Brevibacterium luteolum]. | 148..154 |
| rs:WP_259875854 | [WP_259875854] hypothetical protein [Brevibacterium sp. p3-SID960]. | 148..154 |
| rs:WP_073479901 | [WP_073479901] dienelactone hydrolase family protein [Streptoalloteichus hindustanus]. | 58..64 |
| rs:WP_203995778 | [WP_203995778] ABC transporter ATP-binding protein [Virgisporangium aurantiacum]. | 70..76 |
| rs:WP_152812202 | [WP_152812202] beta-galactosidase [Arthrobacter bussei]. | 460..466 |
| rs:WP_005985233 | [WP_005985233] PP2C family protein-serine/threonine phosphatase [Desulfocurvibacter africanus]. | 121..127 |
| rs:WP_014258315 | [WP_014258315] PP2C family protein-serine/threonine phosphatase [Desulfocurvibacter africanus]. | 121..127 |
| rs:WP_027367875 | [WP_027367875] PP2C family protein-serine/threonine phosphatase [Desulfocurvibacter africanus]. | 121..127 |
| rs:WP_291986577 | [WP_291986577] tripartite tricarboxylate transporter substrate binding protein [Luteitalea sp.]. | 86..92 |
| rs:WP_110137517 | [WP_110137517] hypothetical protein [Acidiferrobacter sp. SPIII_3]. | 466..472 |
| rs:WP_286994809 | [WP_286994809] hypothetical protein [Acinetobacter sp.]. | 123..129 |
| rs:WP_364896329 | [WP_364896329] hypothetical protein [Nonomuraea dietziae]. | 419..425 |
| rs:WP_365020664 | [WP_365020664] hypothetical protein [Nonomuraea dietziae]. | 447..453 |
| rs:WP_151643513 | [WP_151643513] hypothetical protein [Bradyrhizobium betae]. | 341..347 |
| rs:WP_188509008 | [WP_188509008] DUF4192 family protein [Conyzicola nivalis]. | 13..19 |
| rs:WP_344257045 | [WP_344257045] DUF4192 family protein [Terrabacter carboxydivorans]. | 15..21 |
| rs:WP_366634648 | [WP_366634648] ATP-binding protein [Streptomyces griseorubiginosus]. | 642..648 |
| rs:WP_067410210 | [WP_067410210] MULTISPECIES: DeoR/GlpR family DNA-binding transcription regulator [Streptomyces]. | 112..118 |
| rs:WP_162568264 | [WP_162568264] response regulator transcription factor [Variovorax sp. SRS16]. | 29..35 |
| rs:WP_162593278 | [WP_162593278] response regulator transcription factor [Variovorax sp. PBL-E5]. | 29..35 |
| rs:WP_068632018 | [WP_068632018] response regulator transcription factor [Variovorax sp. PAMC 28711]. | 27..33 |
| rs:WP_191725323 | [WP_191725323] hypothetical protein [Brevibacterium gallinarum]. | 148..154 |
| rs:WP_154788061 | [WP_154788061] DUF5134 domain-containing protein [Nocardia aurantiaca]. | 184..190 |
| rs:WP_158060516 | [WP_158060516] DUF4192 domain-containing protein [Ornithinimicrobium pratense]. | 26..32 |
| rs:WP_194255951 | [WP_194255951] hypothetical protein [Gluconobacter cerevisiae]. | 194..200 |
| rs:WP_138565718 | [WP_138565718] 16S rRNA (guanine(527)-N(7))-methyltransferase RsmG [Thiomicrorhabdus sediminis]. | 62..68 |
| rs:WP_168986589 | [WP_168986589] 16S rRNA (guanine(527)-N(7))-methyltransferase RsmG [Azoarcus taiwanensis]. | 60..66 |
| rs:WP_232059330 | [WP_232059330] 16S rRNA (guanine(527)-N(7))-methyltransferase RsmG [Kineobactrum salinum]. | 75..81 |
| rs:WP_201499739 | [WP_201499739] 16S rRNA (guanine(527)-N(7))-methyltransferase RsmG [Psychrobacter arenosus]. | 68..74 |
| rs:WP_236241979 | [WP_236241979] hypothetical protein [Streptomyces sp. CC228A]. | 80..86 |
| rs:WP_358145657 | [WP_358145657] BTAD domain-containing putative transcriptional regulator [Nonomuraea sp. NPDC049152]. | 121..127 |
| rs:WP_234553590 | [WP_234553590] S9 family peptidase [Thermus caliditerrae]. | 80..86 |
| rs:WP_234558076 | [WP_234558076] S9 family peptidase [Thermus tengchongensis]. | 80..86 |
| rs:WP_326849664 | [WP_326849664] glycosyltransferase family 87 protein [Burkholderia cenocepacia]. | 385..391 |
| rs:WP_153432023 | [WP_153432023] hypothetical protein [Gluconobacter aidae]. | 194..200 |

| | | |
|---|---|---|
| rs:WP_313749686 | [WP_313749686] universal stress protein [Streptomyces sp. Li-HN-5-11]. | 307..313 |
| rs:WP_296670873 | [WP_296670873] XrtA system polysaccharide chain length determinant [Sulfuricaulis sp.]. | 467..473 |
| rs:WP_129338301 | [WP_129338301] GNAT family N-acetyltransferase [Cellulomonas endophytica]. | 104..110 |
| rs:WP_033048658 | [WP_033048658] MULTISPECIES: MFS transporter [Sinorhizobium]. | 187..193 |
| rs:WP_276995178 | [WP_276995178] ABC transporter permease [Prevotellamassilia timonensis]. | 258..264 |
| rs:WP_369586608 | [WP_369586608] hypothetical protein [Kingella oralis]. | 6..12 |
| rs:WP_369633977 | [WP_369633977] DUF5134 domain-containing protein [Nocardia sp. JMUB6875]. | 206..212 |
| rs:WP_338294885 | [WP_338294885] molybdopterin cofactor-binding domain-containing protein [Planctobacterium marinum]. | 38..44 |
| rs:WP_084257055 | [WP_084257055] phosphotransferase [Pasteurella testudinis]. | 127..133 |
| rs:WP_107971901 | [WP_107971901] bifunctional diaminohydroxyphosphoribosylaminopyrimidine deaminase/5-amino-6-(5-phosphoribosylamino)uracil reductase RibD [Neisseria elongata]. | 250..256 |
| rs:WP_259307806 | [WP_259307806] GNAT family N-acetyltransferase [Cellulomonas sp. P24]. | 104..110 |
| rs:WP_269844666 | [WP_269844666] DUF6236 family protein [Actinophytocola xanthii]. | 168..174 |
| rs:WP_314849547 | [WP_314849547] DNA methyltransferase [Rothia mucilaginosa]. | 816..822 |
| rs:WP_046546980 | [WP_046546980] glycosyltransferase family 87 protein [Burkholderia contaminans]. | 385..391 |
| rs:WP_047853268 | [WP_047853268] glycosyltransferase family 87 protein [Burkholderia contaminans]. | 385..391 |
| rs:WP_048027266 | [WP_048027266] MULTISPECIES: glycosyltransferase family 87 protein [Burkholderia]. | 385..391 |
| rs:WP_048251872 | [WP_048251872] glycosyltransferase family 87 protein [Burkholderia cepacia]. | 385..391 |
| rs:WP_059485625 | [WP_059485625] glycosyltransferase family 87 protein [Burkholderia cepacia]. | 385..391 |
| rs:WP_059524189 | [WP_059524189] glycosyltransferase family 87 protein [Burkholderia cepacia]. | 385..391 |
| rs:WP_059555507 | [WP_059555507] glycosyltransferase family 87 protein [Burkholderia cepacia]. | 385..391 |
| rs:WP_059587472 | [WP_059587472] MULTISPECIES: glycosyltransferase family 87 protein [Burkholderia]. | 385..391 |
| rs:WP_059666431 | [WP_059666431] glycosyltransferase family 87 protein [Burkholderia cepacia]. | 385..391 |
| rs:WP_059675388 | [WP_059675388] glycosyltransferase family 87 protein [Burkholderia cepacia]. | 385..391 |
| rs:WP_059688227 | [WP_059688227] glycosyltransferase family 87 protein [Burkholderia cepacia]. | 385..391 |
| rs:WP_059699653 | [WP_059699653] glycosyltransferase family 87 protein [Burkholderia cepacia]. | 385..391 |
| rs:WP_059710581 | [WP_059710581] glycosyltransferase family 87 protein [Burkholderia cepacia]. | 385..391 |
| rs:WP_059733139 | [WP_059733139] glycosyltransferase family 87 protein [Burkholderia cepacia]. | 385..391 |
| rs:WP_059813312 | [WP_059813312] glycosyltransferase family 87 protein [Burkholderia cepacia]. | 385..391 |
| rs:WP_059855760 | [WP_059855760] glycosyltransferase family 87 protein [Burkholderia cepacia]. | 385..391 |
| rs:WP_059902890 | [WP_059902890] glycosyltransferase family 87 protein [Burkholderia cepacia]. | 385..391 |
| rs:WP_060063775 | [WP_060063775] glycosyltransferase family 87 protein [Burkholderia cepacia]. | 385..391 |
| rs:WP_060084641 | [WP_060084641] glycosyltransferase family 87 protein [Burkholderia cepacia]. | 385..391 |
| rs:WP_060088793 | [WP_060088793] glycosyltransferase family 87 protein [Burkholderia cepacia]. | 385..391 |
| rs:WP_060125224 | [WP_060125224] glycosyltransferase family 87 protein [Burkholderia cepacia]. | 385..391 |
| rs:WP_060148134 | [WP_060148134] glycosyltransferase family 87 protein [Burkholderia cepacia]. | 385..391 |
| rs:WP_060174395 | [WP_060174395] glycosyltransferase family 87 protein [Burkholderia cepacia]. | 385..391 |
| rs:WP_060197037 | [WP_060197037] glycosyltransferase family 87 protein [Burkholderia cepacia]. | 385..391 |
| rs:WP_060220852 | [WP_060220852] glycosyltransferase family 87 protein [Burkholderia cepacia]. | 385..391 |
| rs:WP_060223641 | [WP_060223641] glycosyltransferase family 87 protein [Burkholderia cepacia]. | 385..391 |
| rs:WP_060309410 | [WP_060309410] glycosyltransferase family 87 protein [Burkholderia cepacia]. | 385..391 |
| rs:WP_060316776 | [WP_060316776] glycosyltransferase family 87 protein [Burkholderia cepacia]. | 385..391 |
| rs:WP_060325613 | [WP_060325613] glycosyltransferase family 87 protein [Burkholderia cepacia]. | 385..391 |
| rs:WP_060342701 | [WP_060342701] glycosyltransferase family 87 protein [Burkholderia cepacia]. | 385..391 |
| rs:WP_060358078 | [WP_060358078] glycosyltransferase family 87 protein [Burkholderia cepacia]. | 385..391 |
| rs:WP_060372596 | [WP_060372596] glycosyltransferase family 87 protein [Burkholderia cepacia]. | 385..391 |
| rs:WP_359520265 | [WP_359520265] LysE family translocator [Streptomyces althioticus]. | 44..50 |
| rs:WP_359744471 | [WP_359744471] LysE family translocator [Streptomyces althioticus]. | 44..50 |
| rs:WP_333525568 | [WP_333525568] SpoIIE family protein phosphatase [Desulfocurvibacter africanus]. | 121..127 |
| rs:WP_327753147 | [WP_327753147] hypothetical protein [Sphingobium sp. SJ10-10]. | 75..81 |
| rs:WP_204664666 | [WP_204664666] aminopeptidase P family protein [Fusibacter tunisiensis]. | 520..526 |
| rs:WP_360376478 | [WP_360376478] hypothetical protein [Dactylosporangium sp. NPDC049140]. | 934..940 |
| rs:WP_069750563 | [WP_069750563] glycosyltransferase family 87 protein [Burkholderia stabilis]. | 385..391 |
| rs:WP_071335923 | [WP_071335923] glycosyltransferase family 87 protein [Burkholderia contaminans]. | 385..391 |
| rs:WP_074807817 | [WP_074807817] glycosyltransferase family 87 protein [Burkholderia cenocepacia]. | 385..391 |
| rs:WP_077179928 | [WP_077179928] glycosyltransferase family 87 protein [Burkholderia cenocepacia]. | 385..391 |
| rs:WP_077217502 | [WP_077217502] glycosyltransferase family 87 protein [Burkholderia cenocepacia]. | 385..391 |
| rs:WP_082303168 | [WP_082303168] DeoR/GlpR family DNA-binding transcription regulator, partial [Streptomyces wadayamensis]. | 112..118 |
| rs:WP_089444321 | [WP_089444321] MULTISPECIES: glycosyltransferase family 87 protein [Burkholderia]. | 385..391 |
| rs:WP_007372467 | [WP_007372467] MULTISPECIES: EAL domain-containing protein [Kosakonia]. | 350..356 |
| rs:WP_039055714 | [WP_039055714] EAL domain-containing protein [Enterobacter sp. Bisph1]. | 350..356 |
| rs:WP_043955652 | [WP_043955652] EAL domain-containing protein [Kosakonia radicincitans]. | 350..356 |
| rs:WP_064568784 | [WP_064568784] EAL domain-containing protein [Kosakonia oryzae]. | 350..356 |
| rs:WP_090121870 | [WP_090121870] EAL domain-containing protein [Kosakonia arachidis]. | 351..357 |
| rs:WP_208657115 | [WP_208657115] EAL domain-containing protein [Kosakonia sp. MUSA4]. | 350..356 |
| rs:WP_264373430 | [WP_264373430] EAL domain-containing protein [Kosakonia radicincitans]. | 233..239 |
| rs:WP_229668354 | [WP_229668354] GGDEF domain-containing protein [Stakelama pacifica]. | 75..81 |
| rs:WP_231727546 | [WP_231727546] MULTISPECIES: GGDEF domain-containing protein [unclassified Sphingomonas]. | 8..14 |
| rs:WP_233420128 | [WP_233420128] GGDEF domain-containing protein [Sphingomonas paucimobilis]. | 8..14 |
| rs:WP_238320221 | [WP_238320221] GGDEF domain-containing protein [Sphingobium sp. YBL2]. | 8..14 |
| rs:WP_239436248 | [WP_239436248] GGDEF domain-containing protein [Sphingomonas sp. ACRSK]. | 75..81 |
| rs:WP_241212735 | [WP_241212735] GGDEF domain-containing protein [Sphingomonas sp. ABOLG]. | 26..32 |

| | | |
|---|---|---|
| rs:WP_255504931 | [WP_255504931] GGDEF domain-containing protein [Novosphingobium sp. EMRT-2]. | 8..14 |
| rs:WP_066550607 | [WP_066550607] MULTISPECIES: GGDEF domain-containing protein [Sphingomonadaceae]. | 75..81 |
| rs:WP_089429732 | [WP_089429732] glycosyltransferase family 87 protein [Burkholderia sp. HI2500]. | 385..391 |
| rs:WP_089433737 | [WP_089433737] glycosyltransferase family 87 protein [Burkholderia sp. AU31652]. | 385..391 |
| rs:WP_089448728 | [WP_089448728] glycosyltransferase family 87 protein [Burkholderia sp. AU33423]. | 385..391 |
| rs:WP_089452627 | [WP_089452627] glycosyltransferase family 87 protein [Burkholderia aenigmatica]. | 385..391 |
| rs:WP_089459443 | [WP_089459443] MULTISPECIES: glycosyltransferase family 87 protein [Burkholderia]. | 385..391 |
| rs:WP_089464052 | [WP_089464052] MULTISPECIES: glycosyltransferase family 87 protein [Burkholderia]. | 385..391 |
| rs:WP_089476200 | [WP_089476200] glycosyltransferase family 87 protein [Burkholderia sp. AU6039]. | 385..391 |
| rs:WP_089487081 | [WP_089487081] MULTISPECIES: glycosyltransferase family 87 protein [Burkholderia]. | 385..391 |
| rs:WP_089498743 | [WP_089498743] glycosyltransferase family 87 protein [Burkholderia sp. AU15512]. | 385..391 |
| rs:WP_091922075 | [WP_091922075] glycosyltransferase family 87 protein [Burkholderia cepacia]. | 385..391 |
| rs:WP_095711822 | [WP_095711822] DeoR/GlpR family DNA-binding transcription regulator [Streptomyces albidoflavus]. | 112..118 |
| rs:WP_016737804 | [WP_016737804] hypothetical protein [Gluconobacter thailandicus]. | 213..219 |
| rs:WP_039355481 | [WP_039355481] MULTISPECIES: glycosyltransferase family 87 protein [Burkholderia]. | 385..391 |
| rs:WP_321880267 | [WP_321880267] glycosyltransferase family 87 protein [Burkholderia cepacia]. | 385..391 |
| rs:WP_321908536 | [WP_321908536] glycosyltransferase family 87 protein [Burkholderia cepacia]. | 385..391 |
| rs:WP_321939193 | [WP_321939193] glycosyltransferase family 87 protein [Burkholderia cepacia]. | 385..391 |
| rs:WP_321953575 | [WP_321953575] glycosyltransferase family 87 protein [Burkholderia cenocepacia]. | 385..391 |
| rs:WP_321965986 | [WP_321965986] glycosyltransferase family 87 protein [Burkholderia cepacia]. | 385..391 |
| rs:WP_337861485 | [WP_337861485] hypothetical protein [Nitrososphaera sp.]. | 181..187 |
| rs:WP_338025022 | [WP_338025022] Na+/H+ antiporter subunit A [Brachybacterium vulturis]. | 537..543 |
| rs:WP_028174318 | [WP_028174318] MULTISPECIES: hypothetical protein [Bradyrhizobium]. | 341..347 |
| rs:WP_104767313 | [WP_104767313] glycosyltransferase family 87 protein [Burkholderia cepacia]. | 385..391 |
| rs:WP_105798116 | [WP_105798116] glycosyltransferase family 87 protein [Burkholderia cenocepacia]. | 385..391 |
| rs:WP_105817113 | [WP_105817113] MULTISPECIES: glycosyltransferase family 87 protein [Burkholderia]. | 385..391 |
| rs:WP_096196259 | [WP_096196259] Na+/H+ antiporter subunit A [Brachybacterium alimentarium]. | 489..495 |
| rs:WP_111941269 | [WP_111941269] glycosyltransferase family 87 protein [Burkholderia cepacia]. | 385..391 |
| rs:WP_323497692 | [WP_323497692] MULTISPECIES: response regulator transcription factor [unclassified Variovorax]. | 27..33 |
| rs:WP_148794314 | [WP_148794314] MULTISPECIES: ABC transporter ATP-binding protein [unclassified Micromonospora]. | 72..78 |
| rs:WP_056397755 | [WP_056397755] MULTISPECIES: hypothetical protein [unclassified Sphingomonas]. | 4..10 |
| rs:WP_140477485 | [WP_140477485] hypothetical protein [Bradyrhizobium symbiodeficiens]. | 341..347 |
| rs:WP_158295518 | [WP_158295518] hypothetical protein [Crenalkalicoccus roseus]. | 167..173 |
| rs:WP_102162388 | [WP_102162388] hypothetical protein [Brevibacterium luteolum]. | 148..154 |
| rs:WP_340625930 | [WP_340625930] glycosyltransferase family 87 protein [Burkholderia arboris]. | 385..391 |
| rs:WP_341065153 | [WP_341065153] GNAT family N-acetyltransferase [Cytobacillus sp. FSL K6-0265]. | 64..70 |
| rs:WP_341381223 | [WP_341381223] glycosyltransferase family 87 protein [Burkholderia contaminans]. | 385..391 |
| rs:WP_341401744 | [WP_341401744] glycosyltransferase family 87 protein, partial [Burkholderia contaminans]. | 301..307 |
| rs:WP_100287786 | [WP_100287786] lipoprotein-releasing ABC transporter permease subunit LolE [Conservatibacter flavescens]. | 58..64 |
| rs:WP_365776916 | [WP_365776916] hypothetical protein [Streptomyces sp. NPDC055692]. | 438..444 |
| rs:WP_366007441 | [WP_366007441] FtsW/RodA/SpoVE family cell cycle protein [Streptomyces sp. NPDC051162]. | 35..41 |
| rs:WP_038040924 | [WP_038040924] S9 family peptidase [Thermus tengchongensis]. | 80..86 |
| rs:WP_038048319 | [WP_038048319] S9 family peptidase [Thermus caliditerrae]. | 80..86 |
| rs:WP_129967692 | [WP_129967692] asparagine synthase (glutamine-hydrolyzing) [Allopusillimonas soli]. | 338..344 |
| rs:WP_067675662 | [WP_067675662] UDP-2,4-diacetamido-2,4,6-trideoxy-beta-L-altropyranose hydrolase [Tsuneonella dongtanensis]. | 278..284 |
| rs:WP_140467163 | [WP_140467163] sensor histidine kinase [Hymenobacter nivis]. | 253..259 |
| rs:WP_140469455 | [WP_140469455] sensor histidine kinase [Hymenobacter nivis]. | 253..259 |
| rs:WP_237206959 | [WP_237206959] GNAT family N-acetyltransferase [Rothia nasimurium]. | 65..71 |
| rs:WP_160869257 | [WP_160869257] ABC transporter ATP-binding protein [Pantoea sp. Taur]. | 195..201 |
| rs:WP_213736772 | [WP_213736772] hypothetical protein [Bradyrhizobium sp. dw_411]. | 342..348 |
| rs:WP_215766749 | [WP_215766749] hypothetical protein [Gluconobacter cerinus]. | 213..219 |
| rs:WP_073693475 | [WP_073693475] hypothetical protein [Mycobacterium sp. ST-F2]. | 160..166 |
| rs:WP_074116774 | [WP_074116774] hypothetical protein [Bradyrhizobium sp. AS23.2]. | 341..347 |
| rs:WP_217005909 | [WP_217005909] hypothetical protein [Brevibacterium luteolum]. | 148..154 |
| rs:WP_114625607 | [WP_114625607] SDR family oxidoreductase [Streptomyces corynorhini]. | 50..56 |
| rs:WP_084809020 | [WP_084809020] hypothetical protein [Bradyrhizobium sp. NAS80.1]. | 341..347 |
| rs:WP_134319600 | [WP_134319600] glycosyltransferase family 87 protein [Burkholderia cepacia]. | 385..391 |
| rs:WP_136470953 | [WP_136470953] glycosyltransferase family 87 protein [Burkholderia sp. LS-044]. | 385..391 |
| rs:WP_193102632 | [WP_193102632] glycosyltransferase family 87 protein [Burkholderia sp. Z1]. | 385..391 |
| rs:WP_200059960 | [WP_200059960] glycosyltransferase family 87 protein [Burkholderia cepacia]. | 385..391 |
| rs:WP_200094707 | [WP_200094707] glycosyltransferase family 87 protein [Burkholderia cenocepacia]. | 385..391 |
| rs:WP_200160329 | [WP_200160329] glycosyltransferase family 87 protein [Burkholderia orbicola]. | 385..391 |
| rs:WP_202755668 | [WP_202755668] glycosyltransferase family 87 protein [Burkholderia cenocepacia]. | 385..391 |
| rs:WP_205675222 | [WP_205675222] glycosyltransferase family 87 protein [Burkholderia cenocepacia]. | 385..391 |
| rs:WP_205788167 | [WP_205788167] glycosyltransferase family 87 protein [Burkholderia sp. Ac-20344]. | 385..391 |
| rs:WP_205823179 | [WP_205823179] glycosyltransferase family 87 protein [Burkholderia sp. Ac-20349]. | 385..391 |
| rs:WP_206123349 | [WP_206123349] glycosyltransferase family 87 protein [Burkholderia sp. Ac-20392]. | 385..391 |
| rs:WP_206133971 | [WP_206133971] glycosyltransferase family 87 protein [Burkholderia sp. Se-20378]. | 385..391 |
| rs:WP_206139817 | [WP_206139817] glycosyltransferase family 87 protein [Burkholderia sp. Se-20373]. | 385..391 |
| rs:WP_206145360 | [WP_206145360] glycosyltransferase family 87 protein [Burkholderia sp. Ac-20384]. | 385..391 |
| rs:WP_212043214 | [WP_212043214] glycosyltransferase family 87 protein [Burkholderia cenocepacia]. | 385..391 |
| rs:WP_212050289 | [WP_212050289] glycosyltransferase family 87 protein [Burkholderia cenocepacia]. | 385..391 |

| | | |
|---|---|---|
| rs:WP_212090517 | [WP_212090517] glycosyltransferase family 87 protein [Burkholderia cenocepacia]. | 385..391 |
| rs:WP_212108750 | [WP_212108750] glycosyltransferase family 87 protein [Burkholderia cenocepacia]. | 385..391 |
| rs:WP_212146117 | [WP_212146117] glycosyltransferase family 87 protein [Burkholderia cenocepacia]. | 385..391 |
| rs:WP_212175493 | [WP_212175493] glycosyltransferase family 87 protein [Burkholderia cenocepacia]. | 385..391 |
| rs:WP_212195288 | [WP_212195288] glycosyltransferase family 87 protein [Burkholderia cenocepacia]. | 385..391 |
| rs:WP_222155008 | [WP_222155008] glycosyltransferase family 87 protein [Burkholderia cepacia]. | 385..391 |
| rs:WP_222212906 | [WP_222212906] glycosyltransferase family 87 protein [Burkholderia cepacia]. | 385..391 |
| rs:WP_222906020 | [WP_222906020] glycosyltransferase family 87 protein [Burkholderia arboris]. | 385..391 |
| rs:WP_226111072 | [WP_226111072] glycosyltransferase family 87 protein [Burkholderia cenocepacia]. | 385..391 |
| rs:WP_226113036 | [WP_226113036] glycosyltransferase family 87 protein [Burkholderia cepacia]. | 385..391 |
| rs:WP_226120399 | [WP_226120399] glycosyltransferase family 87 protein [Burkholderia contaminans]. | 385..391 |
| rs:WP_226135321 | [WP_226135321] glycosyltransferase family 87 protein [Burkholderia cenocepacia]. | 385..391 |
| rs:WP_226151142 | [WP_226151142] glycosyltransferase family 87 protein [Burkholderia cepacia]. | 385..391 |
| rs:WP_226155967 | [WP_226155967] glycosyltransferase family 87 protein [Burkholderia cepacia]. | 385..391 |
| rs:WP_226160146 | [WP_226160146] glycosyltransferase family 87 protein [Burkholderia cepacia]. | 385..391 |
| rs:WP_226181263 | [WP_226181263] glycosyltransferase family 87 protein [Burkholderia cepacia]. | 385..391 |
| rs:WP_226187040 | [WP_226187040] glycosyltransferase family 87 protein [Burkholderia cepacia]. | 385..391 |
| rs:WP_226196041 | [WP_226196041] glycosyltransferase family 87 protein [Burkholderia arboris]. | 385..391 |
| rs:WP_226204823 | [WP_226204823] glycosyltransferase family 87 protein [Burkholderia sp. AU38729]. | 385..391 |
| rs:WP_226215338 | [WP_226215338] glycosyltransferase family 87 protein [Burkholderia sp. AU30198]. | 385..391 |
| rs:WP_226223598 | [WP_226223598] glycosyltransferase family 87 protein [Burkholderia contaminans]. | 385..391 |
| rs:WP_226255407 | [WP_226255407] glycosyltransferase family 87 protein [Burkholderia cepacia]. | 385..391 |
| rs:WP_226265449 | [WP_226265449] glycosyltransferase family 87 protein [Burkholderia arboris]. | 385..391 |
| rs:WP_226274453 | [WP_226274453] glycosyltransferase family 87 protein [Burkholderia sp. AU31624]. | 385..391 |
| rs:WP_226287602 | [WP_226287602] glycosyltransferase family 87 protein [Burkholderia cepacia]. | 385..391 |
| rs:WP_233345151 | [WP_233345151] glycosyltransferase family 87 protein [Burkholderia cepacia]. | 385..391 |
| rs:WP_234621812 | [WP_234621812] glycosyltransferase family 87 protein [Burkholderia cenocepacia]. | 385..391 |
| rs:WP_241293653 | [WP_241293653] glycosyltransferase family 87 protein [Burkholderia stabilis]. | 385..391 |
| rs:WP_241303518 | [WP_241303518] glycosyltransferase family 87 protein [Burkholderia stabilis]. | 385..391 |
| rs:WP_241331627 | [WP_241331627] glycosyltransferase family 87 protein [Burkholderia cenocepacia]. | 385..391 |
| rs:WP_249455512 | [WP_249455512] glycosyltransferase family 87 protein [Burkholderia cepacia]. | 385..391 |
| rs:WP_256087782 | [WP_256087782] glycosyltransferase family 87 protein [Burkholderia arboris]. | 385..391 |
| rs:WP_187054726 | [WP_187054726] response regulator transcription factor [Variovorax sp. PAMC28562]. | 27..33 |
| rs:WP_184206835 | [WP_184206835] iron ABC transporter permease [Prosthecobacter dejongeii]. | 308..314 |
| rs:WP_276363158 | [WP_276363158] hypothetical protein [Amycolatopsis sp. QT-25]. | 28..34 |
| rs:WP_153559282 | [WP_153559282] serine/threonine-protein kinase [Roseimaritima sediminicola]. | 215..221 |
| rs:WP_015863190 | [WP_015863190] MFS transporter [Solidesulfovibrio magneticus]. | 298..304 |
| rs:WP_146253116 | [WP_146253116] glycosyltransferase family 87 protein [Burkholderia cepacia]. | 385..391 |
| rs:WP_153490832 | [WP_153490832] glycosyltransferase family 87 protein [Burkholderia cepacia]. | 385..391 |
| rs:WP_008412597 | [WP_008412597] MULTISPECIES: DeoR/GlpR family DNA-binding transcription regulator [Streptomyces]. | 112..118 |
| rs:WP_018470977 | [WP_018470977] MULTISPECIES: DeoR/GlpR family DNA-binding transcription regulator [Streptomyces]. | 112..118 |
| rs:WP_023420790 | [WP_023420790] MULTISPECIES: DeoR/GlpR family DNA-binding transcription regulator [unclassified Streptomyces]. | 112..118 |
| rs:WP_030309256 | [WP_030309256] DeoR/GlpR family DNA-binding transcription regulator [Streptomyces albidoflavus]. | 112..118 |
| rs:WP_114181826 | [WP_114181826] glycosyltransferase family 87 protein [Burkholderia pyrrocinia]. | 385..391 |
| rs:WP_119338544 | [WP_119338544] MULTISPECIES: glycosyltransferase family 87 protein [Burkholderia]. | 385..391 |
| rs:WP_122172918 | [WP_122172918] glycosyltransferase family 87 protein [Burkholderia stabilis]. | 385..391 |
| rs:WP_122477930 | [WP_122477930] MULTISPECIES: glycosyltransferase family 87 protein [Burkholderia]. | 385..391 |
| rs:WP_124462535 | [WP_124462535] glycosyltransferase family 87 protein [Burkholderia cenocepacia]. | 385..391 |
| rs:WP_124467614 | [WP_124467614] glycosyltransferase family 87 protein [Burkholderia cepacia]. | 385..391 |
| rs:WP_124478017 | [WP_124478017] glycosyltransferase family 87 protein [Burkholderia cenocepacia]. | 385..391 |
| rs:WP_124492880 | [WP_124492880] glycosyltransferase family 87 protein [Burkholderia contaminans]. | 385..391 |
| rs:WP_124535343 | [WP_124535343] glycosyltransferase family 87 protein [Burkholderia cepacia]. | 385..391 |
| rs:WP_124544458 | [WP_124544458] glycosyltransferase family 87 protein [Burkholderia cepacia]. | 385..391 |
| rs:WP_124548528 | [WP_124548528] glycosyltransferase family 87 protein [Burkholderia cenocepacia]. | 385..391 |
| rs:WP_124581133 | [WP_124581133] glycosyltransferase family 87 protein [Burkholderia contaminans]. | 385..391 |
| rs:WP_124620834 | [WP_124620834] glycosyltransferase family 87 protein [Burkholderia cenocepacia]. | 385..391 |
| rs:WP_124627496 | [WP_124627496] glycosyltransferase family 87 protein [Burkholderia cepacia]. | 385..391 |
| rs:WP_124647672 | [WP_124647672] glycosyltransferase family 87 protein [Burkholderia contaminans]. | 385..391 |
| rs:WP_124672403 | [WP_124672403] glycosyltransferase family 87 protein [Burkholderia cepacia]. | 385..391 |
| rs:WP_124676907 | [WP_124676907] glycosyltransferase family 87 protein [Burkholderia cenocepacia]. | 385..391 |
| rs:WP_124700984 | [WP_124700984] glycosyltransferase family 87 protein [Burkholderia cenocepacia]. | 385..391 |
| rs:WP_124815091 | [WP_124815091] glycosyltransferase family 87 protein [Burkholderia cepacia]. | 385..391 |
| rs:WP_124839205 | [WP_124839205] glycosyltransferase family 87 protein [Burkholderia cepacia]. | 385..391 |
| rs:WP_124918895 | [WP_124918895] glycosyltransferase family 87 protein [Burkholderia cepacia]. | 385..391 |
| rs:WP_128463727 | [WP_128463727] DeoR/GlpR family DNA-binding transcription regulator [Streptomyces albidoflavus]. | 112..118 |
| rs:WP_128842453 | [WP_128842453] glycosyltransferase family 87 protein [Burkholderia catarinensis]. | 385..391 |
| rs:WP_129518238 | [WP_129518238] glycosyltransferase family 87 protein [Burkholderia stabilis]. | 385..391 |
| rs:WP_129805189 | [WP_129805189] MULTISPECIES: DeoR/GlpR family DNA-binding transcription regulator [Streptomyces]. | 112..118 |
| rs:WP_129846502 | [WP_129846502] MULTISPECIES: DeoR/GlpR family DNA-binding transcription regulator [Streptomyces]. | 112..118 |
| rs:WP_129863691 | [WP_129863691] DeoR/GlpR family DNA-binding transcription regulator [Streptomyces albidoflavus]. | 112..118 |
| rs:WP_165580040 | [WP_165580040] glycosyltransferase family 87 protein [Burkholderia cepacia]. | 385..391 |
| rs:WP_097803134 | [WP_097803134] hypothetical protein [Pelagimonas varians]. | 159..165 |

| | | |
|---|---|---|
| rs:WP_166895899 | [WP_166895899] MULTISPECIES: glycosyltransferase family 87 protein [unclassified Burkholderia]. | 385..391 |
| rs:WP_174929319 | [WP_174929319] glycosyltransferase family 87 protein [Burkholderia lata]. | 385..391 |
| rs:WP_174933114 | [WP_174933114] glycosyltransferase family 87 protein [Burkholderia lata]. | 385..391 |
| rs:WP_174938789 | [WP_174938789] glycosyltransferase family 87 protein [Burkholderia lata]. | 385..391 |
| rs:WP_174949358 | [WP_174949358] glycosyltransferase family 87 protein [Burkholderia lata]. | 385..391 |
| rs:WP_174970567 | [WP_174970567] glycosyltransferase family 87 protein [Burkholderia contaminans]. | 385..391 |
| rs:WP_174989107 | [WP_174989107] glycosyltransferase family 87 protein [Burkholderia lata]. | 385..391 |
| rs:WP_174992367 | [WP_174992367] glycosyltransferase family 87 protein [Burkholderia arboris]. | 385..391 |
| rs:WP_175014430 | [WP_175014430] glycosyltransferase family 87 protein [Burkholderia lata]. | 385..391 |
| rs:WP_175017801 | [WP_175017801] glycosyltransferase family 87 protein [Burkholderia contaminans]. | 385..391 |
| rs:WP_175020667 | [WP_175020667] glycosyltransferase family 87 protein [Burkholderia aenigmatica]. | 385..391 |
| rs:WP_175037273 | [WP_175037273] glycosyltransferase family 87 protein [Burkholderia contaminans]. | 385..391 |
| rs:WP_175041179 | [WP_175041179] glycosyltransferase family 87 protein [Burkholderia contaminans]. | 385..391 |
| rs:WP_175042461 | [WP_175042461] glycosyltransferase family 87 protein [Burkholderia lata]. | 385..391 |
| rs:WP_175239676 | [WP_175239676] MULTISPECIES: glycosyltransferase family 87 protein [Burkholderia cepacia complex]. | 385..391 |
| rs:WP_175682022 | [WP_175682022] glycosyltransferase family 87 protein [Burkholderia cenocepacia]. | 385..391 |
| rs:WP_175766569 | [WP_175766569] glycosyltransferase family 87 protein [Burkholderia cenocepacia]. | 385..391 |
| rs:WP_175781917 | [WP_175781917] glycosyltransferase family 87 protein [Burkholderia cenocepacia]. | 385..391 |
| rs:WP_175789135 | [WP_175789135] glycosyltransferase family 87 protein [Burkholderia cenocepacia]. | 385..391 |
| rs:WP_175808148 | [WP_175808148] glycosyltransferase family 87 protein [Burkholderia cenocepacia]. | 385..391 |
| rs:WP_175833188 | [WP_175833188] glycosyltransferase family 87 protein [Burkholderia cenocepacia]. | 385..391 |
| rs:WP_175846049 | [WP_175846049] glycosyltransferase family 87 protein [Burkholderia arboris]. | 385..391 |
| rs:WP_175851250 | [WP_175851250] glycosyltransferase family 87 protein [Burkholderia cepacia]. | 385..391 |
| rs:WP_175853972 | [WP_175853972] glycosyltransferase family 87 protein [Burkholderia cepacia]. | 385..391 |
| rs:WP_175861750 | [WP_175861750] glycosyltransferase family 87 protein [Burkholderia cepacia]. | 385..391 |
| rs:WP_175864073 | [WP_175864073] glycosyltransferase family 87 protein [Burkholderia cepacia]. | 385..391 |
| rs:WP_175890543 | [WP_175890543] glycosyltransferase family 87 protein [Burkholderia cepacia]. | 385..391 |
| rs:WP_175893559 | [WP_175893559] glycosyltransferase family 87 protein [Burkholderia cepacia]. | 385..391 |
| rs:WP_175925577 | [WP_175925577] glycosyltransferase family 87 protein [Burkholderia cepacia]. | 385..391 |
| rs:WP_175930776 | [WP_175930776] MULTISPECIES: glycosyltransferase family 87 protein [unclassified Burkholderia]. | 384..390 |
| rs:WP_175936421 | [WP_175936421] glycosyltransferase family 87 protein [Burkholderia cepacia]. | 385..391 |
| rs:WP_175952423 | [WP_175952423] glycosyltransferase family 87 protein [Burkholderia sp. BCC0405]. | 385..391 |
| rs:WP_175997446 | [WP_175997446] glycosyltransferase family 87 protein [Burkholderia stabilis]. | 385..391 |
| rs:WP_176040729 | [WP_176040729] glycosyltransferase family 87 protein [Burkholderia stabilis]. | 385..391 |
| rs:WP_176115173 | [WP_176115173] glycosyltransferase family 87 protein [Burkholderia cepacia]. | 385..391 |
| rs:WP_185921501 | [WP_185921501] glycosyltransferase family 87 protein [Burkholderia cenocepacia]. | 385..391 |
| rs:WP_187579797 | [WP_187579797] sugar transferase [Nocardioides mesophilus]. | 247..253 |
| rs:WP_189058397 | [WP_189058397] dienelactone hydrolase family protein [Longimycelium tulufanense]. | 58..64 |
| rs:WP_191815268 | [WP_191815268] GNAT family N-acetyltransferase [Cytobacillus stercorigallinarum]. | 65..71 |
| rs:WP_191857250 | [WP_191857250] DeoR/GlpR family DNA-binding transcription regulator [Streptomyces fungicidicus]. | 112..118 |
| rs:WP_232818366 | [WP_232818366] site-specific integrase [Elioraea thermophila]. | 198..204 |
| rs:WP_194164922 | [WP_194164922] MULTISPECIES: DeoR/GlpR family DNA-binding transcription regulator [unclassified Pseudactinotalea]. | 119..125 |
| rs:WP_229255982 | [WP_229255982] sensor histidine kinase [Duganella fentianensis]. | 72..78 |
| rs:WP_272544394 | [WP_272544394] glycosyltransferase family 87 protein [Burkholderia cepacia]. | 385..391 |
| rs:WP_135133807 | [WP_135133807] sensor domain-containing diguanylate cyclase [Blastococcus sp. CT_GayMR16]. | 39..45 |
| rs:WP_065971085 | [WP_065971085] hypothetical protein [Acidiferrobacter thiooxydans]. | 466..472 |
| rs:WP_075355059 | [WP_075355059] MFS transporter [Desulfovibrio sp. DV]. | 298..304 |
| rs:WP_259633500 | [WP_259633500] serine/threonine-protein kinase [Stieleria sedimenti]. | 215..221 |
| rs:WP_311011391 | [WP_311011391] glycosyltransferase family 87 protein [Burkholderia cenocepacia]. | 385..391 |
| rs:XP_009319005 | [XP_009319005] PREDICTED: acyl-CoA-binding domain-containing protein 6 [Pygoscelis adeliae]. | 3..9 |
| rs:XP_011262076 | [XP_011262076] UNC93-like protein isoform X1 [Camponotus floridanus]. | 547..553 |
| rs:XP_028432248 | [XP_028432248] LOW QUALITY PROTEIN: ubiquitin-like modifier-activating enzyme 1 [Perca flavescens]. | 576..582 |
| rs:XP_010017002 | [XP_010017002] PREDICTED: amphoterin-induced protein 3 [Nestor notabilis]. | 86..92 |
| rs:XP_029666472 | [XP_029666472] UNC93-like protein isoform X1 [Formica exsecta]. | 547..553 |
| rs:XP_029947903 | [XP_029947903] ubiquitin-like modifier-activating enzyme 1 [Salarias fasciatus]. | 562..568 |
| rs:XP_033938723 | [XP_033938723] ubiquitin-like modifier-activating enzyme 1 isoform X1 [Pseudochaenichthys georgianus]. | 566..572 |
| rs:XP_033938724 | [XP_033938724] ubiquitin-like modifier-activating enzyme 1 isoform X2 [Pseudochaenichthys georgianus]. | 562..568 |
| rs:XP_033994545 | [XP_033994545] ubiquitin-like modifier-activating enzyme 1 isoform X1 [Trematomus bernacchii]. | 566..572 |
| rs:XP_033994546 | [XP_033994546] ubiquitin-like modifier-activating enzyme 1 isoform X2 [Trematomus bernacchii]. | 562..568 |
| rs:XP_030764762 | [XP_030764762] protein NEDD1-like [Sitophilus oryzae]. | 48..54 |
| rs:XP_032369896 | [XP_032369896] ubiquitin-like modifier-activating enzyme 1 [Etheostoma spectabile]. | 562..568 |
| rs:XP_034074972 | [XP_034074972] ubiquitin-like modifier-activating enzyme 1 [Gymnodraco acuticeps]. | 562..568 |
| rs:XP_019909256 | [XP_019909256] dynein regulatory complex subunit 5 [Esox lucius]. | 41..47 |
| rs:XP_023135555 | [XP_023135555] ubiquitin-like modifier-activating enzyme 1 isoform X1 [Amphiprion ocellaris]. | 565..571 |
| rs:XP_023135556 | [XP_023135556] ubiquitin-like modifier-activating enzyme 1 isoform X2 [Amphiprion ocellaris]. | 562..568 |
| rs:XP_054866444 | [XP_054866444] ubiquitin-like modifier-activating enzyme 1 isoform X3 [Amphiprion ocellaris]. | 565..571 |
| rs:XP_034724688 | [XP_034724688] ubiquitin-like modifier-activating enzyme 1 [Etheostoma cragini]. | 562..568 |
| rs:XP_031157362 | [XP_031157362] ubiquitin-like modifier-activating enzyme 1 [Sander lucioperca]. | 562..568 |
| rs:XP_039653648 | [XP_039653648] ubiquitin-like modifier-activating enzyme 1 [Perca fluviatilis]. | 562..568 |
| rs:XP_039831703 | [XP_039831703] calmodulin-binding transcription activator 3-like isoform X1 [Panicum virgatum]. | 9..15 |
| rs:XP_039831704 | [XP_039831704] calmodulin-binding transcription activator 3-like isoform X2 [Panicum virgatum]. | 9..15 |

| | | |
|---|---|---|
| rs:XP_051804143 | [XP_051804143] ubiquitin-like modifier-activating enzyme 1 [Acanthochromis polyacanthus]. | 562..568 |
| rs:XP_040659493 | [XP_040659493] zinc finger domain-containing protein [Drechmeria coniospora]. | 100..106 |
| rs:XP_042363183 | [XP_042363183] ubiquitin-like modifier-activating enzyme 1 [Plectropomus leopardus]. | 562..568 |
| rs:XP_012187514 | [XP_012187514] hypothetical protein PHSY_001495 [Pseudozyma hubeiensis SY62]. | 30..36 |
| rs:XP_038252403 | [XP_038252403] LOW QUALITY PROTEIN: dynein regulatory complex subunit 5 [Dermochelys coriacea]. | 42..48 |
| rs:XP_047472938 | [XP_047472938] hornerin-like [Penaeus chinensis]. | 16..22 |
| rs:XP_047472939 | [XP_047472939] uncharacterized protein LOC125027851 [Penaeus chinensis]. | 16..22 |
| rs:XP_048867380 | [XP_048867380] decorin [Brienomyrus brachyistius]. | 269..275 |
| rs:XP_050446029 | [XP_050446029] UNC93-like protein [Cataglyphis hispanica]. | 547..553 |
| rs:XP_052966990 | [XP_052966990] beta-lactamase/transpeptidase-like protein [Polychytrium aggregatum]. | 204..210 |
| rs:XP_052966175 | [XP_052966175] pyridoxal-dependent decarboxylase [Polychytrium aggregatum]. | 189..195 |
| rs:XP_016269166 | [XP_016269166] uncharacterized protein PV06_01506 [Exophiala oligosperma]. | 14..20 |
| rs:XP_062484263 | [XP_062484263] sulfiredoxin-1 [Pezoporus occidentalis]. | 16..22 |
| **rs:XP_445695** | **[XP_445695] uncharacterized protein GVI51_D06655 [Nakaseomyces glabratus].** | **51..57** |
| tr:A0A4S2MQN4_9PEZI | [A0A4S2MQN4] SubName: Full=Uncharacterized protein {ECO:0000313\|EMBL:TGZ79483.1}; | 6..12 |
| tr:A0A8H7I9B8_9AGAM | [A0A8H7I9B8] RecName: Full=Beta-xylanase {ECO:0000256\|RuleBase:RU361174}; EC=3.2.1.8 {ECO:0000256\|RuleBase:RU361174}; | 15..21 |
| tr:A0A4S9LCT5_AURPU | [A0A4S9LCT5] SubName: Full=Beta-lactamase/transpeptidase-like protein {ECO:0000313\|EMBL:THY26958.1}; | 11..17 |
| tr:A0A9P4MQD0_9PLEO | [A0A9P4MQD0] RecName: Full=Exonuclease domain-containing protein {ECO:0000259\|SMART:SM00479}; | 261..267 |
| tr:A0A4S8KUZ3_DENBC | [A0A4S8KUZ3] SubName: Full=Uncharacterized protein {ECO:0000313\|EMBL:THU79717.1}; | 170..176 |
| tr:A0A9P9JRN0_9HYPO | [A0A9P9JRN0] RecName: Full=Zn(2)-C6 fungal-type domain-containing protein {ECO:0000259\|PROSITE:PS50048}; | 532..538 |
| tr:A0A6A6XAP8_9PLEO | [A0A6A6XAP8] RecName: Full=Zn(2)-C6 fungal-type domain-containing protein {ECO:0000259\|PROSITE:PS50048}; | 170..176 |
| tr:A0A9P9JBG7_9HYPO | [A0A9P9JBG7] RecName: Full=Zn(2)-C6 fungal-type domain-containing protein {ECO:0000259\|PROSITE:PS50048}; | 532..538 |
| tr:A0A2U3E5K3_PURLI | [A0A2U3E5K3] SubName: Full=Uncharacterized protein {ECO:0000313\|EMBL:PWI69766.1}; | 16..22 |
| tr:A0A182K025_9DIPT | [A0A182K025] RecName: Full=LRRCT domain-containing protein {ECO:0008006\|Google:ProtNLM}; | 107..113 |
| tr:A0A9Q1C917_HOLLE | [A0A9Q1C917] RecName: Full=Phospholipase A2 {ECO:0000256\|RuleBase:RU361236}; EC=3.1.1.4 {ECO:0000256\|RuleBase:RU361236}; | 12..18 |
| tr:A0A5B7DUL7_PORTR | [A0A5B7DUL7] SubName: Full=Uncharacterized protein {ECO:0000313\|EMBL:MPC25391.1}; | 74..80 |
| tr:A0A6A3CK20_HIBSY | [A0A6A3CK20] SubName: Full=Putative inactive receptor kinase {ECO:0000313\|EMBL:KAE8729067.1}; | 37..43 |
| tr:A0A8T0MNW6_PANVG | [A0A8T0MNW6] RecName: Full=CG-1 domain-containing protein {ECO:0000259\|PROSITE:PS51437}; | 9..15 |
| tr:A0A8T0MNX7_PANVG | [A0A8T0MNX7] RecName: Full=CG-1 domain-containing protein {ECO:0000259\|PROSITE:PS51437}; | 9..15 |
| tr:A0A8T0MTM4_PANVG | [A0A8T0MTM4] RecName: Full=CG-1 domain-containing protein {ECO:0000259\|PROSITE:PS51437}; | 9..15 |
| tr:A0A8T0Q0S6_PANVG | [A0A8T0Q0S6] SubName: Full=Uncharacterized protein {ECO:0000313\|EMBL:KAG2567490.1}; | 37..43 |
| tr:A0A8T0MM89_PANVG | [A0A8T0MM89] RecName: Full=CG-1 domain-containing protein {ECO:0000259\|PROSITE:PS51437}; | 9..15 |
| tr:A0A8T0MPI3_PANVG | [A0A8T0MPI3] RecName: Full=CG-1 domain-containing protein {ECO:0000259\|PROSITE:PS51437}; | 9..15 |
| tr:A0A8T0MMP0_PANVG | [A0A8T0MMP0] RecName: Full=CG-1 domain-containing protein {ECO:0000259\|PROSITE:PS51437}; | 9..15 |
| tr:A0A8T0MM98_PANVG | [A0A8T0MM98] RecName: Full=CG-1 domain-containing protein {ECO:0000259\|PROSITE:PS51437}; | 9..15 |
| tr:A0A8T0MQK4_PANVG | [A0A8T0MQK4] RecName: Full=CG-1 domain-containing protein {ECO:0000259\|PROSITE:PS51437}; | 9..15 |
| tr:A0A8T0MSG9_PANVG | [A0A8T0MSG9] RecName: Full=CG-1 domain-containing protein {ECO:0000259\|PROSITE:PS51437}; | 9..15 |
| tr:A0A8T0MSF9_PANVG | [A0A8T0MSF9] RecName: Full=CG-1 domain-containing protein {ECO:0000259\|PROSITE:PS51437}; | 9..15 |
| tr:A0A8T0MMZ1_PANVG | [A0A8T0MMZ1] RecName: Full=CG-1 domain-containing protein {ECO:0000259\|PROSITE:PS51437}; | 9..15 |
| tr:A0A8T0MLR9_PANVG | [A0A8T0MLR9] RecName: Full=CG-1 domain-containing protein {ECO:0000259\|PROSITE:PS51437}; | 9..15 |
| tr:A0A383WN18_TETOB | [A0A383WN18] RecName: Full=Eukaryotic translation initiation factor 3 subunit F {ECO:0000256\|HAMAP-Rule:MF_03005}; Short=eIF3f {ECO:0000256\|HAMAP-Rule:MF_03005}; AltName: Full=eIF-3-epsilon {ECO:0000256\|HAMAP-Rule:MF_03005}; | 237..243 |
| tr:A0A371HW29_MUCPR | [A0A371HW29] SubName: Full=Retrovirus-related Pol polyprotein from transposon 17.6 {ECO:0000313\|EMBL:RDY06982.1}; Flags: Fragment; | 260..266 |
| tr:A0A7J8RIR8_GOSDV | [A0A7J8RIR8] SubName: Full=Uncharacterized protein {ECO:0000313\|EMBL:MBA0613473.1}; | 20..26 |
| tr:A0A7J7Q3E2_9CHLO | [A0A7J7Q3E2] RecName: Full=Eukaryotic translation initiation factor 3 subunit F {ECO:0000256\|HAMAP-Rule:MF_03005}; Short=eIF3f {ECO:0000256\|HAMAP-Rule:MF_03005}; AltName: Full=eIF-3-epsilon {ECO:0000256\|HAMAP-Rule:MF_03005}; | 236..242 |
| tr:A0A8S1IS20_9CHLO | [A0A8S1IS20] RecName: Full=BZIP domain-containing protein {ECO:0008006\|Google:ProtNLM}; | 372..378 |
| tr:A0A218XMU8_PUNGR | [A0A218XMU8] RecName: Full=Dynamin-related protein 4C-like {ECO:0008006\|Google:ProtNLM}; | 31..37 |
| tr:A0A2I0JAW2_PUNGR | [A0A2I0JAW2] RecName: Full=GED domain-containing protein {ECO:0000259\|PROSITE:PS51388}; | 31..37 |
| tr:A0AAE0L8F8_9CHLO | [A0AAE0L8F8] RecName: Full=Histidine kinase domain-containing protein {ECO:0000259\|PROSITE:PS50109}; | 176..182 |
| tr:A0A418FVQ7_APHAT | [A0A418FVQ7] SubName: Full=Uncharacterized protein {ECO:0000313\|EMBL:RHZ38728.1}; | 20..26 |
| tr:A0A8J1ZCM9_9DINO | [A0A8J1ZCM9] SubName: Full=Uncharacterized protein {ECO:0000313\|EMBL:CAD7935914.1}; | 26..32 |
| tr:A0AA48KW12_9ALTE | [A0AA48KW12] SubName: Full=Isoquinoline 1-oxidoreductase subunit beta {ECO:0000313\|EMBL:BDX08075.1}; | 22..28 |
| tr:A0A7V4EG93_9DEIN | [A0A7V4EG93] SubName: Full=Dipeptidyl aminopeptidase {ECO:0000313\|EMBL:HGN85626.1}; | 14..20 |
| tr:A0A1L6JI08_9SPHN | [A0A1L6JI08] RecName: Full=diguanylate cyclase {ECO:0000256\|ARBA:ARBA00012528}; EC=2.7.7.65 {ECO:0000256\|ARBA:ARBA00012528}; | 59..65 |
| tr:A0A2E8WT89_UNCCH | [A0A2E8WT89] SubName: Full=Glutaconyl-CoA decarboxylase subunit beta {ECO:0000313\|EMBL:MBS16778.1}; | 391..397 |
| tr:R6AWY4_9BACT | [R6AWY4] SubName: Full=ABC-2 type transporter {ECO:0000313\|EMBL:CDA45627.1}; | 258..264 |
| tr:A0A0S7XCB6_9BACT | [A0A0S7XCB6] RecName: Full=Glycosyltransferase family 1 protein {ECO:0008006\|Google:ProtNLM}; | 109..115 |
| tr:A0A4D7DP15_9SPHN | [A0A4D7DP15] RecName: Full=diguanylate cyclase {ECO:0000256\|ARBA:ARBA00012528}; EC=2.7.7.65 {ECO:0000256\|ARBA:ARBA00012528}; | 59..65 |
| tr:A0A2W6E0M6_9PSEU | [A0A2W6E0M6] SubName: Full=DUF4192 domain-containing protein {ECO:0000313\|EMBL:PZS38150.1}; Flags: Fragment; | 28..34 |
| tr:A0A960AMV1_9ACTN | [A0A960AMV1] SubName: Full=RecQ family ATP-dependent DNA helicase {ECO:0000313\|EMBL:MCB0900665.1}; | 203..209 |
| tr:A0A933LCT7_9ACTN | [A0A933LCT7] SubName: Full=Uncharacterized protein {ECO:0000313\|EMBL:MBI4933521.1}; | 28..34 |
| tr:A0A951KGX7_9ACTN | [A0A951KGX7] SubName: Full=Alpha/beta hydrolase {ECO:0000313\|EMBL:MBW3626672.1}; Flags: Fragment; | 206..212 |
| tr:A0A7X7D618_9ACTN | [A0A7X7D618] SubName: Full=Aminotransferase class I/II-fold pyridoxal phosphate-dependent enzyme {ECO:0000313\|EMBL:NLD77115.1}; | 195..201 |

| | | |
|---|---|---|
| tr:A0A3D5DVE1_UNCAC | [A0A3D5DVE1] SubName: Full=DUF4192 domain-containing protein {ECO:0000313\|EMBL:HCU95765.1}; | 32..38 |
| tr:A0A7V9J0R7_UNCAC | [A0A7V9J0R7] RecName: Full=D-alanine--D-alanine ligase {ECO:0000256\|HAMAP-Rule:MF_00047}; EC=6.3.2.4 {ECO:0000256\|HAMAP-Rule:MF_00047}; AltName: Full=D-Ala-D-Ala ligase {ECO:0000256\|HAMAP-Rule:MF_00047}; AltName: Full=D-alanylalanine synthetase {ECO:0000256\|HAMAP-Rule:MF_00047}; | 105..111 |
| tr:Q92PX7_RHIME | [Q92PX7] SubName: Full=MFS permease {ECO:0000313\|EMBL:CAC46172.1}; | 109..115 |
| tr:A0A430DI86_9SPHN | [A0A430DI86] RecName: Full=diguanylate cyclase {ECO:0000256\|ARBA:ARBA00012528}; EC=2.7.7.65 {ECO:0000256\|ARBA:ARBA00012528}; | 59..65 |
| tr:A0A0S7Z9A8_9SPIR | [A0A0S7Z9A8] SubName: Full=Creatininase {ECO:0000313\|EMBL:KPJ83704.1}; | 172..178 |
| tr:A0A915UEF9_9BACT | [A0A915UEF9] SubName: Full=ABC transporter substrate-binding protein {ECO:0000313\|EMBL:BCS34722.1}; | 66..72 |
| tr:A0A5N8Z7H9_9CHLR | [A0A5N8Z7H9] SubName: Full=Sodium ion-translocating decarboxylase subunit beta {ECO:0000313\|EMBL:MQF83443.1}; | 391..397 |
| tr:A0A5N9G436_9CHLR | [A0A5N9G436] RecName: Full=DUF1700 domain-containing protein {ECO:0008006\|Google:ProtNLM}; | 83..89 |
| tr:A0A0K2W2B2_MESPL | [A0A0K2W2B2] SubName: Full=Uncharacterized protein {ECO:0000313\|EMBL:CDX59948.1}; | 33..39 |
| tr:A0A7V8KJN7_9HYPH | [A0A7V8KJN7] SubName: Full=MFS transporter {ECO:0000313\|EMBL:KAF5884840.1}; | 169..175 |
| tr:A0A356MM88_9FIRM | [A0A356MM88] RecName: Full=RimM N-terminal domain-containing protein {ECO:0000259\|Pfam:PF01782}; | 127..133 |
| tr:A0A935IKW8_9GAMM | [A0A935IKW8] RecName: Full=Ribosomal RNA small subunit methyltransferase G {ECO:0000256\|HAMAP-Rule:MF_00074}; EC=2.1.1.170 {ECO:0000256\|HAMAP-Rule:MF_00074}; AltName: Full=16S rRNA 7-methylguanosine methyltransferase {ECO:0000256\|HAMAP-Rule:MF_00074}; Short=16S rRNA m7G methyltransferase {ECO:0000256\|HAMAP-Rule:MF_00074}; | 67..73 |
| tr:A0A661GHP6_9GAMM | [A0A661GHP6] SubName: Full=Glutamate--cysteine ligase {ECO:0000313\|EMBL:RLA35917.1}; EC=6.3.2.2 {ECO:0000313\|EMBL:RLA35917.1}; | 6..12 |
| tr:A0A538TL55_UNCEI | [A0A538TL55] SubName: Full=Glycosyltransferase family 4 protein {ECO:0000313\|EMBL:TMQ64350.1}; | 214..220 |
| tr:A0A538S848_UNCEI | [A0A538S848] SubName: Full=Glycosyltransferase family 4 protein {ECO:0000313\|EMBL:TMQ47562.1}; | 214..220 |
| tr:A0A538SSM8_UNCEI | [A0A538SSM8] SubName: Full=Glycosyltransferase family 4 protein {ECO:0000313\|EMBL:TMQ54389.1}; | 214..220 |
| tr:A0A538T7U7_UNCEI | [A0A538T7U7] SubName: Full=Glycosyltransferase family 4 protein {ECO:0000313\|EMBL:TMQ59706.1}; | 214..220 |
| tr:A0A538TBF0_UNCEI | [A0A538TBF0] RecName: Full=Glycosyltransferase subfamily 4-like N-terminal domain-containing protein {ECO:0000259\|Pfam:PF13439}; Flags: Fragment; | 186..192 |
| tr:H0G0L8_RHIML | [H0G0L8] SubName: Full=Major facilitator superfamily protein {ECO:0000313\|EMBL:EHK77206.1}; | 169..175 |
| tr:A0A9Q2FND5_GLUJA | [A0A9Q2FND5] RecName: Full=Uracil-DNA glycosylase-like domain-containing protein {ECO:0008006\|Google:ProtNLM}; | 186..192 |
| tr:A0A6C0UBX5_9GAMM | [A0A6C0UBX5] RecName: Full=Ribosomal RNA small subunit methyltransferase G {ECO:0000256\|HAMAP-Rule:MF_00074}; EC=2.1.1.170 {ECO:0000256\|HAMAP-Rule:MF_00074}; AltName: Full=16S rRNA 7-methylguanosine methyltransferase {ECO:0000256\|HAMAP-Rule:MF_00074}; Short=16S rRNA m7G methyltransferase {ECO:0000256\|HAMAP-Rule:MF_00074}; | 58..64 |
| tr:A0A2H5WBM5_UNCXX | [A0A2H5WBM5] SubName: Full=2-methylcitrate dehydratase {ECO:0000313\|EMBL:GBC86369.1}; EC=4.2.1.79 {ECO:0000313\|EMBL:GBC86369.1}; | 179..185 |
| tr:A0A3N5QQP9_9BACT | [A0A3N5QQP9] RecName: Full=1,4-dihydroxy-2-naphthoate octaprenyltransferase {ECO:0000256\|NCBIfam:TIGR00751}; EC=2.5.1.74 {ECO:0000256\|NCBIfam:TIGR00751}; | 233..239 |
| tr:A0A2G4I6S0_9BACT | [A0A2G4I6S0] RecName: Full=Pyridoxine 5'-phosphate synthase {ECO:0000256\|HAMAP-Rule:MF_00279, ECO:0000256\|NCBIfam:TIGR00559}; Short=PNP synthase {ECO:0000256\|HAMAP-Rule:MF_00279}; EC=2.6.99.2 {ECO:0000256\|HAMAP-Rule:MF_00279, ECO:0000256\|NCBIfam:TIGR00559}; | 207..213 |
| tr:A0A9E4FVZ2_9CHLR | [A0A9E4FVZ2] RecName: Full=DUF2157 domain-containing protein {ECO:0008006\|Google:ProtNLM}; | 83..89 |
| tr:A0A3C0YUL8_9CHLR | [A0A3C0YUL8] RecName: Full=DUF1700 domain-containing protein {ECO:0008006\|Google:ProtNLM}; | 83..89 |
| tr:A0A920M5T2_9CHLR | [A0A920M5T2] SubName: Full=Glutaconyl-CoA decarboxylase subunit beta {ECO:0000313\|EMBL:GIS30779.1}; | 391..397 |
| tr:A0A5M8SQC0_9BACT | [A0A5M8SQC0] RecName: Full=Ysc84 actin-binding domain-containing protein {ECO:0000259\|Pfam:PF04366}; | 59..65 |
| tr:A0A291GSE1_9MICO | [A0A291GSE1] SubName: Full=Na+/H+ antiporter subunit A {ECO:0000313\|EMBL:ATG53128.1}; | 486..492 |
| tr:A0A9D2PXR7_9MICO | [A0A9D2PXR7] SubName: Full=DUF4040 domain-containing protein {ECO:0000313\|EMBL:HJC69288.1}; | 317..323 |
| tr:A0A7J4VQA3_9HYPH | [A0A7J4VQA3] SubName: Full=MFS transporter {ECO:0000313\|EMBL:KAA0697161.1}; | 212..218 |
| tr:A0A965P4P4_9CAUL | [A0A965P4P4] RecName: Full=Secreted protein {ECO:0008006\|Google:ProtNLM}; | 61..67 |
| tr:A0A223VWK6_9PSED | [A0A223VWK6] SubName: Full=Uncharacterized protein {ECO:0000313\|EMBL:ASV39725.1}; | 38..44 |
| tr:A0A430BYR9_9SPHN | [A0A430BYR9] RecName: Full=diguanylate cyclase {ECO:0000256\|ARBA:ARBA00012528}; EC=2.7.7.65 {ECO:0000256\|ARBA:ARBA00012528}; | 59..65 |
| tr:A0A8I2GSE2_RHILV | [A0A8I2GSE2] SubName: Full=MFS transporter {ECO:0000313\|EMBL:NKM44544.1}; | 195..201 |
| tr:A0A1F4EVE7_9PROT | [A0A1F4EVE7] RecName: Full=Ribosomal RNA small subunit methyltransferase G {ECO:0000256\|HAMAP-Rule:MF_00074}; EC=2.1.1.170 {ECO:0000256\|HAMAP-Rule:MF_00074}; AltName: Full=16S rRNA 7-methylguanosine methyltransferase {ECO:0000256\|HAMAP-Rule:MF_00074}; Short=16S rRNA m7G methyltransferase {ECO:0000256\|HAMAP-Rule:MF_00074}; | 60..66 |
| tr:A0A919WD43_9ACTN | [A0A919WD43] RecName: Full=GPP34 family phosphoprotein {ECO:0008006\|Google:ProtNLM}; | 211..217 |
| tr:A0A5Q2VXD3_9CELL | [A0A5Q2VXD3] SubName: Full=DeoR family transcriptional regulator {ECO:0000313\|EMBL:QGH68151.1}; | 170..176 |
| tr:A0A258V8F6_9SPHN | [A0A258V8F6] RecName: Full=Peptidase S9 prolyl oligopeptidase catalytic domain-containing protein {ECO:0000259\|Pfam:PF00326}; | 3..9 |
| tr:A0A9D6E9U7_UNCGE | [A0A9D6E9U7] RecName: Full=tRNA dimethylallyltransferase {ECO:0000256\|HAMAP-Rule:MF_00185}; EC=2.5.1.75 {ECO:0000256\|HAMAP-Rule:MF_00185}; AltName: Full=Dimethylallyl diphosphate:tRNA dimethylallyltransferase {ECO:0000256\|HAMAP-Rule:MF_00185}; Short=DMAPP:tRNA dimethylallyltransferase {ECO:0000256\|HAMAP-Rule:MF_00185}; Short=DMATase {ECO:0000256\|HAMAP-Rule:MF_00185}; AltName: Full=Isopentenyl-diphosphate:tRNA isopentenyltransferase {ECO:0000256\|HAMAP-Rule:MF_00185}; Short=IPP transferase {ECO:0000256\|HAMAP-Rule:MF_00185}; Short=IPPT {ECO:0000256\|HAMAP-Rule:MF_00185}; Short=IPTase {ECO:0000256\|HAMAP-Rule:MF_00185}; | 49..55 |
| tr:A0A935QC65_UNCXX | [A0A935QC65] SubName: Full=Glycosyltransferase {ECO:0000313\|EMBL:MBK7702965.1}; | 108..114 |
| tr:A0A496UTY6_UNCXX | [A0A496UTY6] RecName: Full=Glycosyltransferase family 1 protein {ECO:0008006\|Google:ProtNLM}; | 170..176 |
| tr:A0A948EYX4_UNCXX | [A0A948EYX4] SubName: Full=Glycosyltransferase family 4 protein {ECO:0000313\|EMBL:MBU1675327.1}; | 108..114 |
| tr:A0A960Y538_UNCXX | [A0A960Y538] SubName: Full=Glycosyltransferase {ECO:0000313\|EMBL:MCB1149945.1}; | 108..114 |
| tr:A0A353LXQ2_UNCFI | [A0A353LXQ2] RecName: Full=Peptidase S9 prolyl oligopeptidase catalytic domain-containing protein {ECO:0000259\|Pfam:PF00326}; | 39..45 |
| tr:A0A371NZH0_9ACTN | [A0A371NZH0] SubName: Full=Sugar ABC transporter ATP-binding protein {ECO:0000313\|EMBL:REK69074.1}; | 86..92 |
| tr:A0A3D0GLT1_9GAMM | [A0A3D0GLT1] SubName: Full=ABC transporter permease {ECO:0000313\|EMBL:HBZ49443.1}; | 60..66 |

| | | |
|---|---|---|
| tr:A0A9D6FJ47_UNCAI | [A0A9D6FJ47] SubName: Full=TraM recognition domain-containing protein {ECO:0000313\|EMBL:MBI2839648.1}; | 33..39 |
| tr:A0A9E2ZXV4_9ACTN | [A0A9E2ZXV4] SubName: Full=DUF4192 domain-containing protein {ECO:0000313\|EMBL:MBV9095173.1}; | 30..36 |
| tr:A0A954RNE4_9BACT | [A0A954RNE4] RecName: Full=Dystroglycan-type cadherin-like domain-containing protein {ECO:0008006\|Google:ProtNLM}; Flags: Fragment; | 303..309 |
| tr:A0A968HTJ6_9CYAN | [A0A968HTJ6] SubName: Full=Aminotransferase class V-fold PLP-dependent enzyme {ECO:0000313\|EMBL:NJK35816.1}; | 292..298 |
| tr:A0A952EL69_9FLAO | [A0A952EL69] RecName: Full=site-specific DNA-methyltransferase (adenine-specific) {ECO:0000256\|ARBA:ARBA00011900}; EC=2.1.1.72 {ECO:0000256\|ARBA:ARBA00011900}; | 738..744 |
| tr:A0A2U2DLD0_9HYPH | [A0A2U2DLD0] RecName: Full=HTH tetR-type domain-containing protein {ECO:0000259\|PROSITE:PS50977}; | 21..27 |
| tr:A0A928T8Q0_9BACT | [A0A928T8Q0] SubName: Full=Iron ABC transporter permease {ECO:0000313\|EMBL:MBE7496227.1}; | 310..316 |
| tr:A0A933B8W6_UNCLA | [A0A933B8W6] SubName: Full=Glycosyltransferase family 4 protein {ECO:0000313\|EMBL:MBI4364611.1}; | 107..113 |
| tr:A0A7X8FPS9_9FIRM | [A0A7X8FPS9] SubName: Full=Prephenate dehydrogenase/arogenate dehydrogenase family protein {ECO:0000313\|EMBL:NLK57776.1}; | 63..69 |
| tr:A0A3B9YKV5_UNCEL | [A0A3B9YKV5] RecName: Full=AAA domain-containing protein {ECO:0008006\|Google:ProtNLM}; | 332..338 |
| tr:A0A949X662_9PROT | [A0A949X662] RecName: Full=Ribosomal RNA small subunit methyltransferase G {ECO:0000256\|HAMAP-Rule:MF_00074}; EC=2.1.1.170 {ECO:0000256\|HAMAP-Rule:MF_00074}; AltName: Full=16S rRNA 7-methylguanosine methyltransferase {ECO:0000256\|HAMAP-Rule:MF_00074}; Short=16S rRNA m7G methyltransferase {ECO:0000256\|HAMAP-Rule:MF_00074}; | 60..66 |
| tr:A0A0C2ZDK9_9BACT | [A0A0C2ZDK9] RecName: Full=TIGR00341 family protein {ECO:0008006\|Google:ProtNLM}; | 32..38 |
| tr:A0AA43JFF7_9PROT | [A0AA43JFF7] RecName: Full=Ribosomal RNA small subunit methyltransferase G {ECO:0000256\|HAMAP-Rule:MF_00074}; EC=2.1.1.170 {ECO:0000256\|HAMAP-Rule:MF_00074}; AltName: Full=16S rRNA 7-methylguanosine methyltransferase {ECO:0000256\|HAMAP-Rule:MF_00074}; Short=16S rRNA m7G methyltransferase {ECO:0000256\|HAMAP-Rule:MF_00074}; | 60..66 |
| tr:A0A934GBH1_9PROT | [A0A934GBH1] RecName: Full=Ribosomal RNA small subunit methyltransferase G {ECO:0000256\|HAMAP-Rule:MF_00074}; EC=2.1.1.170 {ECO:0000256\|HAMAP-Rule:MF_00074}; AltName: Full=16S rRNA 7-methylguanosine methyltransferase {ECO:0000256\|HAMAP-Rule:MF_00074}; Short=16S rRNA m7G methyltransferase {ECO:0000256\|HAMAP-Rule:MF_00074}; | 60..66 |
| tr:A0A9D6J0G4_9PROT | [A0A9D6J0G4] RecName: Full=Ribosomal RNA small subunit methyltransferase G {ECO:0000256\|HAMAP-Rule:MF_00074}; EC=2.1.1.170 {ECO:0000256\|HAMAP-Rule:MF_00074}; AltName: Full=16S rRNA 7-methylguanosine methyltransferase {ECO:0000256\|HAMAP-Rule:MF_00074}; Short=16S rRNA m7G methyltransferase {ECO:0000256\|HAMAP-Rule:MF_00074}; | 71..77 |
| tr:A0A965VRZ3_9PROT | [A0A965VRZ3] RecName: Full=Type II secretion system protein K {ECO:0000256\|PIRNR:PIRNR002786}; | 222..228 |
| tr:A0A257QCR8_9GAMM | [A0A257QCR8] SubName: Full=Muropeptide transporter AmpG {ECO:0000313\|EMBL:OYV46548.1}; | 314..320 |
| tr:A0A3D3LCA5_9BACT | [A0A3D3LCA5] SubName: Full=ABC transporter permease {ECO:0000313\|EMBL:HCN28513.1}; | 308..314 |
| tr:A0A7X8YZ14_UNCPL | [A0A7X8YZ14] RecName: Full=Bacteriophage Mu GpT domain-containing protein {ECO:0000259\|Pfam:PF10124}; Flags: Fragment; | 545..551 |
| tr:A0A2E0LKK5_9GAMM | [A0A2E0LKK5] RecName: Full=Ribosomal RNA small subunit methyltransferase G {ECO:0000256\|HAMAP-Rule:MF_00074}; EC=2.1.1.170 {ECO:0000256\|HAMAP-Rule:MF_00074}; AltName: Full=16S rRNA 7-methylguanosine methyltransferase {ECO:0000256\|HAMAP-Rule:MF_00074}; Short=16S rRNA m7G methyltransferase {ECO:0000256\|HAMAP-Rule:MF_00074}; | 55..61 |
| tr:A0A0D7PGP0_9BRAD | [A0A0D7PGP0] RecName: Full=Polysaccharide biosynthesis protein {ECO:0008006\|Google:ProtNLM}; | 340..346 |
| tr:A0A535HVV5_UNCCH | [A0A535HVV5] RecName: Full=Cobyric acid synthase {ECO:0000256\|HAMAP-Rule:MF_00028}; | 216..222 |
| tr:A0A6I1K948_9BACT | [A0A6I1K948] RecName: Full=Pyridoxine 5'-phosphate synthase {ECO:0000256\|HAMAP-Rule:MF_00279, ECO:0000256\|NCBIfam:TIGR00559}; Short=PNP synthase {ECO:0000256\|HAMAP-Rule:MF_00279}; EC=2.6.99.2 {ECO:0000256\|HAMAP-Rule:MF_00279, ECO:0000256\|NCBIfam:TIGR00559}; | 207..213 |
| tr:A0A972N5U8_UNCCH | [A0A972N5U8] RecName: Full=Chorismate dehydratase {ECO:0000256\|HAMAP-Rule:MF_00995}; EC=4.2.1.151 {ECO:0000256\|HAMAP-Rule:MF_00995}; AltName: Full=Menaquinone biosynthetic enzyme MqnA {ECO:0000256\|HAMAP-Rule:MF_00995}; | 65..71 |
| tr:A0A522J557_9GAMM | [A0A522J557] SubName: Full=Monovalent cation/H+ antiporter subunit D {ECO:0000313\|EMBL:TBR72774.1}; | 92..98 |
| tr:A0A962HV08_9GAMM | [A0A962HV08] RecName: Full=Ribosomal RNA small subunit methyltransferase G {ECO:0000256\|HAMAP-Rule:MF_00074}; EC=2.1.1.170 {ECO:0000256\|HAMAP-Rule:MF_00074}; AltName: Full=16S rRNA 7-methylguanosine methyltransferase {ECO:0000256\|HAMAP-Rule:MF_00074}; Short=16S rRNA m7G methyltransferase {ECO:0000256\|HAMAP-Rule:MF_00074}; | 67..73 |
| tr:A0A2N8NNC9_STREU | [A0A2N8NNC9] SubName: Full=Cell division protein FtsW {ECO:0000313\|EMBL:PNE30280.1}; | 65..71 |
| tr:A0A7W8B7L1_STREU | [A0A7W8B7L1] SubName: Full=Cell division protein FtsW (Lipid II flippase) {ECO:0000313\|EMBL:MBB5118269.1}; | 70..76 |
| tr:A0A259BAZ5_9GAMM | [A0A259BAZ5] SubName: Full=Muropeptide transporter AmpG {ECO:0000313\|EMBL:OYZ86620.1}; | 314..320 |
| tr:A0A1V2PLD2_9ACTN | [A0A1V2PLD2] RecName: Full=Multidrug efflux pump Tap {ECO:0000256\|ARBA:ARBA00040914}; | 50..56 |
| tr:A0A850DH30_9MYCO | [A0A850DH30] SubName: Full=Carboxymethylenebutenolidase {ECO:0000313\|EMBL:NUS43224.1}; | 63..69 |
| tr:A0AAJ5VTZ8_9HYPH | [A0AAJ5VTZ8] RecName: Full=Protein MgtC {ECO:0000256\|RuleBase:RU365041}; | 145..151 |
| tr:A0AAJ6B0X4_9HYPH | [A0AAJ6B0X4] RecName: Full=Protein MgtC {ECO:0000256\|RuleBase:RU365041}; | 149..155 |
| tr:A0A4Y9F9F9_9DEIN | [A0A4Y9F9F9] SubName: Full=Dipeptidyl aminopeptidase {ECO:0000313\|EMBL:TFU25797.1}; | 14..20 |
| tr:A0A2J0YW67_RHIML | [A0A2J0YW67] SubName: Full=MFS transporter {ECO:0000313\|EMBL:PJR12162.1}; | 169..175 |
| tr:A0A538PVA2_UNCDE | [A0A538PVA2] SubName: Full=Uncharacterized protein {ECO:0000313\|EMBL:TMQ07547.1}; | 298..304 |
| tr:A0A538PKF6_UNCDE | [A0A538PKF6] SubName: Full=Uncharacterized protein {ECO:0000313\|EMBL:TMQ04096.1}; | 313..319 |
| tr:A0A845HWZ6_9BURK | [A0A845HWZ6] SubName: Full=Sensor histidine kinase {ECO:0000313\|EMBL:MYN44051.1}; | 64..70 |
| tr:A0A1J5QLJ1_9ZZZZ | [A0A1J5QLJ1] SubName: Full=Putative acetyltransferase {ECO:0000313\|EMBL:OIQ84186.1}; | 104..110 |
| tr:A0A3Q1EVW6_9TELE | [A0A3Q1EVW6] RecName: Full=E1 ubiquitin-activating enzyme {ECO:0000256\|ARBA:ARBA00012990}; EC=6.2.1.45 {ECO:0000256\|ARBA:ARBA00012990}; AltName: Full=Ubiquitin-activating enzyme E1 {ECO:0000256\|ARBA:ARBA00030371}; | 496..502 |
| tr:A0A7J5Z7A1_DISMA | [A0A7J5Z7A1] RecName: Full=SUMO-activating enzyme subunit 1 {ECO:0000256\|ARBA:ARBA00044187}; EC=6.2.1.45 {ECO:0000256\|ARBA:ARBA00012990}; AltName: Full=Ubiquitin-activating enzyme E1 {ECO:0000256\|ARBA:ARBA00030371}; AltName: Full=Ubiquitin-like 1-activating enzyme E1A {ECO:0000256\|ARBA:ARBA00044354}; | 551..557 |
| tr:A0A484DEK6_PERFV | [A0A484DEK6] RecName: Full=Ubiquitin-activating enzyme E1 C-terminal domain-containing protein {ECO:0000259\|SMART:SM00985}; Flags: Fragment; | 605..611 |
| tr:A0AAN8E6P3_CHAGU | [A0AAN8E6P3] SubName: Full=Uncharacterized protein {ECO:0000313\|EMBL:KAK5934097.1}; | 562..568 |

| | | |
|---|---|---|
| tr:A0A5J5DJA7_9PERO | [A0A5J5DJA7] RecName: Full=Ubiquitin-activating enzyme E1 C-terminal domain-containing protein {ECO:0008006\|Google:ProtNLM}; Flags: Fragment; | 562..568 |
| tr:A0A3P8Y8S7_ESOLU | [A0A3P8Y8S7] SubName: Full=T-complex-associated-testis-expressed 1 {ECO:0000313\|Ensembl:ENSELUP00000013007.1}; | 41..47 |
| tr:A0A8C3H6Z3_CHRPI | [A0A8C3H6Z3] RecName: Full=Phosphodiesterase {ECO:0000256\|RuleBase:RU363067}; EC=3.1.4.- {ECO:0000256\|RuleBase:RU363067}; | 177..183 |
| tr:A0A8C3FA28_CHRPI | [A0A8C3FA28] RecName: Full=Phosphodiesterase {ECO:0000256\|RuleBase:RU363067}; EC=3.1.4.- {ECO:0000256\|RuleBase:RU363067}; | 103..109 |
| tr:A0A8C3H6J3_CHRPI | [A0A8C3H6J3] RecName: Full=Phosphodiesterase {ECO:0000256\|RuleBase:RU363067}; EC=3.1.4.- {ECO:0000256\|RuleBase:RU363067}; | 157..163 |
| tr:A0A8C3F8K2_CHRPI | [A0A8C3F8K2] RecName: Full=Phosphodiesterase {ECO:0000256\|RuleBase:RU363067}; EC=3.1.4.- {ECO:0000256\|RuleBase:RU363067}; | 179..185 |
| tr:A0A8C3F818_CHRPI | [A0A8C3F818] RecName: Full=Phosphodiesterase {ECO:0000256\|RuleBase:RU363067}; EC=3.1.4.- {ECO:0000256\|RuleBase:RU363067}; | 177..183 |
| tr:A0A8C3F8C6_CHRPI | [A0A8C3F8C6] RecName: Full=Phosphodiesterase {ECO:0000256\|RuleBase:RU363067}; EC=3.1.4.- {ECO:0000256\|RuleBase:RU363067}; | 179..185 |
| tr:A0A8C3H6Z4_CHRPI | [A0A8C3H6Z4] RecName: Full=Phosphodiesterase {ECO:0000256\|RuleBase:RU363067}; EC=3.1.4.- {ECO:0000256\|RuleBase:RU363067}; | 103..109 |
| tr:A0A8C8S5F3_9SAUR | [A0A8C8S5F3] SubName: Full=T-complex-associated-testis-expressed 1 {ECO:0000313\|Ensembl:ENSPCEP00000014461.1}; | 42..48 |
| tr:A0A8C3F9U9_CHRPI | [A0A8C3F9U9] RecName: Full=Phosphodiesterase {ECO:0000256\|RuleBase:RU363067}; EC=3.1.4.- {ECO:0000256\|RuleBase:RU363067}; | 81..87 |
| tr:A0AAN8D4M3_9TELE | [A0AAN8D4M3] SubName: Full=Uncharacterized protein {ECO:0000313\|EMBL:KAK5912598.1}; | 562..568 |
| tr:A0A3Q1BYD9_AMPOC | [A0A3Q1BYD9] RecName: Full=E1 ubiquitin-activating enzyme {ECO:0000256\|ARBA:ARBA00012990}; EC=6.2.1.45 {ECO:0000256\|ARBA:ARBA00012990}; AltName: Full=Ubiquitin-activating enzyme E1 {ECO:0000256\|ARBA:ARBA00030371}; | 562..568 |
| tr:A0A672I4K8_SALFA | [A0A672I4K8] RecName: Full=E1 ubiquitin-activating enzyme {ECO:0000256\|ARBA:ARBA00012990}; EC=6.2.1.45 {ECO:0000256\|ARBA:ARBA00012990}; AltName: Full=Ubiquitin-activating enzyme E1 {ECO:0000256\|ARBA:ARBA00030371}; | 602..608 |
| tr:A0A672I9D2_SALFA | [A0A672I9D2] RecName: Full=E1 ubiquitin-activating enzyme {ECO:0000256\|ARBA:ARBA00012990}; EC=6.2.1.45 {ECO:0000256\|ARBA:ARBA00012990}; AltName: Full=Ubiquitin-activating enzyme E1 {ECO:0000256\|ARBA:ARBA00030371}; | 597..603 |
| tr:A0A672I373_SALFA | [A0A672I373] RecName: Full=E1 ubiquitin-activating enzyme {ECO:0000256\|ARBA:ARBA00012990}; EC=6.2.1.45 {ECO:0000256\|ARBA:ARBA00012990}; AltName: Full=Ubiquitin-activating enzyme E1 {ECO:0000256\|ARBA:ARBA00030371}; | 602..608 |
| tr:A0A672I8V9_SALFA | [A0A672I8V9] RecName: Full=E1 ubiquitin-activating enzyme {ECO:0000256\|ARBA:ARBA00012990}; EC=6.2.1.45 {ECO:0000256\|ARBA:ARBA00012990}; AltName: Full=Ubiquitin-activating enzyme E1 {ECO:0000256\|ARBA:ARBA00030371}; | 596..602 |
| tr:A0A672I7E0_SALFA | [A0A672I7E0] RecName: Full=E1 ubiquitin-activating enzyme {ECO:0000256\|ARBA:ARBA00012990}; EC=6.2.1.45 {ECO:0000256\|ARBA:ARBA00012990}; AltName: Full=Ubiquitin-activating enzyme E1 {ECO:0000256\|ARBA:ARBA00030371}; | 652..658 |
| tr:A0A672I950_SALFA | [A0A672I950] RecName: Full=E1 ubiquitin-activating enzyme {ECO:0000256\|ARBA:ARBA00012990}; EC=6.2.1.45 {ECO:0000256\|ARBA:ARBA00012990}; AltName: Full=Ubiquitin-activating enzyme E1 {ECO:0000256\|ARBA:ARBA00030371}; | 528..534 |
| tr:A0A672IA53_SALFA | [A0A672IA53] RecName: Full=E1 ubiquitin-activating enzyme {ECO:0000256\|ARBA:ARBA00012990}; EC=6.2.1.45 {ECO:0000256\|ARBA:ARBA00012990}; AltName: Full=Ubiquitin-activating enzyme E1 {ECO:0000256\|ARBA:ARBA00030371}; | 562..568 |
| tr:A0A672I919_SALFA | [A0A672I919] RecName: Full=E1 ubiquitin-activating enzyme {ECO:0000256\|ARBA:ARBA00012990}; EC=6.2.1.45 {ECO:0000256\|ARBA:ARBA00012990}; AltName: Full=Ubiquitin-activating enzyme E1 {ECO:0000256\|ARBA:ARBA00030371}; | 571..577 |
| tr:A0A672I7H1_SALFA | [A0A672I7H1] RecName: Full=E1 ubiquitin-activating enzyme {ECO:0000256\|ARBA:ARBA00012990}; EC=6.2.1.45 {ECO:0000256\|ARBA:ARBA00012990}; AltName: Full=Ubiquitin-activating enzyme E1 {ECO:0000256\|ARBA:ARBA00030371}; | 614..620 |
| tr:A0A8C0IN79_CHEAB | [A0A8C0IN79] RecName: Full=Phosphodiesterase {ECO:0000256\|RuleBase:RU363067}; EC=3.1.4.- {ECO:0000256\|RuleBase:RU363067}; | 166..172 |
| tr:A0A8C3XLK4_CHESE | [A0A8C3XLK4] RecName: Full=Phosphodiesterase {ECO:0000256\|RuleBase:RU363067}; EC=3.1.4.- {ECO:0000256\|RuleBase:RU363067}; | 194..200 |
| tr:A0A8C0IN77_CHEAB | [A0A8C0IN77] RecName: Full=Phosphodiesterase {ECO:0000256\|RuleBase:RU363067}; EC=3.1.4.- {ECO:0000256\|RuleBase:RU363067}; | 98..104 |
| tr:A0A8C0GJB6_CHEAB | [A0A8C0GJB6] RecName: Full=Phosphodiesterase {ECO:0000256\|RuleBase:RU363067}; EC=3.1.4.- {ECO:0000256\|RuleBase:RU363067}; | 98..104 |
| tr:A0A8C0INW5_CHEAB | [A0A8C0INW5] RecName: Full=Phosphodiesterase {ECO:0000256\|RuleBase:RU363067}; EC=3.1.4.- {ECO:0000256\|RuleBase:RU363067}; | 87..93 |
| tr:A0A3Q1ETW4_9TELE | [A0A3Q1ETW4] RecName: Full=E1 ubiquitin-activating enzyme {ECO:0000256\|ARBA:ARBA00012990}; EC=6.2.1.45 {ECO:0000256\|ARBA:ARBA00012990}; AltName: Full=Ubiquitin-activating enzyme E1 {ECO:0000256\|ARBA:ARBA00030371}; | 572..578 |
| tr:A0A3Q1GBE2_9TELE | [A0A3Q1GBE2] RecName: Full=E1 ubiquitin-activating enzyme {ECO:0000256\|ARBA:ARBA00012990}; EC=6.2.1.45 {ECO:0000256\|ARBA:ARBA00012990}; AltName: Full=Ubiquitin-activating enzyme E1 {ECO:0000256\|ARBA:ARBA00030371}; | 596..602 |
| tr:A0A3Q1EUD5_9TELE | [A0A3Q1EUD5] RecName: Full=E1 ubiquitin-activating enzyme {ECO:0000256\|ARBA:ARBA00012990}; EC=6.2.1.45 {ECO:0000256\|ARBA:ARBA00012990}; AltName: Full=Ubiquitin-activating enzyme E1 {ECO:0000256\|ARBA:ARBA00030371}; | 596..602 |
| tr:A0AAD9CG94_DISEL | [A0AAD9CG94] RecName: Full=E1 ubiquitin-activating enzyme {ECO:0000256\|ARBA:ARBA00012990}; EC=6.2.1.45 {ECO:0000256\|ARBA:ARBA00012990}; AltName: Full=Ubiquitin-activating enzyme E1 {ECO:0000256\|ARBA:ARBA00030371}; | 553..559 |
| tr:A0AAD6ALL4_9TELE | [A0AAD6ALL4] RecName: Full=E1 ubiquitin-activating enzyme {ECO:0000256\|ARBA:ARBA00012990}; EC=6.2.1.45 {ECO:0000256\|ARBA:ARBA00012990}; AltName: Full=Ubiquitin-activating enzyme E1 {ECO:0000256\|ARBA:ARBA00030371}; | 562..568 |

| | | |
|---|---|---|
| tr:A0A3P8SVT2_AMPPE | [A0A3P8SVT2] RecName: Full=SUMO-activating enzyme subunit 1 {ECO:0000256\|ARBA:ARBA00044187}; AltName: Full=Ubiquitin-like 1-activating enzyme E1A {ECO:0000256\|ARBA:ARBA00044354}; | 561..567 |
| tr:A0A8C9ZLD1_SANLU | [A0A8C9ZLD1] SubName: Full=Ubiquitin-like modifier activating enzyme 7 {ECO:0000313\|Ensembl:ENSSLUP00000042929.1}; | 557..563 |
| tr:A0A8D0D9F9_SANLU | [A0A8D0D9F9] SubName: Full=Ubiquitin-like modifier activating enzyme 7 {ECO:0000313\|Ensembl:ENSSLUP00000042933.1}; | 557..563 |
| tr:A0A8C9ZN52_SANLU | [A0A8C9ZN52] SubName: Full=Ubiquitin-like modifier activating enzyme 7 {ECO:0000313\|Ensembl:ENSSLUP00000042919.1}; | 559..565 |
| tr:A0A8C9ZSX2_SANLU | [A0A8C9ZSX2] SubName: Full=Ubiquitin-like modifier activating enzyme 7 {ECO:0000313\|Ensembl:ENSSLUP00000042925.1}; | 557..563 |
| tr:A0A8C9ZPE4_SANLU | [A0A8C9ZPE4] RecName: Full=SUMO-activating enzyme subunit 1 {ECO:0000256\|ARBA:ARBA00044187}; AltName: Full=Ubiquitin-like 1-activating enzyme E1A {ECO:0000256\|ARBA:ARBA00044354}; | 565..571 |
| tr:A0A8C9ZQD1_SANLU | [A0A8C9ZQD1] SubName: Full=Ubiquitin-like modifier activating enzyme 7 {ECO:0000313\|Ensembl:ENSSLUP00000042940.1}; | 394..400 |
| tr:A0A8C9ZPQ8_SANLU | [A0A8C9ZPQ8] RecName: Full=SUMO-activating enzyme subunit 1 {ECO:0000256\|ARBA:ARBA00044187}; AltName: Full=Ubiquitin-like 1-activating enzyme E1A {ECO:0000256\|ARBA:ARBA00044354}; | 555..561 |
| gpu:CP085224_4095 | [CP085224] hypothetical protein [Caenispirillum salinarum] | 206..212 |
| gp:CP108344_2752 | [CP108344] LysE family translocator [Streptomyces gancidicus] | 44..50 |
| gp:CP003926_1593 | [CP003926] hypothetical protein [Gluconobacter oxydans H24] | 186..192 |
| gp:CP013732_421 | [CP013732] hypothetical protein [Burkholderia cepacia JBK9] | 385..391 |
| gp:CP021793_1302 | [CP021793] MFS transporter [Sinorhizobium meliloti] | 169..175 |
| gp:CP021797_3330 | [CP021797] MFS transporter [Sinorhizobium meliloti] | 122..128 |
| gp:CP021800_786 | [CP021800] MFS transporter [Sinorhizobium meliloti] | 169..175 |
| gp:CP021808_479 | [CP021808] MFS transporter [Sinorhizobium meliloti] | 169..175 |
| gp:CP022564_778 | [CP022564] MFS transporter [Rhizobium leguminosarum bv. viciae] | 195..201 |
| gp:CP032010_160 | [CP032010] hypothetical protein [Burkholderia cepacia] | 349..355 |
| gp:CP070702_1854 | [CP070702] glycosyltransferase family 4 protein [Candidatus Eisenbacteria bacterium] | 109..115 |
| gp:CP070767_863 | [CP070767] ribosome biogenesis/translation initiation ATPase RLI [Euryarchaeota archaeon] | 440..446 |
| gp:CP089641_295 | [CP089641] Hypothetical protein [Nakaseomyces glabratus] | 49..55 |
| gp:CP126201_420 | [CP126201] hypothetical protein [Tetradesmus obliquus] | 237..243 |
