## Supplementary Table S4 for "A novel *Candida glabrata* protein regulated by mating signalling pathway shapes inter-species interaction"

*C. albicans, C. glabrata* and *S. cerevisiae* background recipient strains used in this study. Strain HTL was used (Schwarzmüller et al., 2014) as recipient strain for the generation of the deletion mutants in the strain library described in **Supplementary Table S1 and S2.**

| Strain Name | Strain Description | Genotype | Reference | Link |
|---|---|---|---|---|
| *C. albicans* SC5314 | Wild type strain | Wild-type strain | | NCBI Link |
| *C. glabrata* CBS138 | Wild type strain | **MATα** wild-type strain | Dujon et al., 2004 | Article Link |
| *C. glabrata* CBS138 HTL | Wild type strain (with auxotrophies) | **MATα** *his3Δ::FRT leu2Δ::FRT trp1Δ::FRT* | Schwarzmüller et al., 2014 | Article Link |
| *C. glabrata* BG2 | Wild type strain | **MATa** wild-type strain | Cormack & Falkow, 1999 | Article Link |
| *S. cerevisiae* S288c | Wild type strain (parent strain) | **MATα** SUC2 gal2 mal2 mel flo1 flo8-1 hap1 ho bio1 bio6 | Mortimer & Johnston, 1986 | Article Link |
| *S. cerevisiae* BY4741 | Wild type strain (specified mating type) | **MATa** *his3Δ1 leu2Δ0 met15Δ0 ura3Δ0* | Brachmann et al., 1998 | Article Link |
| *S. cerevisiae* BY4742 | Wild type strain (specified mating type) | **MATα** *his3Δ1 leu2Δ0 lys2Δ0 ura3Δ0* | Brachmann et al., 1998 | Article Link |
| *C. tropicalis* MYA-3404 | Wild type strain | Wild type strain | | NCBI Link |
| *C. dubliniensis* CD36 | Wild type strain | Wild type strain | | NCBI Link |
| *C. parapsilosis* CDC317 | Wild type strain | Wild type strain | | NCBI Link |
| *C. parapsilosis* CLIB 214 | Wild type strain | Wild type strain | | ATCC Link |
| *E. coli* DH5α | *E. coli* strain for cloning and plasmid maintenance | F– Δ(argF-lac)169, φ80dlacZ58(M15), ΔphoA8, glnX44(AS), λ-, deoR481, rfbC1, gyrA96(NalR), recA1, endA1, thiE1, hsdR17 | Hanahan, 1983<br>Raleigh et al., 1988 | Article Link<br>Article Link |
| *E. coli* BL21 (DE3) pLysS | *E. coli* strain for expression of recombinant protein | F– ompT, hsdSB (rB–, mB–), gal, dcm, λ(DE3), pLysS, CamR | Studier & Moffatt, 1986<br>Moffatt & Studier, 1987 | Article Link<br>Article Link |
| RPL-YS-54 | *S. cerevisiae* BY4741+*CgYHI1* | **MATa** *his3Δ1 leu2Δ0 met15Δ0 ura3Δ0* pRPL033 | This study | |
| RPL-YS-27 | *S. cerevisiae* BY4742+*CgYHI1* | **MATα** *his3Δ1 leu2Δ0 lys2Δ0 ura3Δ0* pRPL033 | This study | |
| RPL-YS-81 | *C. glabrata mfa2Δ* | **MATα** *his3Δ::FRT leu2Δ::FRT trp1Δ::FRT mfa2::LEU2* | This study | |
| RPL-YS-35 | *C. glabrata yhi1Δ+CgYHI1*$^{1-66}$ | **MATα** *his3Δ::FRT leu2Δ::FRT trp1Δ::FRT* pRPL035 | This study | |
| RPL-YS-30 | *C. glabrata yhi1Δ+CgYHI1*$^{3-66}$ | **MATα** *his3Δ::FRT leu2Δ::FRT trp1Δ::FRT* pRPL034 | This study | |
| RPL-YS-74 | *C. glabrata yhi1Δ+CgYHI1*$^{1-32}$ | **MATα** *his3Δ::FRT leu2Δ::FRT trp1Δ::FRT* pRPL100 | This study | |
| RPL-YS-75 | *C. glabrata yhi1Δ+CgYHI1*$^{33-66}$ | **MATα** *his3Δ::FRT leu2Δ::FRT trp1Δ::FRT* pRPL101 | This study | |
| RPL-YS-87 | *C. glabrata yhi1Δ+CgYHI1*$^{1-18}$ | **MATα** *his3Δ::FRT leu2Δ::FRT trp1Δ::FRT* pRPL123 | This study | |
| RPL-YS-88 | *C. glabrata yhi1Δ+CgYHI1*$^{21-52}$ | **MATα** *his3Δ::FRT leu2Δ::FRT trp1Δ::FRT* pRPL124 | This study | |
| RPL-YS-98 | *C. glabrata yhi1Δ+CgYHI1*$^{1-32(A18G,\ H22A)}$ | **MATα** *his3Δ::FRT leu2Δ::FRT trp1Δ::FRT* pRPL126 | This study | |
| RPL-YS-89 | *C. glabrata yhi1Δ*+SP-yeGFP-AxVxH | **MATα** *his3Δ::FRT leu2Δ::FRT trp1Δ::FRT* pRPL127 | This study | |
| RPL-YS-90 | *C. glabrata yhi1Δ*+SP-yeGFP | **MATα** *his3Δ::FRT leu2Δ::FRT trp1Δ::FRT* pRPL128 | This study | |
| RPL-ECS-04 | *E. coli* BL21 (DE3) pLysS + *CgYHI1* | F– ompT, hsdSB (rB–, mB–), gal, dcm, λ(DE3), pLysS, CamR, pRPL130 | This study | |

**Supplementary Table S4.**

Plasmids used or generated in this study.

| Plasmid | Description | Reference | Link |
|---|---|---|---|
| p426-GPD | Shuttle expression vector, containing a GPD promoter flanked by cloning array and the CYC1 terminator. Constructed in the 2µ plasmid background of pRS series [Sikorski & Hieter, 1989; Christianson et al., 1992] containing a URA3 selectable marker. | Mumberg, 1995 | Link |
| pBEVY-L | Bi-directional expression vector for yeast, constitutively active bi-directional promoter consisting a fusion of the GPD promoter and a short derivative of the ADH1 promoter, LEU2 selectable marker. | Miller III, 1998 | Link |
| pET32b(+) | Plasmid for cloning and expression of proteins/peptides fused with a thioredoxin protein tag, containing cleavable His•Tag and S•Tag sequences for detection and purification of fusion protein. | LaVallie et al., 1993 | Link |
| pKT0209 | pFA6a–link–yEGFP–CaURA3. A codon-optimized linker added into the yEGFP to improve expression level. *C. albicans* URA3 (*CaURA3*) introduced by subcloning the *BglII/SacI* fragment of pAG60 (Goldstein et al., 1999) into the plasmid, in place of SpHIS5. | Sheff & Thorn, 2004 | Link |
| pRPL033 | *CgYHI1*$^{1-66}$ cloned in p426-GPD between *BamHI/EcoRI* for expression of WT small protein (Yhi1) under GPD promoter, for expression in yeast | This study | |
| pRPL034 | *CgYHI1*$^{3-66}$ cloned in pBEVY-L between *BamHI/SalI* for expression of truncated version (Yhi1$^{3-66}$) of the small protein Yhi1 under GPD promoter, for expression in yeast | This study | |
| pRPL035 | *CgYHI1*$^{1-66}$ cloned in pBEVY-L between *BamHI/SalI* for expression of the small protein Yhi1 under GPD promoter, for expression in yeast | This study | |
| pRPL092 | CgMFA2-5'-UTR (546 bp) in pRPL091 at *BamHI/XbaI* site. CgMFA2-3'-UTR (572 bp) in pBEVY-L between *KpnI/SacI*. Final plasmid for deletion of *C. glabrata* mating factor **a** *CgMFA2* (*CAGL0C01919g*) in *C. glabrata* HTL strain. | This study | |
| pRPL100 | *CgYHI1*$^{1-32}$ (truncated N-terminal part of small protein Yhi1) cloned in pBEVY-L between *BamHI/SalI* for expression of truncated version of small protein (Yhi1$^{1-32}$) under GPD promoter, for expression in yeast | This study | |
| pRPL101 | *CgYHI1*$^{33-66}$ (truncated C-terminal part of small protein Yhi1) cloned in pBEVY-L between *BamHI/SalI* for expression of truncated version of small protein under GPD promoter, for expression in yeast | This study | |
| pRPL124 | *CgYHI1*$^{21-52}$ (truncated/disrupted sequence between the two AxVxH motifs of Yhi1) cloned in pBEVY-L between *BamHI/SalI* for expression of truncated version (Yhi1$^{21-52}$) of the small protein Yhi1 under GPD promoter, for expression in yeast | This study | |
| pRPL126 | *CgYHI1*$^{1-32(A18G,\ H22A)}$ (truncated part of Yhi1 with mutated ADVWH motif) cloned in pBEVY-L between *BamHI/SalI* for expression of truncated version (Yhi1$^{1-32(A18G,\ H22A)}$) of the small protein Yhi1 under GPD promoter, for expression in yeast | This study | |
| pRPL127 | yeGFP (pKT209) cloned in-frame with a N-terminal signal peptide and a pentapeptide motif (ADVWH motif of *Cg* Yhi1) after the 5th aa residue of yeGFP (accessible loop region), in pBEVY-L between *BamHI/SalI* for expression of **SP-yeGFP-ADVWH** fusion protein under GPD promoter, for expression in yeast. | This study | |
| pRPL128 | yeGFP (pKT209) cloned with a N-terminal signal peptide, in pBEVY-L between *BamHI/SalI* for expression of **SP-yeGFP** fusion protein under GPD promoter, for expression in yeast. | This study | |
| pRPL130 | *CgYHI1*$^{1-66}$ cloned in frame with TrxA+6xHis+Thrombin Cleavage Site between *XbaI/BamHI* in pET32b+, removing the enterokinase site of pET32b+, as well as relocating the thrombin cleavage site to reduce the number of linker residues. | This study | |

**Supplementary Table S4.**

List of oligonucleotide primers used in this study

| Primer | Sequence (5' - 3') | Comments |
|---|---|---|
| CgYHI1-F | TAAGCAggatccATGAAGATGTATAAGCTAGCCTTG | Forward primer for ORF ampification of *CgYHI1* , for cloning in p426GPD/pBEVY-L between *BamHI/EcoRI* or *BamHI/SalI* under the GPD promoter, for expression in yeast. |
| CgYHI1-R1 | TAAGCAgaattcTCAGAGCATGATACTTGGAGTAAAC | Reverse primer for ORF ampification of *CgYHI1* , for cloning in p426GPD between *BamHI/EcoRI* under the GPD promoter, for expression in yeast. |
| CgYHI1-R2 | TAAGCAgtcgacTCAGAGCATGATACTTGGAGTAAAC | Reverse primer for ORF ampification of *CgYHI1* , for cloning in pBEVY-L between *BamHI/SalI* under the GPD promoter, for expression in yeast. |
| Yhi1$^{3-66}$-F | TAAGCAggatccATGTATAAGCTAGCCTTGATAATAAAG | Forward primer for ampification of *CgYHI1* without the first two amino acid residues, for cloning in pBEVY-L between *BamHI/SalI* under the GPD promoter, for expression in yeast. |
| CgYHI1-qPCR-F | CAAAGCCTCTTTATTCATTTC | Forward primer for ORF amplification of *CgYHI1* - For qPCR |
| CgYHI1-qPCR-R | GAAAACTTGTTATGACACTATC | Reverse primer for ORF amplification of *CgYHI1* - For qPCR |
| CgARP2-F | GGATCCACACAACCCAATTG | Forward primer for limited ORF amplification of *CgARP2* - For detection of genomic DNA contamination and qPCR |
| CgARP2-R | CCTCGTCACCAATCATTATG | Reverse primer for limited ORF amplification of *CgARP2* - For detection of genomic DNA contamination and qPCR |
| CgSTE6-qPCR-F | CAAAGCCTCTTTATTCATTTC | Forward primer for limited ORF amplification of *CgSTE6* - For qPCR |
| CgSTE6-qPCR-R | GAAAACTTGTTATGACACTATC | Reverse primer for limited ORF amplification of *CgSTE6* - For qPCR |
| CgMFA2-F | ATGCAACCAACTATTGAAGCC | Forward primer for ORF ampification of *CgMFA2* |
| CgMFA2-R | CTAAGCGATTACACAATCTGG | Reverse primer for ORF ampification of CgMFA2 |
| CgMFA2-5'UTR-F | TAAGCAggatccCACTTGAGGTTGTCAGAGATAG | Forward primer for cloning of 5' flank of *CgMFA2* in pBEVY-L between *BamHI/XbaI* , for deletion of *CgMFA2* in *C. glabrata* . |
| CgMFA2-5'UTR-R | TAAGCAtctagaTTGTTATGTGATCACCATTTGAAAATG | Reverse primer for cloning of 5' flank of *CgMFA2* in pBEVY-L between *BamHI/XbaI* , for deletion of *CgMFA2* in *C. glabrata* . |
| CgMFA2-3'UTR-F | TAAGCAgagctcGAAATACGCTGACGATAATTATCTC | Forward primer for cloning of 3' flank of *CgMFA2* in pBEVY-L between *SacI/KpnI* , for deletion of *CgMFA2* in *C. glabrata* . |
| CgMFA2-3'UTR-R | TAAGCAggtaccGGATTTAAGAATCAGAGAAATAAATGG | Forward primer for cloning of 3' flank of *CgMFA2* in pBEVY-L between *SacI/KpnI* , for deletion of *CgMFA2* in *C. glabrata.* |
| CgMFA2-LS-F | CAGTACATGCTCAAGACAGTTG | Forward primer for confirmation of locus-specific integration of deletion casette for deletion of *CgMFA2* in *C. glabrata.* |
| CgMFA2-LS-R | ATGCAAGCTTTGGACTTCTTCG | Reverse primer for confirmation of locus-specific integration of deletion casette for deletion of *CgMFA2* in *C. glabrata.* |
| Yhi1$^{1-32}$-F | TAAGCAggatccATGAAGATGTATAAGCTAGCC | Forward primer for cloning of Yhi1$^{1-32}$ in pBEVY-L between BamHI/SalI under the GPD promoter, for expression in yeast. |
| Yhi1$^{1-32}$-R | TAAGCAgtcgacTCAGCATTTCACTTTGTAGGCATG | Reverse primer for cloning of Yhi1$^{1-32}$ in pBEVY-L between *BamHI/SalI* under the GPD promoter, for expression in yeast. |
| Yhi1$^{33-66}$-F | TAAGCAggatccATGACACCGTTCAGAGGTTTCTTG | Forward primer for cloning of Yhi1$^{33-66}$ in pBEVY-L between *BamHI/SalI* under the GPD promoter, for expression in yeast. |
| Yhi1$^{33-66}$-R | TAAGCAgtcgacTCAGAGCATGATACTTGGAGTAAAC | Reverse primer for cloning of Yhi1$^{33-66}$ in pBEVY-L between *BamHI/SalI* under the GPD promoter, for expression in yeast. |
| Yhi1$^{21-52}$-F | TAAGCAggatccATGTGGCATGACAAATTACATGCCTAC | Forward primer for amplification of sequence between the two pentapeptide A*V*H motifs from Yhi1 sequence leading to disruption of both motifs, with a Met (ATG) added, for cloning between *BamHI/SalI* in pBEVY-L, under GPD Promoter, for expression in yeast. |
| Yhi1$^{21-52}$-R | TAAGCAgtcgacTCAAACTGCCAACGGAGGCCTCTT | Reverse primer for amplification of sequence between the two pentapeptide A*V*H motifs from Yhi1 sequence leading to disruption of both motifs, with an opal stop codon (TGA) added in the reverse primer, for cloning between *BamHI/SalI* in pBEVY-L, under GPD Promoter, for expression in yeast. |
| SP-yeGFP-ADVWH-F | TAGCggatccATGGTTTCTTTCTCTTTTGCTACTATTGTAGCAGCGCTTTCTGTAAGTGCGTTGGCTAAGATGTCTAAAGGTGAAGCTGATGTTTGGCATGAATTATTC | Forward primer for amplification of yeGFP from pKT209, containing a Signal peptide from Cg484 at the N-terminus (20 aa) and a pentapeptide motif (ADVWH) after the 5th aa residue, for cloning between *BamHI/SalI* in pBEVY-L, for expression in yeast. |
| SP-yeGFP-ADVWH-R | TAAGCAgtcgacTTATTTGTACAATTCATCCATACCATGGGTAATACCAG | Reverse primer for amplification of yeGFP from pKT209, containing a Signal peptide from Cg484 at the N-terminus (20 aa) and a pentapeptide motif (ADVWH) after the 5th aa residue, for cloning between *BamHI/SalI* in pBEVY-L, for expression in yeast. |
| SP-yeGFP-F | TAGCggatccATGGTTTCTTTCTCTTTTGCTACTATTGTAGCAGCGCTTTCTGTAAGTGCGTTGGCTAAGATGTCTAAAGGTGAAGAATTATTCACTGGTGTG | Forward primer for amplification of yeGFP from pKT209, containing a Signal peptide from Cg484 at the N-terminus (20 aa), without a pentapeptide motif (ADVWH), for cloning between *BamHI/SalI* in pBEVY-L. |
| SP-yeGFP-R | TAAGCAgtcgacTTATTTGTACAATTCATCCATACCATGGGTAATACCAG | Reverse primer for amplification of yeGFP from pKT209, containing a Signal peptide from Cg484 at the N-terminus (20 aa), for cloning between *BamHI/SalI* in pBEVY-L, for expression in yeast. |
| Yhi1-pET-PCR1-F | TAAGCAtctagaAATAATTTTGTTTAACTTTAAGAAGGAGATATACAT | Forward Primer for amplification of TrxA + 6xHis + Thrombin site for PCR1, with *XbaI* site, in sequential PCR for amplification of "TrxA+6xHis+Thrombin+Yhi1" fusion casette, for cloning in pET32b+ between *XbaI/BamHI* , for bacterial expression. |
| Yhi1-pET-PCR1-R | CTTTATTATCAAGGCTAGCTTATACATCTTCATTTTCATACCAGAACCGCGTGGCACCAG | Reverse Primer for amplification of TrxA + 6xHis + Thrombin site for PCR1, in sequential PCR for amplification of "TrxA+6xHis+Thrombin+Yhi1" fusion casette, for cloning in pET32b+ between *XbaI/BamHI* , for bacterial expression. |
| Yhi1-pET-PCR2-F | CTGGTGCCACGCGGTTCTGGTATGAAAATGAAGATGTATAAGCTAGCCTTGATAATAAAG | Forward Primer for amplification of Yhi1 for PCR2, with overhang of Thrombin site+linker(3aa), in sequential PCR for amplification of "TrxA+6xHis+Thrombin+Yhi1" fusion casette, for cloning in pET32b+ between *XbaI/BamHI* , for bacterial expression. |
| Yhi1-pET-PCR2-R | TAAGCAggatccTCAGAGCATGATACTTGGAGTAAACTTTAATAAG | Reverse Primer for amplification of Yhi1 for PCR2, with *BamHI* site, in sequential PCR for amplification of "TrxA+6xHis+Thrombin+Yhi1" fusion casette, for cloning in pET32b+ between *XbaI/BamHI* , for bacterial expression. |
