## Supplementary Table S5 for "A novel *Candida glabrata* protein regulated by mating signalling pathway shapes inter-species interaction"

List of reagents used in this study.

| Technique | Reagent | Source | Catalog No. |
|---|---|---|---|
| Media Components | Yeast extract | HiMedia | RM027 |
| | Proteose Peptone | HiMedia | RM005 |
| | Glucose | HiMedia | GRM016 |
| | Agar agar | HiMedia | GRM666 |
| | Luria Broth | HiMedia | M575 |
| | Yeast Nitrogen Base, with ammonium sulphate, without amino acids | HiMedia | M878 |
| | L-Histidine HCl | Sigma-Aldrich | H6034 |
| | L-Leucine | Sigma-Aldrich | L8912 |
| | L-Lysine HCl | Sigma-Aldrich | L8662 |
| | L-Methionine | Sigma-Aldrich | M5308 |
| | L-Tryptophan | Sigma-Aldrich | T8941 |
| | Uracil | Sigma-Aldrich | U1128 |
| Buffer Components | Tris | Sisco Research Laboratories | 71033 |
| | Glycine | HiMedia | MB013 |
| | Sodium dodecyl sulphate | Sigma-Aldrich | L3771 |
| | Sodium dihydrogen phosphate, monobasic | Sisco Research Laboratories | 61707 |
| | Sodium chloride | Sisco Research Laboratories | 41721 |
| | Imidazole | Sisco Research Laboratories | 61510 |
| | Potassium chloride | Sisco Research Laboratories | 38630 |
| | Disodium hydrogen phosphate | Sisco Research Laboratories | 21669 |
| | Potassium dihydrogen phosphate | HiMedia | TC001 |
| | Hydrochloric acid | Supelco (Merck) | 1.93001 |
| | Methanol | Sigma-Aldrich | 179337 |
| | Tween 20 | HiMedia | MB067 |
| Filter sterilization | 0.22 µm PES syringe filter | Millipore (Merck) | SLGPR33RS |
| Protein Expression & Purification | Isopropyl β-1-thiogalactopyranoside (IPTG) | HiMedia | RM2578 |
| | Protease Inhibitor Cocktail | Sigma-Aldrich | P2714 |
| | Ni-NTA His•Bind Resin | Millipore (Merck) | 70666 |
| | Imidazole | Sigma-Aldrich | I5513 |
| | SnakeSkin™ Dialysis Tubing | Thermo Scientific | 88244 |

| | | | |
|---|---|---|---|
| Ultrafiltration | Amicon Ultra Centrifugal Filter, 3 kDa MWCO | Millipore (Merck) | UFC9003 |
| | Amicon Ultra Centrifugal Filter, 30 kDa MWCO | Millipore (Merck) | UFC9030 |

| | | | |
|---|---|---|---|
| Cloning | GoTaq Flexi DNA Polymerase | Promega Corporation | M8295 |
| | 5X Colorless GoTaq Flexi Reaction Buffer | Promega Corporation | M8901 |
| | Phusion High-Fidelity DNA Polymerase | Thermo Scientific | F350S |
| | dNTP Set (100 mM) | Thermo Scientific | 10297018 |
| | BamHI, FastDigest | Thermo Scientific | FD0054 |
| | EcoRI, FastDigest | Thermo Scientific | FD0274 |
| | KpnI, FastDigest | Thermo Scientific | FD0524 |
| | SacI, FastDigest | Thermo Scientific | FD1134 |
| | SalI, FastDigest | Thermo Scientific | FD0644 |
| | XbaI, FastDigest | Thermo Scientific | FD0685 |
| | T4 DNA Ligase | Thermo Scientific | EL0011 |

| | | | |
|---|---|---|---|
| RT-PCR | TRIzol Reagent | Invitrogen (Thermo Scientific) | 15596026 |
| | Protector RNase Inhibitor | Roche (Merck) | 3335399001 |
| | DNase I, RNase-free | Thermo Scientific | EN0521 |
| | iScript cDNA Synthesis Kit | BioRad | 1708891 |
| | SsoAdvanced Uuniversal SYBR Green Supermix | BioRad | 1725270 |

| | | | |
|---|---|---|---|
| Immunoblotting | Glass beads, acid-washed (425-600 µm) | Sigma-Aldrich | G8772-500G |
| | Protease Inhibitor Cocktail | Sigma-Aldrich | P8340 |
| | PVDF Western Blotting Membranes, pore size 0.2 µm | Roche (Merck) | 3010040001 |
| | Ponceau S | Sigma-Aldrich | P3504 |
| | Anti-Yhi1 Polyclonal Antibody, raised in rabbit | Biotech Desk Pvt Ltd | Custom Order |
| | Goat anti-Rabbit IgG (H+L) Cross-Adsorbed Secondary Antibody, HRP | Invitrogen (Thermo Scientific) | G21234 |
| | WESTAR ANTARES ECL substrate | Cyanagen | XLS142 |

| | | | |
|---|---|---|---|
| Markers | GeneRuler 1 kb DNA Ladder | Thermo Scientific | SM0311 |
| | BLUeye Prestained Protein Ladder | GeneDireX | PM007-0500 |

| | | | |
|---|---|---|---|
| Transwell Assay | Nunc 24-well Carrier Plate with Cell Culture Inserts (0.4 µm) | Thermo Scientific | 141002 |

| | | | |
|---|---|---|---|
| Hyphal Induction Assay | Nunc BioLite 96 Well Multidish, Cell culture treated, Flat bottom | Thermo Scientific | 130188 |

| | | | |
|---|---|---|---|
| Fluorescence Imaging | Confocal Dish, cell culture treated glass-bottom petridish | SPL Life Sciences | 200350 |

**Supplementary Table S5.**

List of buffers, media, and solutions used in this study, with their compositions.

| Buffer/Media/Solution | Chemical/Media Component | Concentration | Reference | Link |
|---|---|---|---|---|
| 1x Ni-NTA Binding Buffer (pH 8.0) | Sodium dihydrogen phosphate, monobasic | 50 mM | Ni-NTA His•Bind Resin (Merck) Handbook | Link |
| | Sodium chloride | 300 mM | | |
| | Imidazole | 10 mM | | |
| 1x Ni-NTA Wash Buffer (pH 8.0) | Sodium dihydrogen phosphate, monobasic | 50 mM | Ni-NTA His•Bind Resin (Merck) Handbook | Link |
| | Sodium chloride | 300 mM | | |
| | Imidazole | 20 mM | | |
| 1x Ni-NTA Elution Buffer (pH 8.0) | Sodium dihydrogen phosphate, monobasic | 50 mM | Ni-NTA His•Bind Resin (Merck) Handbook | Link |
| | Sodium chloride | 300 mM | | |
| | Imidazole | 250 mM | | |
| 1x SDS-PAGE Running Buffer (1 L) | Tris | 3.03 g/L | Cold Spring Harbor Protocols, 2015 | Link |
| | Glycine | 14.44 g/L | | |
| | SDS | 1 g/L | | |
| 1x Western Transfer Buffer (1 L) | Tris | 25 mM | Cold Spring Harbor Protocols, 2014 | Link |
| | Glycine | 192 mM | | |
| | Methanol | 20% (v/v) | | |
| 1x Tris-buffered Saline with Tween 20 (TBST) [pH 7.4] | Tris | 19 mM | Cold Spring Harbor Protocols, 2013 | Link |
| | Potassium chloride | 2.7 mM | | |
| | Sodium chloride | 137 mM | | |
| | Tween 20 | 0.1% (w/v) | | |
| Tris-Cl (pH 7) | Tris | 0.5 M | Modified from: Cold Spring Harbor Protocols, 2006 | Link |
| 1x PBS (pH 7.4) | Sodium chloride | 137 mM | Cold Spring Harbor Protocols, 2006 | Link |
| | Potassium chloride | 2.7 mM | | |
| | Disodium hydrogen phosphate | 10 mM | | |
| | Potassium dihydrogen phosphate | 1.8 mM | | |
| YPD | Yeast extract | 10 g/L | Cold Spring Harbor Protocols, 2017 | Link |
| | Proteose Peptone | 20 g/L | | |
| | Glucose | 20 g/L | | |
| Lysogeny Broth | Luria Broth | 20 g/L | HiMedia Technical Datasheet | Link |
| Glucose Minimal Medium (GMM) | Glucose | 20 g/L | Xu et al., 2008 | Link |
| | Yeast Nitrogen Base, with ammonium sulphate, without amino acids | 6.79 g/L | | |
| | Glucose | 20 g/L | | |

| | | | | |
|---|---|---|---|---|
| Synthetic Defined Medium (for *C. glabrata* CBS138 HTL) | Yeast Nitrogen Base, with ammonium sulphate, without amino acids | 6.79 g/L | Schwarzmüller et al., 2014 | Link |
| | L-Histidine HCl | 20 mg/L | | |
| | L-Tryptophan | 20 mg/L | | |
| | L-Leucine | 100 mg/L | | |
| Synthetic Defined Medium (for *S. cerevisiae* BY4741) | Glucose | 20 g/L | Cold Spring Harbor Protocols, 2015 | Link |
| | Yeast Nitrogen Base, with ammonium sulphate, without amino acids | 6.79 g/L | | |
| | L-Histidine HCl | 20 mg/L | | |
| | L-Leucine | 100 mg/L | | |
| | L-Methionine | 20 mg/L | | |
| | Uracil | 20 mg/L | | |
| Synthetic Defined Medium (for *S. cerevisiae* BY4742) | Glucose | 20 g/L | Cold Spring Harbor Protocols, 2015 | Link |
| | Yeast Nitrogen Base, with ammonium sulphate, without amino acids | 6.79 g/L | | |
| | L-Histidine HCl | 20 mg/L | | |
| | L-Leucine | 100 mg/L | | |
| | L-Lysine HCl | 30 mg/L | | |
| | Uracil | 20 mg/L | | |
| Tris-buffered Saline with Tween 20 (TBST) [pH 7.5] | Tris | 20 mM | Towbin et al., 1979 | Link |
| | Sodium chloride | 150 mM | | |
| | Tween 20 | 0.1% (w/v) | | |
| 1x TE Buffer [pH 8.0] | Tris-Cl | 10 mM | Cold Spring Harbor Protocols, 2009 | Link |
| | EDTA | 1 mM | | |
